# Alphavirus replicon particle expressing IL-12 reprograms tumor-associated macrophages and neutrophils and induces anti-tumor immunity

**DOI:** 10.1101/2025.08.07.666585

**Authors:** Momoko Ishikawa, Dhanya K Nambiar, Jaya Sastri, Forrest Bowling, Keiko Ishimoto, Fumiya Tao, Koyo Takahashi, Ivan Stephanek, Hongbin Cao, Davis Leitner, Kenta Matsuda, Fred M Baik, John B. Sunwoo, Quynh-Thu Le, Jonathan F Smith, Wataru Akahata

## Abstract

Cancers evade anti-tumor immunity by the infiltration of immunosuppressive myeloid cells such as macrophages and neutrophils into the tumor microenvironment (TME). The limited efficacy of current immunotherapy is attributed to the presence of these immunosuppressive cells. Here, we describe a novel immunotherapy approach, using a single-cycle alphavirus replicon particle (VRP) that carries a self-amplifying RNA encoding interleukin-12 (IL-12). Leveraging the natural tropism of alphavirus to myeloid cells, single cell transcriptomic analyses of mouse tumor tissues demonstrated that intratumoral (i.t.) injection of the vector effectively reprogrammed tumor-associated macrophages and neutrophils toward a pro-inflammatory, anti-tumor phenotype via interferon signaling, further enhanced by IL-12 expression. The treatment also induced cytotoxic NK and T cell responses, resulting in a significant reduction of tumor growth and effectively preventing nodal and distant metastases in solid tumor mouse models. By targeting tumor-supporting myeloid cells to transform the TME into an immune-active state, our myeloid modulation approach has the potential to enhance the efficacy of current immunotherapies.

## Introduction

Immune checkpoint therapies (ICT) have transformed cancer therapy by harnessing the host’s immune system to eliminate malignant cells. However, some tumors do not respond to ICT, and this limited efficacy can be attributed to multiple factors including the presence of immunosuppressive cells in the tumor microenvironment (TME)^1,2^. Many solid tumors develop TME that promotes tumor progression by suppressing the function of T cells^3,4^, natural killer (NK) cells^5–8^, macrophges^9,10^ and neutrophils^11,12^, thus allowing tumor cells to evade immune control.

Among the immune cell populations in the TME, myeloid cells such as tumor-associated macrophages (TAMs) are the most abundant in most tumor types and often serve as central mediators of immune suppression^13–17^. Chronic exposure to tumor antigens and inflammatory signals can polarize myeloid cells into immunosuppressive phenotypes^13–17^. This typically results in a shift to M2-like macrophages, which are characterized by the secretion of anti-inflammatory cytokines that inhibit T cell function^18^. While a balance between pro-inflammatory M1 and anti-inflammatory M2 phenotypes is critical for maintaining tissue homeostasis under physiological conditions, an M2-skewed state in the TME is associated with worse clinical outcomes, including poor response to therapy and lower recurrence-free survival^13–17^. Neutrophils are also a key component of the immune myeloid cell populations in the TME. Recent studies have shown that tumor-associated neutrophils (TANs), like macrophages, can both inhibit (N1 TANs) and promote (N2 TANs) tumor growth^19,20^. Most human tumors have a high rate of TANs infiltration, and they are associated with poor prognoses^21–23^. Therefore, modulating the plasticity of these TAMs and TANs phenotypes as dynamic participants in tumor immunity could improve treatment outcomes.

Interleukin-12 (IL-12) has re-emerged among immunomodulators as a potent therapeutic candidate, especially in reprogramming immunosuppressive myeloid cells. IL-12 is a heterodimeric cytokine composed of two disulfide-linked subunits: p35, encoded by the *Il12a* gene, and p40, encoded by the *Il12b* gene^24^. IL-12 promotes anti-tumor immunity by stimulating the proliferation and activation of cytotoxic T cells and NK cells, inducing interferon-gamma (IFN-γ) production to enhance Th1 responses, reducing Treg activity, reprogramming macrophages toward an M1-like phenotype, and suppressing angiogenesis^25^. Although active, systemic delivery of IL-12 is complicated by its toxicity^26–28^, thus recent studies have utilized intratumoral (i.t.) gene delivery such as plasmid IL-12 delivered with electroporation^29–31^ or RNA expressing IL-12 formulated with LNP^32–35^.

One promising delivery platform is the viral replicon particle (VRP) system that can be used to express IL-12 in the TME. VRP consists of self-amplifying RNA (saRNA) packaged in a virus-like particle, which enables high transgene expression via RNA replication and transcription within target cells^36^. VRP systems are derived from alphaviruses such as Venezuelan Equine Encephalitis Virus (VEEV) and Chikungunya virus, which are positive-sense, single-stranded RNA viruses that naturally infect myeloid cells and fibroblasts^37–39^. Thus, VRP systems are particularly attractive for this purpose because they offer a mechanism for targeted and efficient delivery of RNA cargo to immunosuppressive myeloid cells in the TME. This unique characteristic of cell tropism enables localized transgene expression in target cells.

Here we present a preclinical study that utilizes VRP to deliver IL-12, enabling robust and localized cytokine expression in the TME while leveraging the natural tropism of the vector to target tumor-associated macrophages and neutrophils to promote immune reprogramming. By shifting these cells toward a pro-inflammatory, anti-tumor phenotype, we demonstrate an alternative immunotherapeutic approach that acts through myeloid modulation.

## Results

### Alphavirus VRP with a Capsid Nuclear Localization Signal Mutation Efficiently Transduces Mammalian Macrophages

We first developed a VRP system based on the VEEV for gene delivery. The VRP constitutes a replication-incompetent, single-cycle vector system lacking the genes encoding the viral structural proteins. The production platform utilizes two plasmids for transient transfection of mammalian cells (**Fig. 1a**). The first plasmid includes the VEEV 5’ untranslated region (UTR), non-structural protein sequences (nsPs) 1–4, a subgenomic promoter (26S), a gene of interest, and the 3’ UTR. The nsPs form the RNA-dependent RNA polymerase (RdRp) complex, which allows for self-amplification of the replicon RNA. The second plasmid encodes the VEEV structural proteins, capsid and envelope (E3, E2, 6k and E1). Transfection of HEK293T cells with both plasmids resulted in the production of VRPs with a mean measured diameter of 76.8 ±6.3nm and were morphologically indistinguishable from native VEEV particles (**Fig. 1b**).

**Fig. 1.**
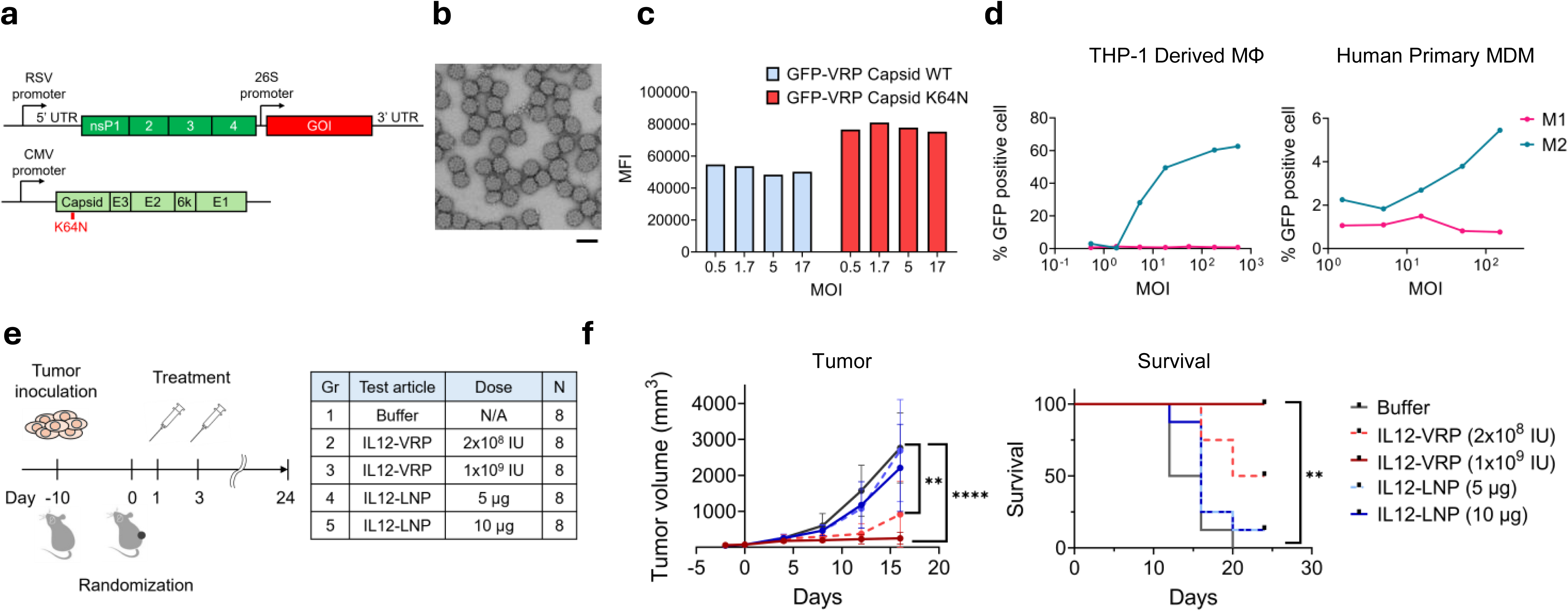
Virus replicon particle (VRP) with IL-12 payload transduces macrophages and inhibits mouse tumor growth. **a,** Replicon (top) and helper (bottom) plasmid maps. UTR: untranslated region; nsP: non-structure protein; GOI: gene of interest; E1-3: envelope 1-3. **b,** Transmission electron microscopy imaging of GFP-VRPs. Scale bar indicates 100nm. **c,** GFP expression levels in mouse macrophage cell line J774 transduced by GFP-VRP with or without capsid mutation. MFI: mean fluorescent intensity; IU: infectious unit; MOI: multiplicity of infection. **d,** Flowcytometry analysis of percent transduced M1-like or M2-like macrophages by GFP-VRP. Human THP-1 derived macrophages (left) and human CD14^+^ monocyte derived macrophages (right). **e,** Study design using MC38 mouse syngeneic tumor model to evaluate tumor growth inhibition by IL12-VRP or IL12-LNP. **f,** Tumor growth curve of each treatment group (left). Asterisks indicate statistical significance between Group 1 and Group 2, or Group 1 and Group 3 on Day 16. **, *p*<0.01; ****, *p<*0.0001 by *t*-test. Survival proportion of each treatment group (right) Asterisks indicate statistical significance between Group 1 and Group 3 on Day 24. **, *p<*0.01 by Fisher’s exact test.

To improve transgene expression, we introduced a mutation in the nuclear localization signal (NLS) of the capsid protein (K64N) (**Fig. 1a**). Previous studies have shown that the wild-type VEEV capsid can induce host gene shutdown in an NLS-dependent manner^40^, and that NLS-deficient replicons display reduced cytotoxicity and virulence *in vivo*^41^. Modification of the NLS prevents nuclear accumulation of the capsid protein, thereby reducing the interference with host transcription and cell cycle regulation. VRPs with the wild type and K64N capsid mutation encoding green fluorescent protein (GFP) were generated (GFP-VRP), and the degree of transgene expression was compared in the murine macrophage J774 cell line, since macrophages are susceptible to VEEV infection^37^. Both VRP variants were successfully transduced and expressed GFP in the J774 cell line. When compared to the wild-type capsid VRP, VRP with the capsid mutation showed higher GFP expression, measured by mean fluorescent intensity (MFI) (**Fig. 1c**). The capsid mutant VRP variant is used in subsequent experiments.

To further characterize the cell tropism of VRP to macrophages, human monocyte cell line THP-1 was differentiated and polarized to M1-like or M2-like macrophages *in vitro*. Notably, M2 polarized THP-1 cells showed GFP expression in a dose-dependent manner after transduction, while no expression was seen in M1 polarized cells (**Fig. 1d, left**). To confirm this in primary cells, human monocyte-derived macrophages (hMDMs) were differentiated to M1-like or M2-like macrophages and transduced with GFP-VRP. Although the frequencies of GFP positive cells were lower in hMDMs compared to THP-1 cells, GFP expression was observed in a dose dependent manner but only in M2 polarized cells, consistent with the result in THP-1 cell line (**Fig. 1d, right**). These results indicate that VRP maintains the cell tropism of wild type VEEV, preferentially transducing M2-like macrophages.

### Efficient gene transduction by Alphavirus VRP induced improved control of tumor growth relative to transduction by Lipid Nanoparticle

To deliver the IL-12 transgene to the tumor microenvironment, we generated VRP expressing IL-12 by including genetically linked *Il12a* (p35) and *Il12b* (p40) subunits in the VRP replicon to encode functional murine IL-12 (IL12-VRP). For comparison purposes, we also generated an IL-12 saRNA by *in vitro* transcription that has identical coding elements, and formulated this saRNA with a lipid nanoparticle (IL12-LNP). IL12-VRP and IL12-LNP were transduced to HEK293T and THP-1-derived M2 macrophages with dose-dependent IL-12 secretion measured in both treatments (**Extended Data Fig. 1**). The anti-tumor efficacies of two delivery approaches were then evaluated in the MC38 syngeneic murine colorectal cancer model using i.t. injections. Two different doses of IL12-VRP (1×10^8^ infectious unit, IU, and 1×10^9^ IU) and IL12-LNP (5µg or 10µg total RNA) were injected i.t. and the rate of tumor growth was compared (**Fig. 1e**). Animals treated with IL12-LNP partially inhibited tumor growth in a dose dependent manner without a significant impact on the survival rate compared to the buffer-treated control group. Treatment with low dose IL12-VRP treatment resulted in a significant inhibition (*p*=0.0089, *t*-test) of tumor growth and achieved a 50% survival rate at day 24 post treatment. Treatment with the high dose IL12-VRP achieved complete inhibition of tumor growth with 100% survival rate (**Fig. 1f**). Notably, the effective dose of IL12-VRP (1×10⁹ IU) theoretically contained only ∼5ng of saRNA (**Extended Data Table. 1**) —1,000-fold lower than the 5µg dose of IL12-LNP (containing 5µg of saRNA), suggesting that IL12-VRP is more efficient in tumor inhibition than IL12-LNP. Altogether, IL12-VRP with the capsid NLS mutation demonstrated superiority in inducing IL-12 expression *in vitro* and controlling tumor growth *in vivo* compared to IL12-LNP.

### Immune Cell Population Dynamics in Response to IL12-VRP Treatment

To assess changes in the cell composition within the TME from VRP treatment and to determine which cell types were transduced by VRP, we analyzed tumors and draining lymph nodes (D-LNs) by single-cell RNA sequencing (scRNA-seq) using the 4T1 syngeneic mouse model of breast cancer, an aggressive, poorly immunogenic tumor with substantial myeloid infiltration^42^. BALB/c mice were subcutaneously inoculated with 4T1 cells and received i.t. injections of 1×10⁹ IU of either GFP-VRP or IL12-VRP when tumors reached injectable size. Tumors and D-LNs were harvested one day and three days post-injection, and processed into single cell suspensions for transcriptomic analysis (**Fig. 2a**). The merged scRNA-seq dataset (no. of cells, n=236,794) that includes all the conditions and tissue types was clustered into cell types by transcriptomic signature, and annotated based on canonical marker genes (see Methods) (**Fig. 2b).**

**Fig. 2.**
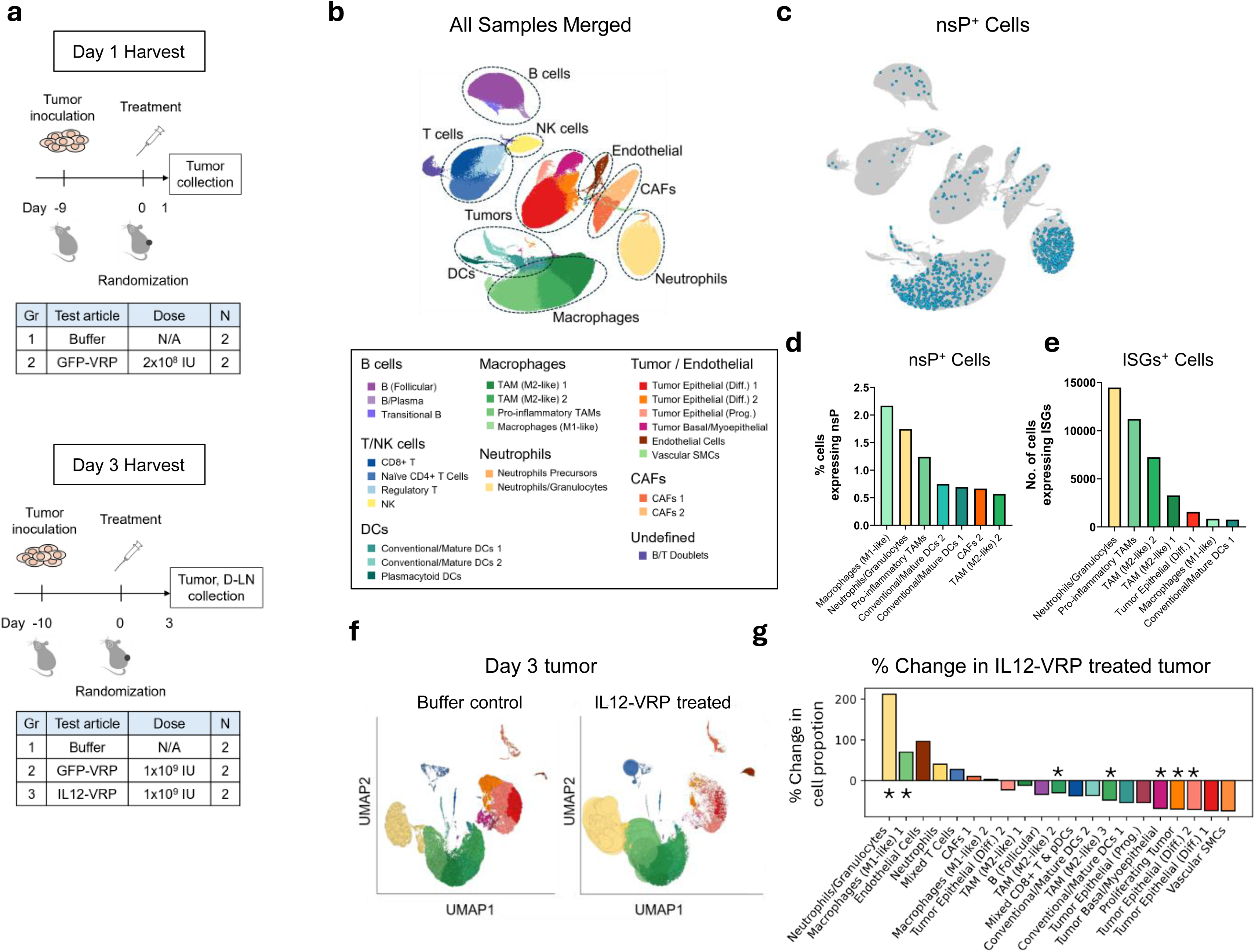
VRP uptake and cell population dynamics in 4T1 mouse tumor microenvironment. **a,** Study design for single cell transcriptomic analysis using 4T1 mouse syngeneic tumor model. **b,** Uniform Manifold Approximation and Projection (UMAP) and cell type annotation of the merged datasets from tumors harvested day 1 and tumors/draining lymph nodes (D-LNs) harvested day 3 post treatment. **c,** VRP uptake in cells in 4T1 tumor and D-LN. Blue dots indicate cells that harbored nsP transcripts. **d,** Proportion of cells expressing nsP transcripts per cell type. **e,** Number of cells in the tumor that expressed ISGs by cell type. **f,** Mapping score of cell types in day 3 tumor. Buffer control (left) and IL12-VRP treated (right). **g,** Proportional change in cell population upon IL12-VRP treatment, compared to buffer control. Asterisk indicates *p*<1e-20 by Chi-squared test.

We assessed frequency of VRP uptake by cell type by identifying cells harboring a non-structural protein (nsP) gene, a vector-derived transcript. No nsP transcript was detected in the buffer control group, confirming the specificity of the signal (**Extended Data Fig. 2a**). Notably, nsP transcripts were detected at high frequency in Macrophages (M1-like) and Neutrophils/Granulocytes (**Fig. 2c, d**), suggesting that these cell populations are primary targets of VRP. Signals of type I and II interferon-stimulated genes (ISGs) were observed in both GFP-VRP and IL12-VRP treated mouse tumors tissues, with a more pronounced effect seen in the IL12-VRP group (*p*<0.0001, Mann–Whitney *U* test) (**Extended Data Fig. 2b**). Neutrophils and macrophages were the cell types that most frequently expressed ISGs (**Fig. 2e**), consistent with the high VRP uptake by these cell types.

Using the day 3 dataset, we assessed the population dynamics of tumor-infiltrating cells following IL12-VRP treatment by mapping cells (see Methods) from IL12-VRP treated tumors onto buffer treated tumors. (**Fig. 2f**). There was a significant enrichment of Neutrophils/Granulocytes expanding +212.3% (from 10.51% to 32.83%, *p*<1e-300, Chi-squared test) and Macrophages (M1-like) increasing +70.0% (from 8.17% to 13.9%, *p*<4.1e-115, Chi-squared test) compared to the buffer control. In contrast, there was a remarkable decrease in multiple epithelial tumor cell clusters and TAM (M2-like) clusters (**Fig. 2g**). Similar trends were found in tumors treated with the GFP-VRP, though the effects were more pronounced in the IL12-VRP treated tumors (**Extended Data Fig. 2c**) suggesting that the replicon vector itself contributes to the remodeling of the tumor microenvironment with IL-12 enhancing its effectiveness.

Overall, i.t. injection of VRP resulted in pronounced VRP uptake in neutrophils and macrophages in the tumor. On day 3 post injection, there was a significant enrichment of neutrophils and M1-like macrophages in the tumor from the IL12-VRP treated mice, with a remarkable decrease in tumor cells and M2-like macrophages.

### Macrophage Phenotypic Reprogramming in Response to IL12-VRP Treatment

Next, we performed a detailed cellular analysis of the macrophage compartment (n=66,540 cells) using the same scRNA-seq dataset (**Fig. 3a**). Pooled across all the samples (tumors harvested day 1 and tumors/D-LNs harvested day 3 post injection, treated and untreated), we observed a diverse TAM populations in the TME, which include Pro-Tumorigenic TAMs (*Vegfa* and *Arg1* high), Locally Proliferating Immunosuppressive TAMs (*Sapcd2* and *Kif14* high), pro-inflammatory TAMs such as M1-like macrophages (*Nos2* and *Jak2* high), and Transitional Monocytic TAMs (*Ly6c1/2* and *Ccr2* high) (**Extended Data Table. 2**).

**Fig. 3.**
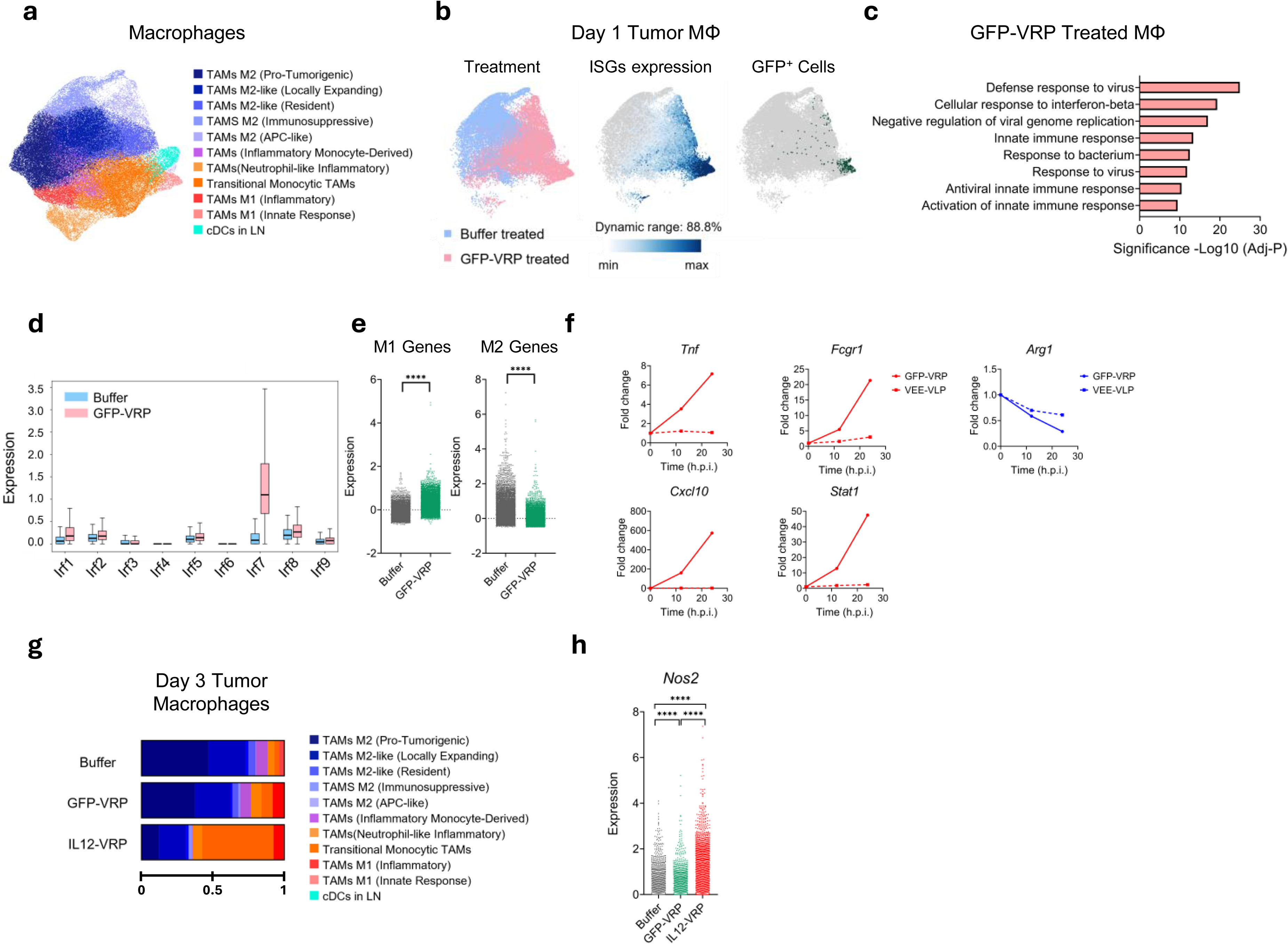
Intratumoral injection of IL12-VRP reprograms tumor-associated macrophages. **a,** Phenotypic analysis of macrophage compartment extracted from day 1 and day 3 merged datasets. Eleven macrophage subsets were identified by gene signatures and color coded. **b,** Cellular status analysis on the macrophage subsets from day 1 tumor. GFP-VRP treated and buffer treated (left), ISGs expression (middle), and GFP expressing (right) macrophage subsets were color coded. **c,** Functional annotation analysis of top 95 upregulated genes in macrophages from GFP-VRP treated tumor, compared to the buffer control. **d,** *Irf* gene expressions in macrophages from GFP-VRP treated or buffer control tumor. **e,** Expression of M1- and M2-marker genes in macrophages from GFP-VRP treated or buffer control tumor. Asterisk indicates statistical significance. ****, *p*<0.0001 by Mann– Whitney *U* test. **f,** qPCR analysis of M1- or M2-marker gene expression kinetics upon *in vitro* transduction of GFP-VRP (MOI 10, solid line) or empty virus-like particle (VLP, dot line) to M2-polarized J774 macrophage cell lines. **g,** Proportional distribution of different macrophage phenotypes in day 3 tumor upon treatment **h,** Gene expression of *Nos2* in macrophages in day 3 tumor upon treatment. Asterisk indicates statistical significance. ****, *p*<0.0001 by Mann–Whitney *U* test.

In these macrophage populations in the TME, we examined whether VRP’s transduction of M2-like macrophages initiates a phenotypic shift to an M1 phenotype *in vivo*. To dissect early molecular signaling by VRP independent of IL-12, we analyzed scRNA-seq dataset from 4T1 tumor harvested 1 day (18 hour) post GFP-VRP treatment (**Fig. 3b, left**). Compared to buffer-treatment, GFP-VRP treated macrophages displayed a distinct transcriptional profile of robust induction of ISGs (**Fig. 3b, middle**). Interestingly, the strong ISGs expression was also observed in the GFP negative macrophages from GFP-VRP treated tumors, suggesting that GFP-VRP treatment indirectly induces activation of non-transduced cells (**Fig. 3b, right**). Functional annotation analysis using the Database for Annotation, Visualization, and Integrated Discovery (DAVID) confirmed elevation of genes involved in the innate response to viral infection including different interferon pathways in the GFP-VRP treated macrophages. (**Fig. 3c**). Notably, when we focused on genes coding for Interferon Regulatory Factors (IRFs), the upregulation of innate responses appeared to be mediated primarily through the virus-induced IRF7 axis (**Fig. 3d**). Moreover, the macrophages from GFP-VRP treated tumors had lower mean intensity of gene expression associated with the M2 phenotype (*p*<0.0001, Mann–Whitney *U* test) and higher expression of genes associated with the M1 phenotype (*p* < 0.0001, Mann–Whitney *U* test), compared to buffer control (**Fig. 3e**).

From the above observations, we hypothesized that VRP transduction could repolarize M2-like macrophages to M1 status. To test this, we examined the kinetics of M1-hallmark gene expression in M2-polarized mouse J774 macrophage cell line upon GFP-VRP transduction *in vitro*. The expression of M1-hallmark genes (*Tnf, Cxcl10, Fcgr1, Stat1*) increased while that of M2-associated gene *Arg1* decreased, indicating a capacity for phenotypic repolarization by GFP-VRP transduction (**Fig. 3f**). These changes were not observed in the M2-polarized macrophages treated with VEE virus-like particles which share the same structural proteins but lack replicon RNA, suggesting that the phenotypic shift to M1 is driven by replicon RNA rather than viral proteins. In addition, the changes above were observed within 24 hours post transduction, which is in alignment with M2 to M1 shift observed *in vivo* as shown in **Fig. 3e**.

Next, we investigated the impact of IL12-VRP treatment on the macrophage populations in the TME on day 3 post treatment. Upon IL12-VRP treatment, we observed a pronounced depletion of M2-like macrophages (from 80.1% to 33.0% collectively), especially Pro-Tumorigenic TAMs (from 46.7% to 11.2%) (**Fig. 3g**), resulting in the macrophage landscape shifting towards pro-inflammatory status. Notably, IL12-VRP had its strongest impact in the increase of Transitional Monocytic TAMs (from 3.5% to 48.1%). This overall pro-inflammatory shift of macrophage population induced by IL12-VRP was further characterized by higher expression of *Nos2* compared to either buffer control or GFP-VRP treatment (**Fig. 3h**). This suggests that IL12-VRP promotes a drastic change in the macrophage populations in the tumor, accompanied by upregulation of proinflammatory genes.

Altogether, these results suggest that i.t. injection of VRP leads to efficient uptake by M2-like macrophages, resulting in initiation of antiviral innate immune response against the VRP. This response, enhanced by IL-12, drives changes from M2- to M1-like macrophages within the TME as well as recruitment of monocytes, reshaping the landscape of tumor-associated macrophage populations towards a pro-inflammatory status.

### Neutrophil Phenotypic Reprogramming in Response to IL12-VRP Treatment

Given that neutrophils exhibited the highest level of VRP uptake (**Fig. 2d**) and there was a significant increase in the Neutrophils/Granulocytes population in the TME following IL12-VRP treatment (**Fig. 2g**), we conducted a detailed analysis of the neutrophil compartment (n=21,611 cells) from the 4T1 scRNA-seq dataset. We identified four distinct neutrophil populations in the TME based on gene expression signatures: *Siglecf^high^*Cancer-Promoting Neutrophils, *Hif1a^high^* Hypoxic Neutrophils, *Sell^high^* Interferon-Stimulated Neutrophils, and *Nos2^high^*Stress-Adapted Neutrophils (**Fig. 4a, Extended Data Table. 3**).

**Fig. 4.**
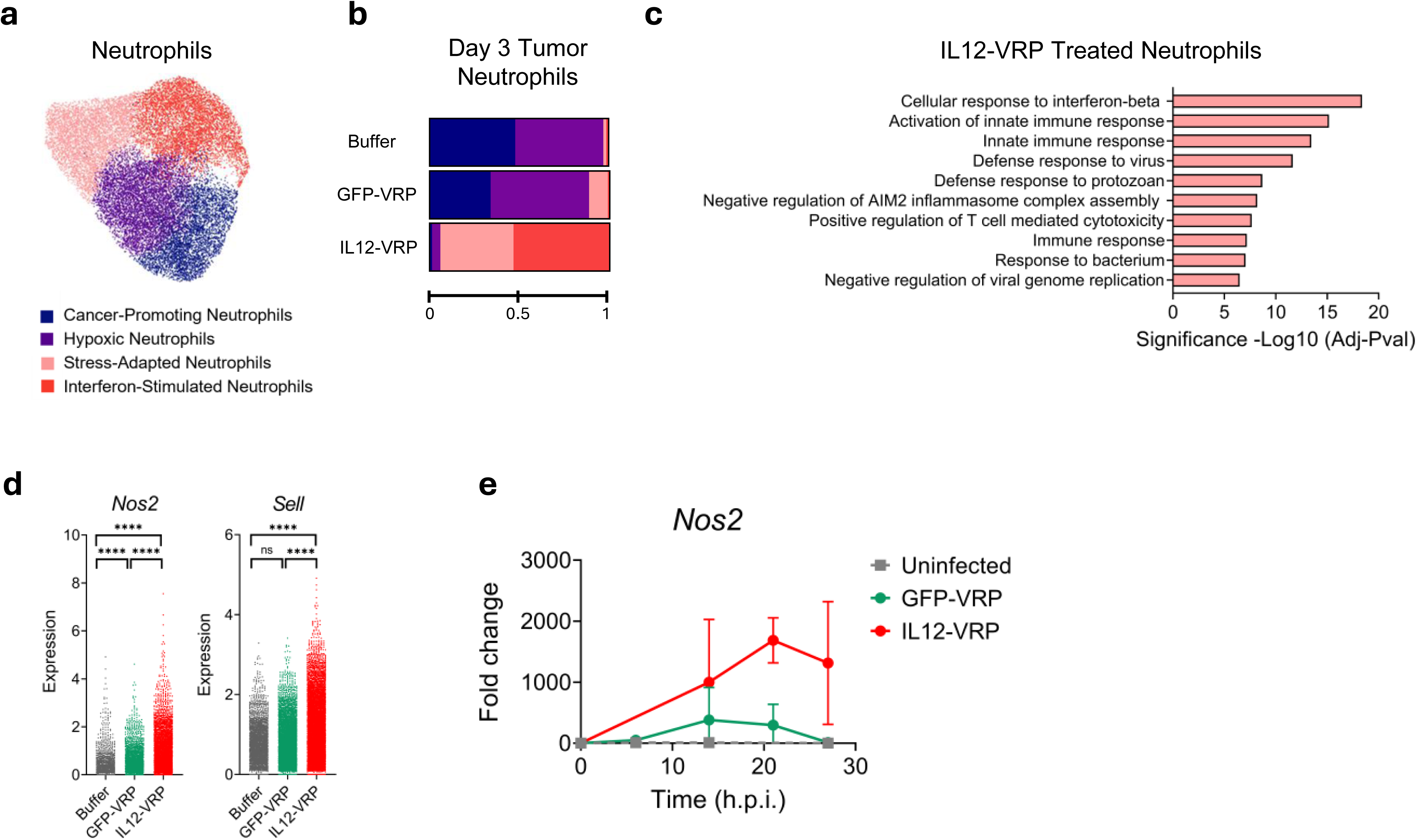
Intratumoral injection of IL12-VRP reprograms tumor-associated neutrophils. **a,** Phenotypic analysis of neutrophil compartment using day 1 and day 3 merged datasets. Four neutrophil subsets were identified by gene signatures. **b,** Proportional distribution of different neutrophil phenotypes in day 3 tumor upon treatment. **c,** Functional annotation analysis of top 216 upregulated genes in neutrophils from IL12-VRP treated tumor, compared to the buffer control. **d,** Gene expression of *Nos2* and *Sell* in neutrophils in day 3 tumor upon treatment. Asterisk indicates statistical significance. ***, *p*<0.001; ****, *p*<0.0001 by Mann–Whitney *U* test. **e,** qPCR analysis on *Nos2* gene expression kinetics upon *in vitro* transduction of VRP (MOI 5) to neutrophils isolated from human peripheral blood.

IL12-VRP treatment, compared to the buffer control, significantly enriched *Sell^high^* Interferon-stimulated (from 1.3% to 53.4%) and *Nos2^high^*Stress-adapted (1.7% to 40.6%) subsets were observed, while *Siglecf^high^*Cancer-Promoting (from 48.1% to 1.4%) and *Hif1a^high^* Hypoxic (from 49.0% to 4.6%) subsets were depleted (**Fig. 4b**), indicating an altered functional state characterized by innate immune activation. DAVID analysis confirmed elevation of genes involved in the innate response to viral infection including interferon pathways in the IL12-VRP treated neutrophils (**Fig. 4c**). This overall pro-inflammatory shift of neutrophil population induced by IL12-VRP was further characterized by higher expression of *Nos2* and *Sell* compared to either buffer control or GFP-VRP treatment (**Fig. 4d**). We confirmed that VRP treatment induced the *Nos2^high^* Stress-adapted phenotype *in vitro* by measuring *Nos2* expression using qPCR in human blood derived neutrophils following transduction with either GFP-VRP and IL-12-VRP (**Fig. 4e**). IL12-VRP elicited a stronger induction than GFP-VRP in alignment with the *in vivo* data. We next explored whether VRP transduction could modulate the phenotypic plasticity of neutrophils, shifting them from an immunosuppressive state to a pro-inflammatory phenotype. Human blood derived neutrophils were cultured in a TME-mimicking cytokine and chemical cocktail to polarize them to the tumor-associated phenotype of N2-like TAN, then treated with GFP-VRP. Analysis of cell surface markers in these cells revealed upregulation of N1-associated markers after GFP-VRP treatment (**Extended Data Fig. 3**), suggesting that VRP treatment could drive phenotypic polarization of neutrophils toward a pro-inflammatory, anti-tumor phenotype.

Taken together, these findings indicate that VRP uptake in neutrophils induces strong interferon signaling, enhanced by transgene IL-12, that triggers reprogramming towards *Nos2*-high and *Sell*-high pro-inflammatory, immune active states.

### IL12-VRP Induces IFN-γ Secretion and Cytotoxicity in NK and T Cells

IL-12 produced by antigen-presenting cells (APCs) is known to stimulate IFN-γ secretion from cytotoxic immune cells, such as NK cells and T cells, thereby enhancing anti-tumor activity within the TME. IFN-γ can further stimulate APCs, forming a positive feedback loop to produce additional IL-12 that amplifies local immune responses^43^. We evaluated whether IL12-VRP initiates this IL-12–IFN-γ signaling cascade using the same 4T1 tumor and D-LN scRNA-seq dataset.

By day 3 post-treatment, a robust induction of *Ifng* expression was observed in NK cells from IL12-VRP treated tumors, with 85.4% of NK cells expressing *Ifng,* compared to 37.1% in the GFP-VRP treated tumors and 20.6% in the buffer control (**Fig. 5a**). This indicates a strong NK cell response to IL-12 transgene expression within the TME. Cell state profiling of NK cell compartment (n=4,272 cells) through inference of the top 50 expressed genes in these cell subsets (**Fig. 5b, Extended Data Table. 4**), further revealed an enrichment of effector/cytotoxic NK cells in IL12-VRP treated tumors, compared to the buffer control (from 0.0% to 56.9%, **Fig. 5c**). Among T cell compartments, 13.0 % of the T cells expressed *Ifng* in IL12-VRP, much higher than the GFP-VRP (5.7%) and buffer control (4.6%) groups (**Fig. 5d**). T cells from IL12-VRP treated tumors also displayed enhanced cytotoxic markers (*Gzma, Gzmb, Prf1, Ifng, Cxcl9*) compared to the buffer control (**Fig. 5e**).

**Fig. 5.**
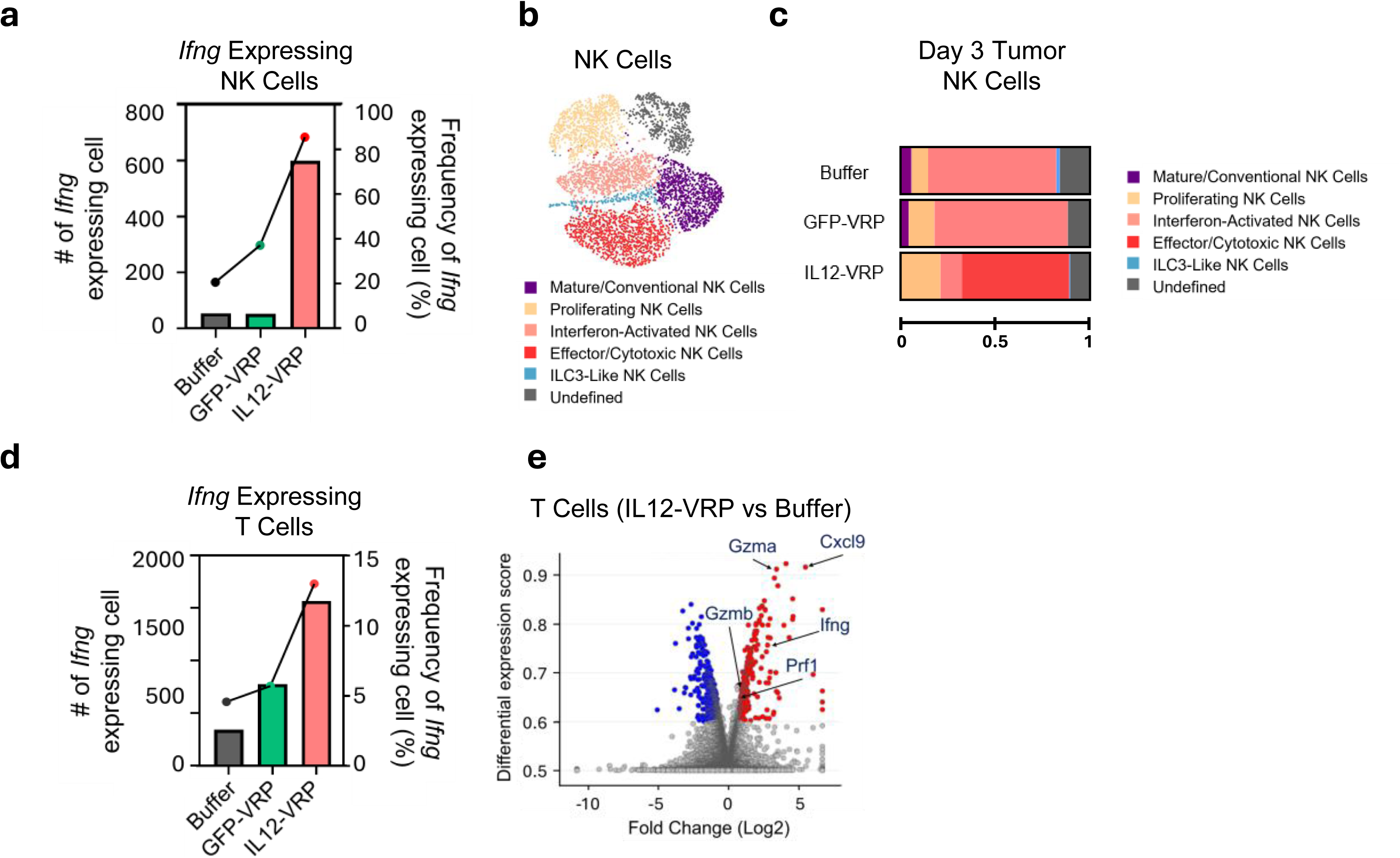
IL12-VRP induces IFN-γ expression and cytotoxicity in NK cells and T cells *in vivo*. **a,** Absolute number (bar) and frequency (line) of *Ifng* expressing cells in the NK cells upon treatment. **b,** Phenotypic analysis of NK compartment using day 1 and day 3 merged datasets. Six NK subsets were identified by gene signatures and color coded. **c,** Distribution of different NK cell phenotypes in the day 3 tumor upon treatment. **d,** Absolute number (bar) and frequency (line) of *Ifng* expressing cells in the T cell cohort upon treatment. **e,** Differentially expressed gene analysis of T cell population (IL12-VRP treatment vs buffer control). The red, blue, and gray dots indicate genes that were upregulated, downregulated, and no change by IL12-VRP treatment, respectively. Genes associated with T cell cytotoxicity are labeled.

Altogether, these data indicate that NK and T cells population expanded and expressed cytotoxic molecules in response to the IL-12 expressed by the transgene.

### Intratumoral Injection of IL12-VRP Prevents Nodal and Distant Metastasis by Reprogramming the Tumor Microenvironment and Draining Lymph Nodes

We hypothesized that the reprogrammed TME by i.t. injection of IL12-VRP would lead to enhanced systemic adaptive immunity capable of suppressing and preventing both nodal and distant metastases. To test this hypothesis, we employed the MOC2 metastatic head and neck cancer (HNC) model in C57BL/6 mice (**Fig. 6a**). Treatment with IL12-VRP i.t. injection led to significant inhibition of tumor growth (*p*<0.0001, *t*-test), whereas treatment with either anti-PD-1 antibody or GFP-VRP exhibited only moderate inhibition effects (*p*=0.0005 and *p*=0.0097 respectively, *t*-test) (**Fig. 6b**). We further assessed nodal and lung metastases by histological analysis. MOC2 tumors typically develop nodal and lung metastases at ∼20 days and ∼30 days post-inoculation, respectively^44^. On Day 21, 100% of the mice in the buffer treated group developed nodal metastases while 33% of the mice in the GFP-VRP treated group developed nodal metastases. Lung metastases were observed in 66% of the buffer group and 50% of the anti-PD-1 antibody treatment group. Remarkably, mice treated with IL12-VRP exhibited no nodal and lung metastases, demonstrating potent anti-metastatic activity (**Fig. 6c**).

**Fig. 6.**
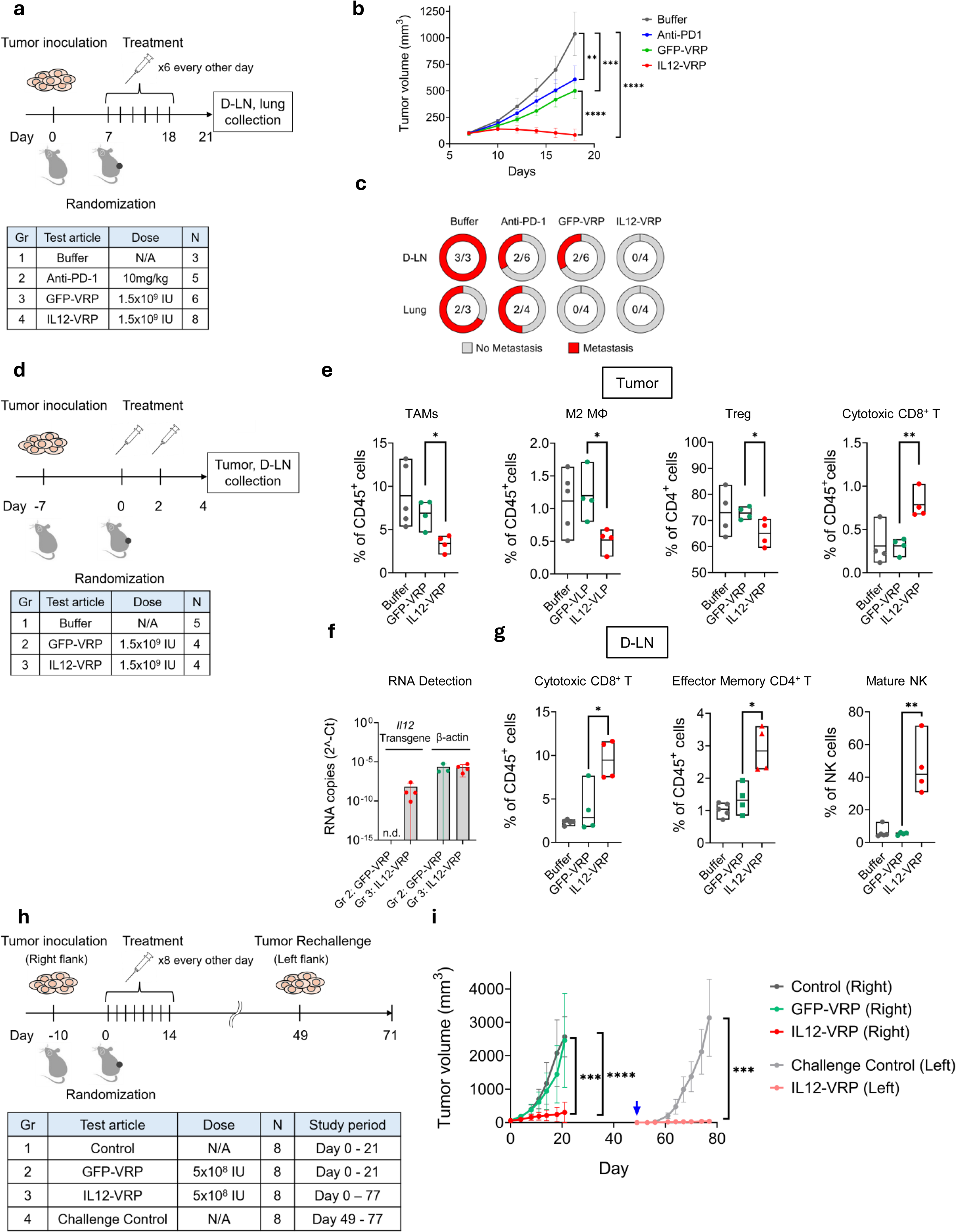
IL12-VRP prevents mouse tumor growth, nodal and distant metastasis, as well as tumor re-challenge by reprogramming local and systemic immune environment. **a,** Study design using MOC2 mouse syngeneic tumor model to evaluate inhibition of primary tumor growth and metastasis. **b,** Tumor growth curve of each treatment group. Asterisk indicates statistical significance on day 18. **, *p*<0.01; ***, *p*<0.001, ****, *p*<0.0001 by *t*-test. **c,** Percentage of metastasis observed in D-LN and lung tissue. **d,** Study design using MOC2 mouse syngeneic tumor model to assess immune microenvironment of primary tumor and D-LN. **e,** Flow cytometry analysis on immune cell population in primary tumor. Tumor associated macrophage (TAM) population (% CD11b^+^ F4/80^+^ Ly6G^-^ from isolated CD45^+^ cells). M2 macrophage population (% CD11b^+^ F4/80^+^ Ly6G^-^ CD206^+^ MHC Class II ^low^ from isolated CD45^+^ cells). Regulatory T cell population (% FoxP3^+^ from isolated CD45^+^ CD4^+^ cells). Cytotoxic CD8^+^ T cell population (% CD8^+^ NFATc1^+^ from isolated CD45^+^ cells). **f,** qPCR analysis showing IL-12 transgene detection. PCR primers were designed to distinguish IL-12 transgene from endogenous IL-12. The house-keeping gene *Actb* was also detected as assay control. n.d. = not detected. **g,** Flow cytometry analysis on immune cell population in D-LN. Cytotoxic CD8^+^ T cell population (% CD8^+^ NFATc1^+^ from isolated CD45^+^ cells). Effector Memory CD4^+^ T cell population (% CD4^+^ Tbet^+^ from isolated CD45^+^ cells). Mature NK population (% CD27^-^ CD11b^+^ from isolated CD19^-^/CD3^-^/CD45^+^/NKp46^+^/NK1.1^+^/DX5^+^ cells). *, *p*<0.05; **, *p<*0.01 by *t*-test. **h,** Study design using CT26 mouse syngeneic tumor model to evaluate IL12-VRP’s ability to protect animals from repeated tumor inoculation. **i,** Tumor growth curve of the initial tumor on the right flank (day 0 to 21) and the rechallenging tumor on the left flank (day 49 to 77). Arrow indicates the day of tumor rechallenge (day 49). Asterisks indicate statistical significance between Group 1 and 3 and Group 2 and 3 on day 21; Group 3 and 4 on Day 77. ***, *p*<0.001; ****, *p*<0.0001 by *t*-test.

To explore the mechanisms underlying this effect, we performed an additional *in vivo* study with the MOC2 model, focusing on immune landscape remodeling in both the primary tumor and D-LNs (**Fig. 6d**). Flow cytometry analysis of the primary tumors showed that IL12-VRP treatment reduced the frequency of tumor-associated macrophages (TAMs) (*p*=0.0107, *t*-test), particularly M2-like immunosuppressive macrophages (*p*=0.0178, *t*-test) compared to GFP-VRP treatment. It also reduced regulatory T cells (Tregs) (*p*=0.0336, *t*-test) and increased the abundance of cytotoxic CD8⁺ T cells (*p*=0.0023, *t*-test), indicating the establishment of a more immune-permissive, anti-tumor TME, especially after IL12-VRP treatment (**Fig. 6e**). Similar results were confirmed in the MC38 colon cancer model (**Extended Data Fig. 4**).

In D-LN, VRP-derived IL-12 RNA was detected following i.t. injection (**Fig. 6f**), suggesting that either the cells transduced in the tumor migrated to the D-LNs, or IL12-VRP directly reached the D-LNs. Thus, immunomodulatory effects from IL12-VRP were expected in the D-LNs. Flow cytometry analysis of cells in the D-LN revealed an increase in cytotoxic CD8⁺ T cells (*p*=0.0179, *t*-test), effector memory CD4⁺ T cells (*p*=0.0103, *t*-test), and a notable shift toward mature NK cell phenotype (*p*=0.0037, *t*-test) by IL12-VRP treatment, compared to GFP-VRP treatment (**Fig. 6g**). These findings suggest that injected VRPs not only modifies the TME, but also impacts the D-LN microenvironment, which may result in suppression of tumor growth and potential metastasis by induction of systemic immune activation.

### IL12-VRP Induces Systemic Tumor-Specific Immune Memory

Given the observed increase in cytotoxic CD8⁺ T cells in the D-LNs following i.t. IL12-VRP injection, we next investigated whether this enhanced immunity could protect the animals from re-challenge with tumor inoculation. To evaluate this, we utilized the CT26 colon carcinoma model in BALB/c mice (**Fig. 6h**). IL12-VRP treatment again significantly reduced tumor volumes compared to buffer control (*p*<0.0001, *t*-test) (**Fig. 6i**). To assess whether the treatment induced durable systemic immunity, four mice that survived the initial tumor challenge were re-challenged with CT26 cells into the contralateral flank on Day 49. Eight treatment-naïve mice were used as tumor growth controls and received the same number of CT26 cells on the same day. All the control mice developed rapidly growing tumors. In contrast, none of the animals previously treated with IL12-VRP exhibited measurable tumor growth after re-challenge (*p*=0.0004, *t*-test) (**Fig. 6i**).

These results demonstrate that IL12-VRP therapy not only reduces primary tumor growth but also induces durable systemic immune memory that confers protection against subsequent tumor re-challenge.

## Discussion

To address limitations of the current immunotherapies, we developed a novel therapy that reprograms the TME by targeting tumor-associated macrophages and neutrophils leveraging the natural tropism of alphavirus. These cells, upon VRP uptake, initiate robust innate immune responses via the induction of ISGs, further enhanced by localized expression of IL-12. This cascade facilitates the activation of cytotoxic NK and T cells, resulting in promotion of durable systemic immunity that prevents dissemination of cancer cells.

Among TAMs-targeted therapies, inhibitors for phosphatidylinositol 3-kinase gamma (PI3K-γ)^45,46^ are currently in clinical development. PI3K-γ inhibition has been shown to reprogram TAMs and suppress tumor growth^47–49^. These inhibitors enhanced the expression of pro-inflammatory molecules such as IL-12, IFN-γ and NOS2, while suppressing immune suppressive molecules such as Arg1 in TAM. Notably, blockade of IL-12 or NOS2 in PI3K-γ-deficient macrophages promoted tumor progression, demonstrating the importance of IL-12 and NOS2 from the macrophage-derived immune factors for the reprogramming of TME. Given the marked elevation of *Nos2* in both TAMs and TANs following IL12-VRP treatment, our approach might concurrently accelerate to reprogram TME.

Recent studies have highlighted the crucial role of crosstalk among TAMs and TANs in shaping effective anti-tumor immunity^19,20,50,51^. In a mouse model, interactions between TAMs and TANs were shown to promote the development of intrahepatic cholangiocarcinoma cells^52^. Conversely, neutrophil–macrophage interactions were shown to increase macrophage production of IL-12, which in turn induced IFN-γ expression and enhanced their anti-tumor activity^53^. Building on this biology, our approach is distinctive in that it simultaneously targets both TAMs and TANs, enabling coordinated myeloid activation and potentially amplifying the IL-12–IFN-γ axis that drives adaptive anti-tumor immunity.

Further investigation into the mechanism by which VRP efficiently transduces M2-like macrophages, while sparing M1-like macrophages, would be valuable. Alphaviruses enter cells through receptor-mediated endocytosis and are released from endosomes via a pH-depended fusion with the endosomal membrane^54^. M2-like macrophages express distinct surface molecules such as the DC-SIGN and mannose receptor^55,56^, which are known to be a receptor or have enhanced binding for viruses such as HIV-1 and dengue virus^57–59^. Additionally, the endosomal pH, lipid composition, and protease profile in M2 macrophages may create a more favorable environment for efficient viral uncoating and membrane fusion, as well as support viral RNA replication and translation. The differences between M2 and M1-like macrophages may lead to preferential VRP transduction of the M2 type; however, further work is needed to confirm this hypothesis.

In conclusion, IL12-VRP is a novel immunomodulatory approach that targets immunosuppressive myeloid populations. While further validation in both preclinical and clinical settings is necessary, including our ongoing Phase I clinical trial in patients with head and neck cancer (NCT06736379), this approach holds promise, particularly for solid tumors characterized by myeloid-dominant TMEs. Given the central role of immunosuppressive myeloid cells in resistance to immune check-point therapies, strategies that directly reprogram these populations may offer an alternative therapeutic approach. Future studies will define predictive biomarkers of response and test combinations to broaden clinical benefit across multiple tumor types.

## Materials and Methods

### Plasmids

The VEEV replicon plasmid was generated by two steps: The 5’ untranslated region (UTR), nonstructural protein (nsP) 1-4, subgenomic promoter and 3’ UTR of the Venezuelan equine encephalitis virus (VEEV) TC-83 strain (GenBank# L01443.1) were inserted into a mammalian expression vector with RSV promoter; either synthesized green fluorescent protein (GFP) or murine IL-12p40 and p35 linked by peptide was inserted into the VEEV replicon plasmid between the subgenomic promotor and 3’ UTR. The VEEV replicon plasmid, designed for in vitro transcription was generated by replacing the RSV promoter with a T7 promoter. The VEEV helper plasmid was generated by inserting the capsid and envelope sequences of the VEEV TC-83 strain into a mammalian expression vector with CMV promoter.

### Cell lines

293T/17, Vero, THP-1 and J774A.1 cell lines were obtained from ATCC and maintained in culture at 37℃ and CO_2_ concentration at 5%. The growth medium used for each cell line was: DMEM (Gibco) supplemented with 10% FBS (GeminiBio) and 1% penicillin/streptomycin (pen/strep, Gibco) for 293T/17 and J774A.1 cells. RPMI1640 (Gibco) supplemented with 10% FBS and 1% pen/strep for THP-1 cell. EMEM (ATCC) supplemented with 10% FBS and 1% pen/strep for Vero cell. MOC2 cells were obtained from Kerafast and maintained in culture in IMDM/F12 (2:1) with 5% FCS, 1% penicillin/streptomycin, amphotericin, 5 ng/mL EGF (Millipore), 40 ng/mL hydrocortisone, and 5 μg/mL insulin.

### VRP Production, purification, and quantification

The VEEV replicon plasmid and VEEV helper plasmids were co-transfected into 293T/17 cells using polyethyrenimine (Polysciences, US). The culture supernatant was collected at 42-hour and 66-hour post transfection and purified by combination of tangential flow filtration (Repligen), CaptoCore700 chromatography (Cytiva), and sterilized by polyethersulfone membrane filter with pore size of 0.22µm.

### Isolation of Human Monocytes and Neutrophils

Anti-coagulated human peripheral blood obtained from healthy donors was segmented using Histopaque-1077 (Sigma Aldrich) according to the manufacturer’s instruction. Briefly, the peripheral blood was poured onto the same amount of Histopaque in a 50 mL conical tube. The tube was centrifuged at 400x g for 45 minutes at room temperature. The resulting PBMC layer was collected and the CD14^+^ monocytes/macrophages were isolated by positive selection using MacroBeads conjugated CD14 antibody (Miltenyi #130-050-201) and column (Militenyi #130-042-041). Separately, the granulocyte/erythrocyte sediment on the bottom was collected and the erythrocytes further sedimented by adding 2% dextran and letting the tube stand upright at room temperature for 20 minutes. The upper erythrocyte-poor layer was collected, and the residual erythrocytes were destroyed by treating cells in reduced tonicity for 30 seconds. The isolation of CD14 monocytes and neutrophils was confirmed by flowcytometry analysis. The panel of antibodies used for cell surface staining included fluorophore-conjugated anti-human CD45 (HI30, #416-0459-41), CD14 (61D3, #12-0149-41), CD16 (3G8, #64-0166-42), CD15 (MMA, #46-0158-41) from ThermoFisher; CD66b (6/40c, #392903) from BioLegend.

### In Vitro Macrophage Polarization

J774 mouse macrophage cell line was treated with recombinant mouse macrophage colony stimulating factor (M-CSF, Peprotech) at 50ng/mL. Twenty-four hours later, the media was replaced with fresh media containing either recombinant mouse IFN-γ (Peprotech) at 20 ng/mL and lipopolysaccharide (LPS, Sigma) at 100ng/mL for M1 polarization, or recombinant mouse IL-4 (Peprotech) at 20ng/mL for M2 polarization.

Polarization was confirmed by gene expression analysis by qPCR, using gene-specific primers for mouse *Il1b*, *Il6*, *Arg1*, *Tgfb2,* and *Vegfa*. Primer sequences are listed in **Extended Data Table 5**. THP-1 human monocyte cell line was treated with phorbol 12-myristate 13-acetate (PMA, Sigma) at 50nM. Twenty-four hours later, the media was replaced with fresh media containing either recombinant human IFN-γ (Peprotech) at 20 ng/mL and LPS (Sigma) at 10ng/mL for M1 polarization, or recombinant human IL-4 (Peprotech) at 20ng/mL for M2 polarization. Primary human CD14^+^ cells were treated with recombinant human M-CSF (Peprotech) at 20ng/mL for 5∼6 days, then media was replaced by fresh media containing either recombinant human IFN-γ at 20ng/mL and LPS at 10ng/mL for M1 polarization, or recombinant human IL-4 at 20ng/mL for M2 polarization. Polarization was confirmed by gene expression analysis by qPCR, using gene-specific primers for human *Cxcl10, Calhm6, Cd209, Tgm2,* and *Ccl26*. Primer sequences are listed in **Extended Data Table 5**.

### In Vitro Neutrophil Polarization

Neutrophils isolated from human peripheral blood were polarized to N1- or N2-like status following a protocol adapted from Ohms et al ^60^. Briefly, 1.5×10^7^ neutrophils were cultured in six-well plates in 3mL RPMI1640 supplemented with 10% FBS, 1% pen/strep, and 3 µM pan-caspase inhibitor QVD-Oph (R&D Systems). For polarization, either N1 cocktail containing 100ng/mL lipopolysaccharide (SigmaAldrich), 50 ng/mL IFN-γ (PeproTech), and 50 ng/mL IFN-β (Peprotech), or N2 polarization cocktail containing 25mM L-lactate (SigmaAldrich), 10 µM adenosine (Merck), 20ng/mL TGF-β (PeproTech), 10ng/mL IL-10 (PeproTech), 20 ng/mL prostaglandin E2 (PGE2; Tocris), and 100ng/mL granulocytecolony-stimulatingfactor (PeproTech) was added to the culture and incubated at 37℃ for 24 hours. The polarization neutrophils were confirmed by flowcytometry analysis. The panel of antibodies used for cell surface staining included fluorophore-conjugated anti-human CD182 (5E8-C7-F10, #46-1829-42) and CD62L (DREG-56, #407-0629-41) from ThermoFisher; CD66b (6/40c, #392903), CD54 (HA58, #353131), CD95 (DX2, #305627), and CD11b (M1/70, #101216) from BioLegend.

### ELISA

IL-12 concentration in the culture supernatant was measured by sandwich ELISA using Mouse IL-12 p70 DuoSet ELISA kit (R&D Systems DY419) following user manual. Briefly, microplates were coated with mouse IL-12 capturing antibody overnight. After blocking, samples and recombinant standard IL-12 were applied to the plate in serial dilution. After 2-hour incubation at room temperature (RT), the captured IL-12 was probed by detecting IL-12 antibody, followed by streptavidin-HRP. The resulting signal from the substrate application, followed by stop solution, was measured as absorbance at 450nm by a BioTek plate reader.

### qPCR

Vector-derived RNA or endogenous RNA in the transduced cells was quantified by qRT-PCR method. Briefly, cells were collected at the indicated time points and RNA was extracted using RNeasy Mini Kit (Qiagen #74106). Reverse transcription was conducted using iScript cDNA Synthesis Kit (BioRad #1708891), and resulting cDNA was used as a template of quantitative PCR using PowerTrack SYBR Green Master Mix for qPCR (ThermoFisher #A46110). Primer sets used for gene expression analysis included human IL-12 transgene, GFP transgene, mouse *Tnf, Stat1, Cxcl10, Fcgr1, Arg1*, and human *Nos2*. Primer sequences are listed in **Extended Data Table 5**. RNA copy number was calculated using standard curve made with GFP replicon plasmid with known concentration, as a qPCR template.

### Flow cytometry

Infectious titer: The infectious titer of GFP/IL12-VRP was determined through infectivity of Vero cells using a flow cytometry-based assay. Briefly, Vero cells were plated at 2×10^5^ cells per well in 24-well plate 6 hours prior to VRP transduction. GFP/IL12-VRP were serial-diluted and added to the cells. Eighteen hours post transduction, cells transduced with IL12-VRP were collected, fixed, permeabilized and probed by rat anti-mouse IL12 (p40/p70) antibody conjugated with PE (BDbiosciences #554479); cells transduced with GFP-VRP were collected and washed in PBS. The percentage of GFP/IL-12 positive cells were measured by flow cytometry, and infectivity curve was drawn to determine the infectious titer.

In vivo Tumor Study (MOC2 Model): Tumors were digested using the mouse tumor dissociation kit (Miltenyi #130-096-730), following the manufacturer’s guidelines. Spleen and lymph nodes were mechanically dissociated through a 70 μm cell strainer and washed with RPMI media. The cells were further washed with PBS and then stained with Live/Dead Zombie NIR™ (ZNIR) Fixable Viability Kit (1:500; BioLegend #423105, BioLegend) for 10 minutes at 4°C. After staining, the cells were washed in FACS buffer (PBS supplemented with 2% FCS) and Fc receptors were blocked using TruStain FcX™ (1:100, anti-mouse CD16/32, BioLegend #101319d). Subsequently, the cells were stained with a panel of antibodies diluted in FACS buffer for 30 minutes at 4°C in the dark. The panel of antibodies used for cell surface staining included fluorophore-conjugated anti-mouse CD45.2 (30-F11, #103147), TCR-beta (H57-597), CD4 (GK1.5, # 100411), CD8 (53-6.7, #100712), NK1.1 (PK136), CD11b (M1/70, #101206), CD11c (N418, #117333), MHCII (M5/114.15.2), Ly6G (1A8, #127607), Ly6C (HK1.4, # 128041), F4/80 (BM8,#123131) from BioLegend (San Diego, CA). For intracellular staining, the cells were fixed using the Fixation-Permeabilization kit protocol (eBioscience #88-8824-00), and then intracellular staining was performed for CD206 (MMR, BioLegend #141720), NFATc-1 (BioLegend # 649606), Foxp3 (Invitrogen # 72-5775) and IL-10 (BioLegend #505028). Following staining, the cells were washed with FACS buffer and analyzed using

Flow cytometry data were acquired using a BD LSR II flow cytometer and analyzed using FlowJo software (Tree Star).

### Electron Microscopy

Samples (0.4 mg/ml, 4.5 µL) were adsorbed to glow discharged (EMS GloQube) ultra-thin (UL) carbon coated 400 mesh copper grids (EMS CF400-Cu-UL), by floatation for 10 seconds. Grids were quickly blotted then rinsed in 3 drops (1 min each) of TBS. Grids were negatively stained in 2 consecutive drops of 0.75% uranyl formate (UF), then quickly aspirated. Grids were imaged on a ThermoFisher Talos L120C TEM operating at 1200 kV with a ThermoFisher Ceta (cooled 16 Mpixel CMOS). The diameter of 50 representative particles was measured in ImageJ.

### In vitro RNA Transcription and Lipid Nanoparticle Formulation

The IL12 VEEV replicon plasmid with T7 promoter was linearized by digestion with the Nrul or BspQ1 restriction enzymes at 37 °C or 50 °C for 3 h. The linearized plasmid was then purified using the Wizard Plus SV Miniprep DNA Purification System (Promega), and saRNA was transcribed in vitro using the T7 RiboMAX Express Large-Scale RNA Production System (Promega). After DNase treatment, the saRNA was purified with RNeasy midi kit (Qiagen), and subsequently modified by the addition of a 7-methylguanosine cap with the Vaccinia Capping System (New England Biolabs [NEB]) using the NEB Capping protocol (NEB, M20280). After purification of the capped saRNA using the Monarch kit (NEB), LNP-saRNA was formulated with GenVoy-ILM (Precision Nano Systems) with the NanoAssemblr Ignite (Precision Nano Systems). The mean hydrodynamic diameters were measured by dynamic light scattering instrument (Zetasizer ELSZ-200ZS, Malvern Instruments Ltd.) for material consistency.

### Syngeneic Mouse Tumor Models

MC38: The animal study was conducted at Crown Bioscience San Diego (San Diego, California, USA) in accordance with IACUC guideline. Cryogenic vials containing MC38 tumor cells were thawed and cultured according to the manufacturer’s protocol. On the day of injection, cells were counted and resuspended in cold RPMI. Six- to eight-week-old female C57BL/6J mice were purchased from the Jackson Laboratory and prepared for injection using standard approved anesthesia. A suspension of 1×10^6^ viable cells/100μL was subcutaneously injected into the rear flank of the mouse. Once tumors are palpable, tumors were measured 2 times a week using calipers. Tumor volume was calculated using the following equation: (longest diameter x shortest diameter^2^)/2. Once tumors were of appropriate size to begin the study, tumors and body weights were measured 2 times per week for the duration of the study. When the average tumor volume reached 50-100 mm^3^, mice were randomly assigned to the respective treatment groups and dosed the same day. All the animals received either PBS + 4% sucrose buffer, IL12-VRP, or IL12-LNP in a 50µL volume, intratumorally. The animals were euthanized if the tumor burden reached 2000mm^3^ or the study period ended.

MOC2: The animal study was conducted at Stanford University School of Medicine (Stanford, California, USA) in accordance with APLAC guidelines. Cryogenic vials containing MOC2 tumor cells were thawed and cultured according to the manufacturer’s protocol. On the day of injection, cells were counted and resuspended in PBS with 20% reduced growth factor Matrigel (Corning). Seven- to nine-week-old male C57BL/6 mice were purchased from the Jackson Laboratory and prepared for injection using standard approved anesthesia. A suspension of 3×10^5^ viable cells/100μL was subcutaneously injected into the rear flank of the mouse. Once tumors are palpable, tumors were measured 2 times a week using calipers. Tumor volume was calculated using the following equation: (longest diameter x shortest diameter^2^)/2. Once tumors were of appropriate size to begin the study, tumors were measured 2 times per week for the duration of the study. When the average tumor volume reached 90-100 mm^3^, mice were randomly assigned to the respective treatment groups and dosed after 24 hours of randomization. All the animals received either TNE+5% sucrose buffer, GFP-VRP or IL12-VRP in a 50µL volume, intratumorally. The anti-PD-1 monoclonal antibody (Clone J43) was obtained from Bio X Cell and was administered at a dose of 10mg/kg, twice a week for three weeks. The animals were euthanized if the tumor burden reached 1700mm^3^ or the study period ended. Tumors and draining lymph nodes were collected at sacrifice and analyzed with flow cytometry for immune cell changes (short-term study). The lungs and draining lymph nodes were collected and subjected to H&E staining to evaluate nodal and lung metastasis (long-term study).

CT26: The animal study was conducted at Crown Bioscience San Diego (San Diego, California, USA) in accordance with IACUC guideline. Cryogenic vials containing CT26 tumor cells were thawed and cultured according to the manufacturer’s protocol. On the day of injection, cells were counted and resuspended in cold RPMI. Six- to nine-week-old female Balb/c mice were purchased from the Jackson Laboratory and prepared for injection using standard approved anesthesia. A suspension of 5×10^5^ viable cells/100 μL was subcutaneously injected into the rear flank of the mouse. Once tumors are palpable, tumors were measured 2 times a week using calipers. Tumor volume was calculated using the following equation: (longest diameter x shortest diameter^2^)/2. Once tumors were of appropriate size to begin the study, tumors and body weights were measured 2 times per week for the duration of the study. When the average tumor volume reached 50-100 mm^3^, mice were randomly assigned to the respective treatment groups and dosed the same day. All the animals received either PBS + 4% sucrose buffer, GFP-VRP, or IL12-VRP in a 50µL volume, intratumorally. The animals were euthanized if the tumor burden reached 2000mm^3^ or the study period ended. Survived animals (N=4) from the IL12-VRP treatment group, and treatment-naïve animals (N=8) were challenged by subcutaneously injecting CT26 cells on the left flank on Day 49.

4T1: The animal study was conducted at Noble Life Sciences (Sykesville, Maryland, USA) in accordance with IACUC guideline (NLS-23-10-032.A2). Cryogenic vials containing 4T1 tumor cells were thawed and cultured according to the manufacturer’s protocol. On the day of injection, cells were counted and resuspended in PBS containing 20% Matrigel. Six- to eight-week-old female BALB/c mice were purchased from Envigo and prepared for injection using standard approved anesthesia. A suspension of 5×10^5^ viable cells/100μL was subcutaneously injected into the rear flank of the mouse. Once tumors are palpable, tumors were measured 2 times a week using calipers. Tumor volume was calculated using the following equation: (longest diameter x shortest diameter^2^)/2. Once tumors were of appropriate size to begin the study, tumors and body weights were measured 2 times per week for the duration of the study. When the average tumor volume reached 200-500 mm^3^, mice were randomly assigned to the respective treatment groups and dosed the same day. All the animals received either PBS + 4% sucrose buffer, GFP-VRP, or IL12-VRP in a 50µL volume, intratumorally. Sixteen hours after dosing, following euthanasia, tumors were collected aseptically (Day 1 harvest). On Day 3, following euthanasia, tumors and draining lymph nodes were collected aseptically (Day 3 harvest). The tissues were processed for Tumor Dissociation and Single Cell RNA Sequencing on the day of harvest.

### Tumor Dissociation and Single Cell RNA Sequencing

4T1 mouse tumors were collected aseptically and digested following the protocol ^61^. Briefly, tissue was chopped into pieces and went through a sequence of enzymatic and chemical digestion including collagenase (Sigma Aldrich #C9891), hyaluronidase (Sigma Aldrich #H3506), DNase I (Roche #10104159001), dispase II (Roche #04942078001), TEG (trypsin (Gibco #15090-046), Ethylene glycol-bis(2-aminoethylether)-N,N,N’,N’-tetraacetic acid (Sigma #E3889), polyvinyl alcohol (Sigma #P8136)). Red blood cells were lysed and washed out. The resulting single cell suspension was adjusted to 1∼3 x 10^6^ cells/mL in PBS. The sequence library was prepared for the 10x Genomics 5’ gene expression system (10x Genomics) according to the instruction and sequenced by NovaSeq X Plus (Illumina).

### Single Cell RNA Sequence Data Analysis

Quality control and preprocessing: Raw count matrices were imported into the scarf analysis environment ^62^. Initial quality control metrics were calculated for each cell, including total UMI counts, number of detected genes, and percentage of mitochondrial and ribosomal gene expression. Cells were retained if they met the following criteria: 5,000-100,000 total UMI counts, 1,000-10,000 detected genes, <10% mitochondrial gene expression, and <40% ribosomal gene expression. Low-quality cells failing these thresholds were excluded from downstream analysis.

Feature selection, Clustering and cell type annotation: The top highly variable genes (HVGs) were identified using variance-stabilized selection. Principal component analysis was performed using the selected HVGs, and the optimal number of principal components was determined using elbow plot analysis. Uniform Manifold Approximation and Projection (UMAP) was performed for two-dimensional visualization using 2,000 epochs with parallel processing. Leiden clustering was applied to identify distinct cell populations. Marker gene identification and differential gene expression (DEG) between cell sets were performed using ‘run_marker_search’ function of scarf 0.31.3. The marker score between 0 and 1 was ascribed to each cell with a score of 1 reflecting that the gene was present in all the cells of the cluster (cell set) and not in any other cluster (or the other cell set). The top 50 marker genes from each of the clusters used for cell type annotation.

Integration and comparative analysis: For integrated analysis across all samples, individual datasets were merged using scarf’s dataset integration framework, resulting in a combined dataset of 236,794 cells. Changes in cell type frequencies between conditions were assessed using Chi-squared tests of independence. For each cell type, contingency tables were constructed comparing cell counts between control and treatment groups relative to total cell numbers.

Reference mapping analysis: To assess treatment-specific effects while controlling for cross-dataset heterogeneity, reference mapping analyses were performed for each tissue and timepoint combination. Buffer control samples served as reference datasets, with GFP-VRP and IL12-VRP treatment samples mapped to the reference using established cell type annotations from Leiden clustering. For each mapping comparison, two complementary statistical analyses were performed to quantify treatment effects. First, paired t-tests were conducted on mapping scores between control and treatment conditions within each cluster, with log2 fold changes calculated and p-values adjusted using Bonferroni correction for multiple comparisons. Second, Chi-squared tests of independence were applied to assess changes in cell type proportions between conditions, calculating percent changes in cellular composition and testing for statistical significance of shifts in cluster representation.

Interferon response and macrophage polarization signature analysis: A z-score-based gene set aggregation approach was implemented. For interferon pathway analysis, tumor samples at day 3 post-treatment and Type I and Type II interferon response gene sets GO:0034340 and GO:0034341 respectively, were used. For macrophage polarization analysis, macrophage subset from tumor samples at day 1 post-treatment and M1 and M2 gene sets manually curated from established polarization markers were used: M1 genes included *Nos2, Il1b, Il6, Tnf, Il12b, Il18, Il15, Ccr7, Cd38, Fpr2, Gpr18, Cd80, Cd86, Cd68, H2-Ab1, H2-Eb1, Psme2, Irf1, Irf9, Stat1, Stat2, Jak2, Ifnar2, Ifi35, Ifitm3, Tap1, Psmb8, Rela, Trem1, Socs3*; M2 genes included *Arg1, Mrc1, Retnla, Egr2, Myc, Cd163, Clec10a, Pparg, Csf1r, Pdcd1lg2, Fn1, Msx3, Itgb3*. For both analyses, normalized expression values for all genes within each gene set were extracted, with a small pseudocount (1×10⁻⁷) added to handle zero-inflation. Expression values were z-score standardized across all cells for each gene individually, then aggregated by calculating the mean z-score across all genes within each pathway for every cell. This approach generates cell-level scores representing relative pathway activity, where positive scores indicate above-average pathway activation and negative scores indicate below-average activity. The resulting aggregated expression scores were plotted.

Statistical analysis and data visualization: All statistical analyses were performed in Python using scipy and statsmodels scientific computing libraries. Cell type proportion differences between treatment groups were assessed using appropriate statistical methods with multiple hypothesis correction. Data visualization was performed using matplotlib and custom plotting functions.

## Acknowledgements

We thank Parashar Dhapola and Yi Su from Nygen Analytics for their contributions to the analysis of the single-cell RNA sequencing data. Their expertise in bioinformatics supported the interpretation and visualization of the results.

## Author contributions

M.I., J.S., F.B., K.M., F.M.B., J.B.S., Q.T.L., J.F.S. W.A. conceived the project. M.I., D.K.N., J.S., F.B., J.B.S., Q.T.L., J.F.S. and W.A. designed the experiments. M.I., D.K.N., J.S., F.B., K.I., F.T., K.T., I.S., H.C., D.L. performed experiments and data analysis. K.M., F.M.B., J.B.S., Q.T.L., J.F.B. and W.A. provided supervision and resources. All authors discussed the results and contributed to writing and/or editing the manuscript.

---

**Extended Data Figure 1.**
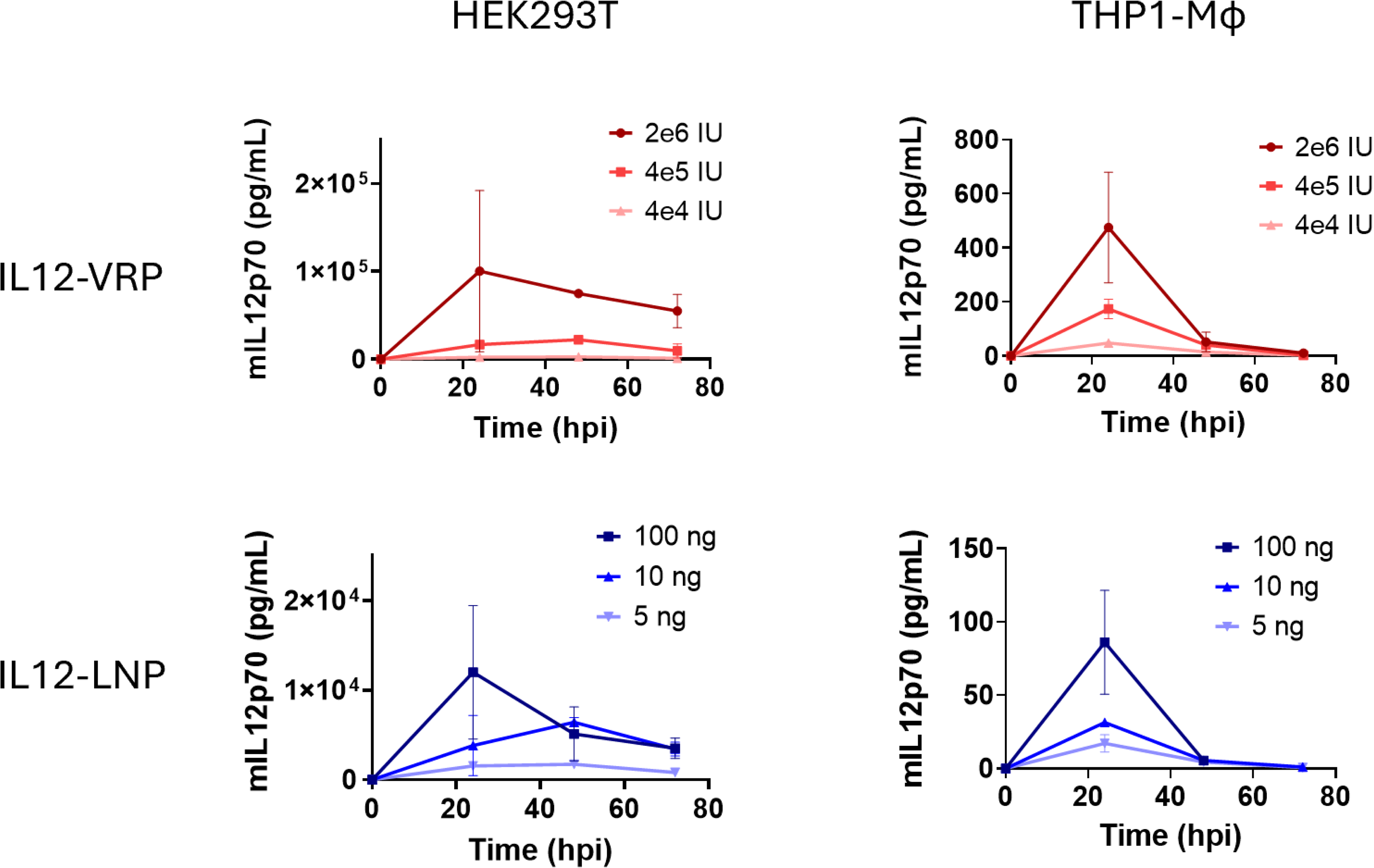
Dose dependent transduction by IL12-VRP and IL12-LNP *in vitro.* Human HEK293T (Left panels) or human THP-1 (Right panels), polarized into M2-like macrophages, were treated with either IL12-VRP (Upper panels) or IL12-LNP (Lower panels) at several dose levels. At the indicated timepoints, cell culture supernatant was collected and mIL-12p70 amount was measured by ELISA. Each timepoint was tested in triplicates.

**Extended Data Figure 2.**
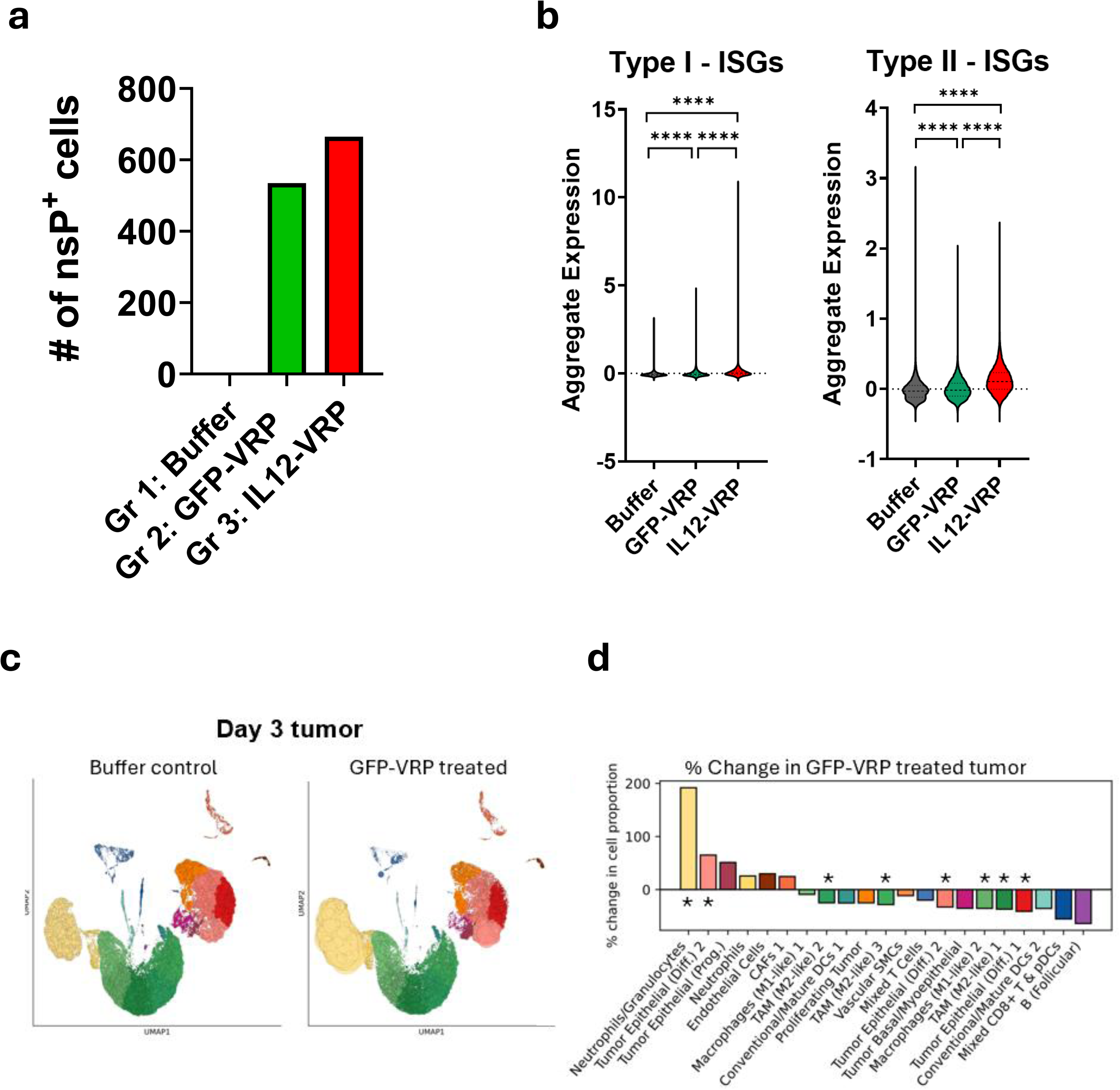
**a,** Absolute number of cells harboring nsP transcripts in each treatment group. Pooled datasets from tumors harvested day 1 and tumors/D-LNs harvested day 3 post treatment are tested. **b,** Expression levels of type I interferon stimulated genes (ISGs) and type II ISGs in the cells from tumors harvested day 3 post treatment. Asterisk indicates statistical significance. ****, *p*<0.0001 by Mann–Whitney *U* test. **c,** Mapping score of cell types in day 3 tumor. Buffer control (left) and GFP-VRP treated (right). **d,** Proportional change in cell population upon GFP-VRP treatment, compared to buffer control. Asterisk indicates *p*<1e-20 by Chi-squared test.

**Extended Data Figure 3.**
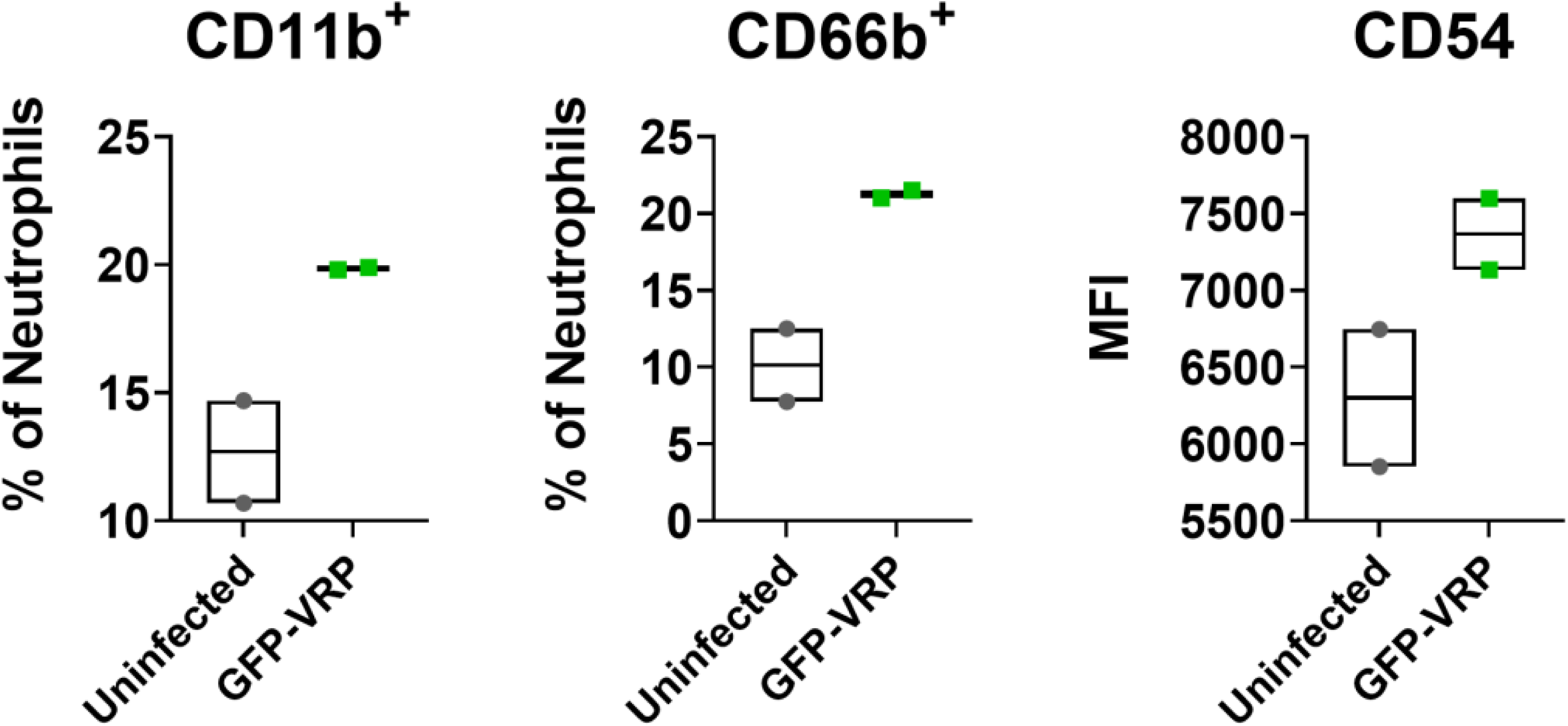
N2 to N1 shift by GFP-VRP treatment to human neutrophils *in vitro*. Human neutrophils were isolated and induced into N2-like status in N2 cocktail for 24 hours then transduced with GFP-VRP at MOI 150. The markers associated with N1-neutrophils (CD11b^+^; left panel, CD66b^+^; middle panel, and CD54; right panel) on N2 polarized neutrophils were then assessed 24 hours later by flow cytometry.

**Extended Data Figure 4.**
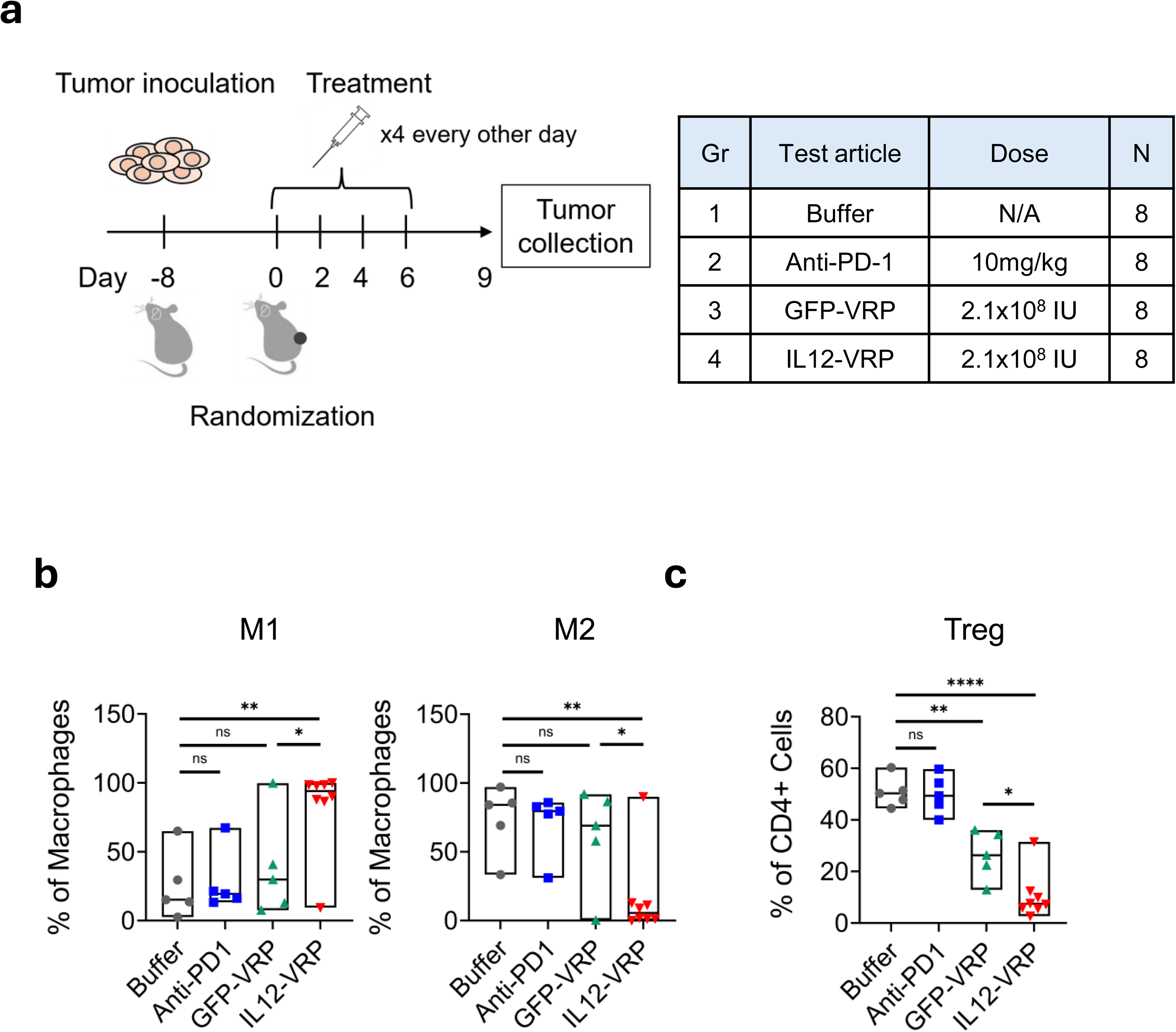
IL12-VRP reduces immune suppressive cells in TME in MC38 colorectal cancer model. **a,** study design using MC38 mouse syngeneic tumor model to assess changes in immune cells in the injected tumor. **b,** Flow cytometry analysis on macrophage populations in primary tumor. M1 macrophage population (%CD11b^+^ F4-80^+^ IA/IE^+^ from isolated CD45^+^ cells). M2 macrophage population (%CD11b^+^ F-80^+^ CD206^+^ from isolated CD45^+^ cells). **c,** Flow cytometry analysis on regulatory T cell (Treg) population (% FoxP3^+^ from isolated CD45^+^ CD11b^-^ CD3^+^ CD4^+^ cells) in primary tumor. *p* value indicates statistical significance. *, *p*<0.05; **, *p*<0.01; ****, *p*<0.0001; ns, not significant by *t*-test.

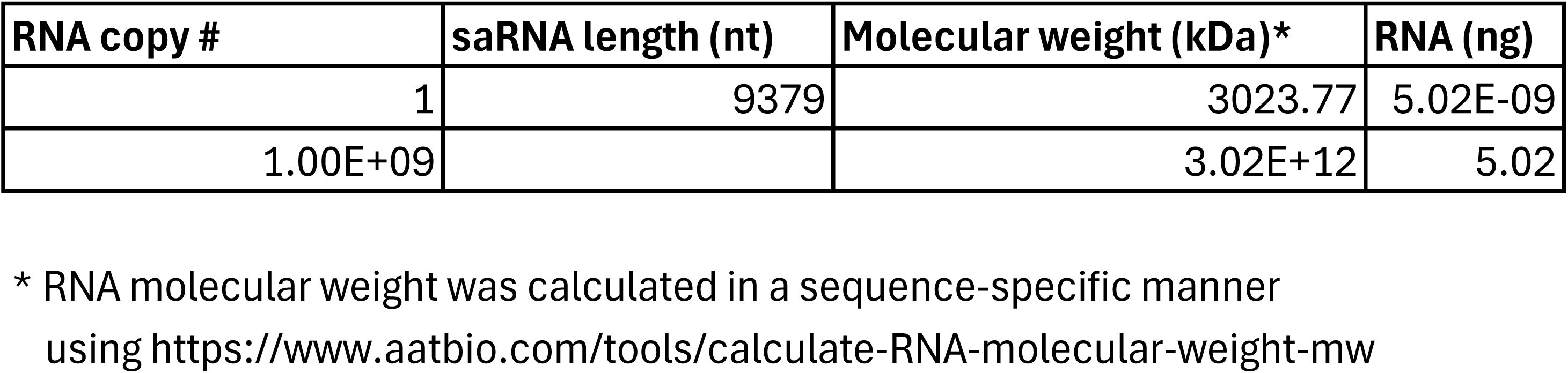
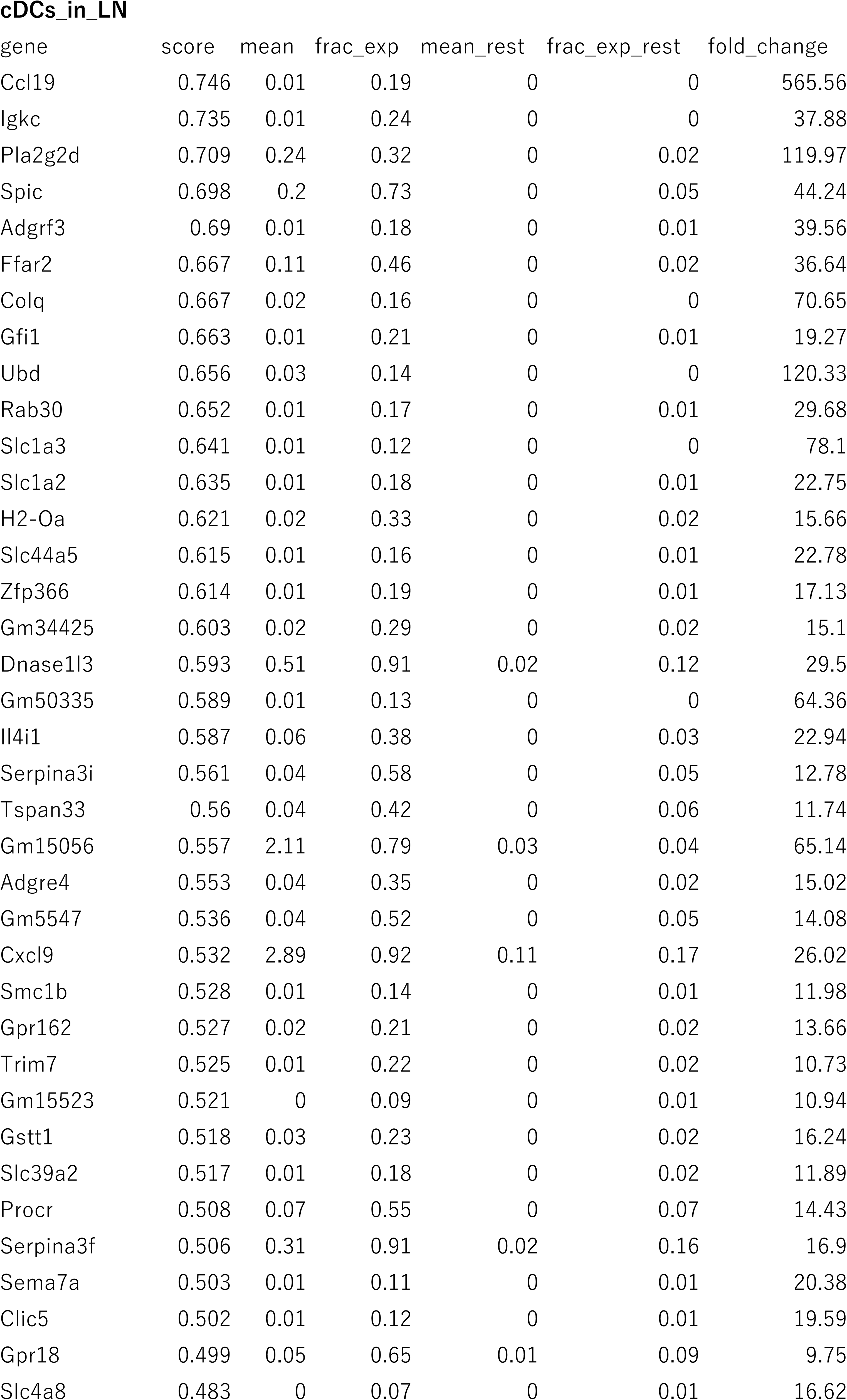
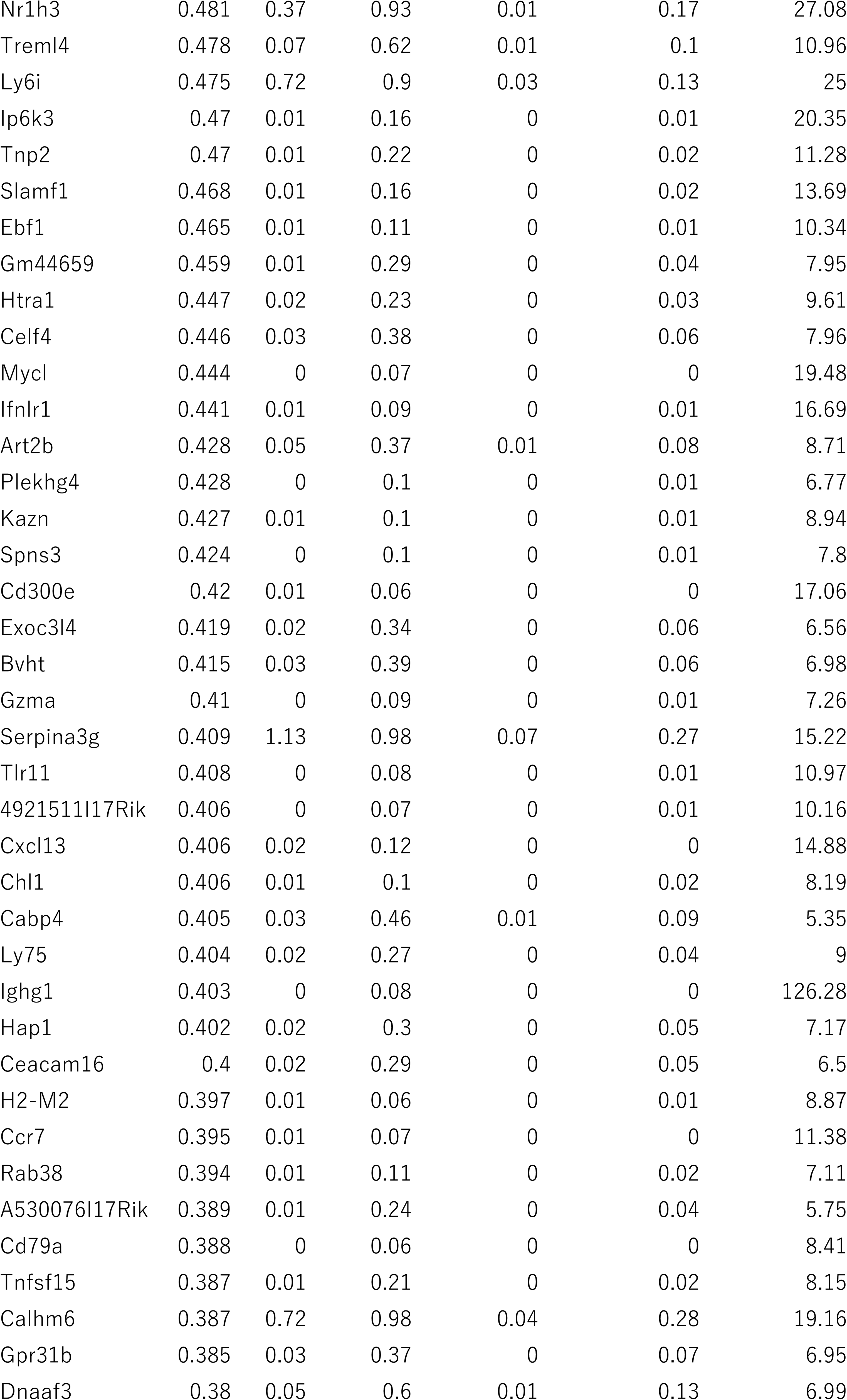
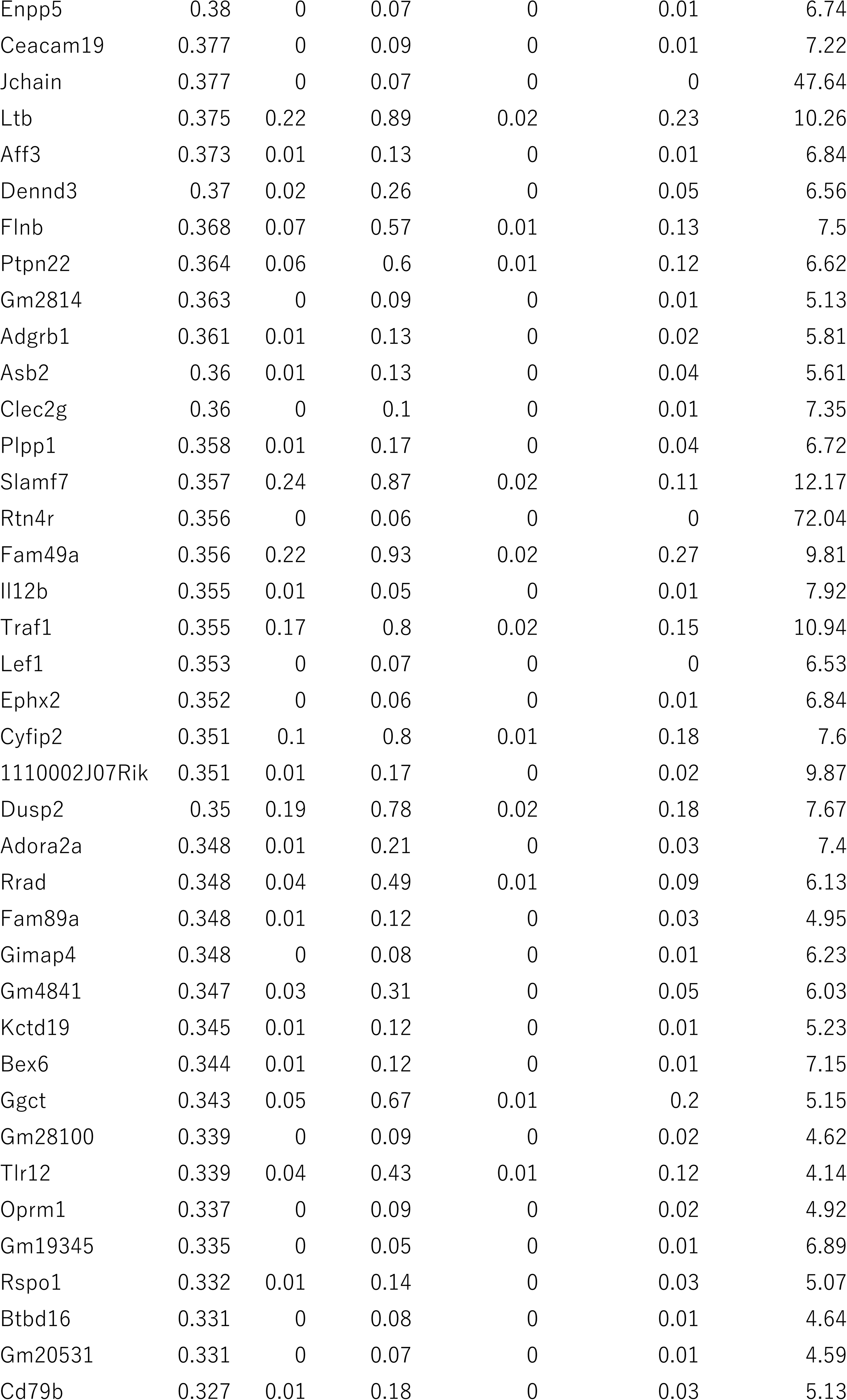
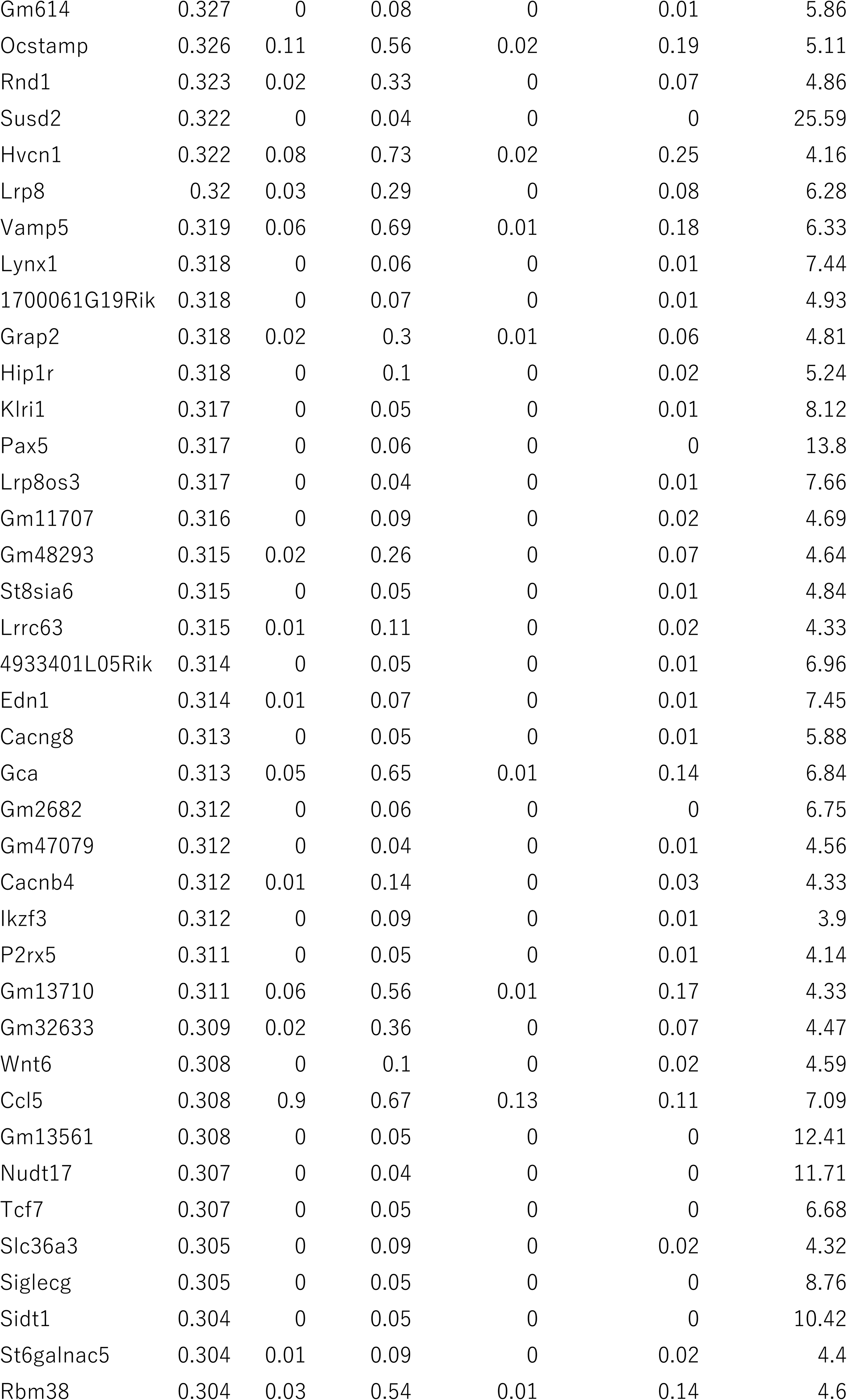
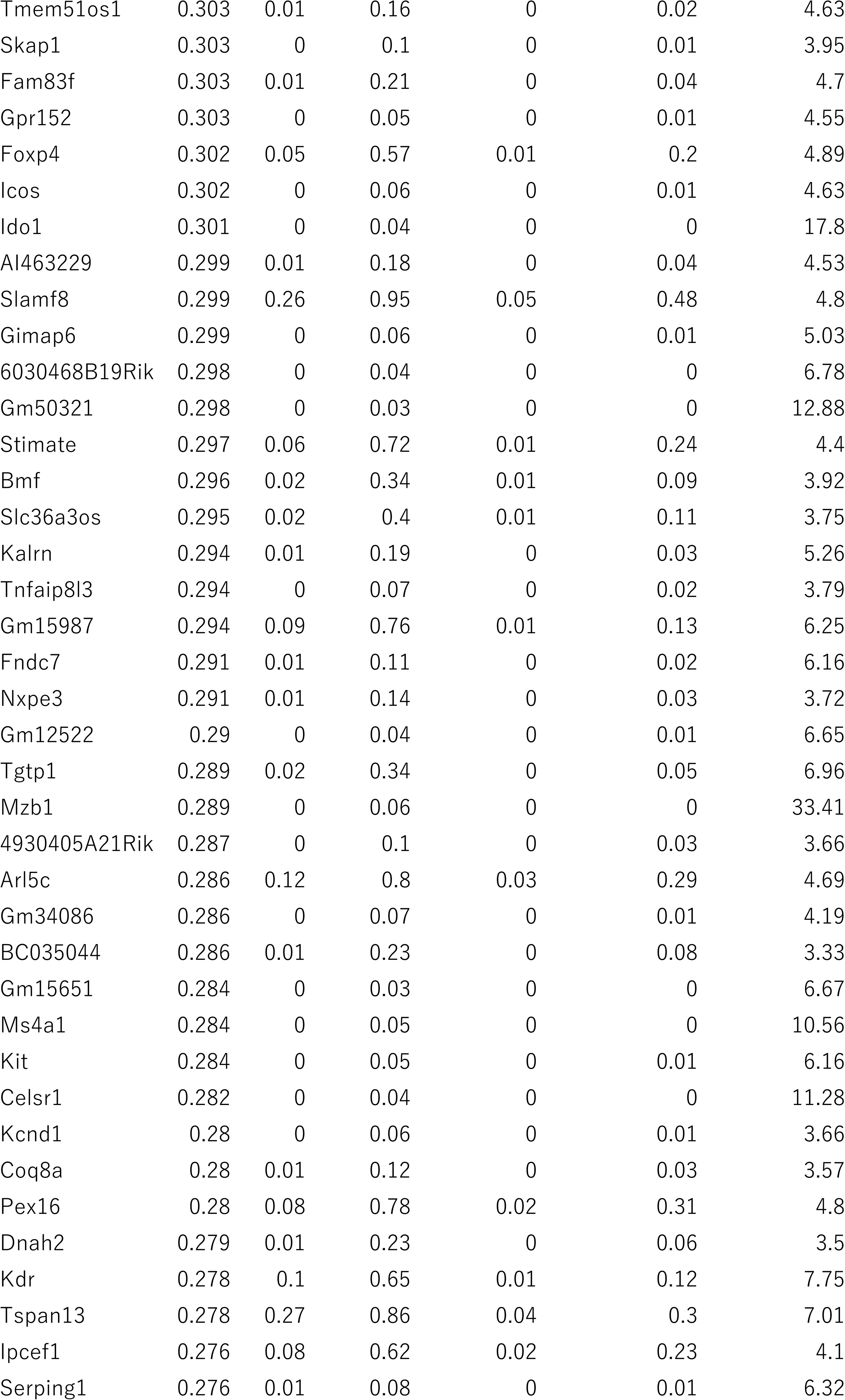
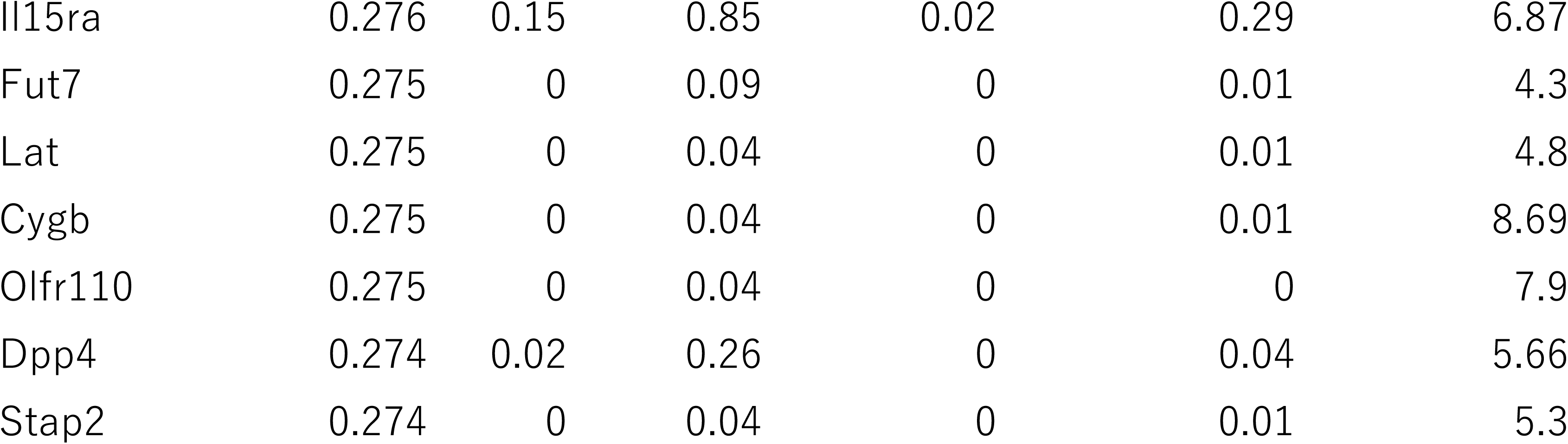
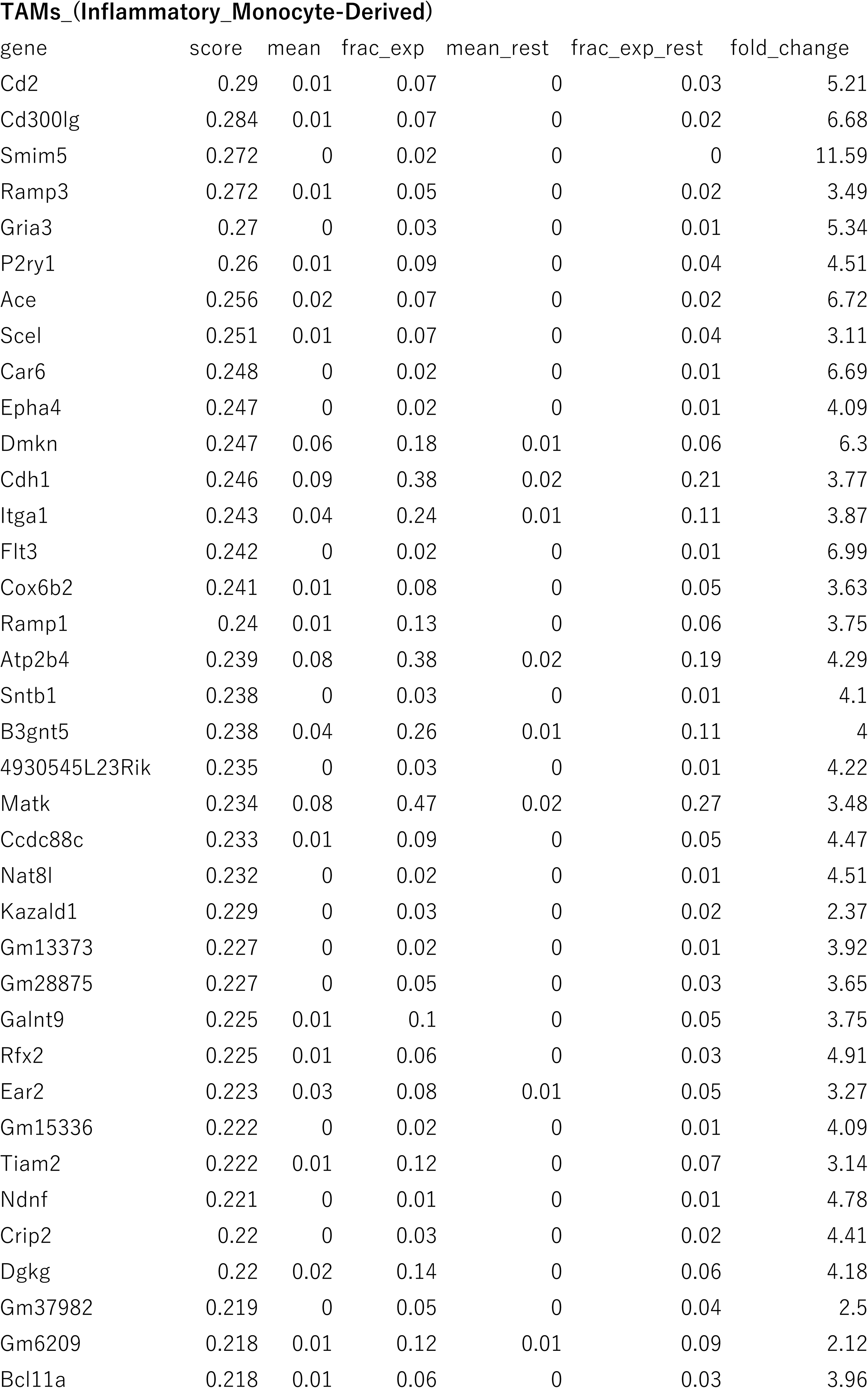
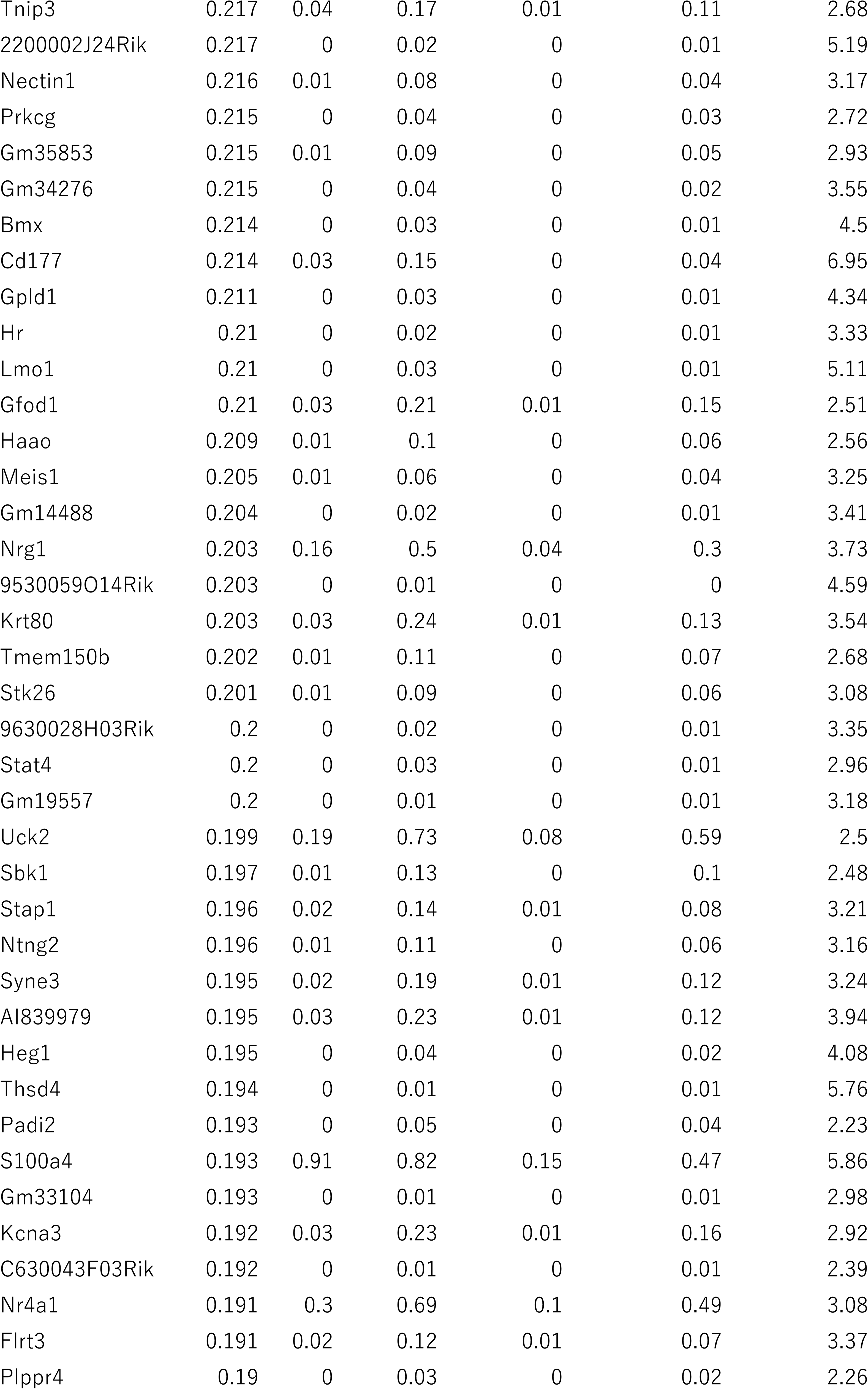
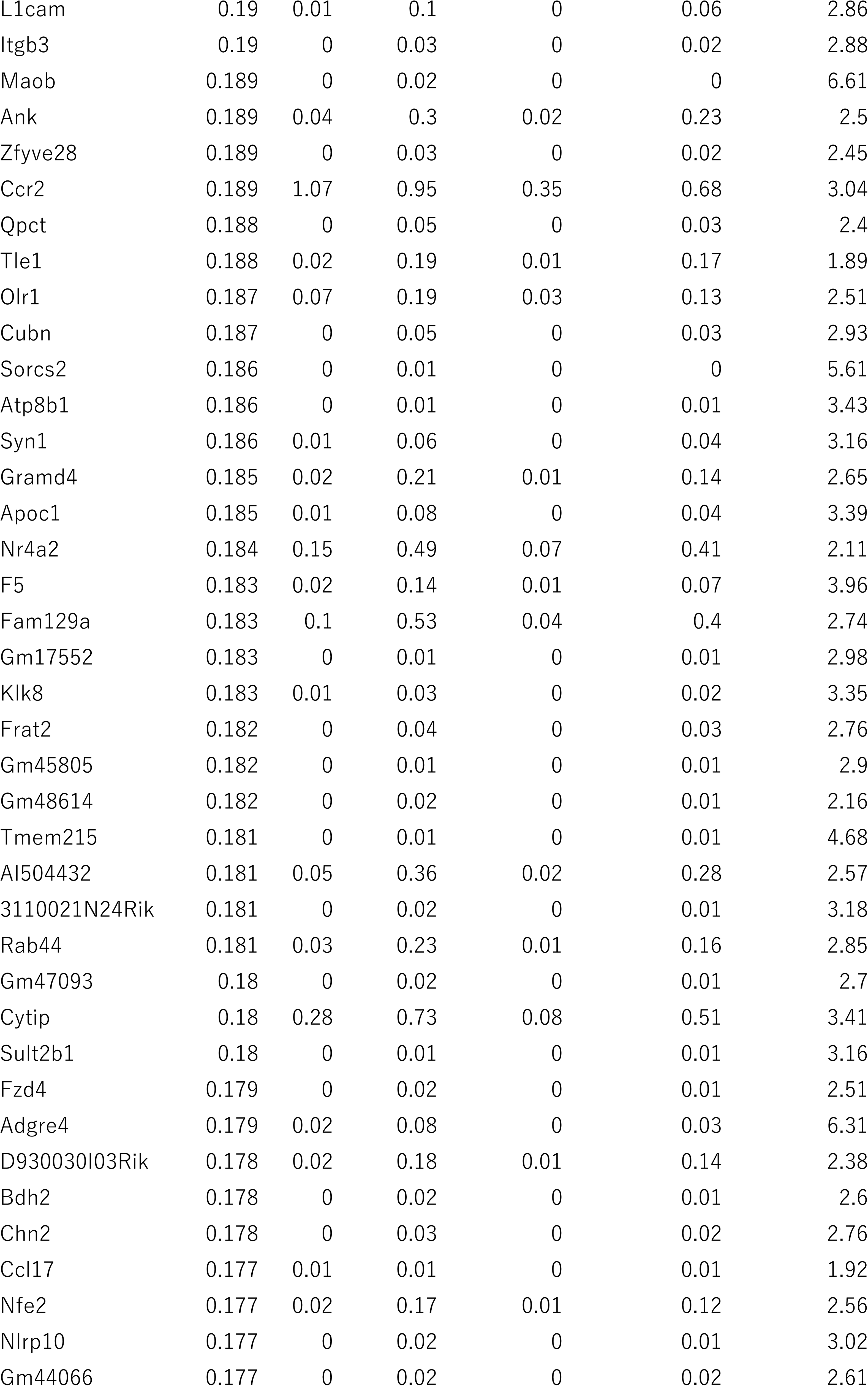
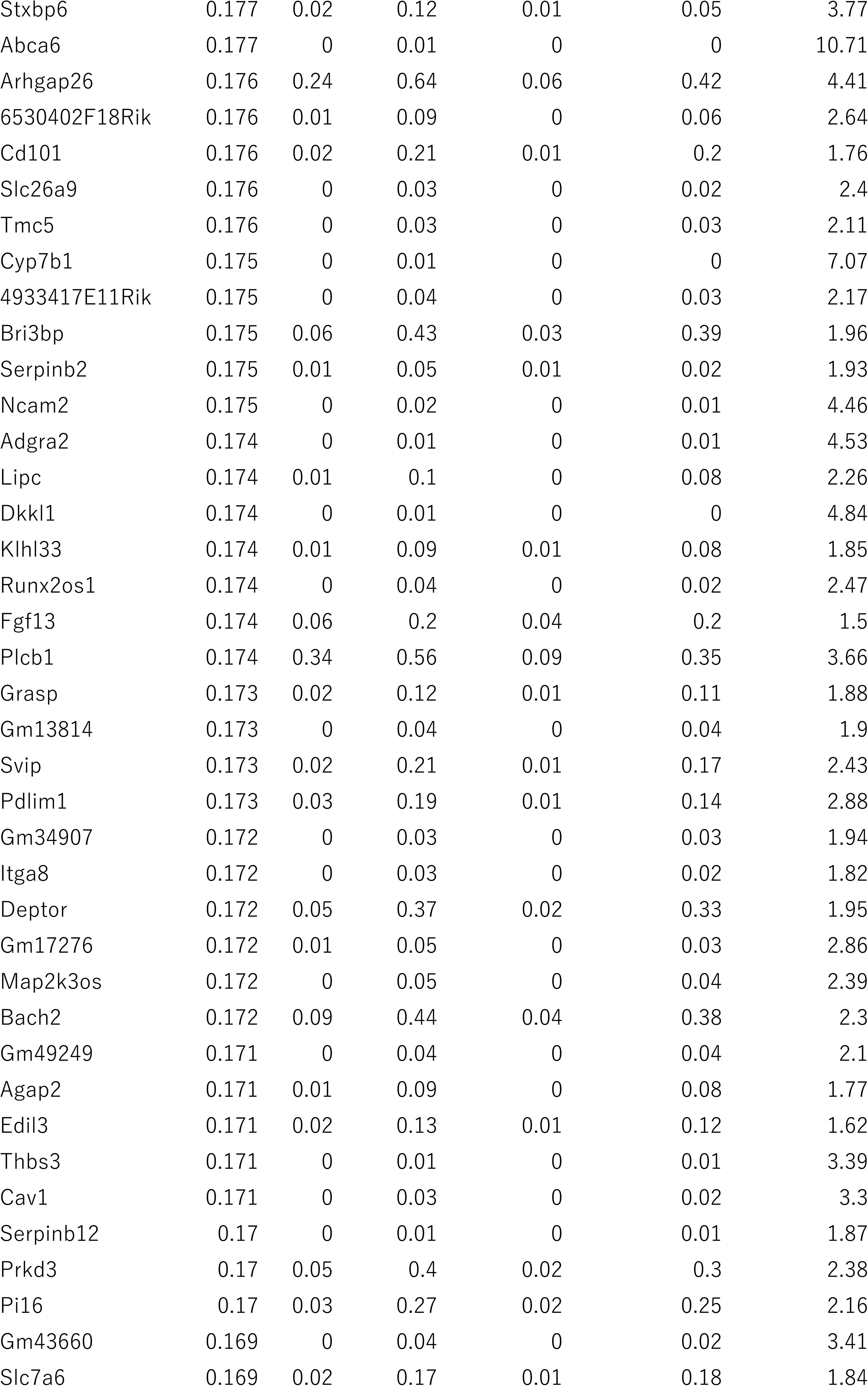
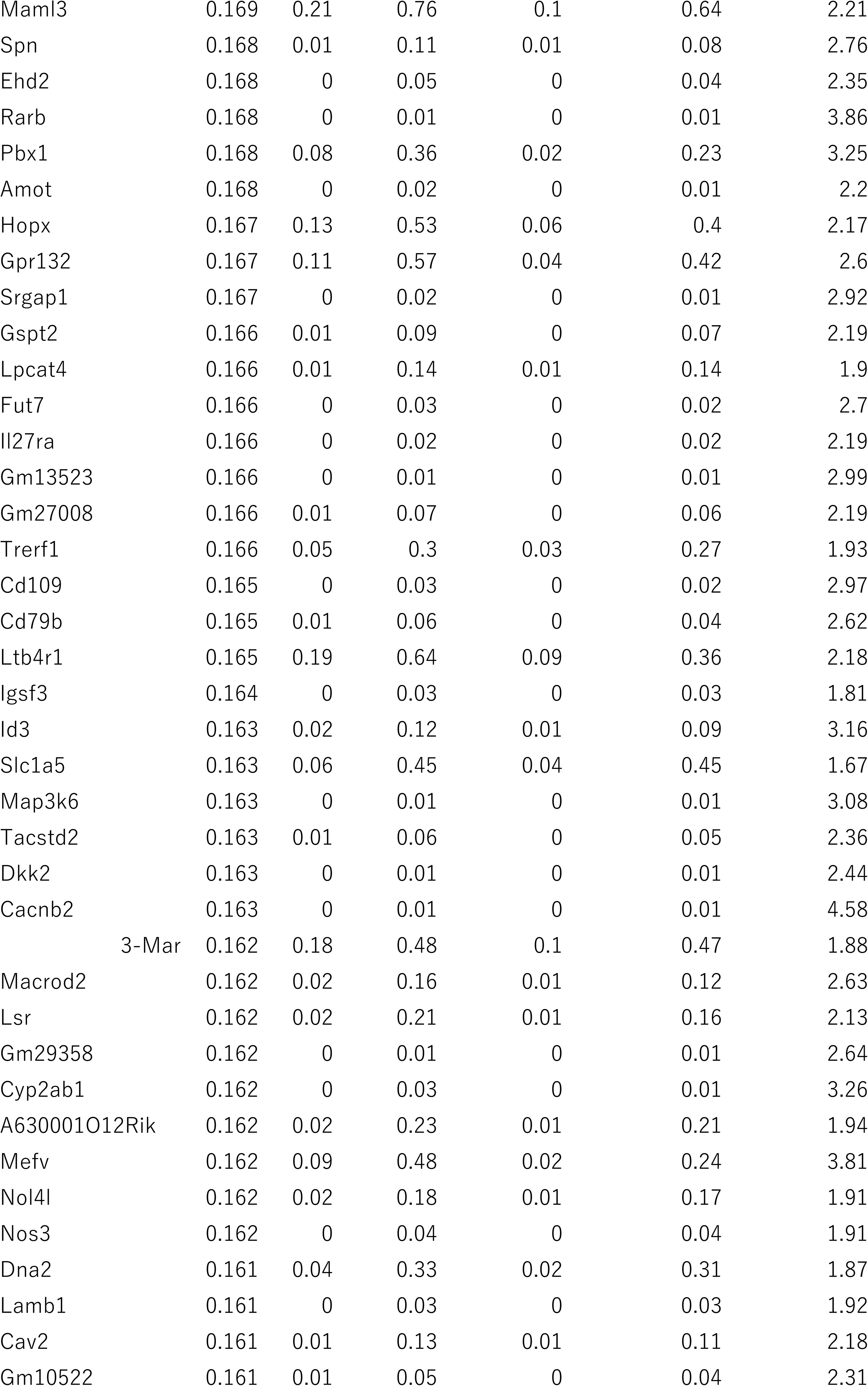
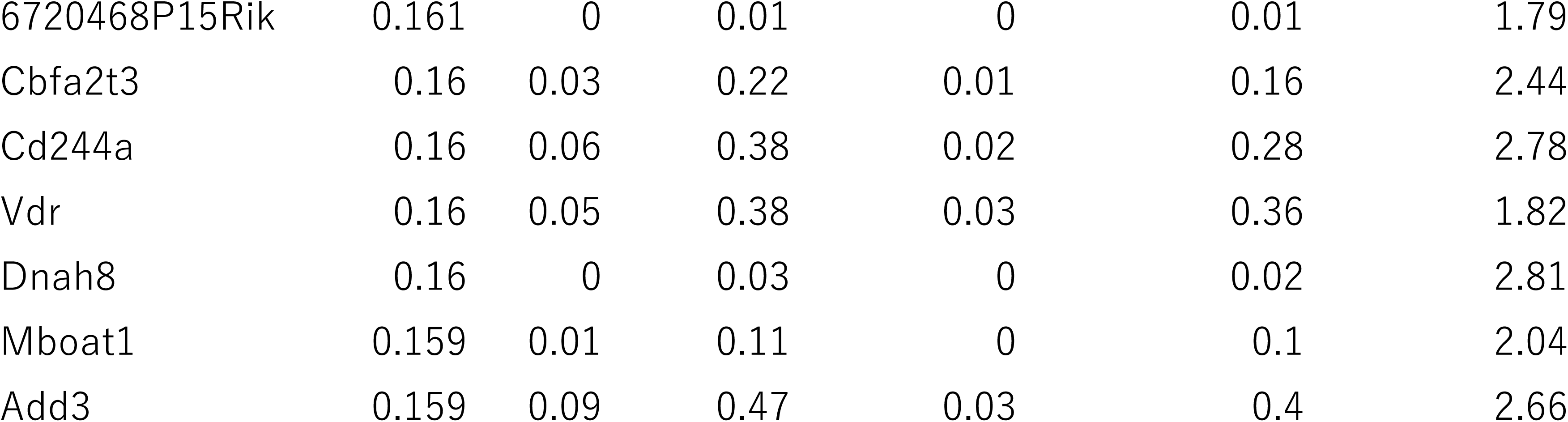
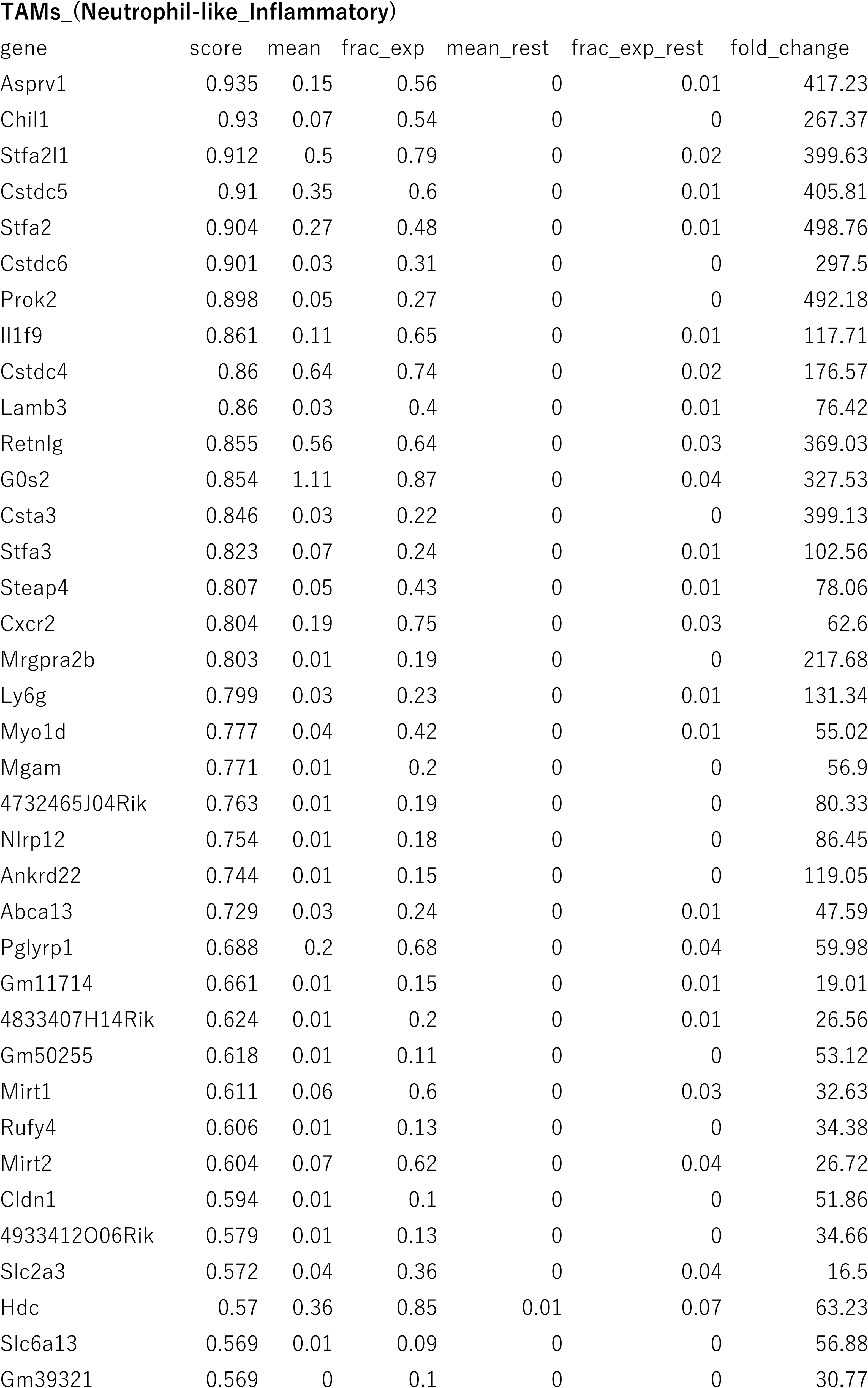
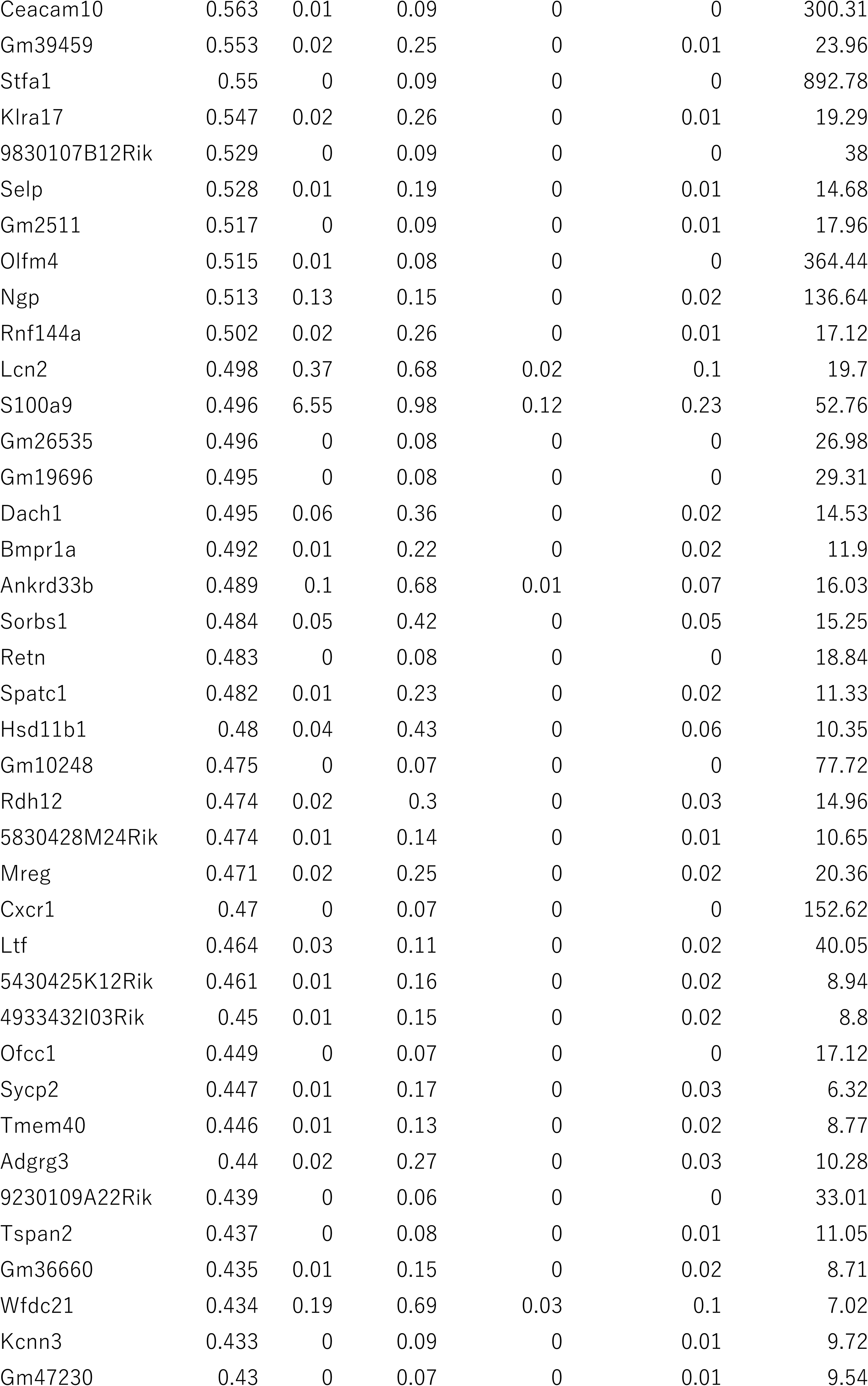
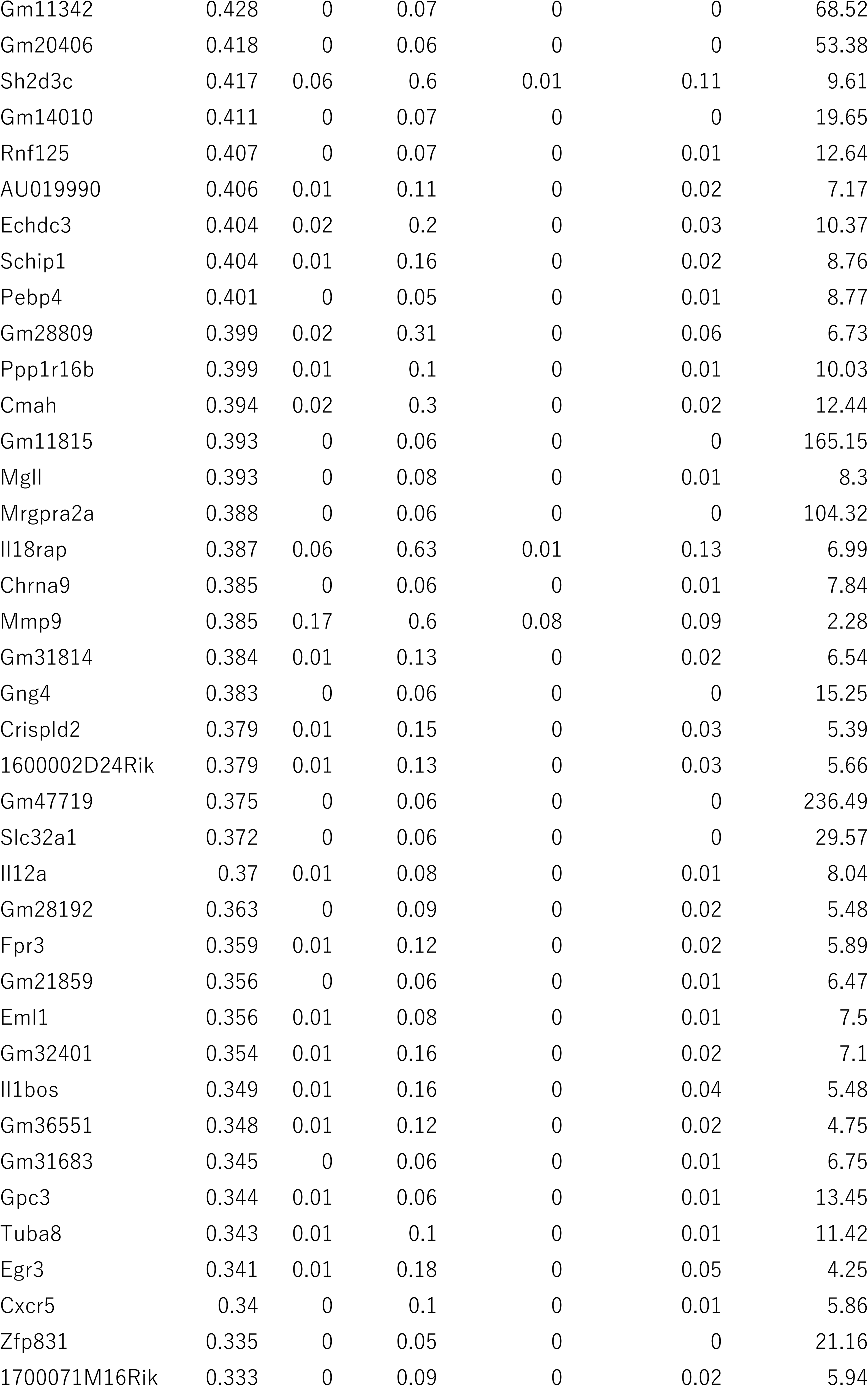
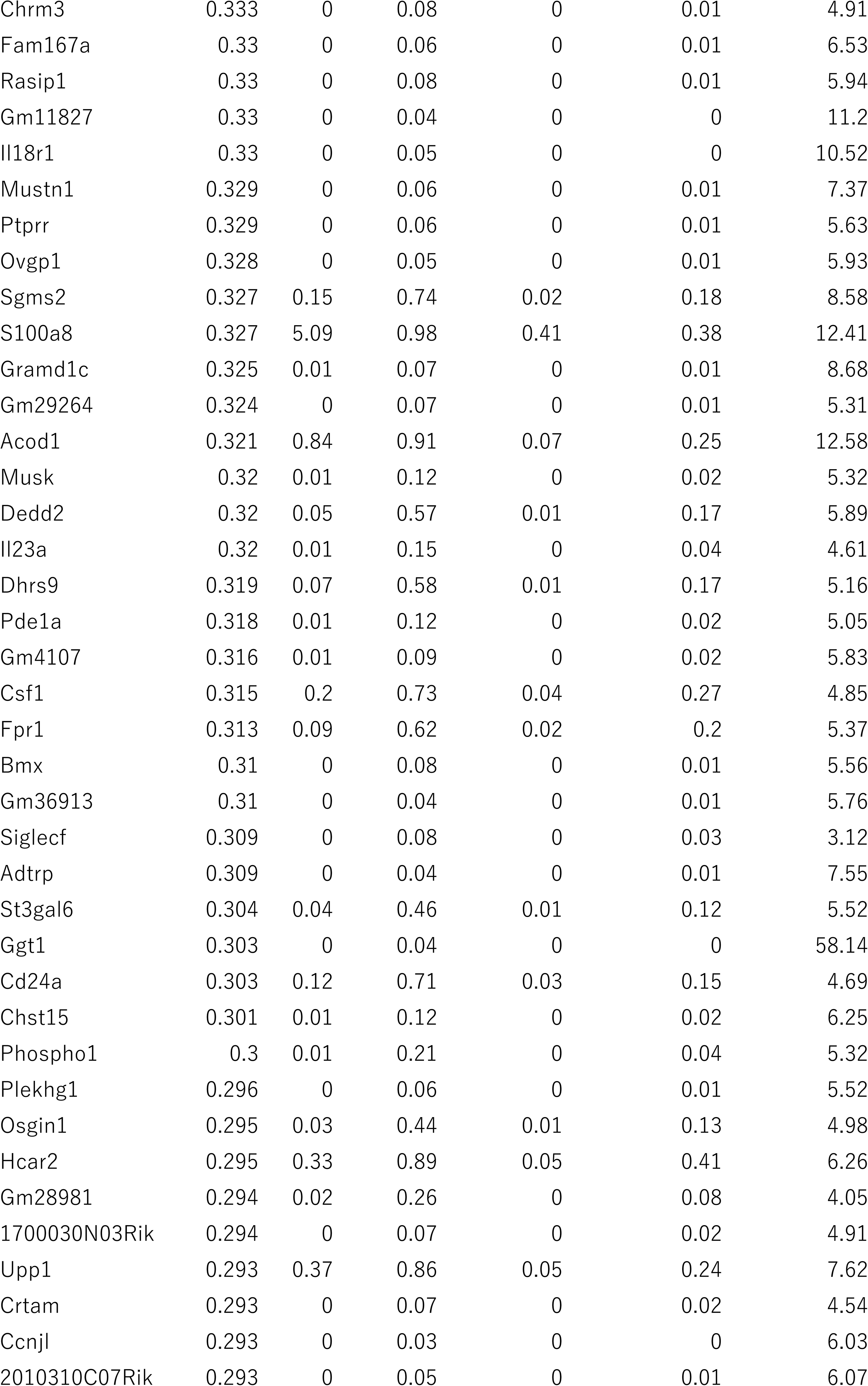
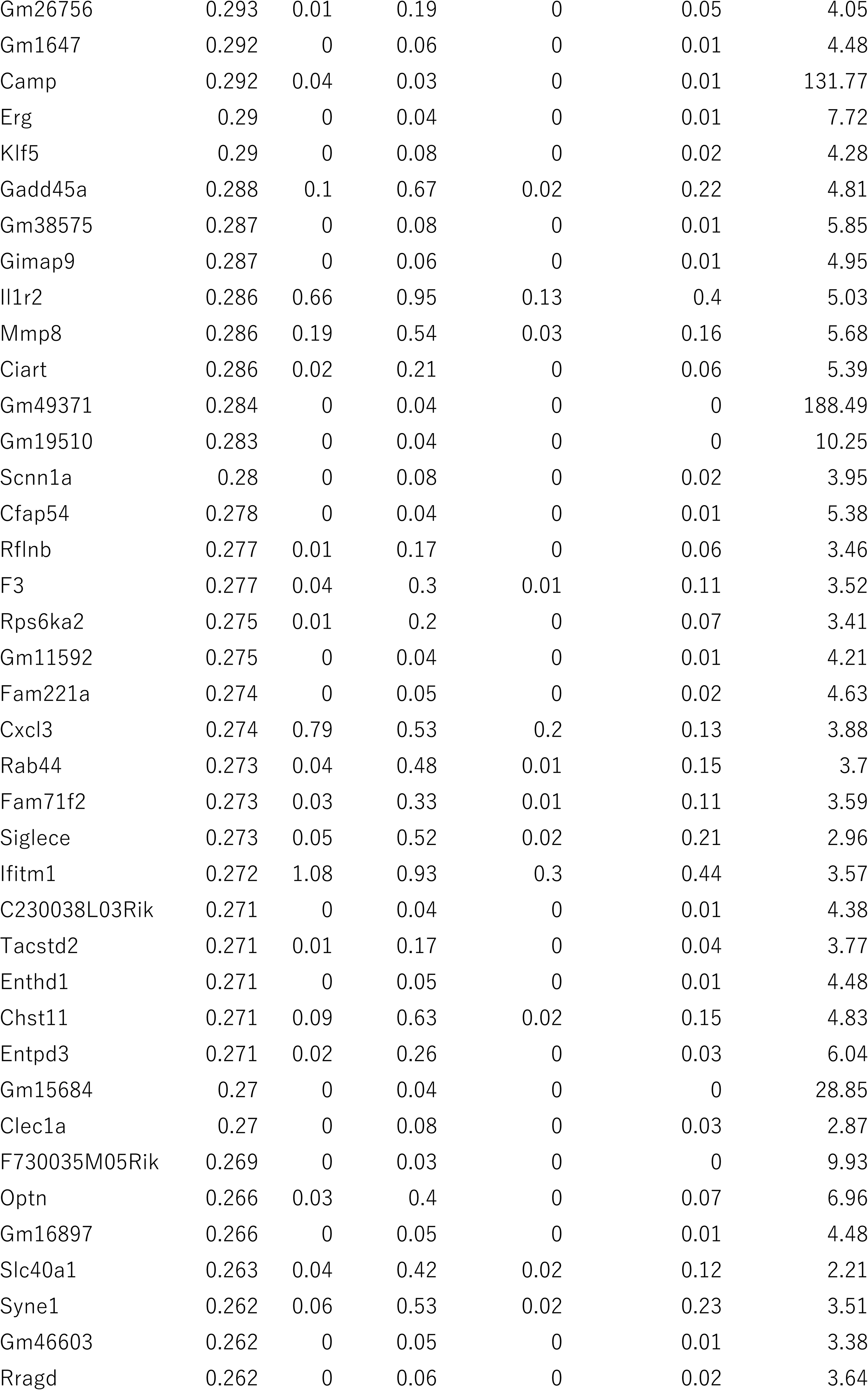
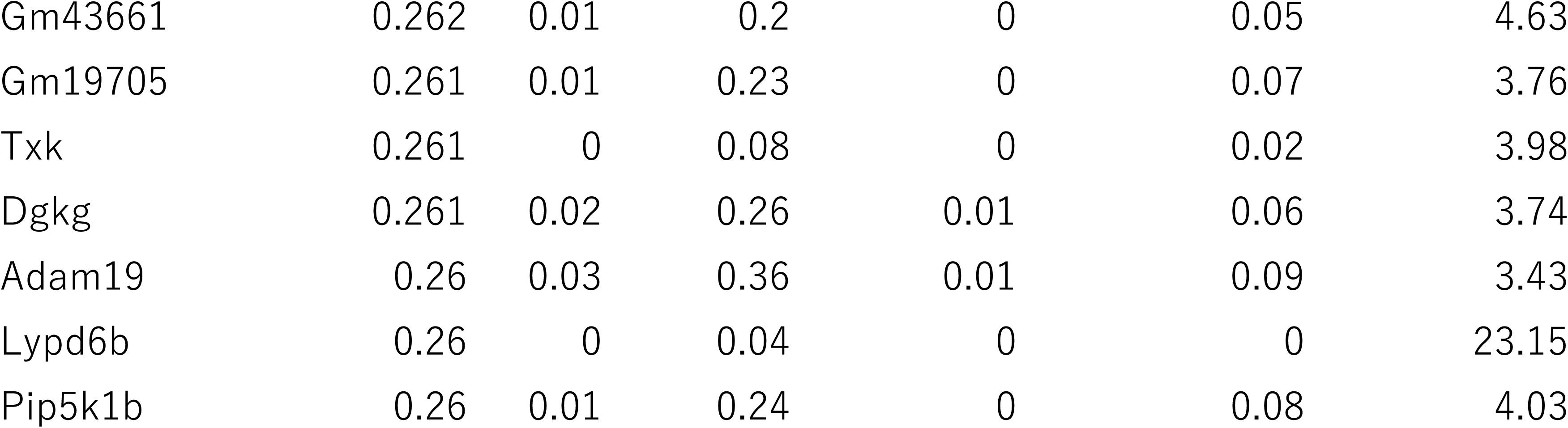
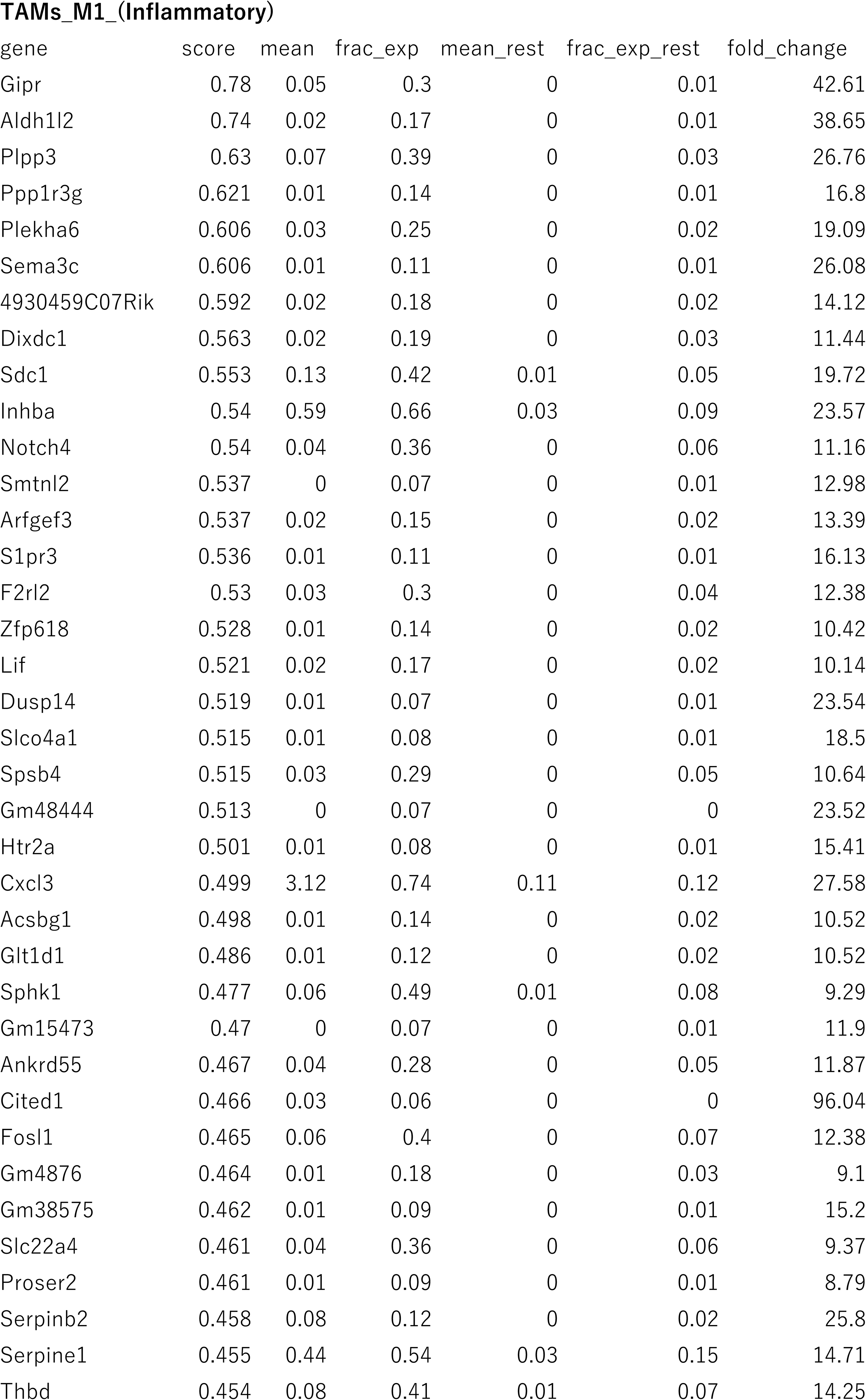
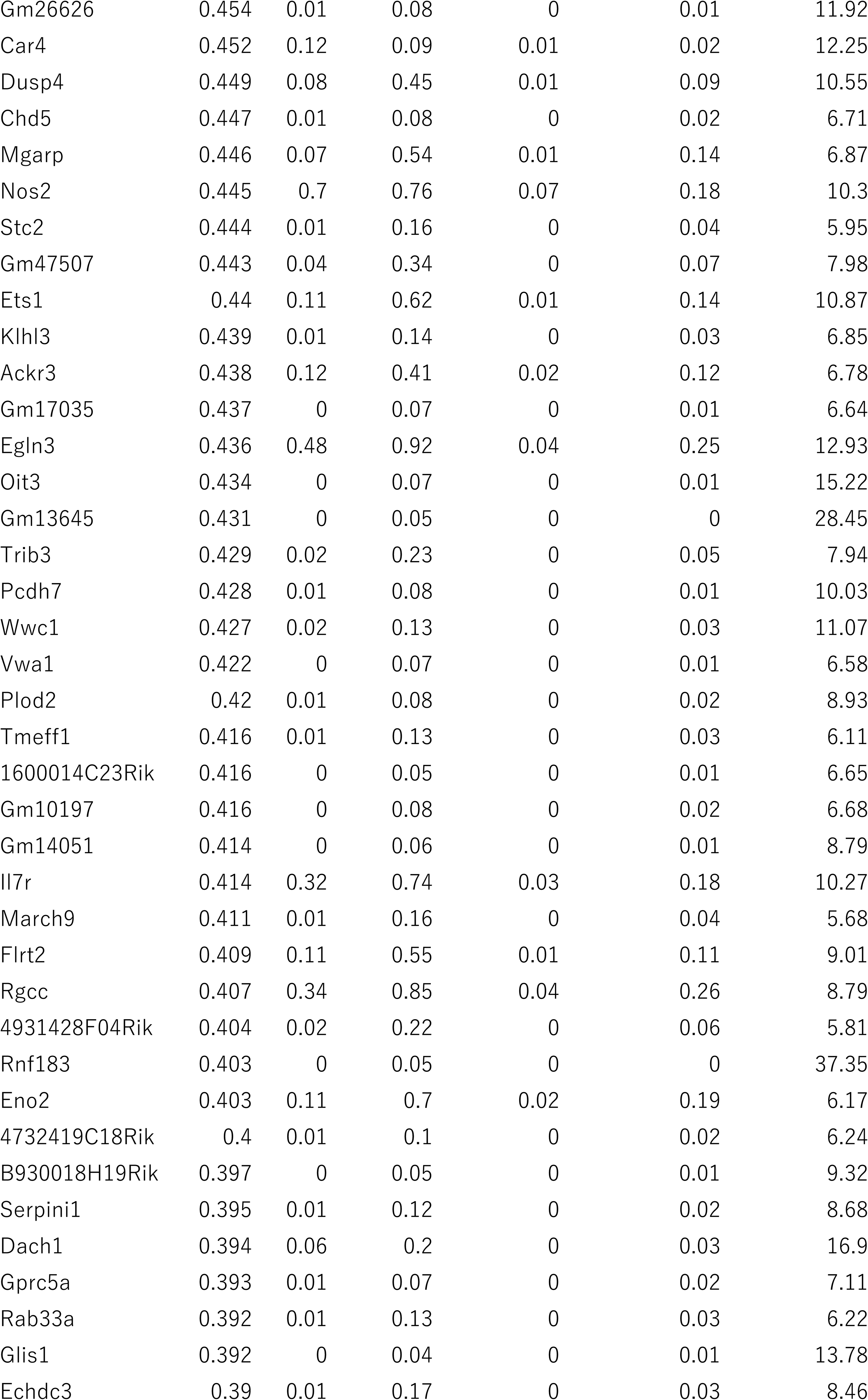
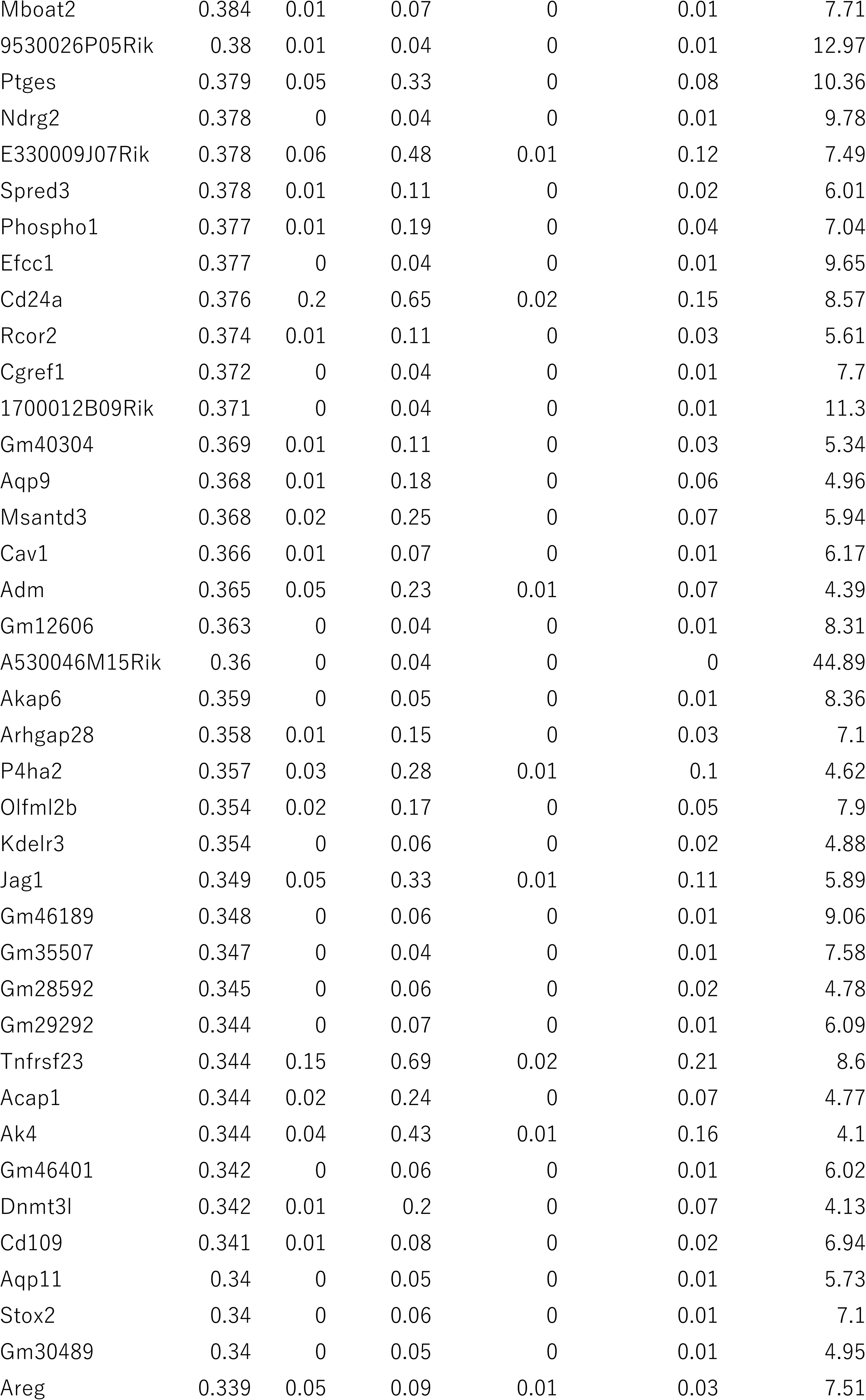
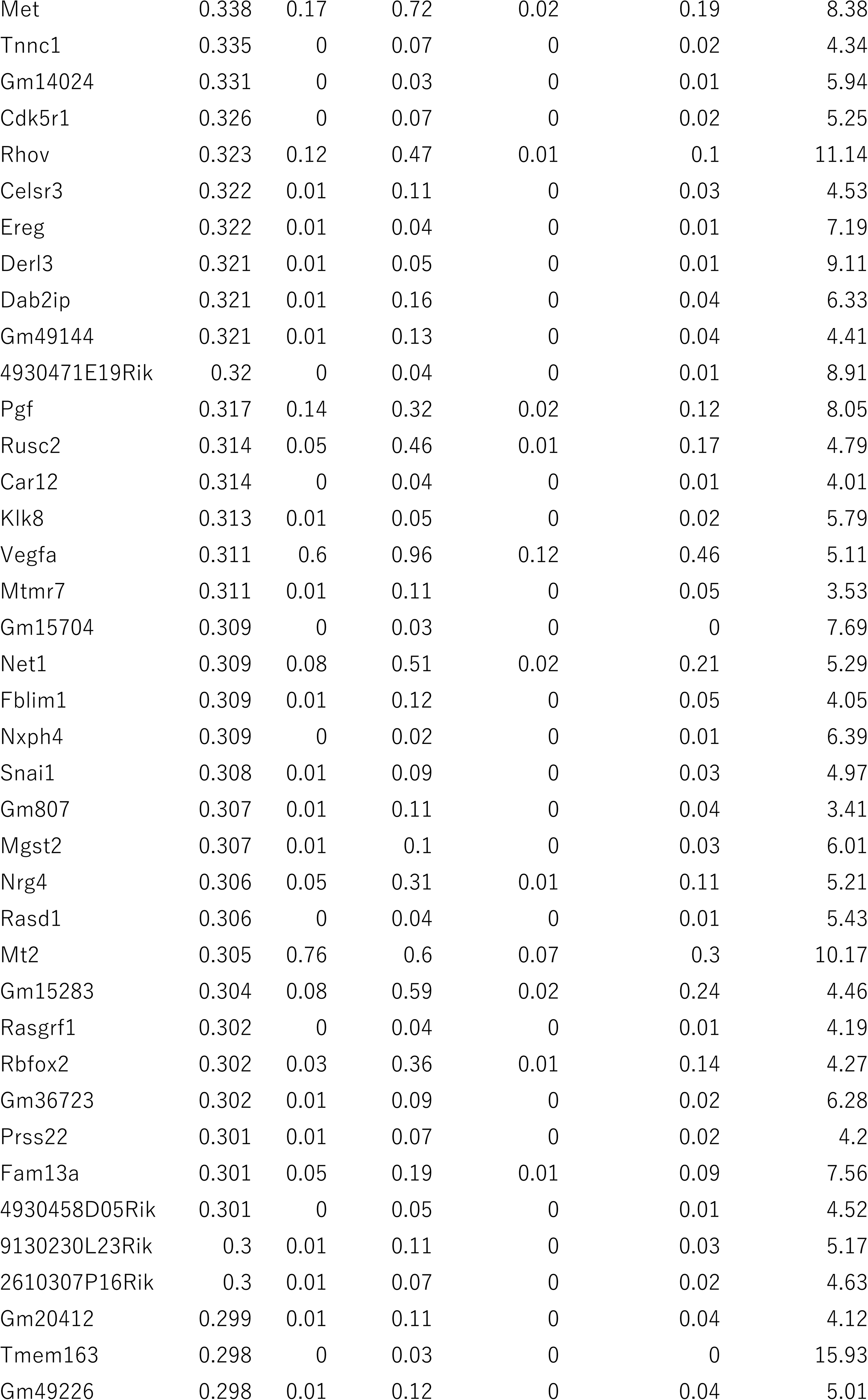
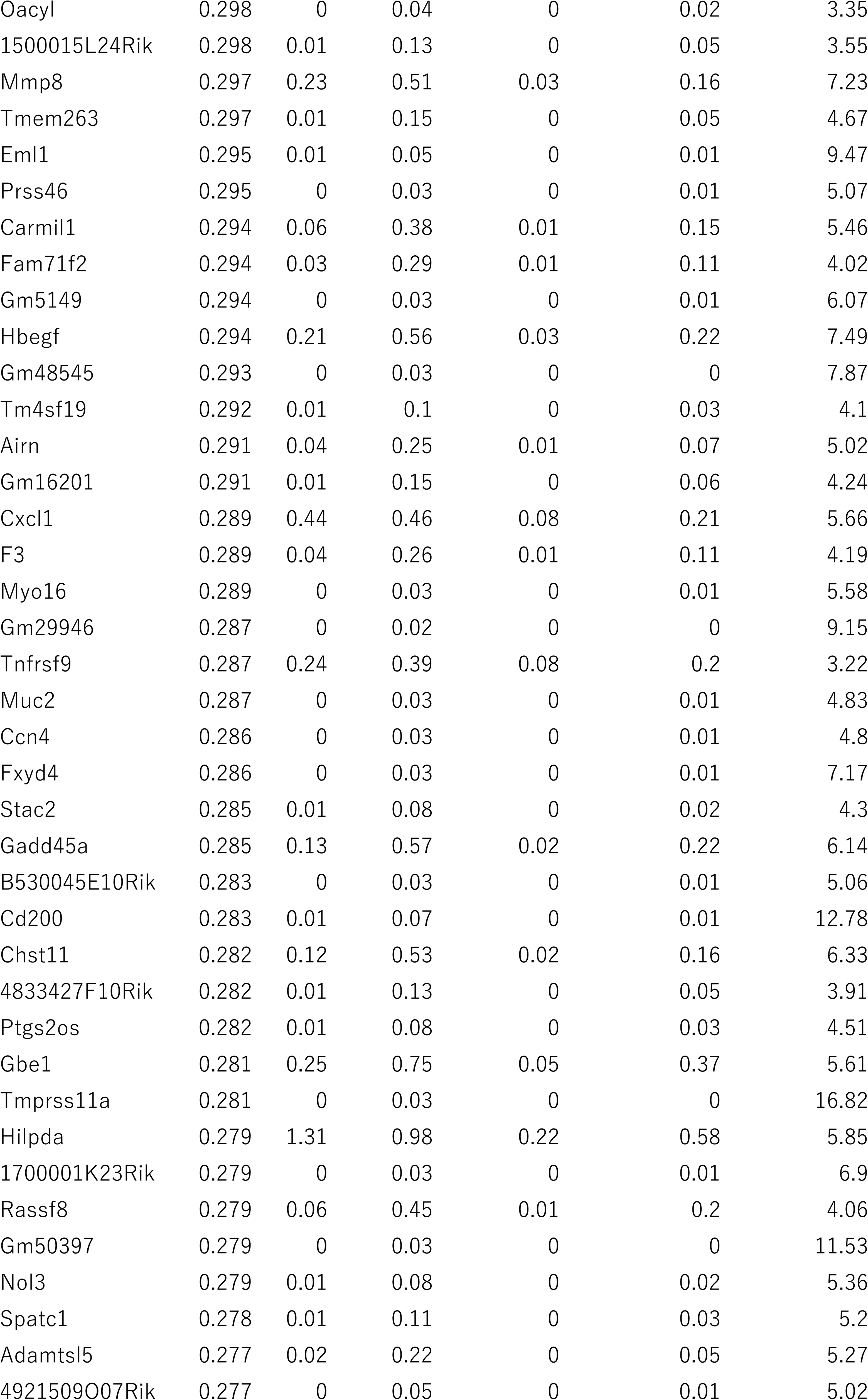
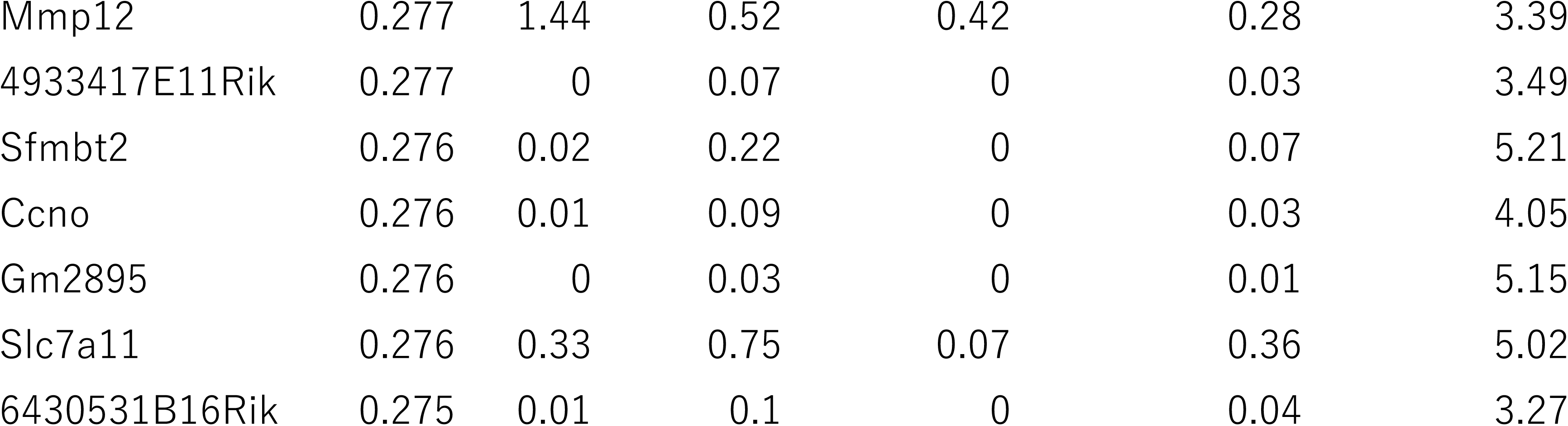
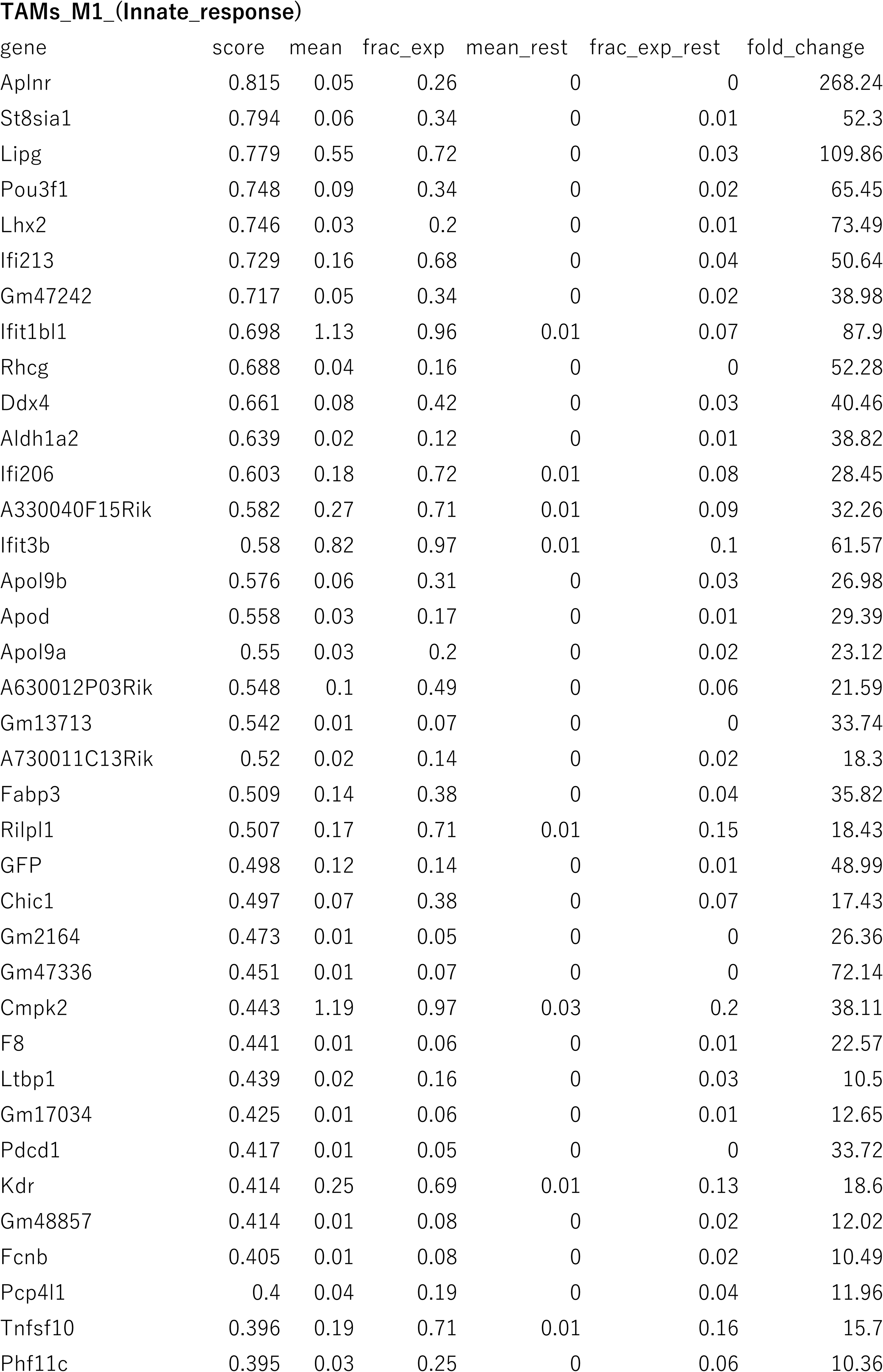
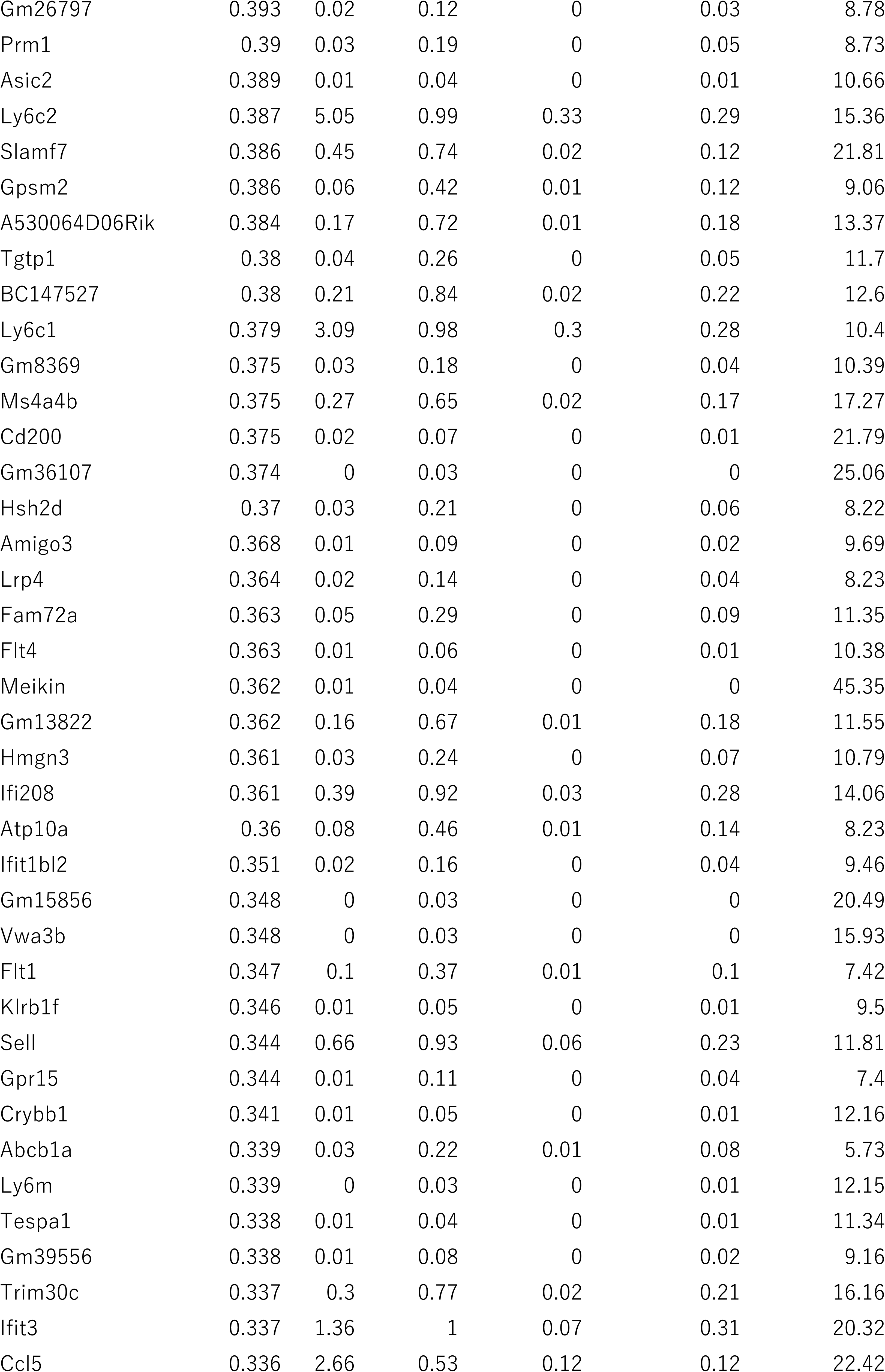
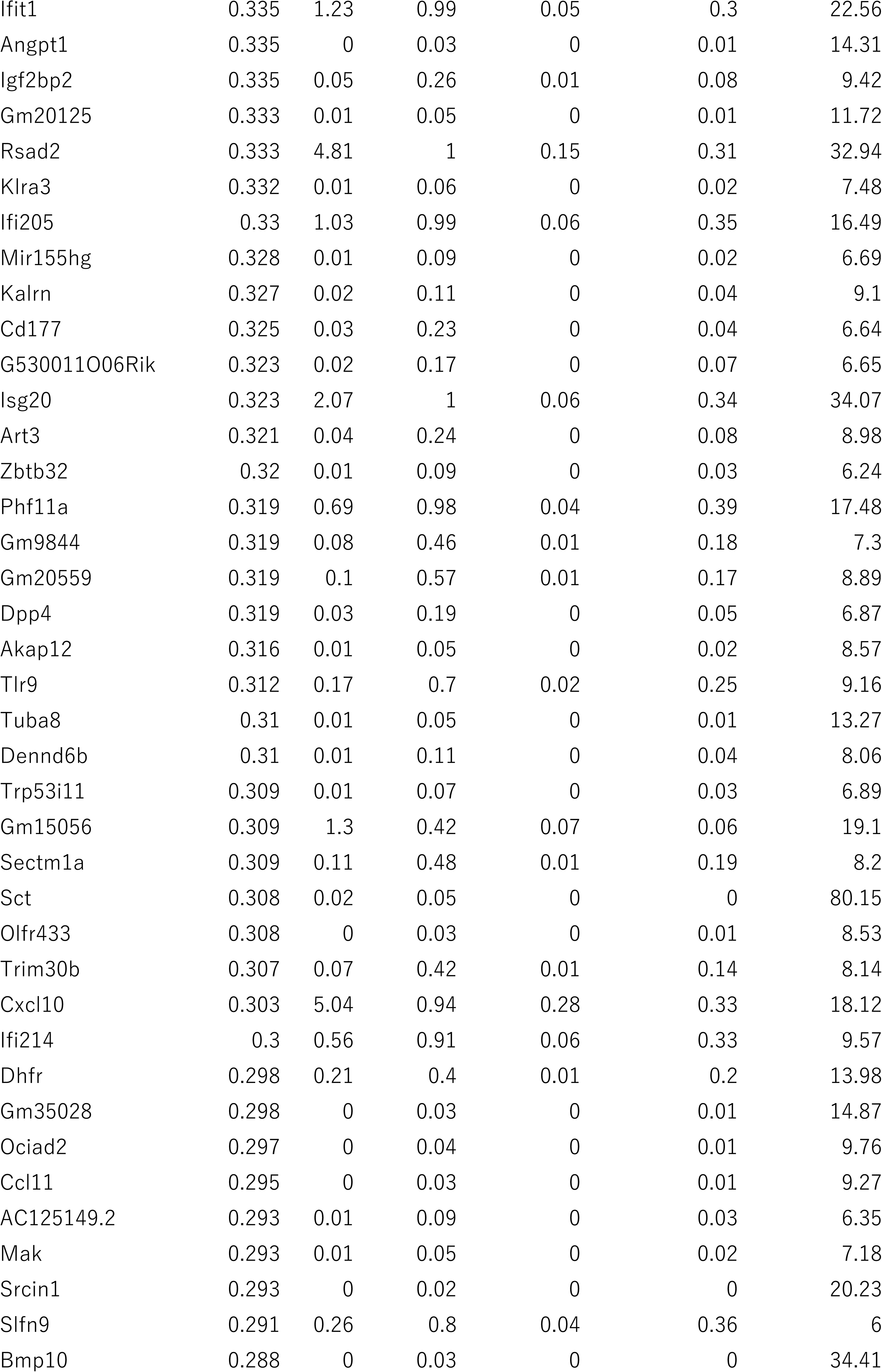
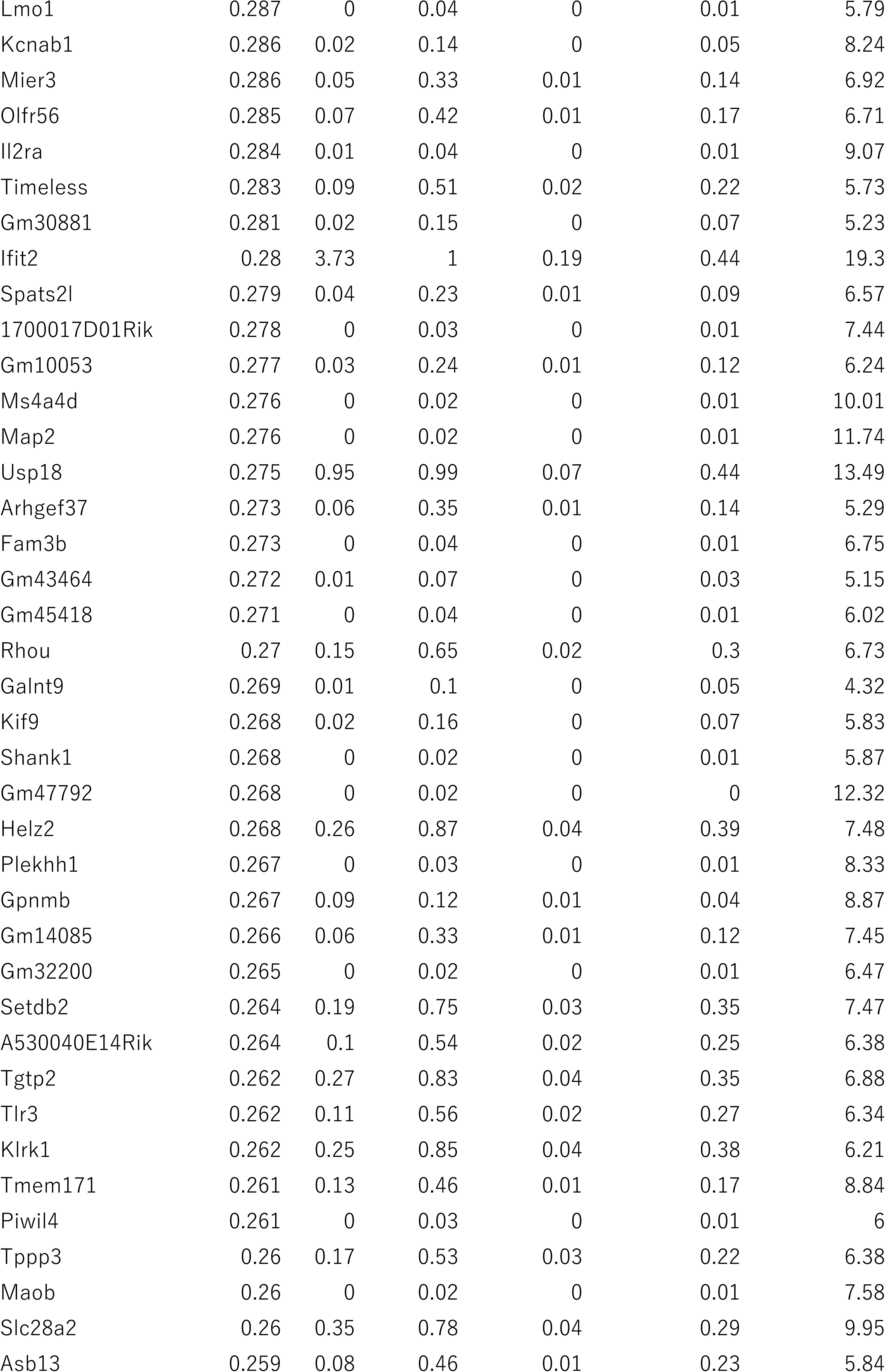
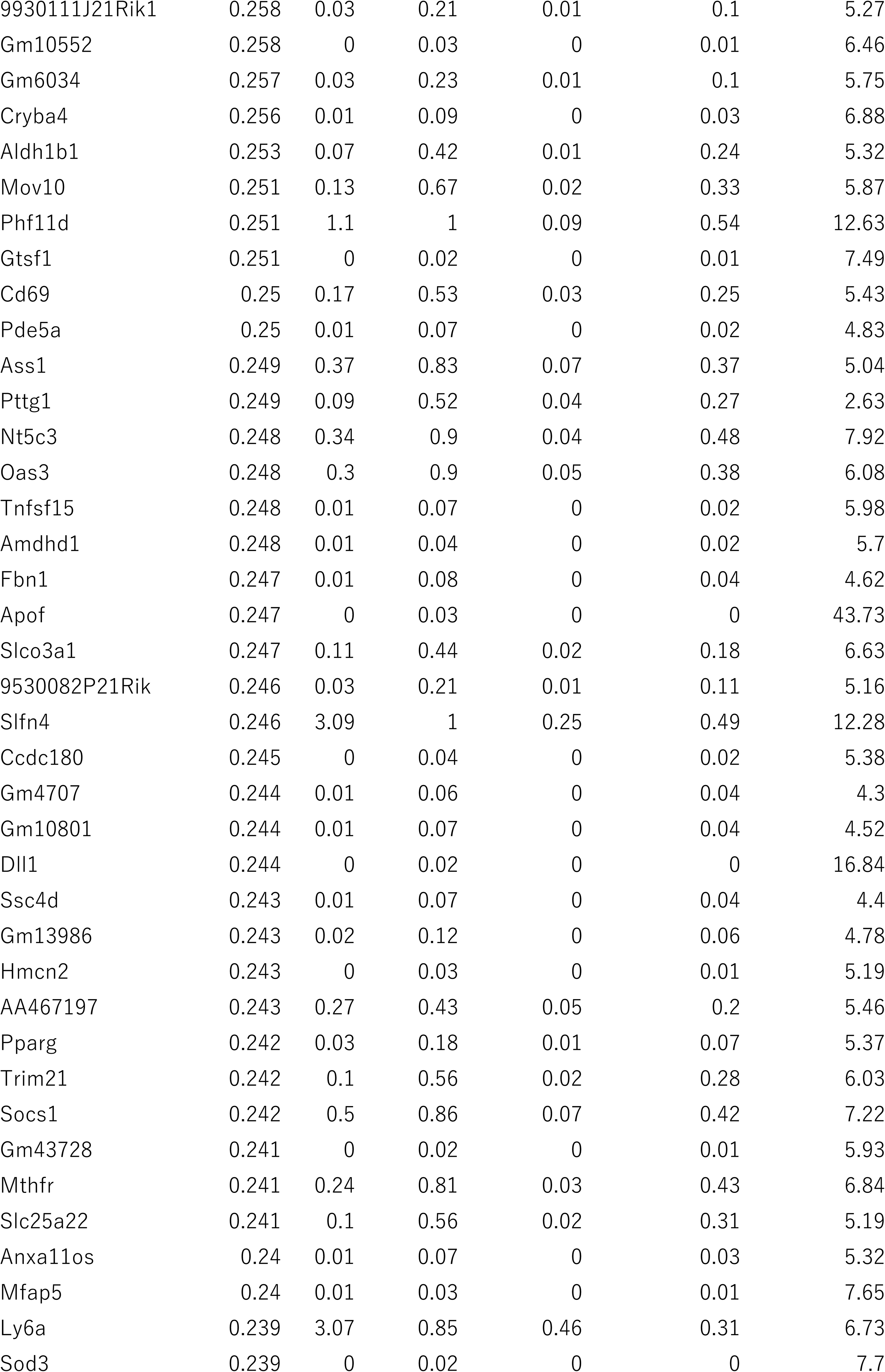
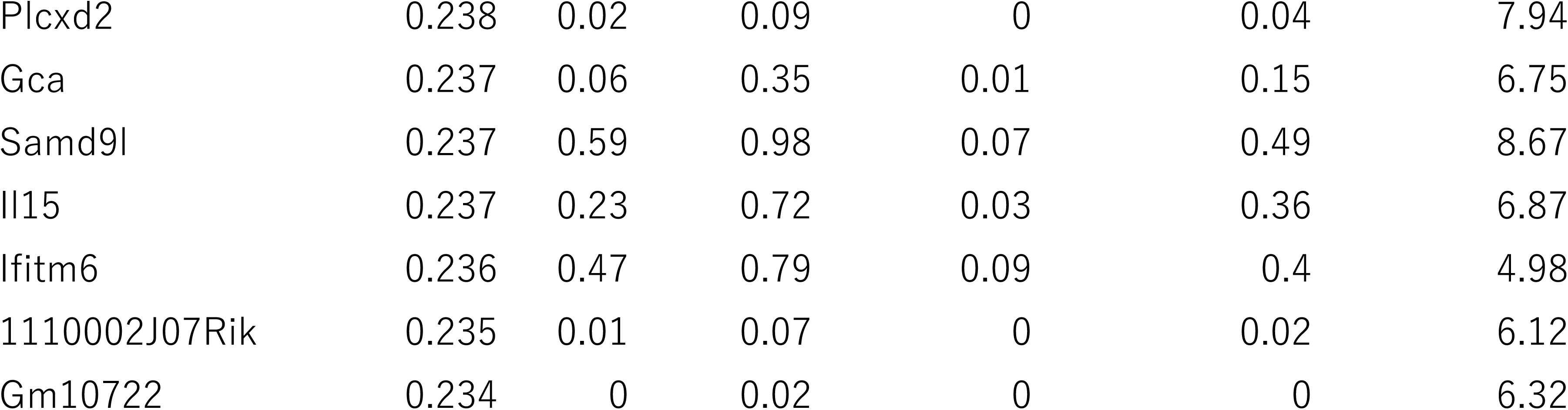
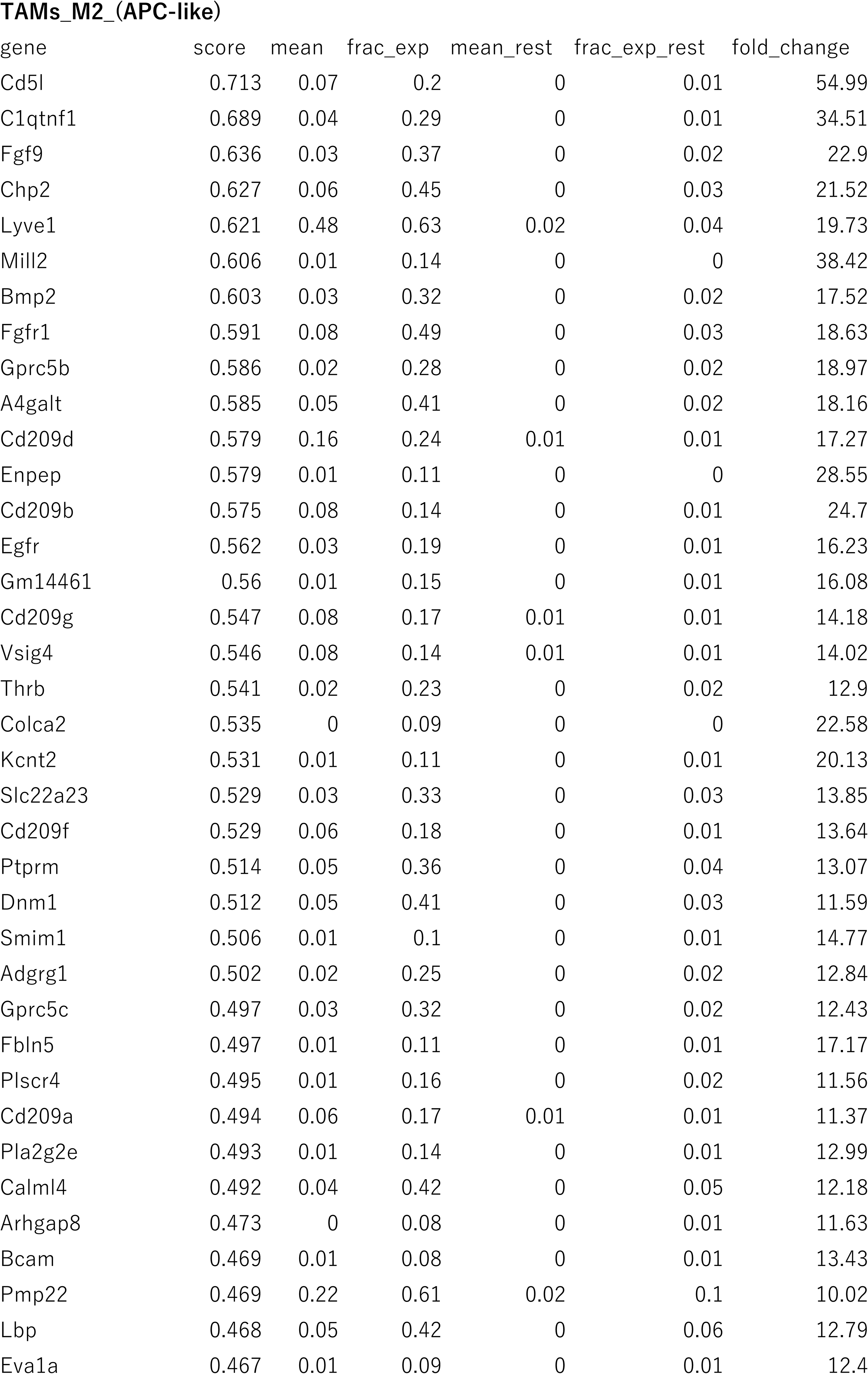
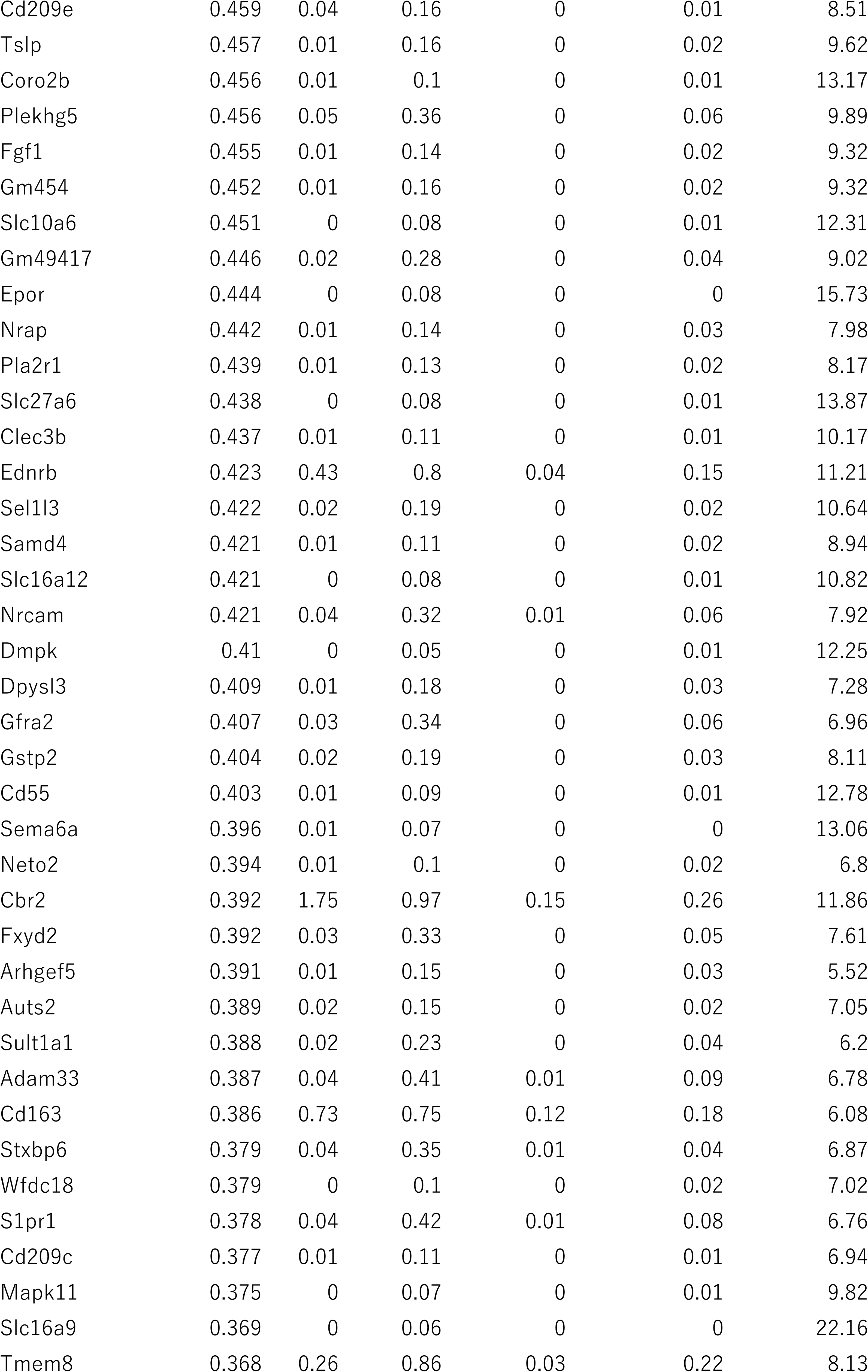
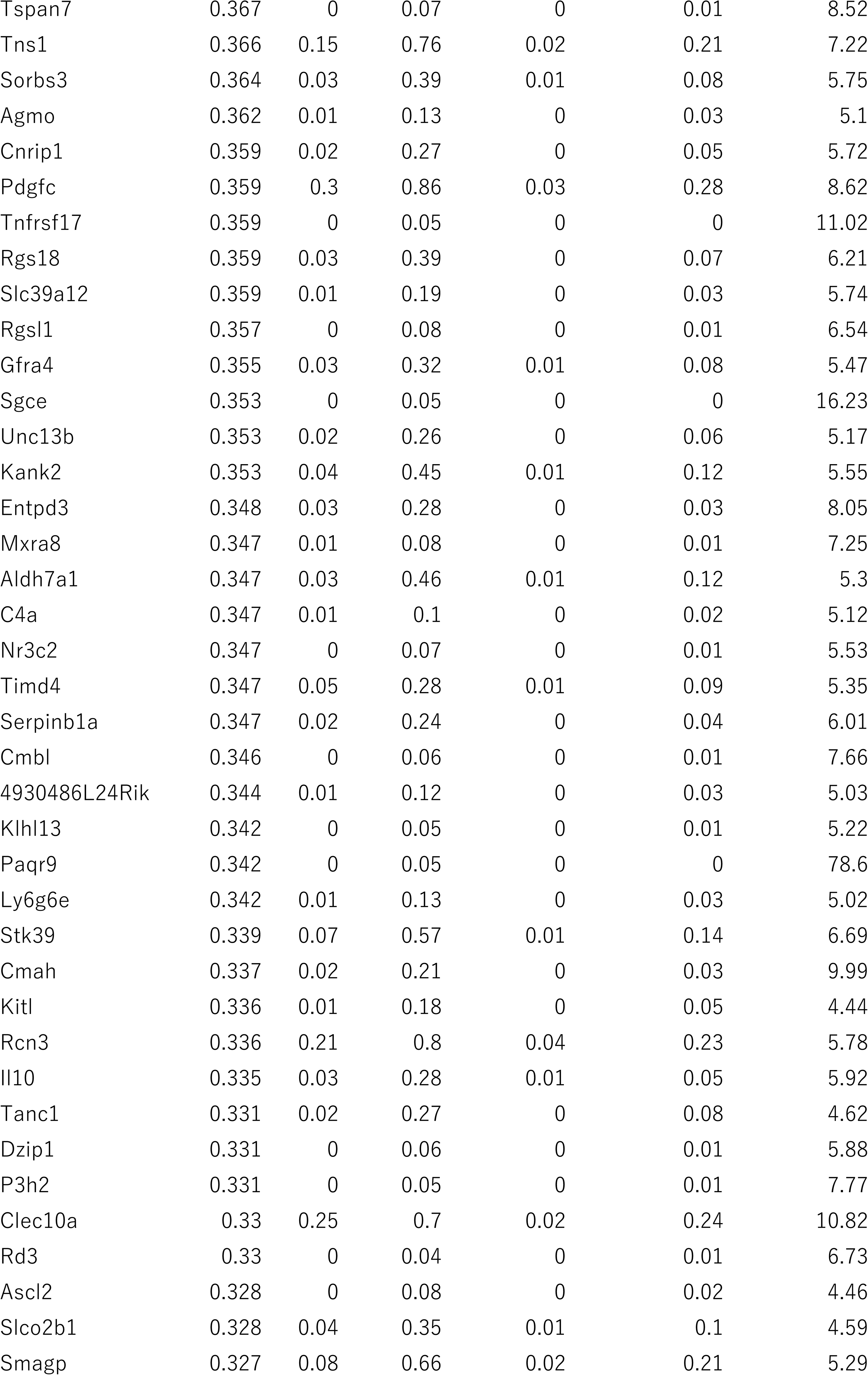
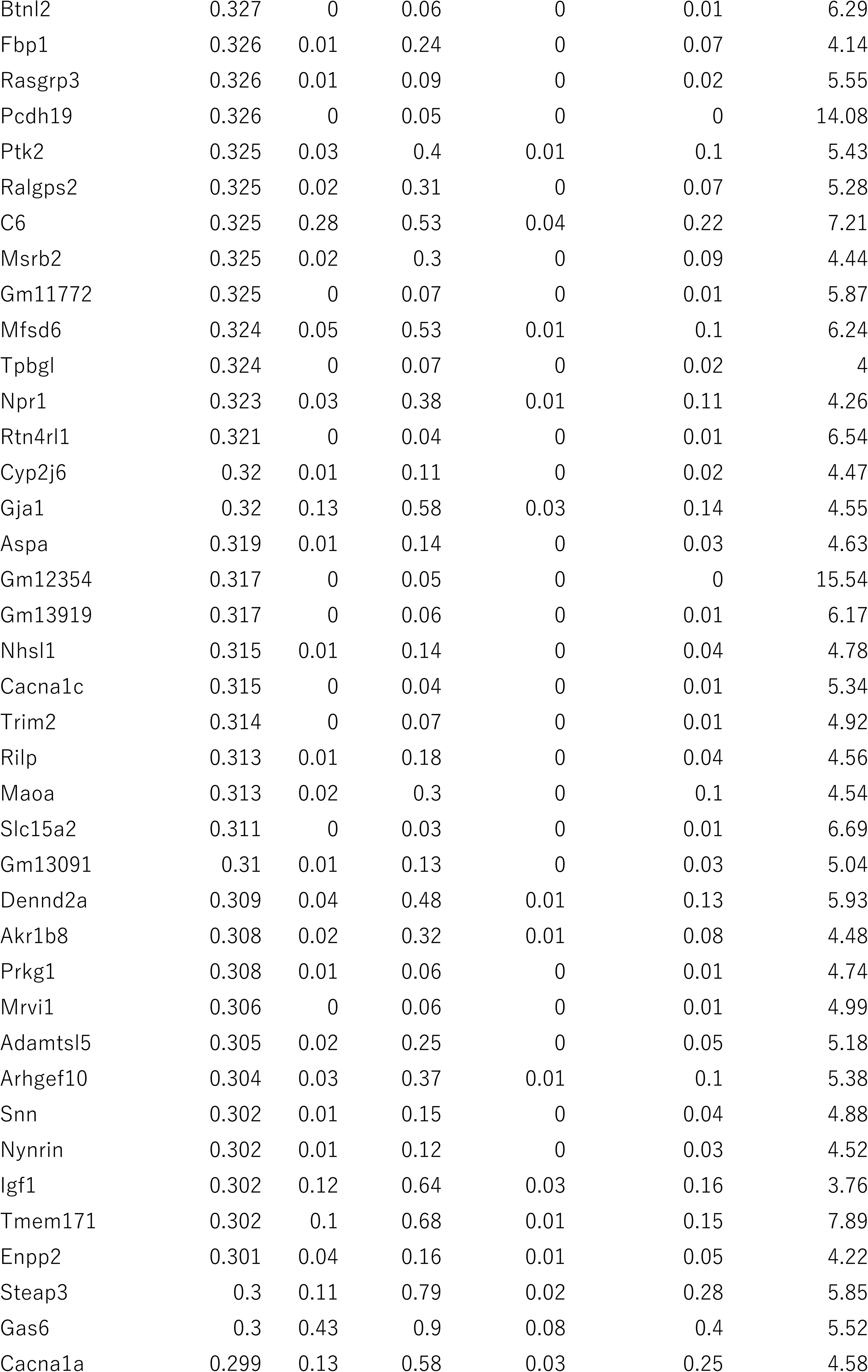
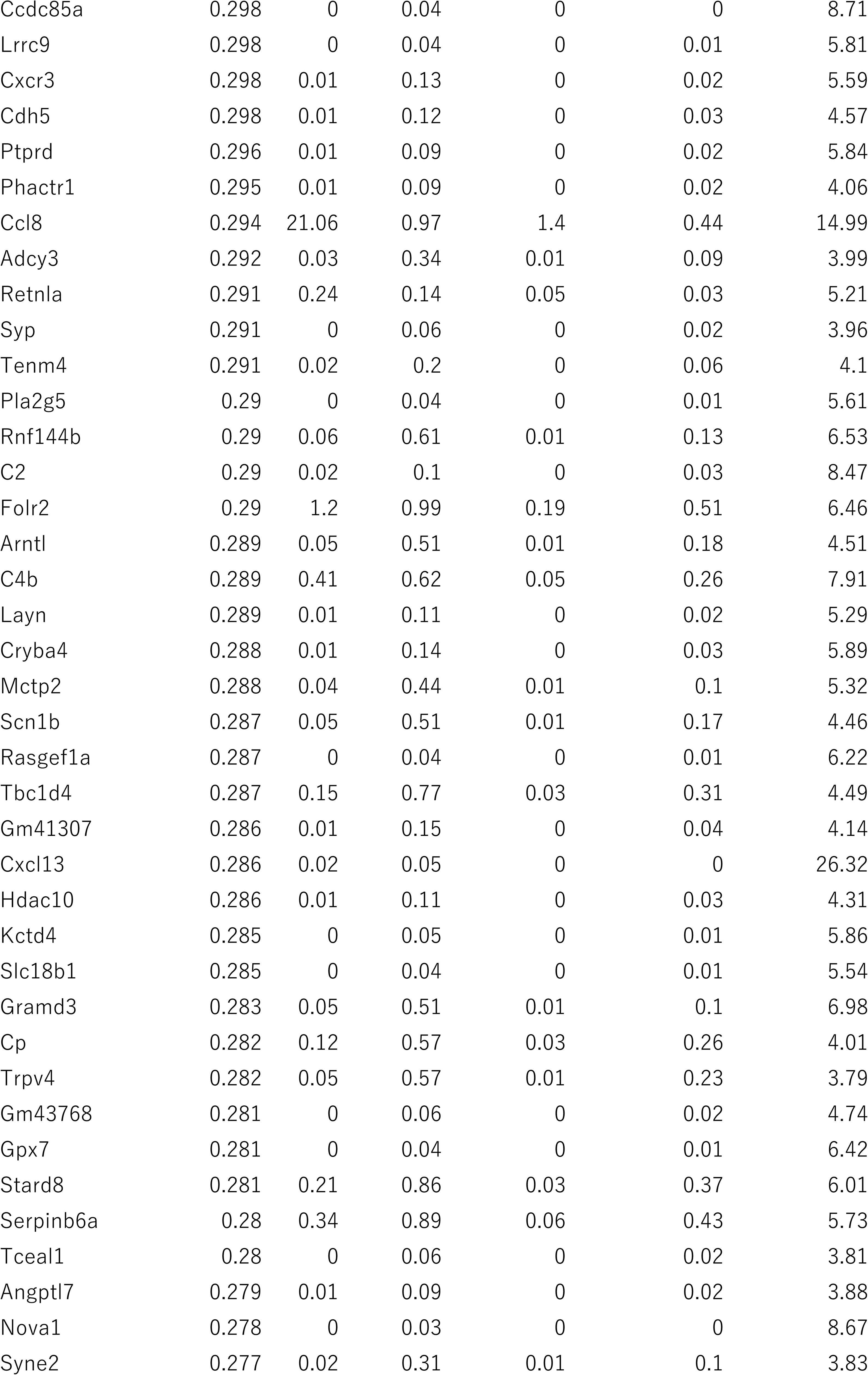
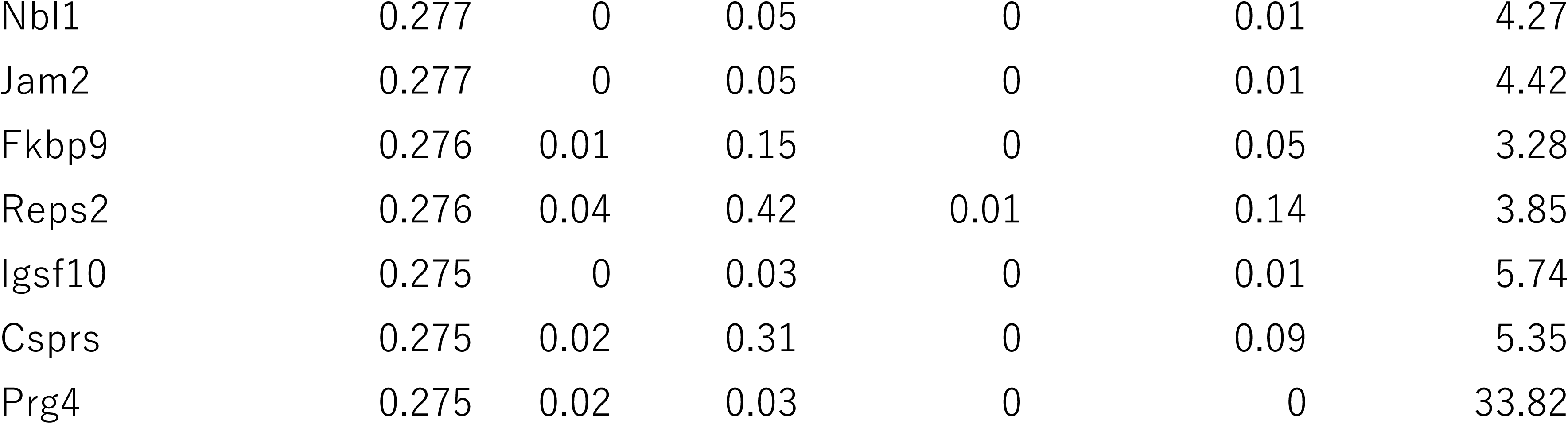
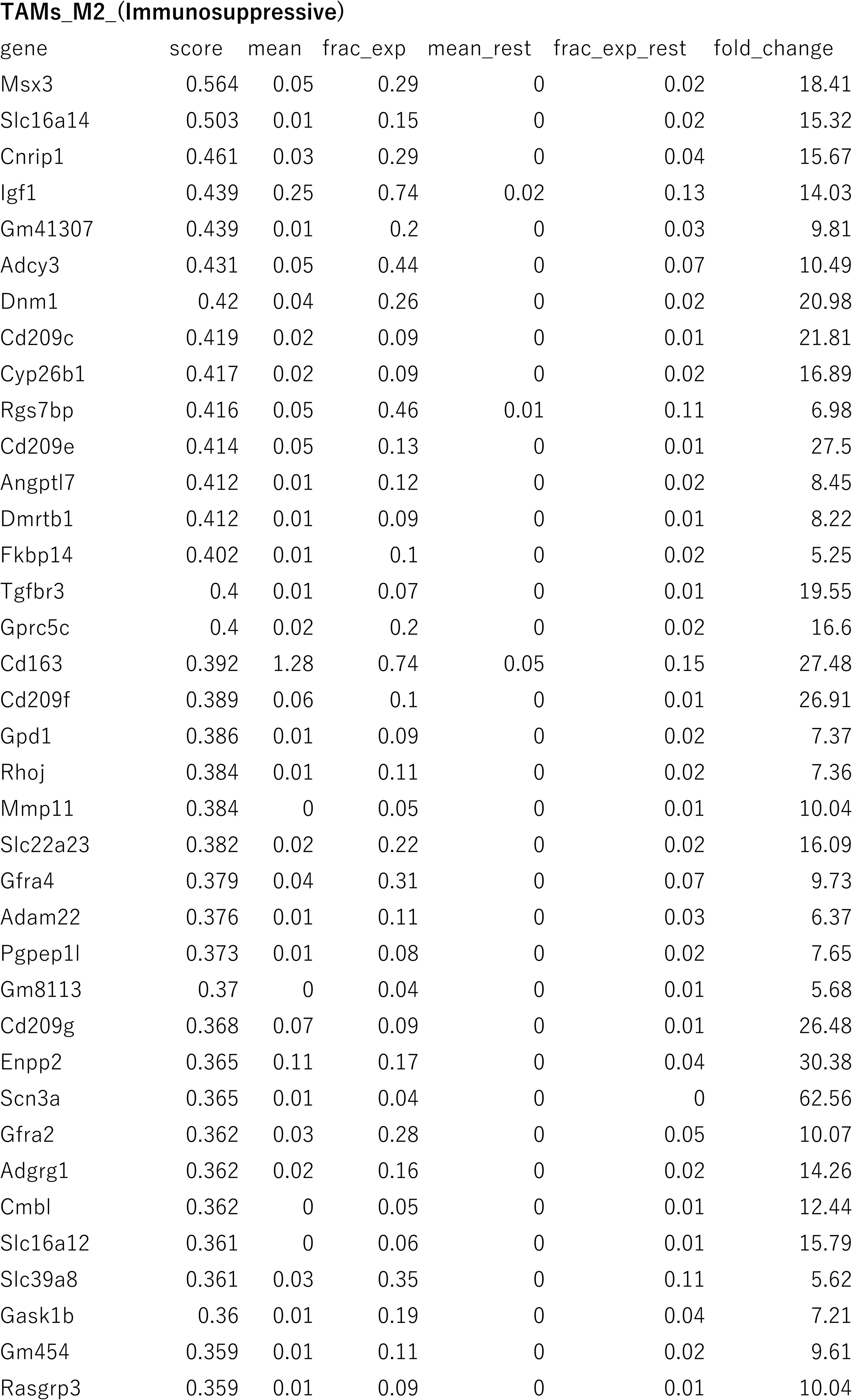
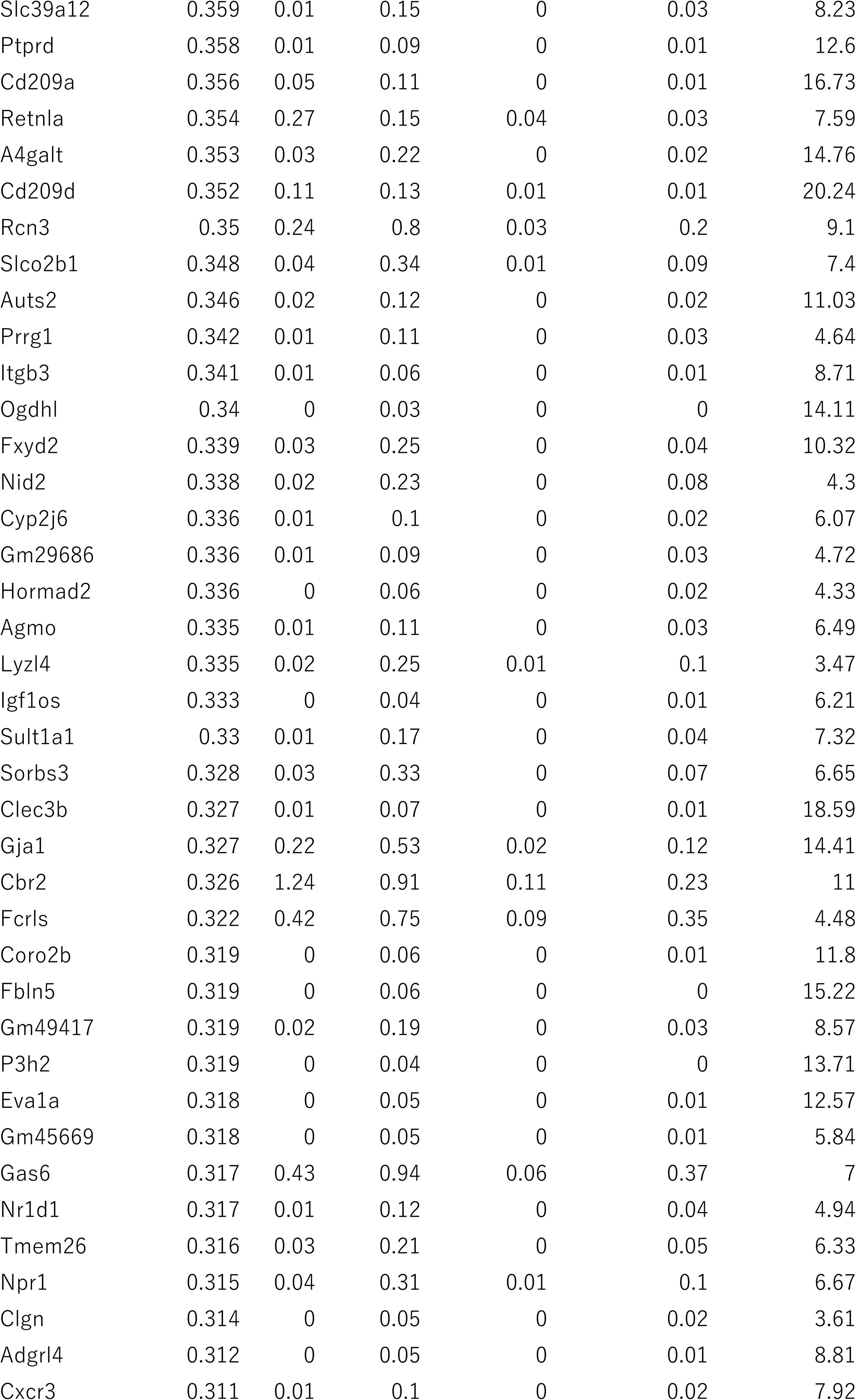
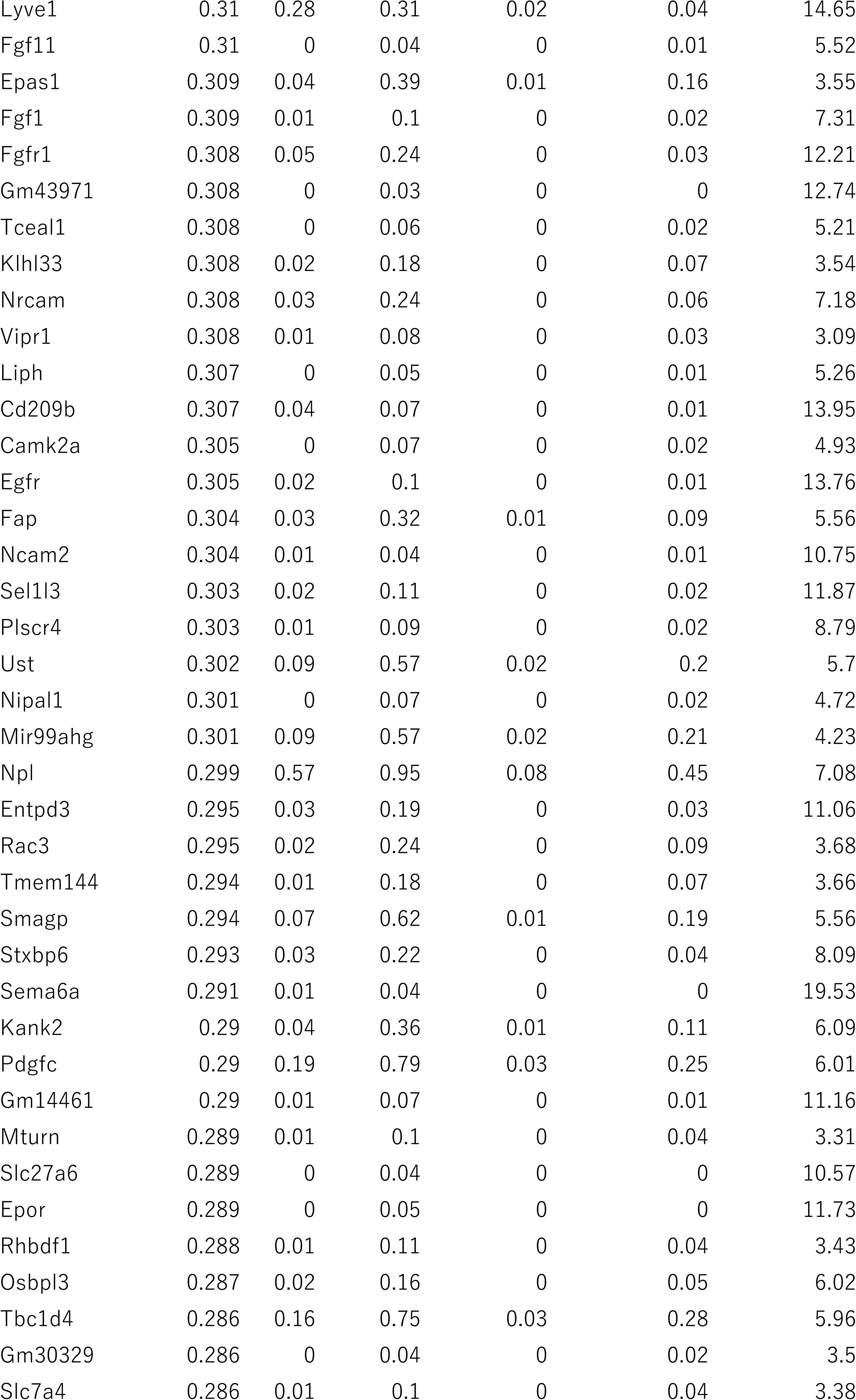
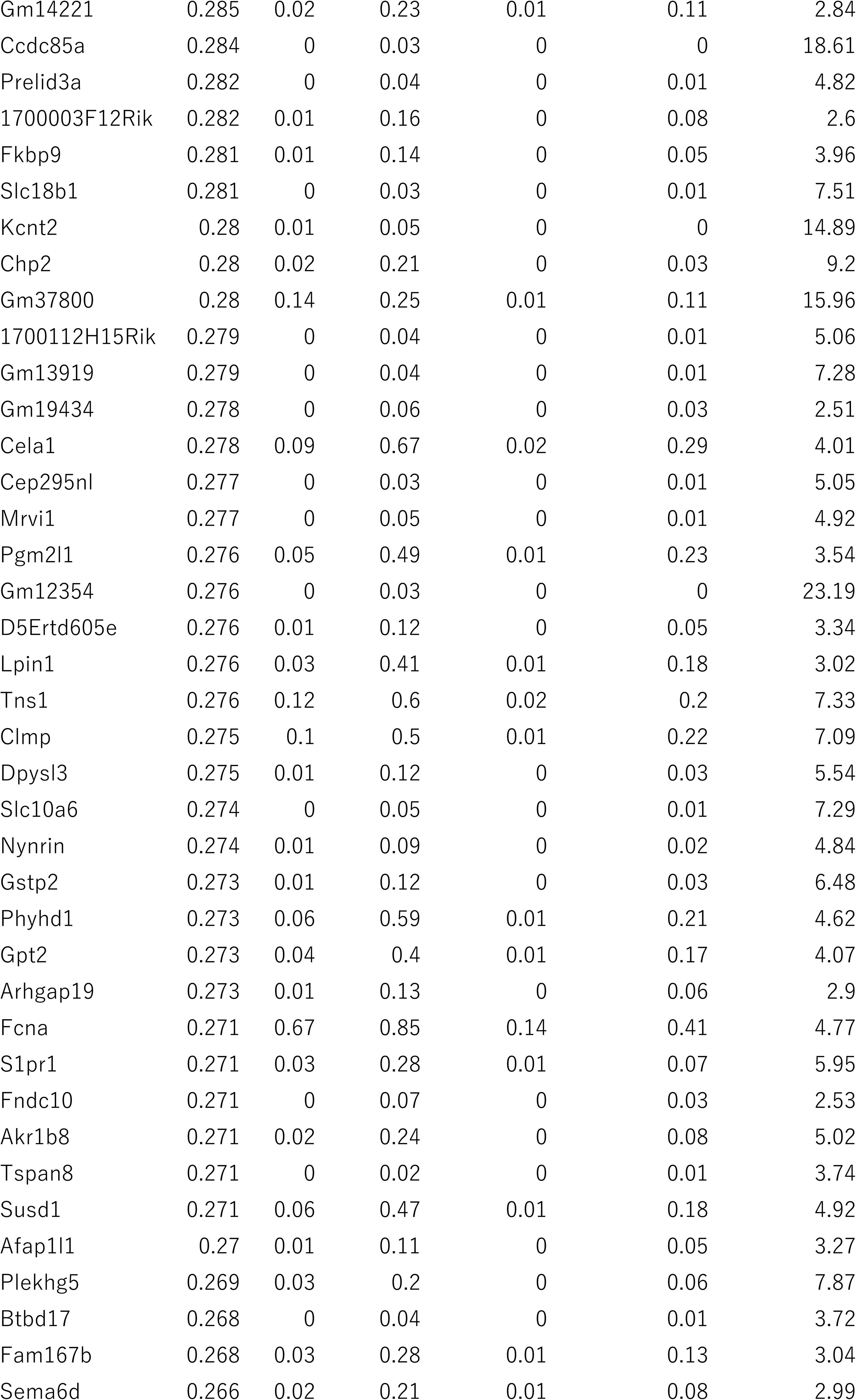
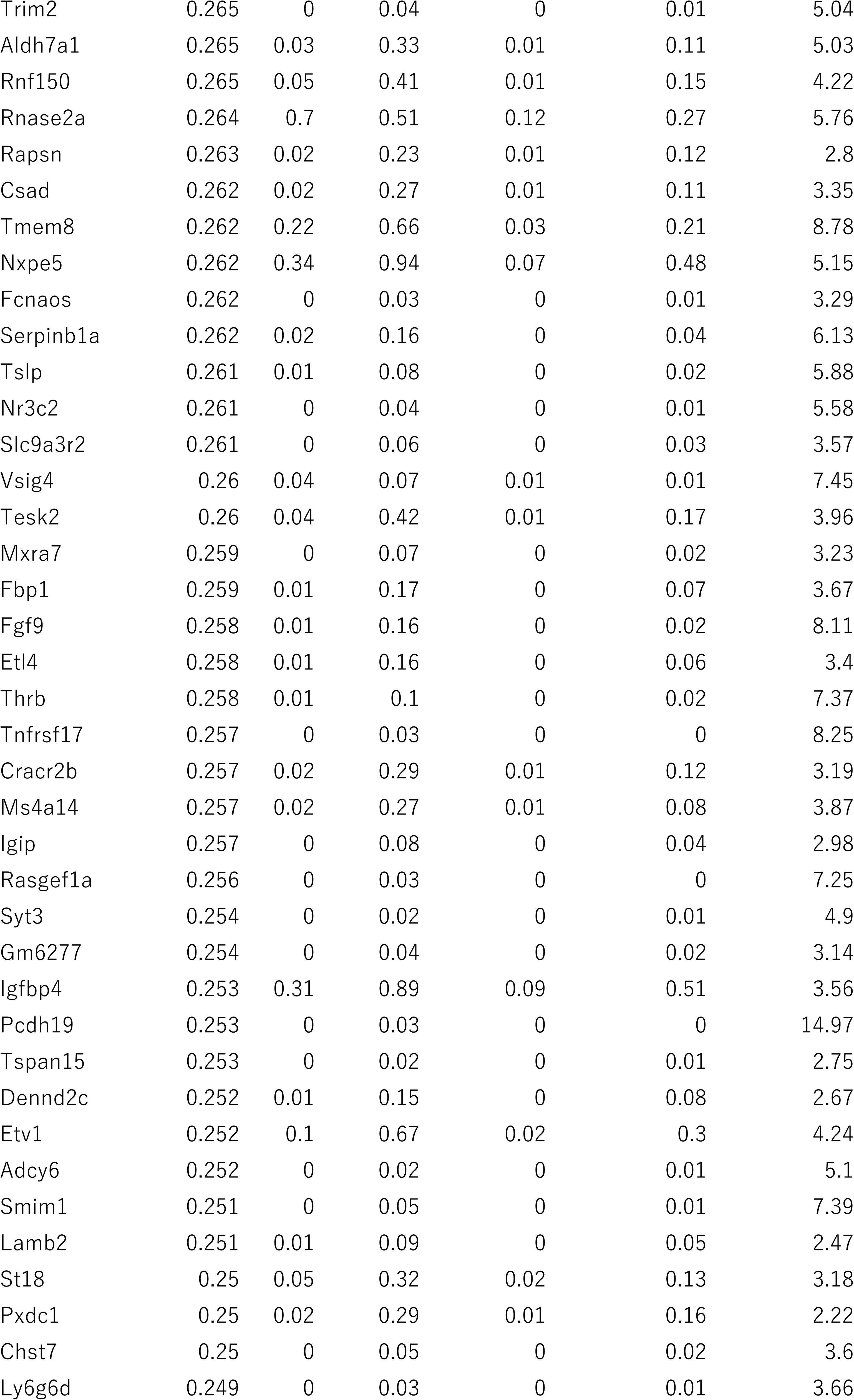
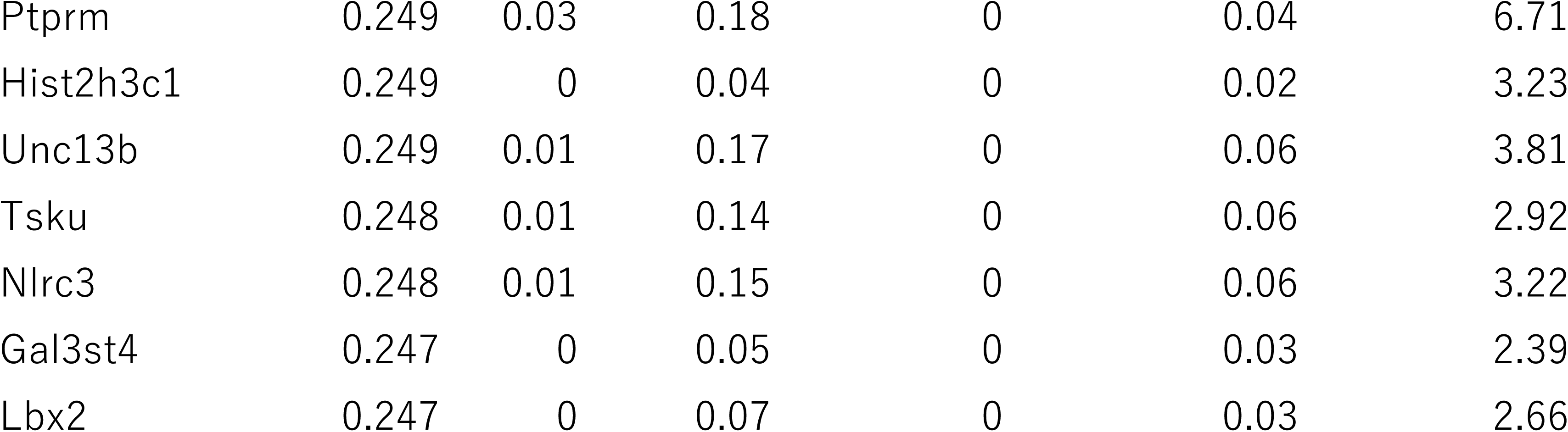
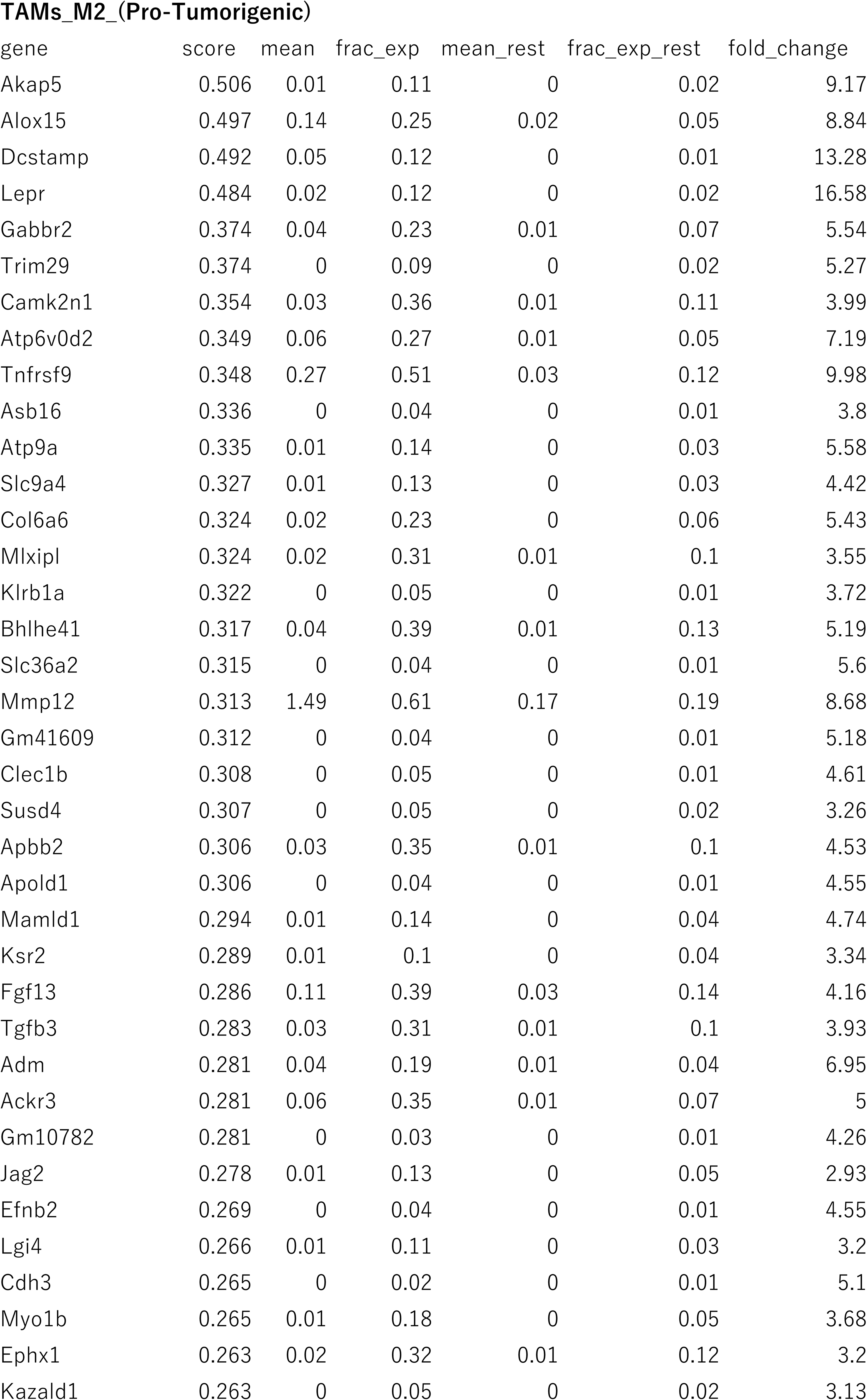
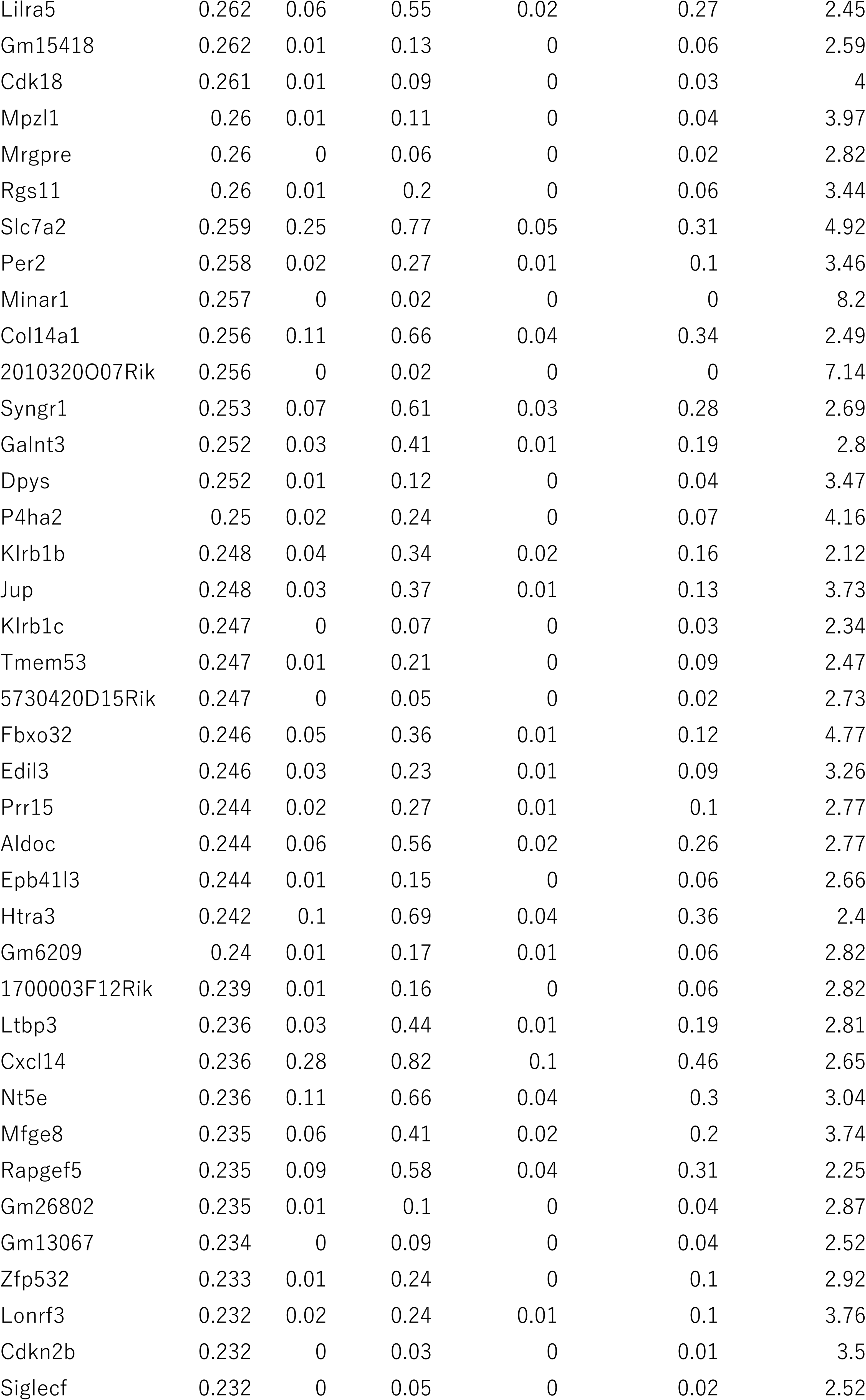
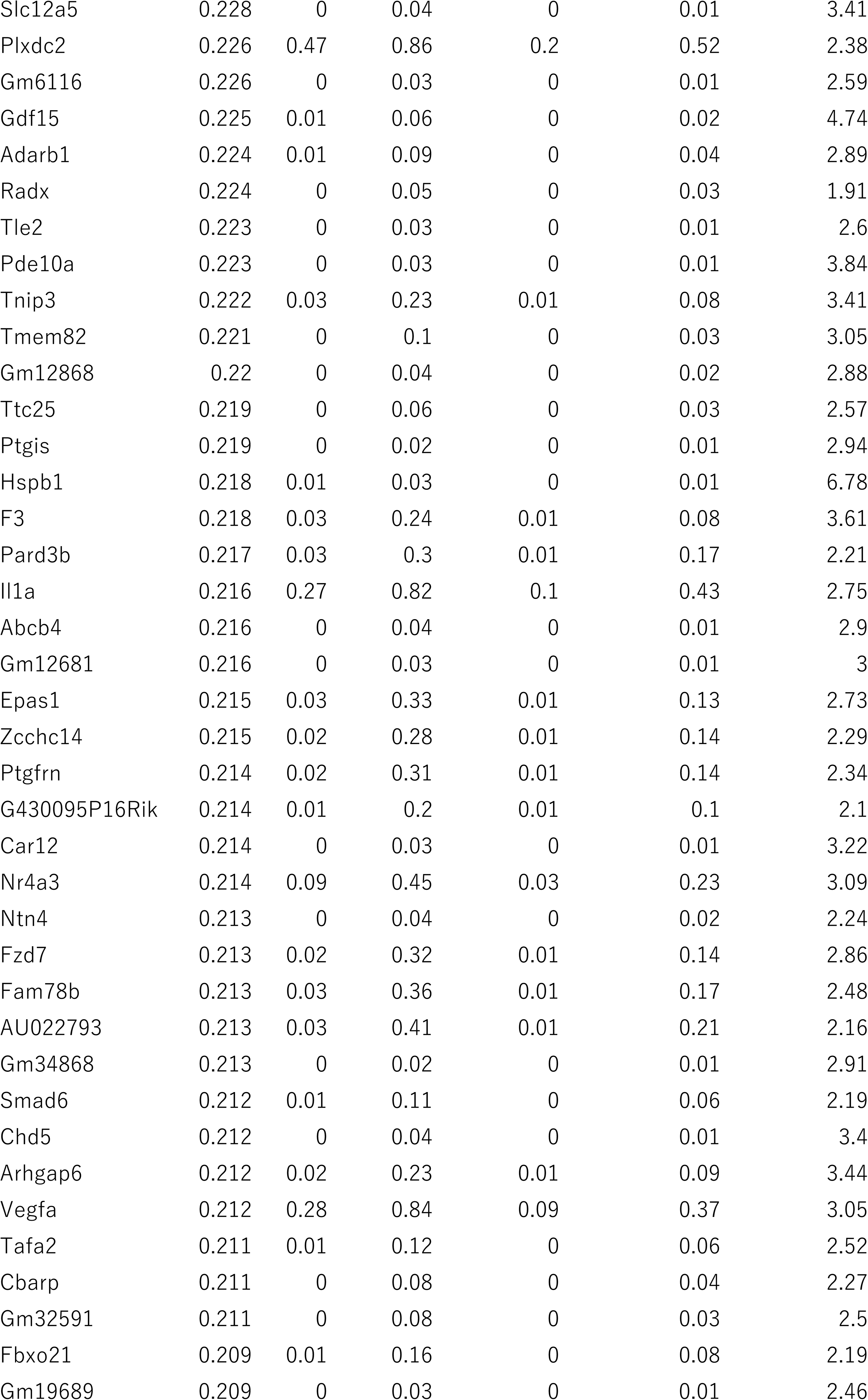
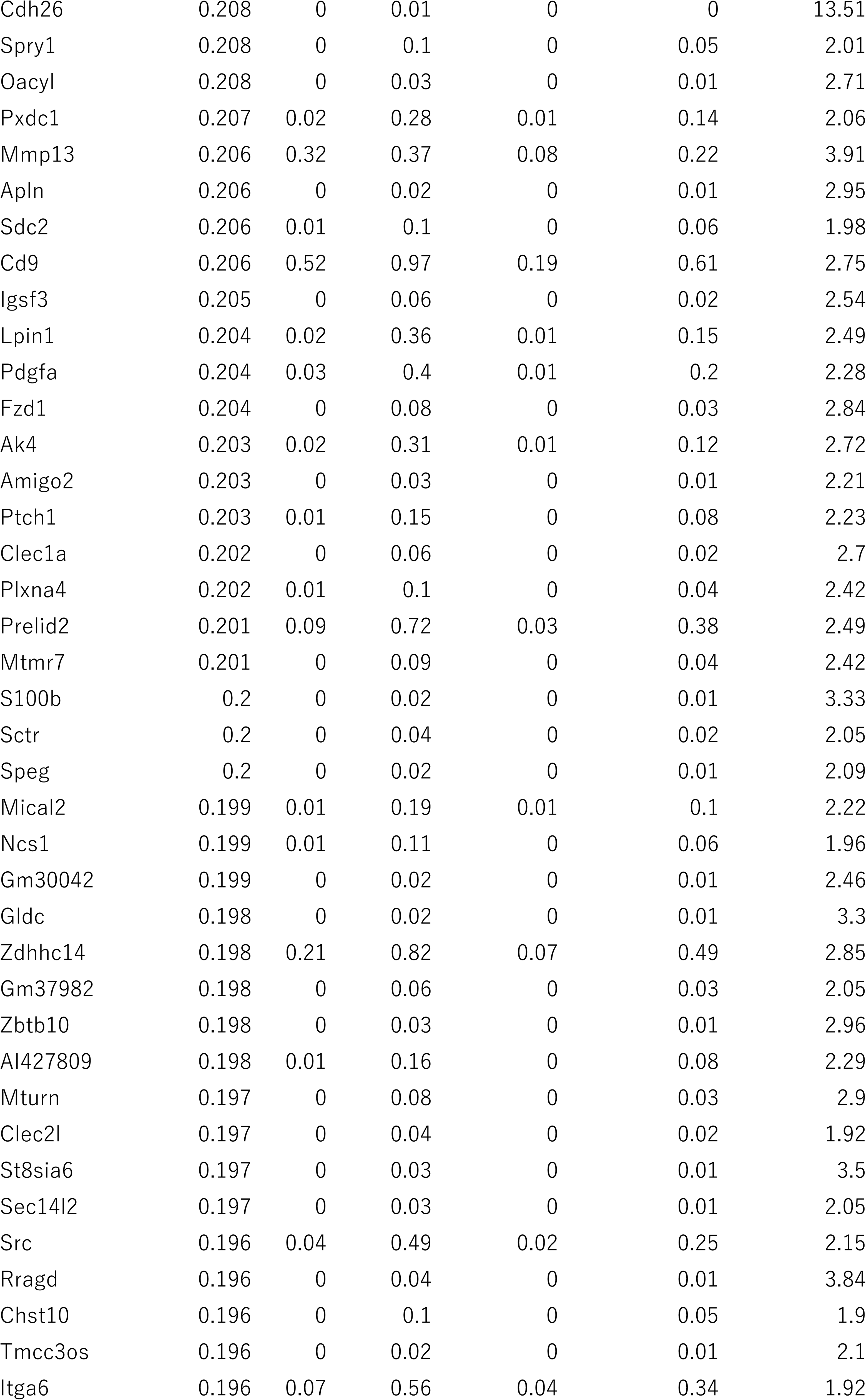
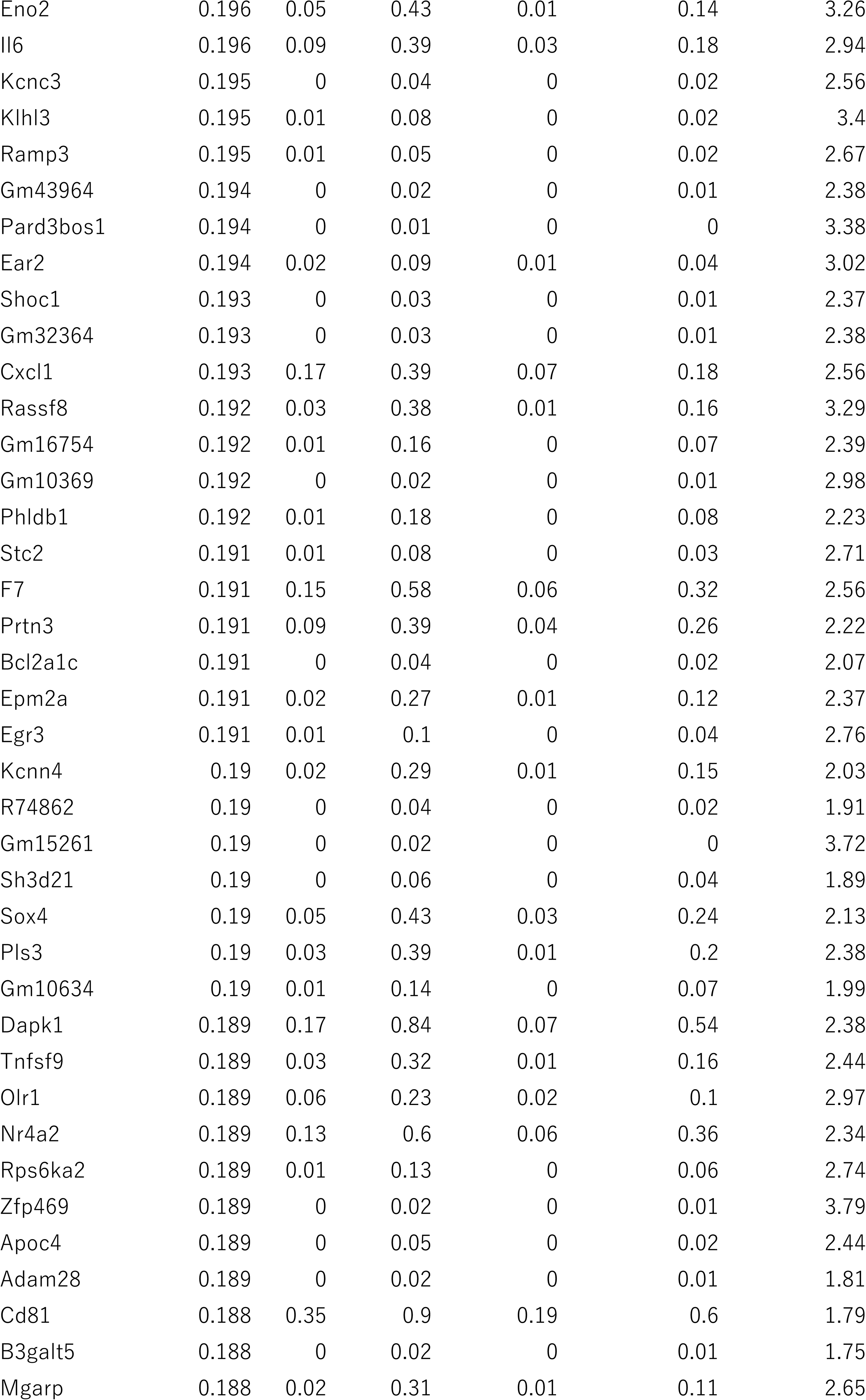
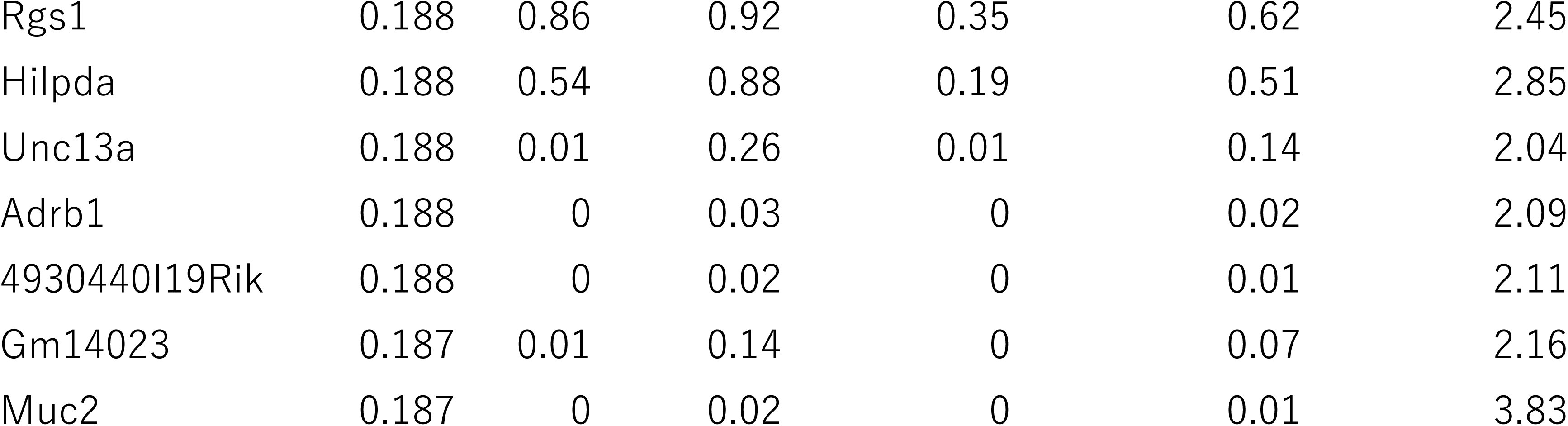
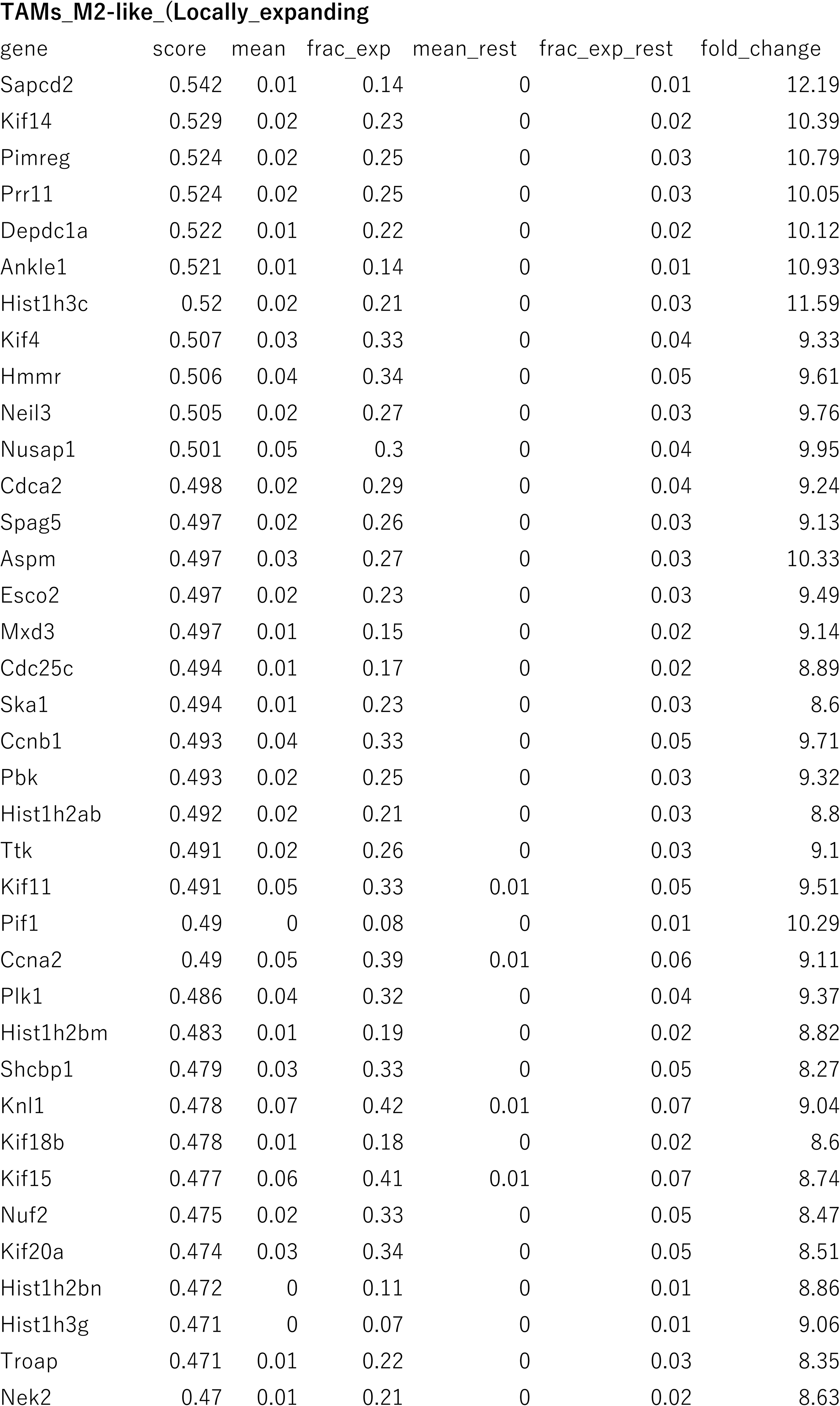
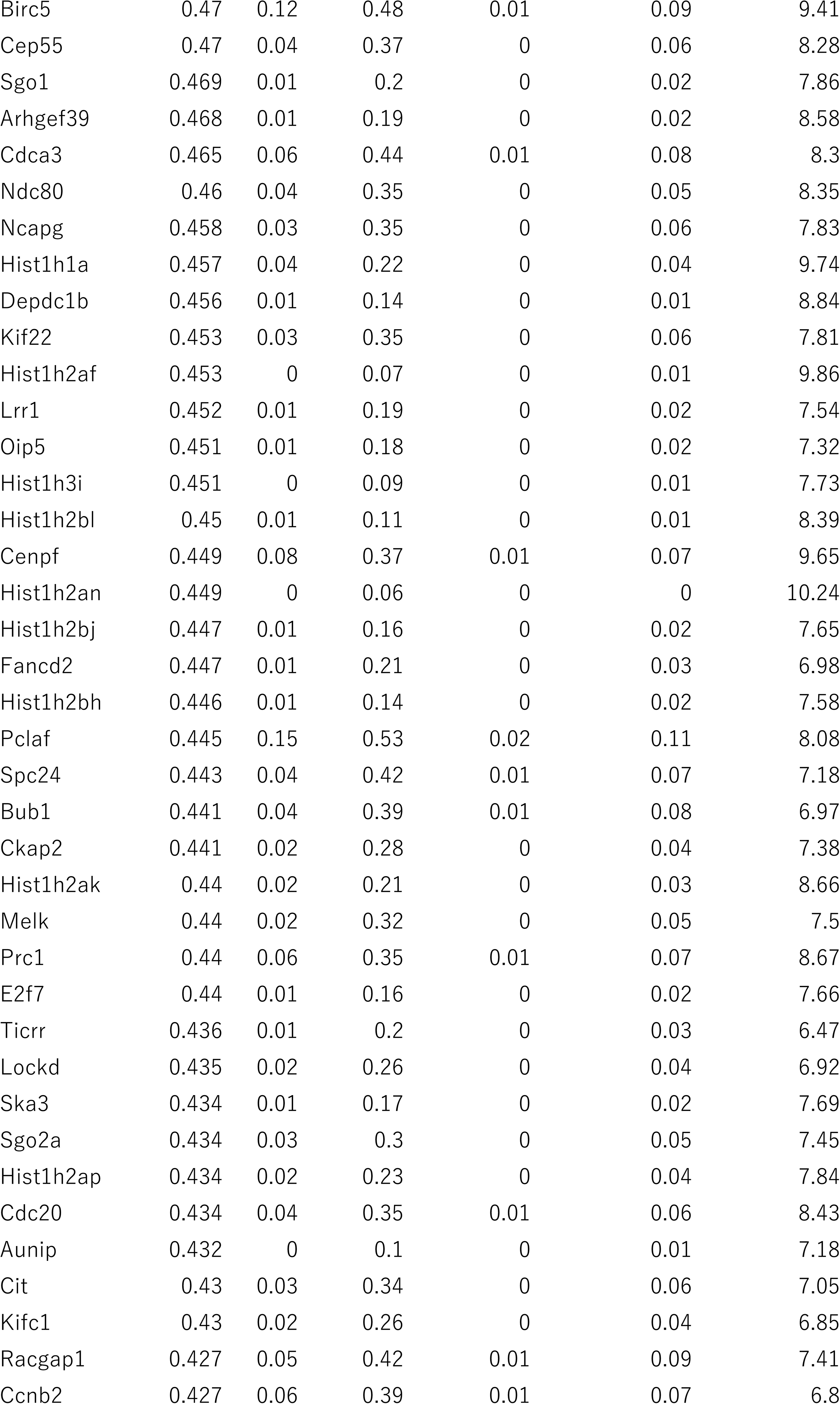
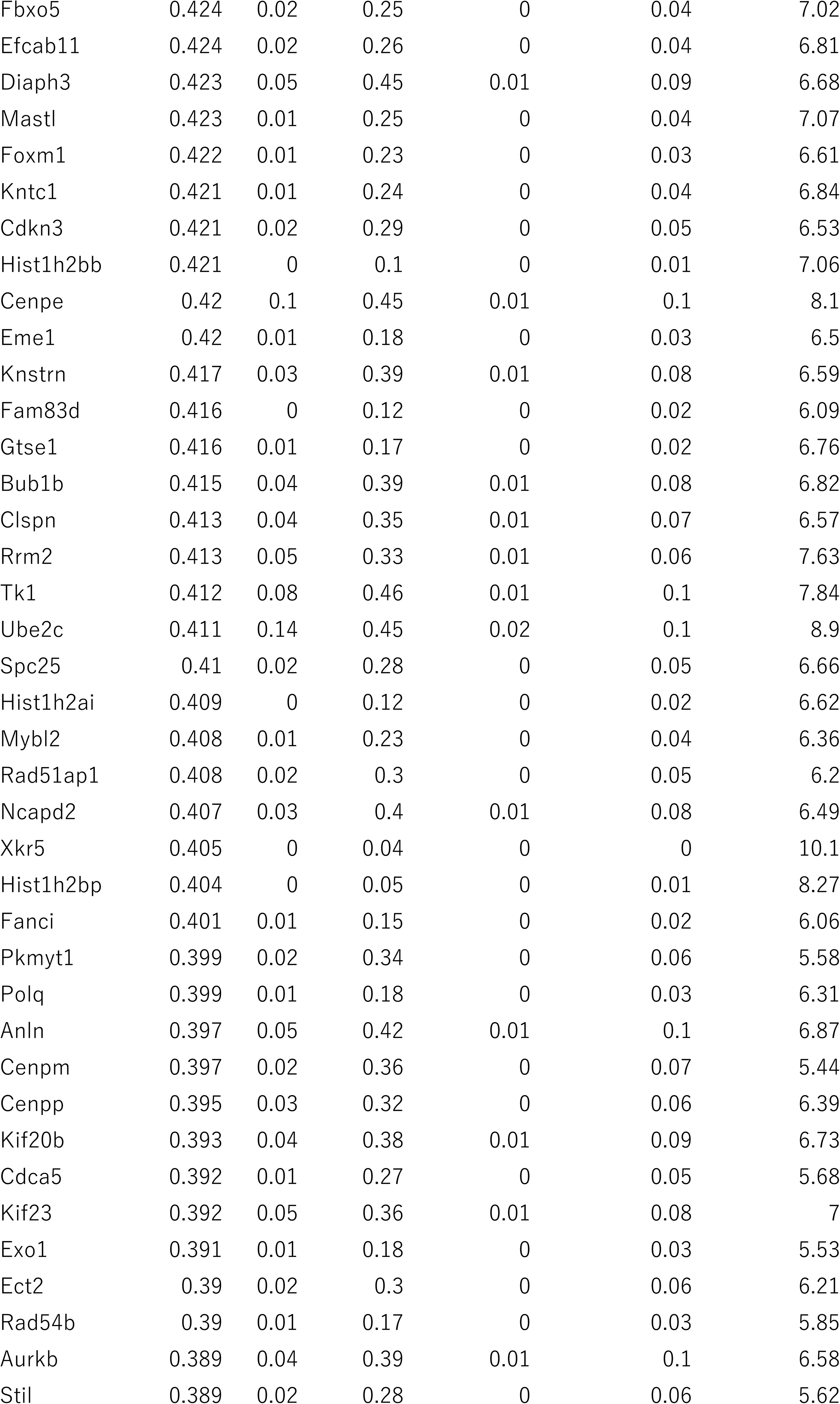
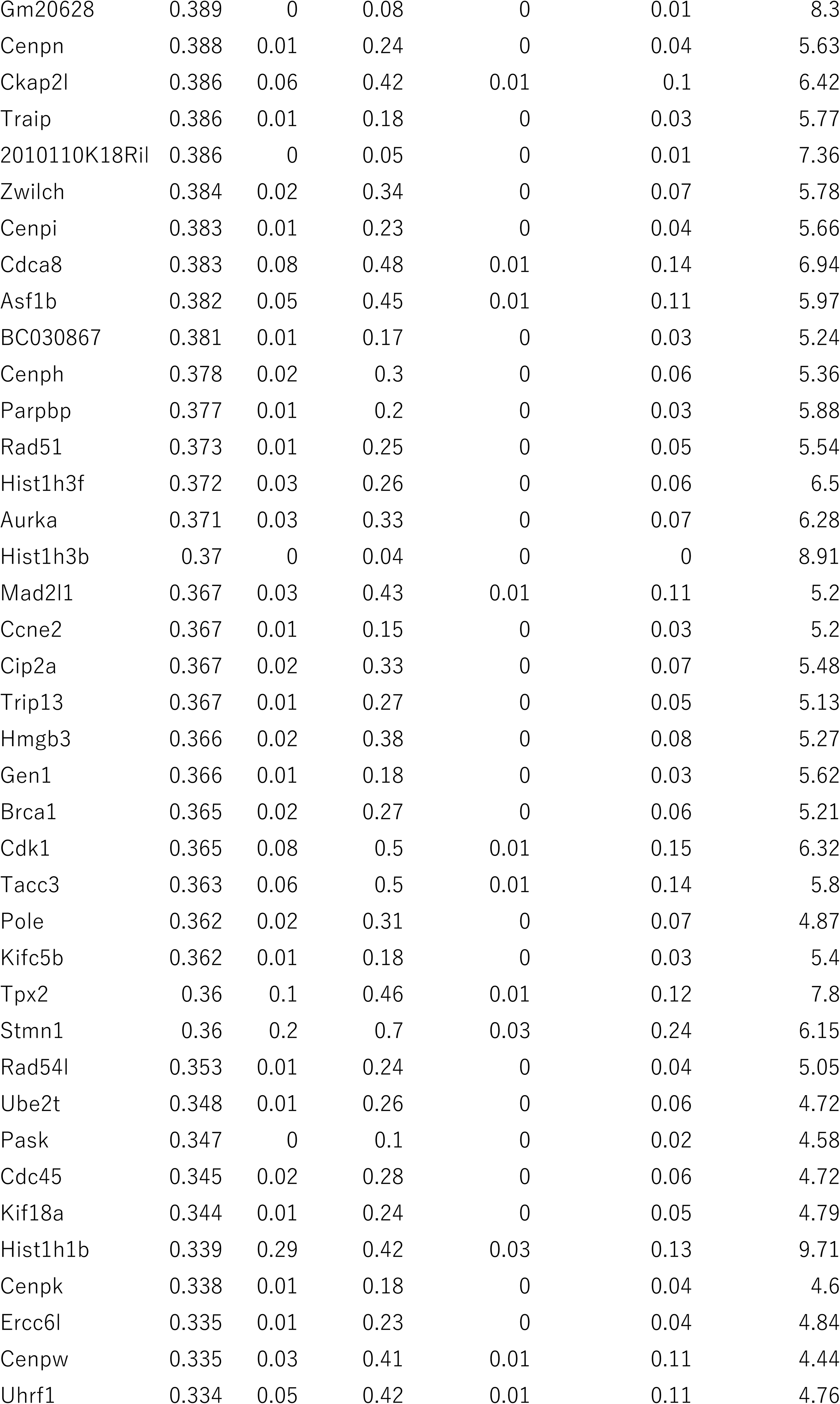
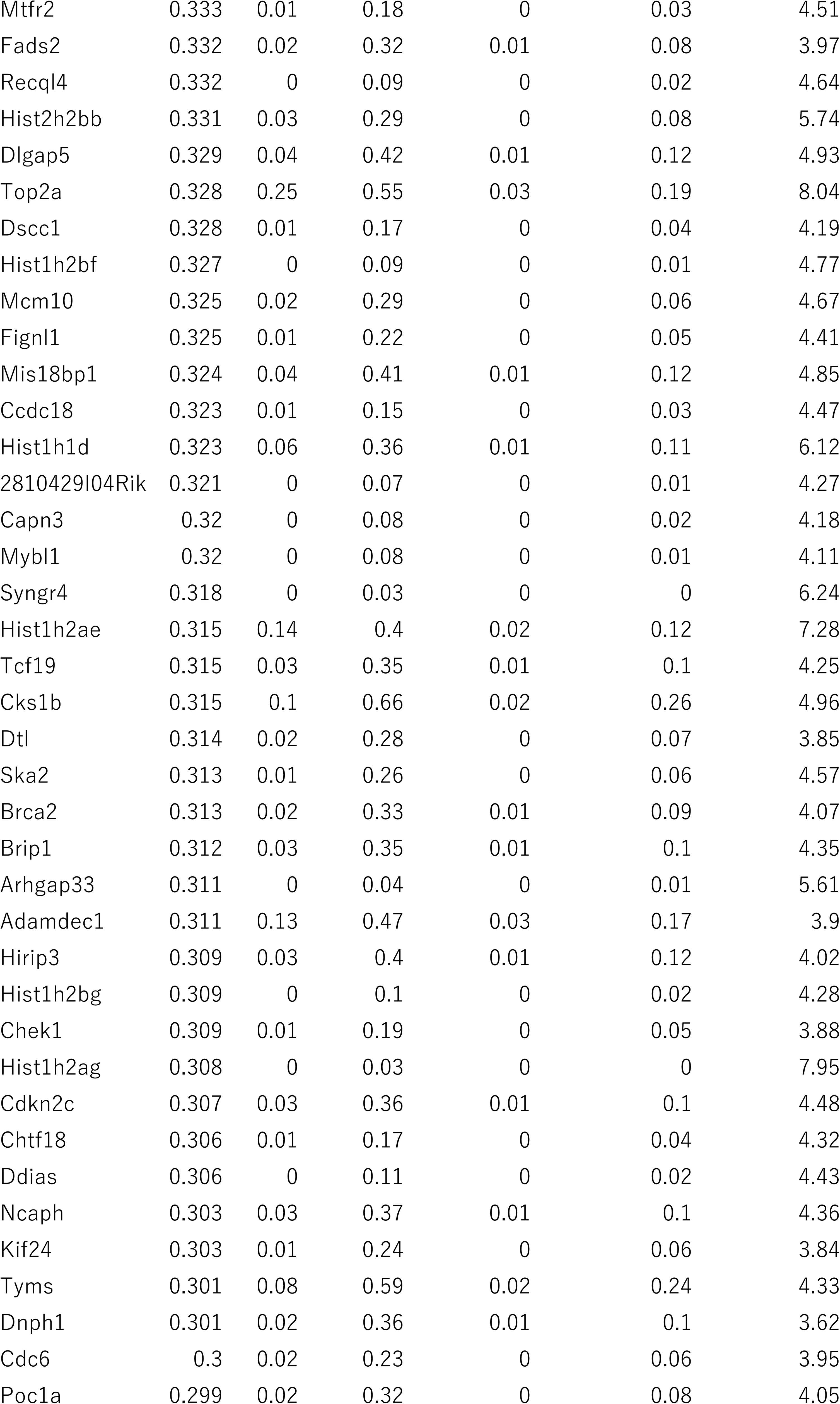
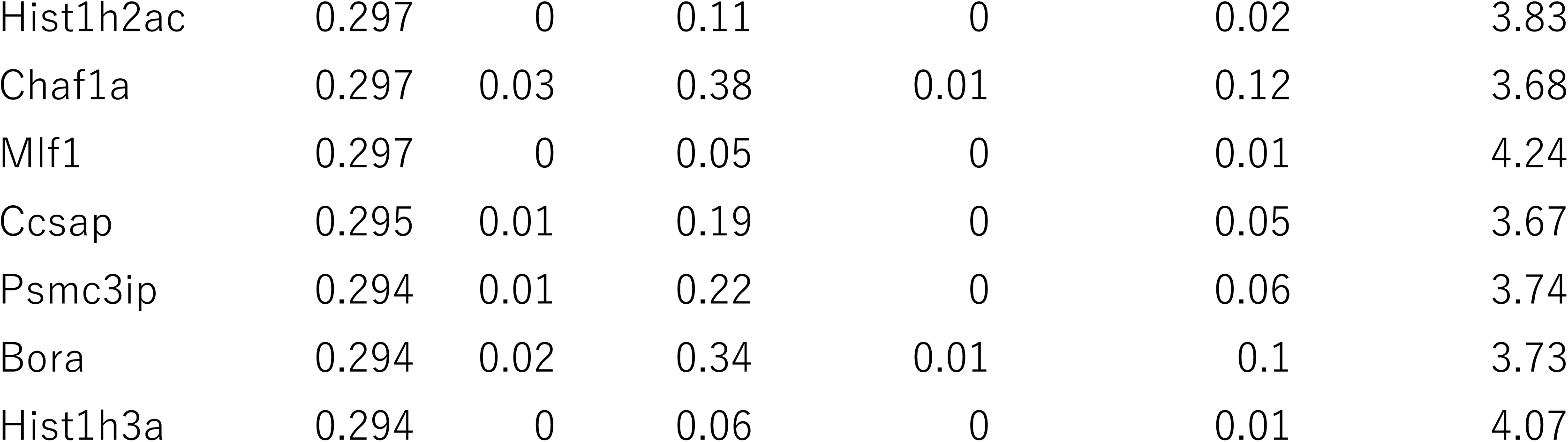
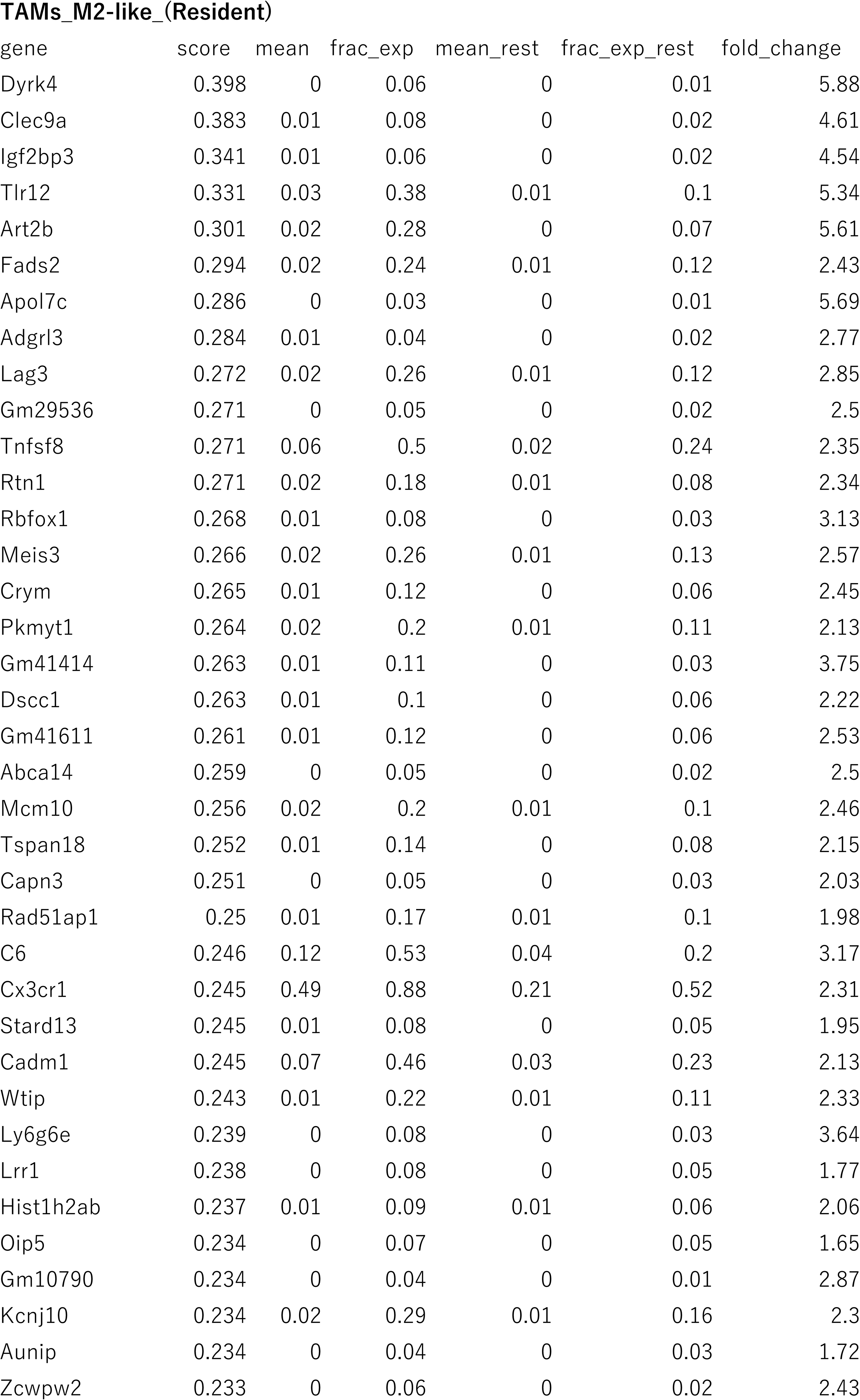
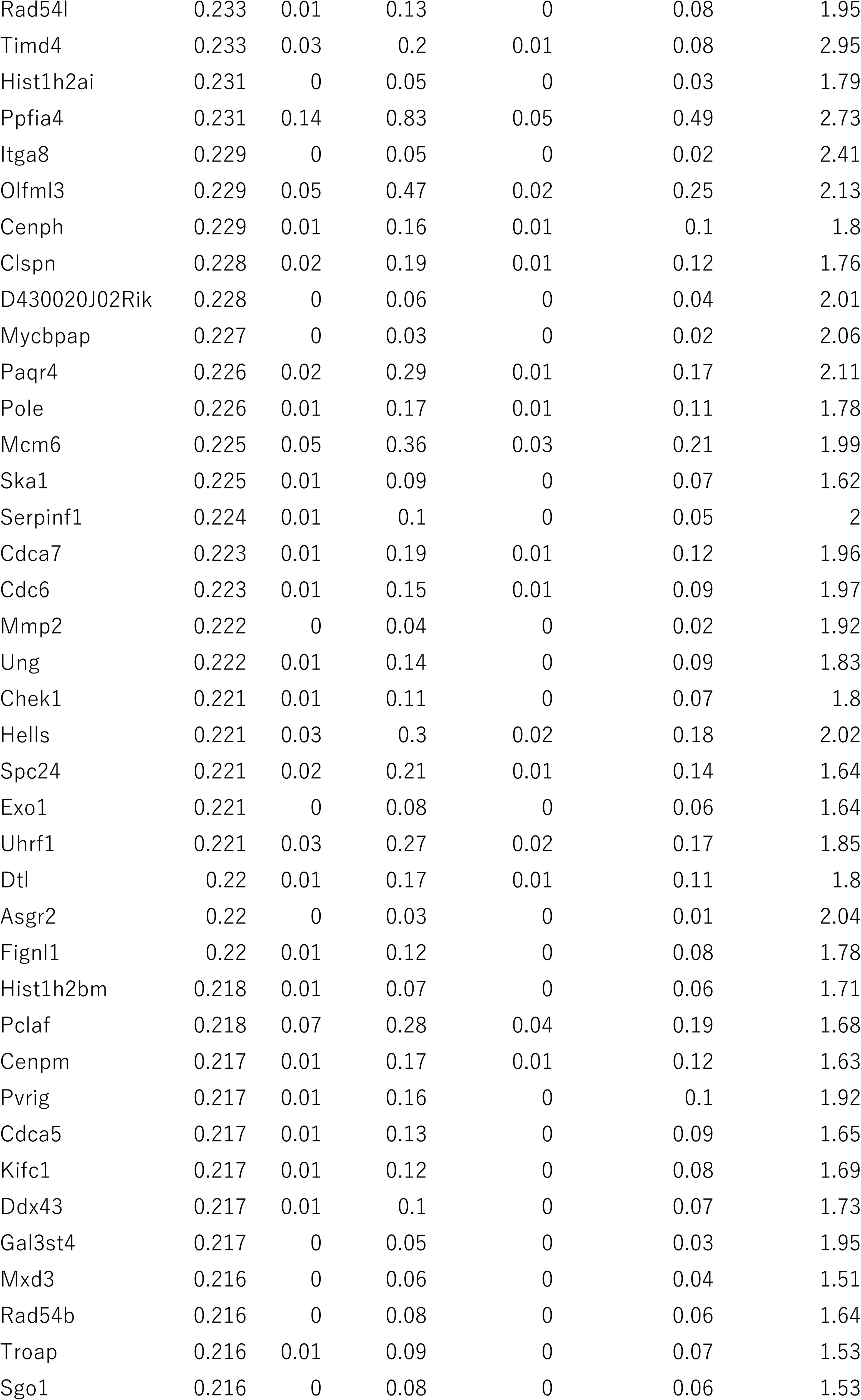
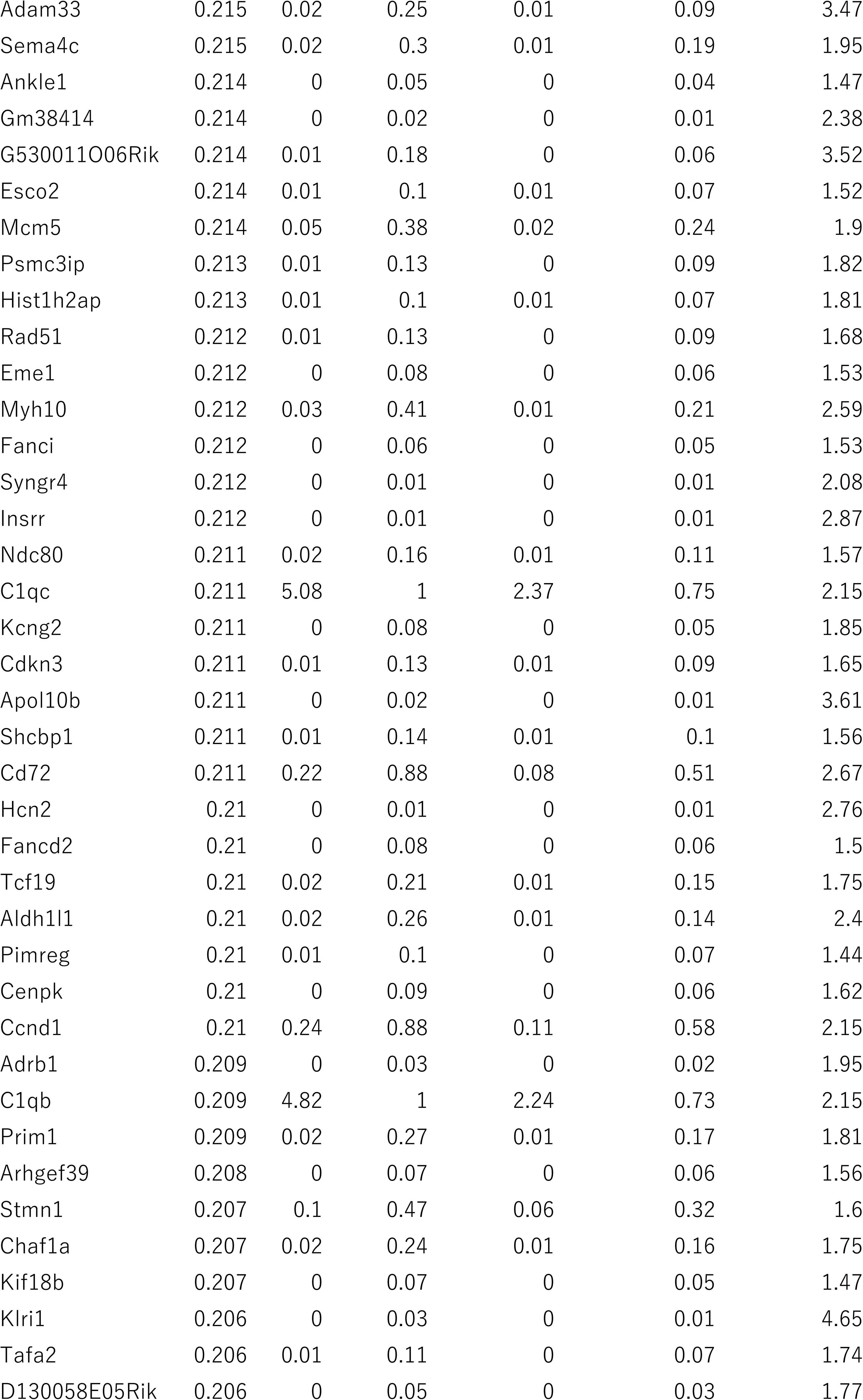
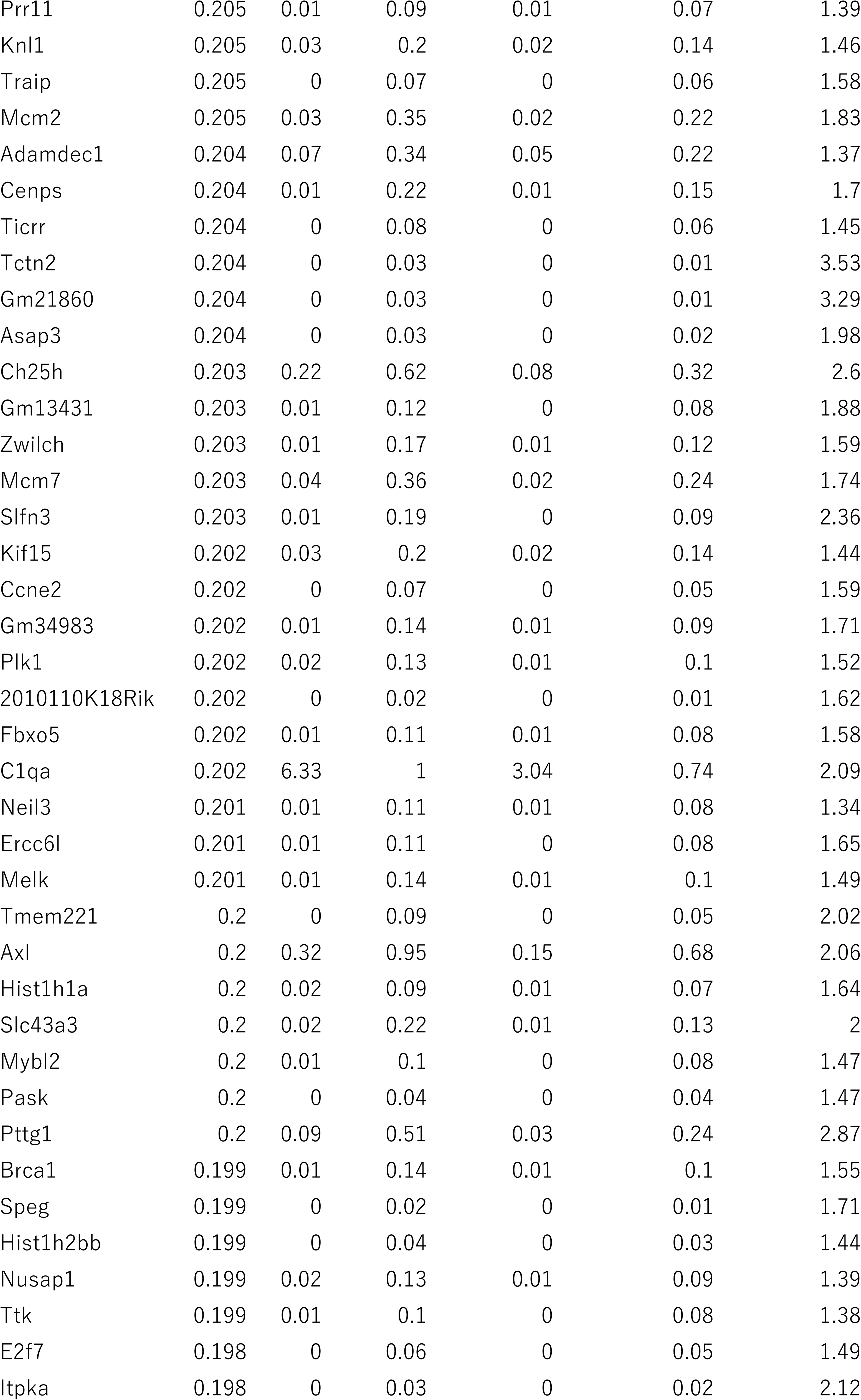
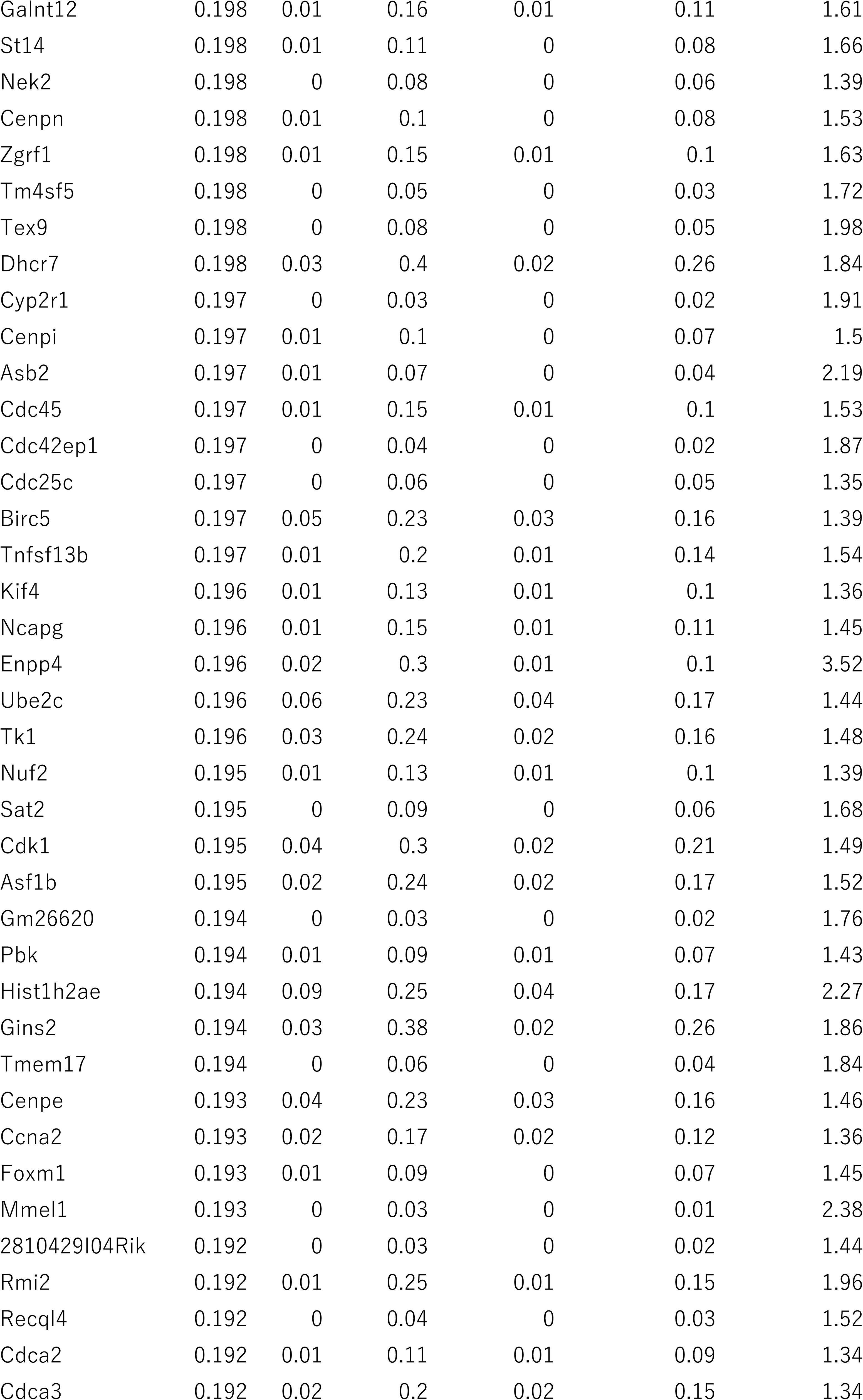
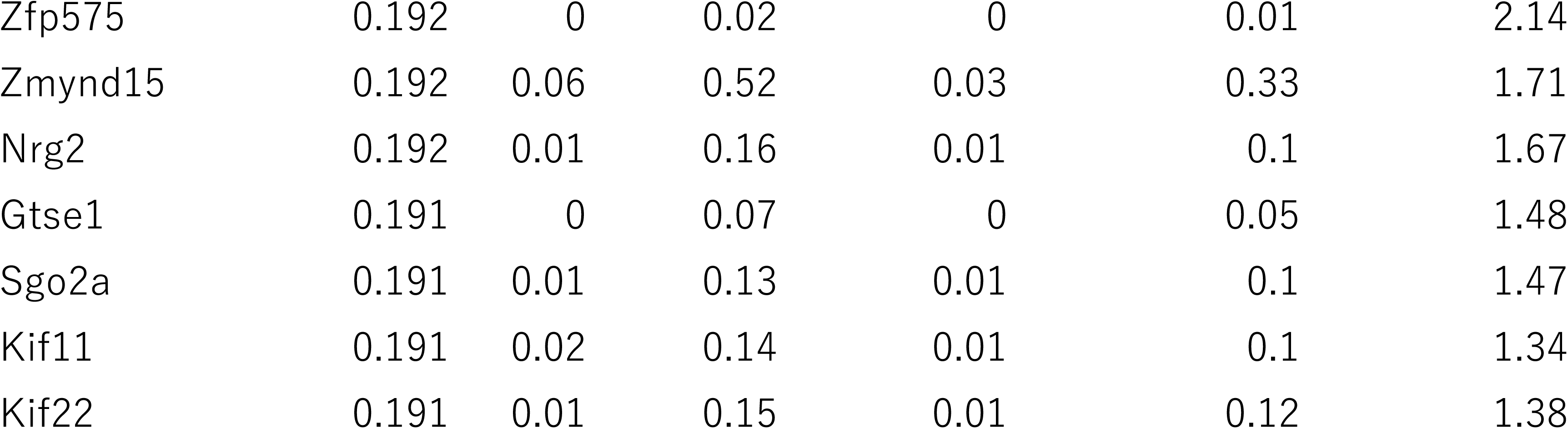
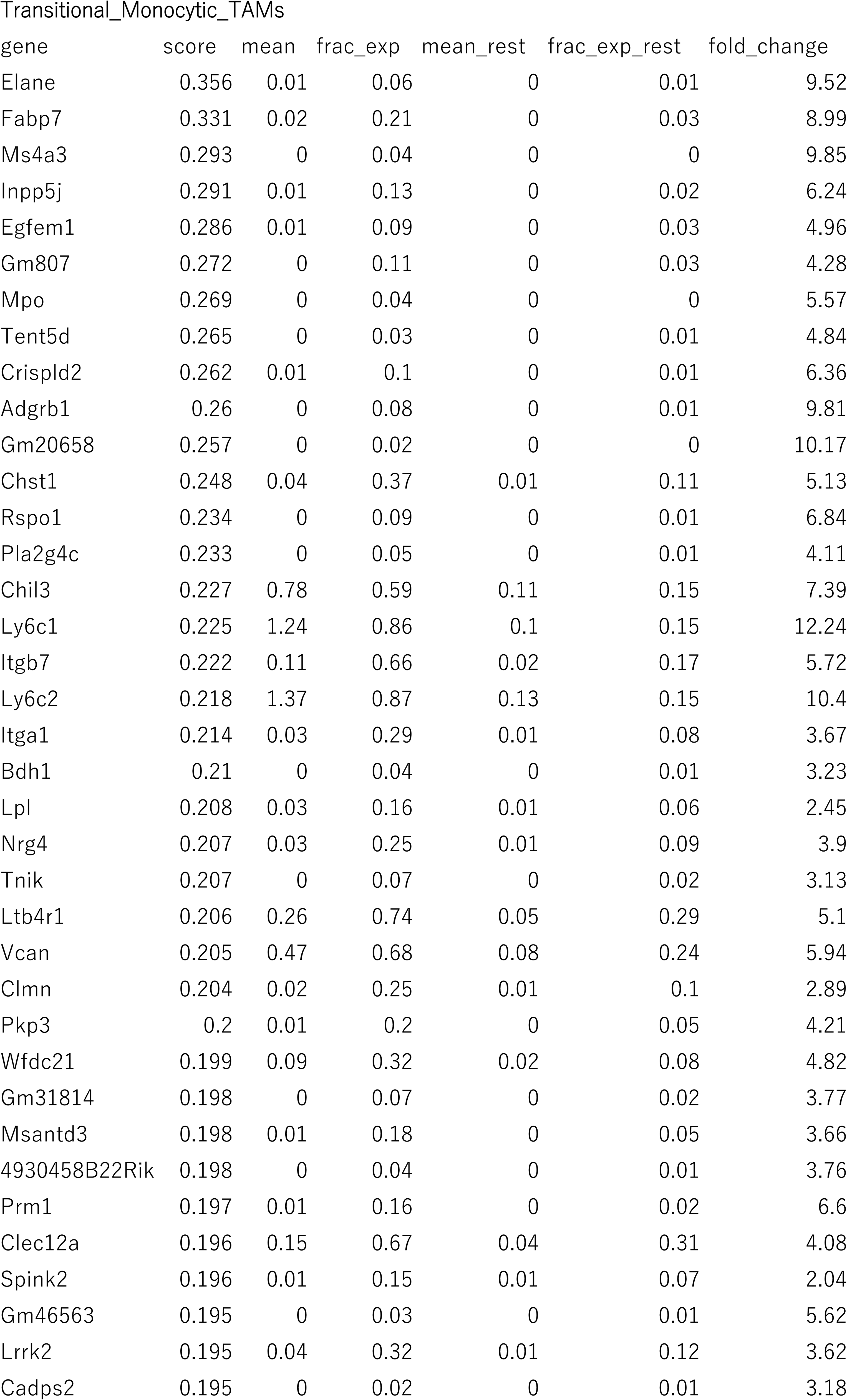
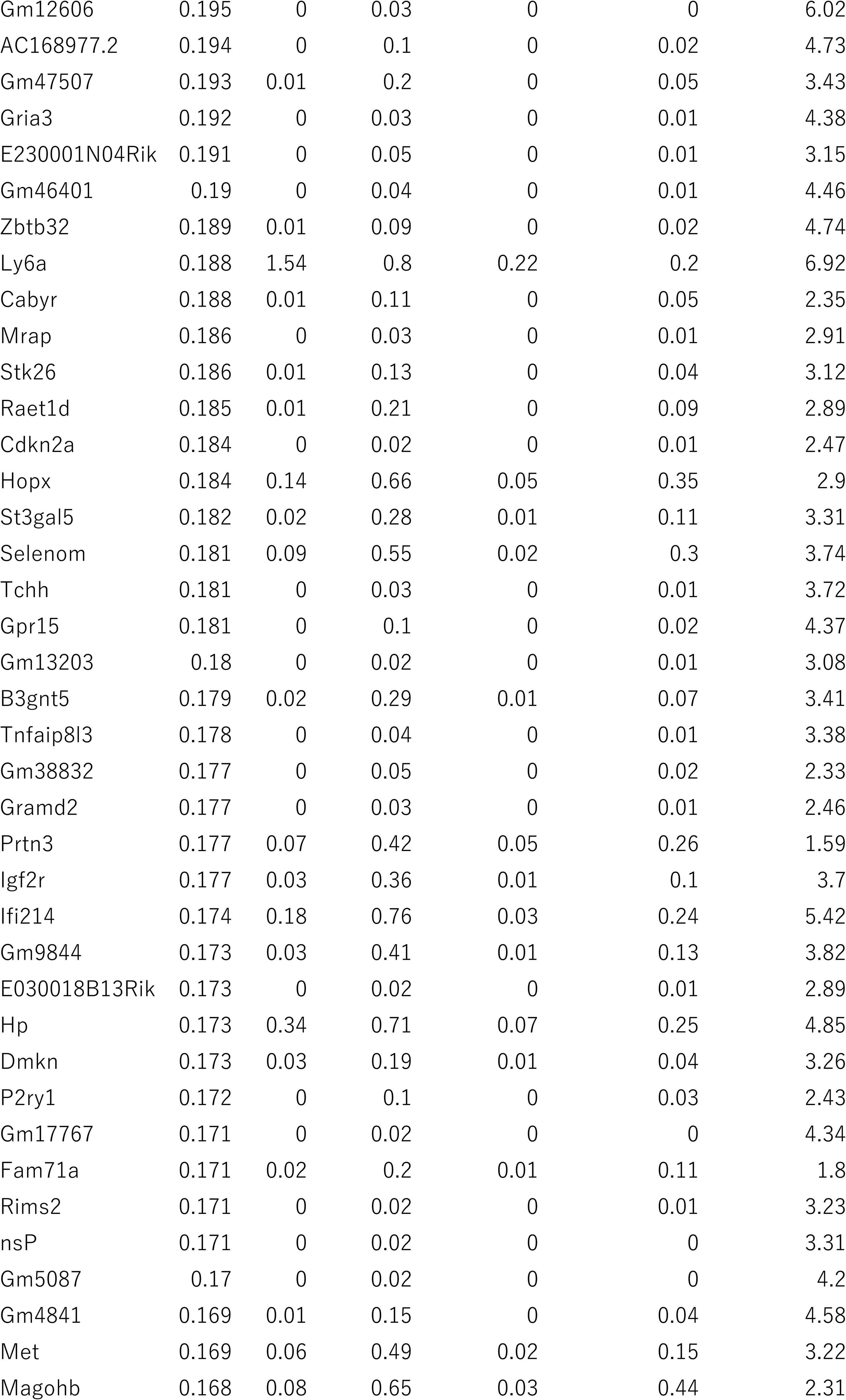
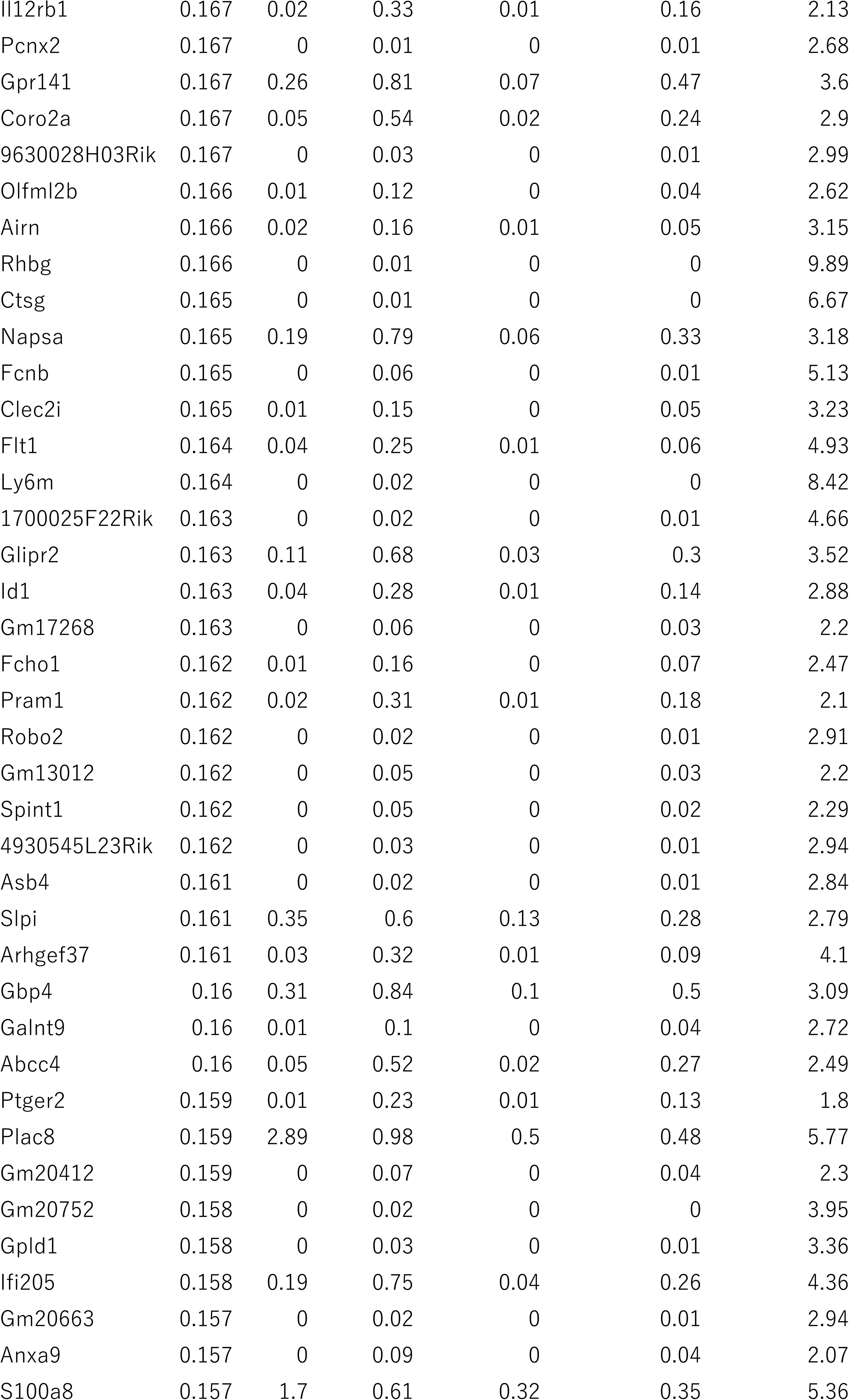
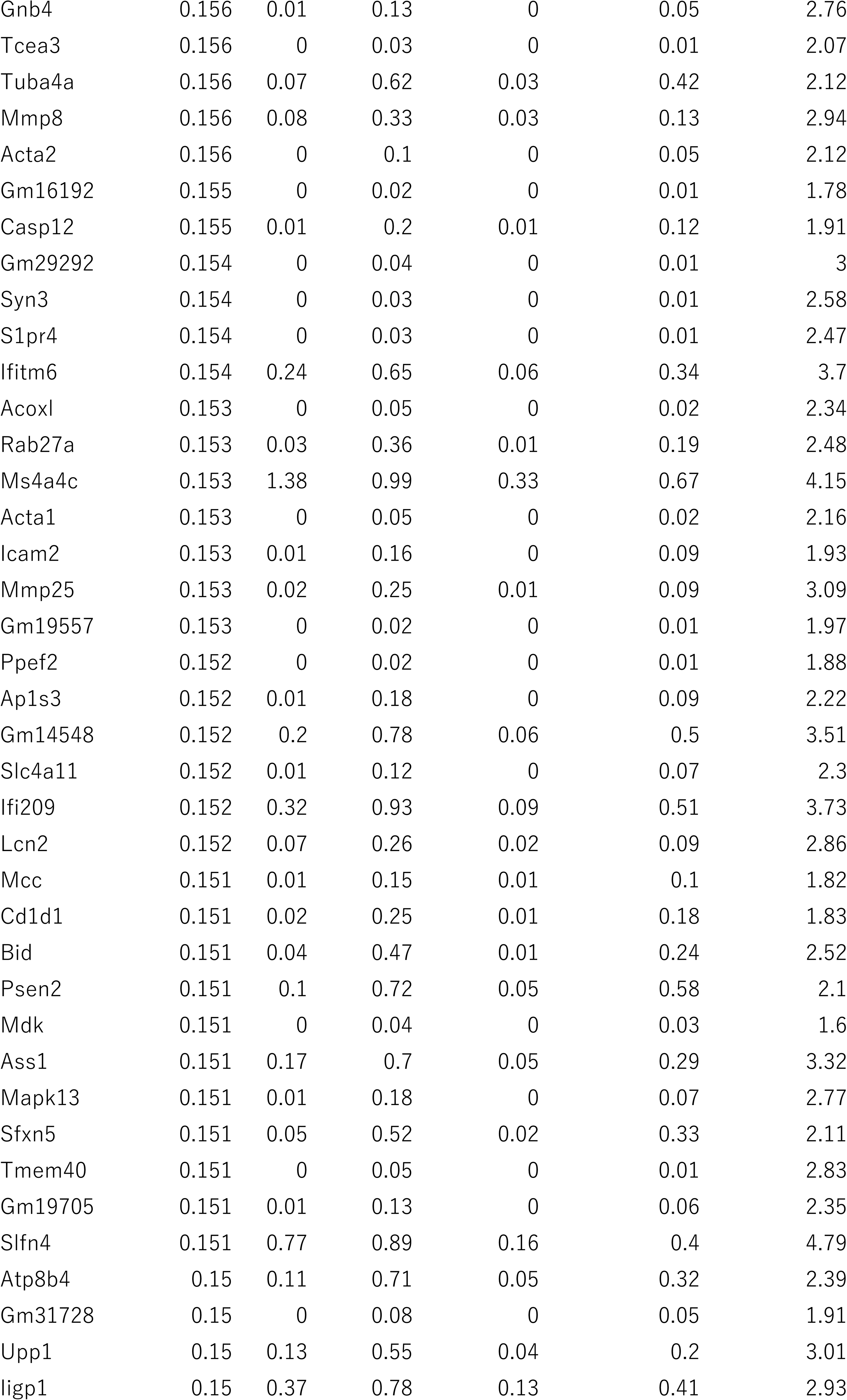
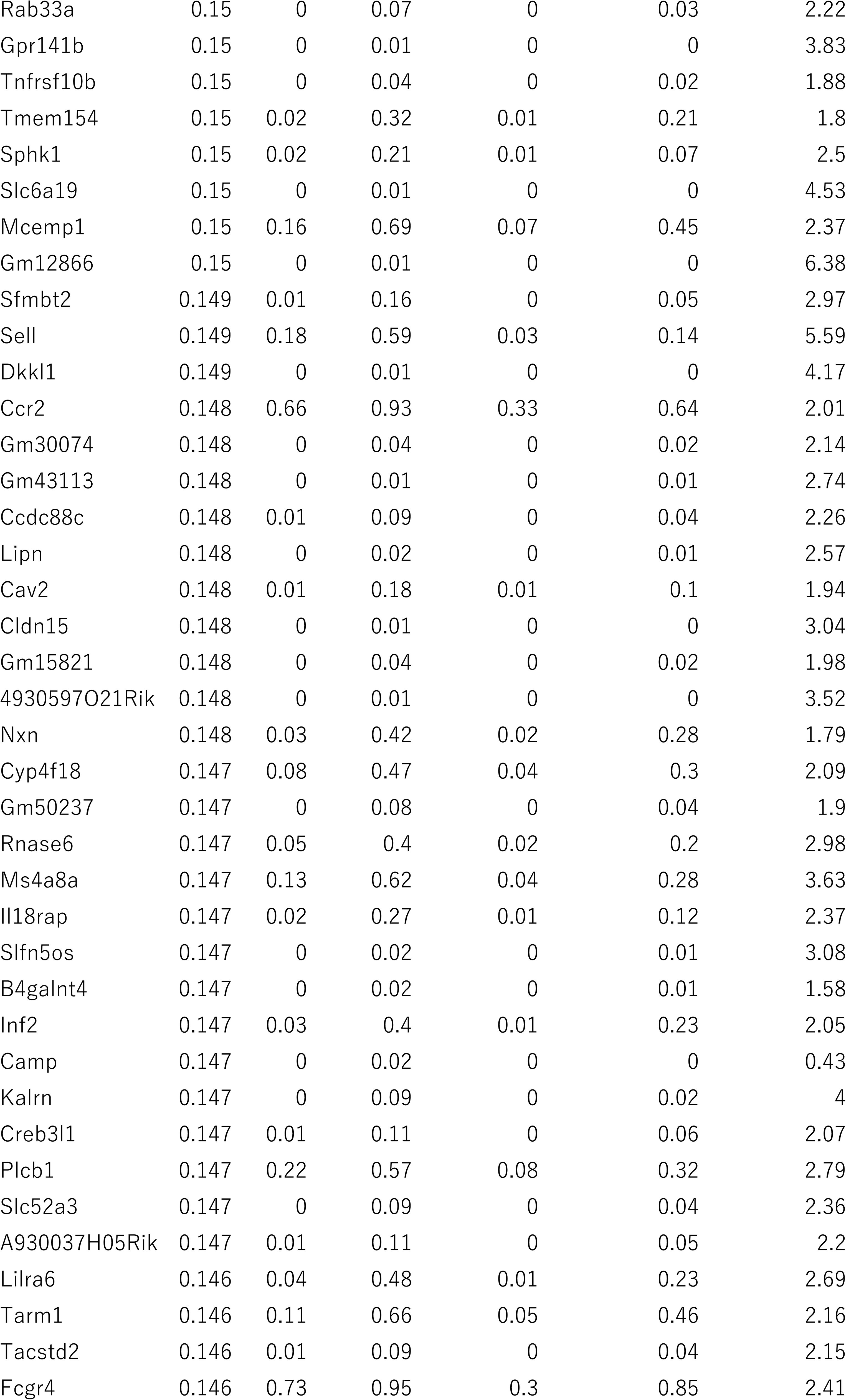
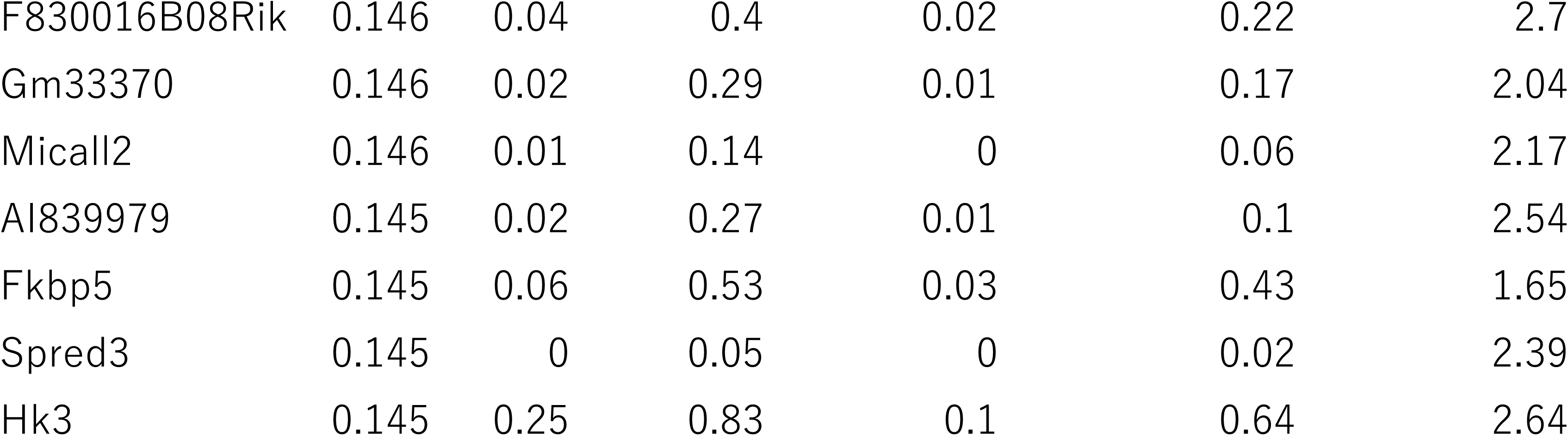
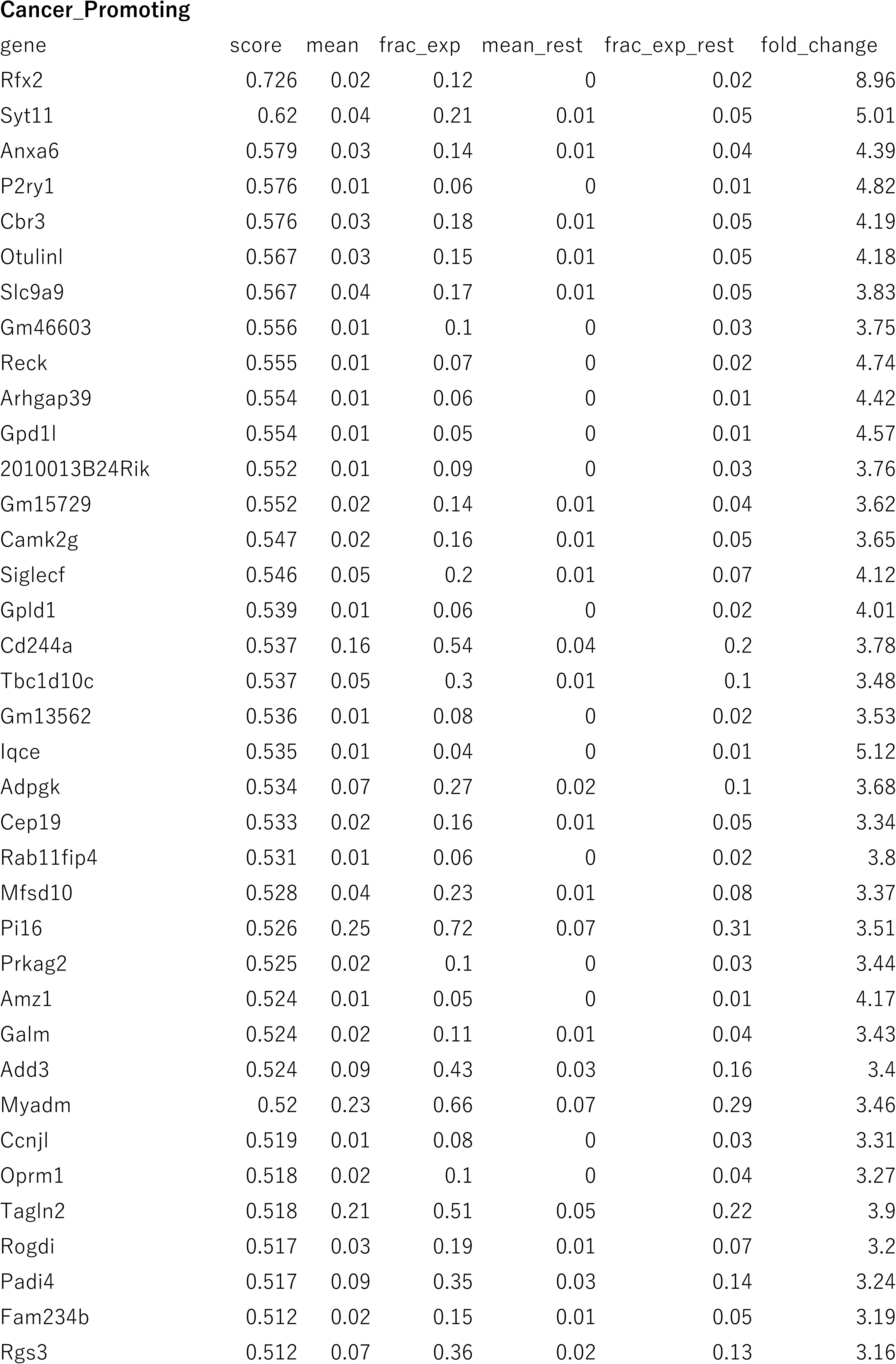
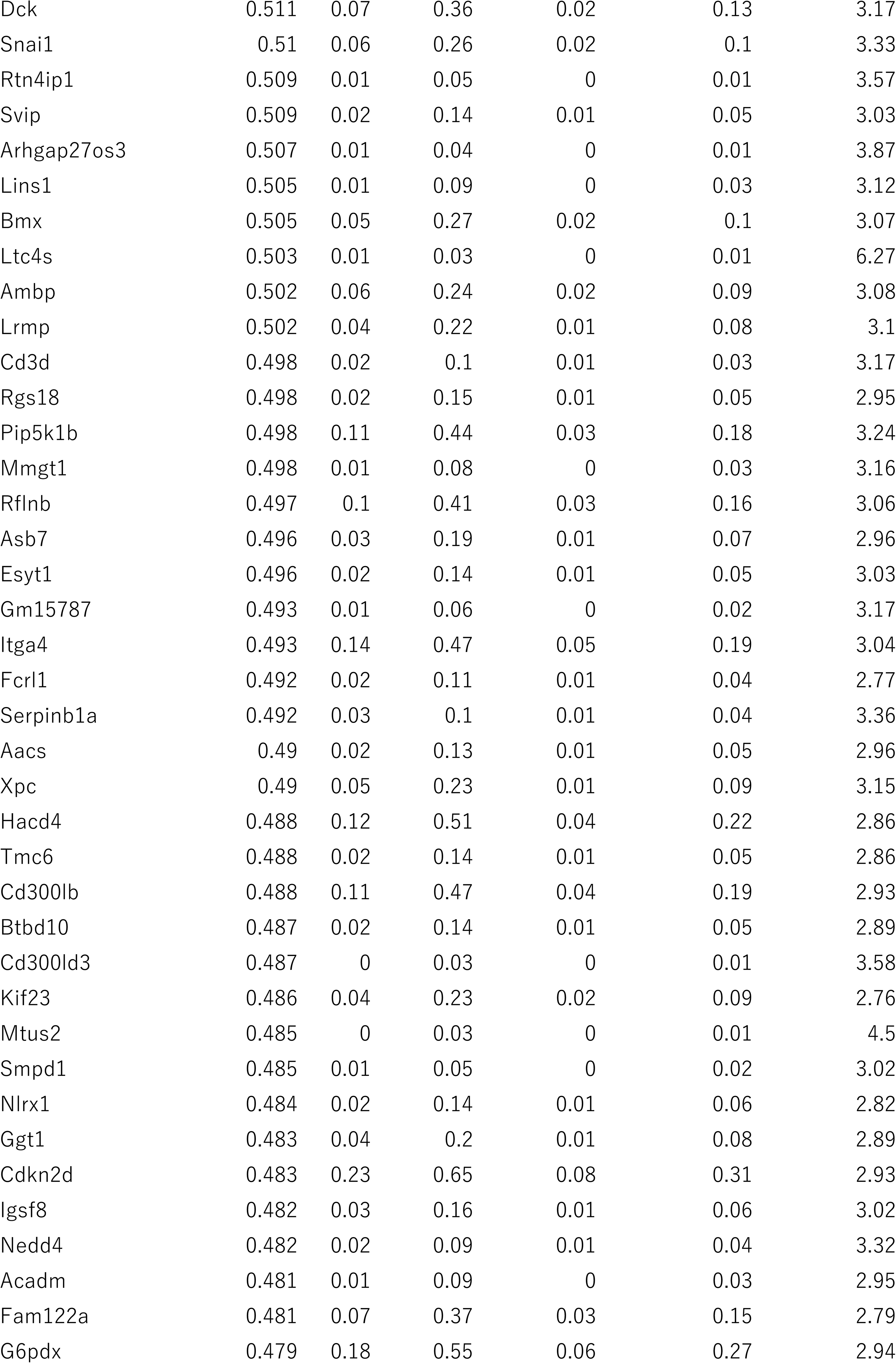
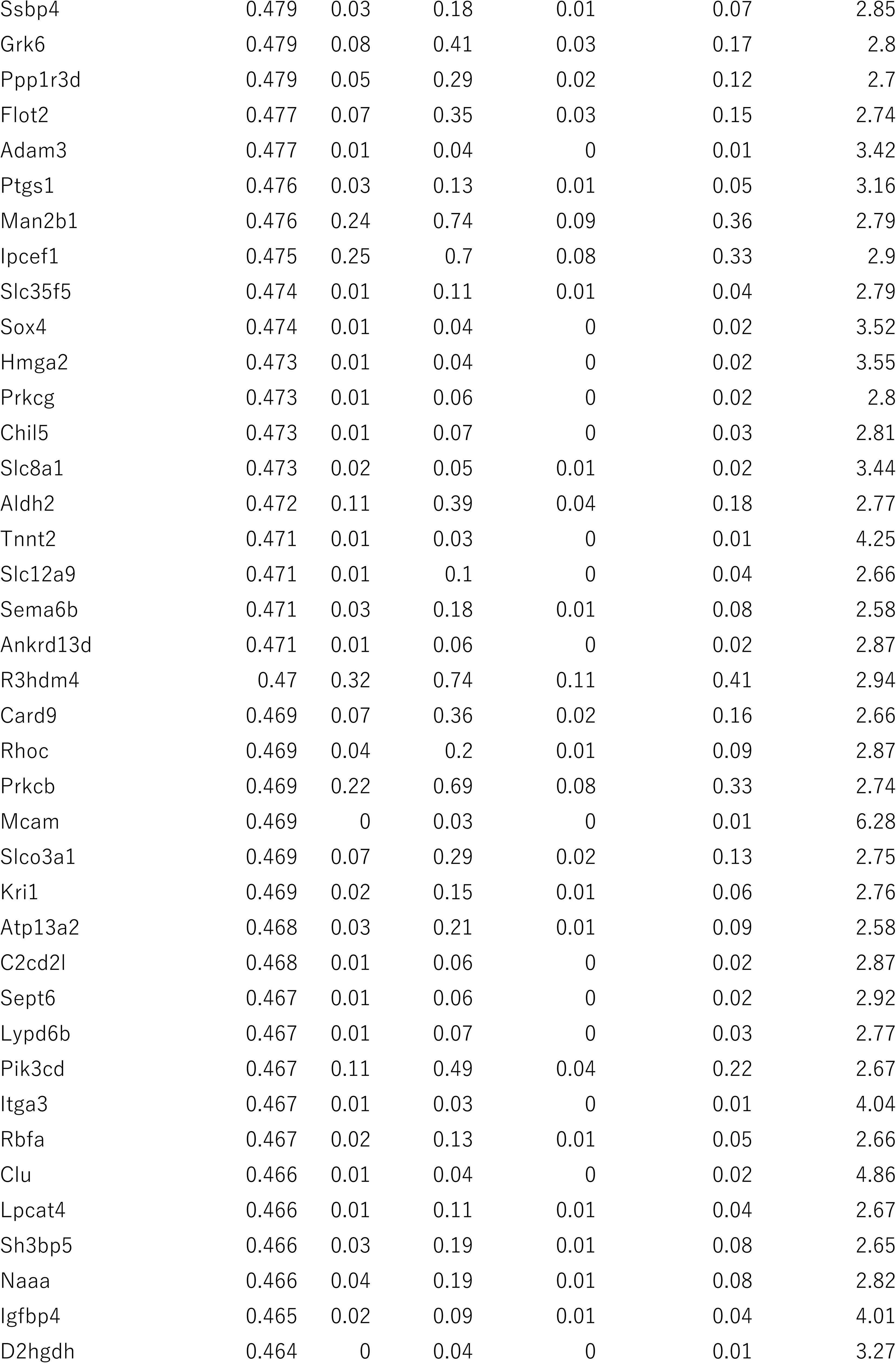
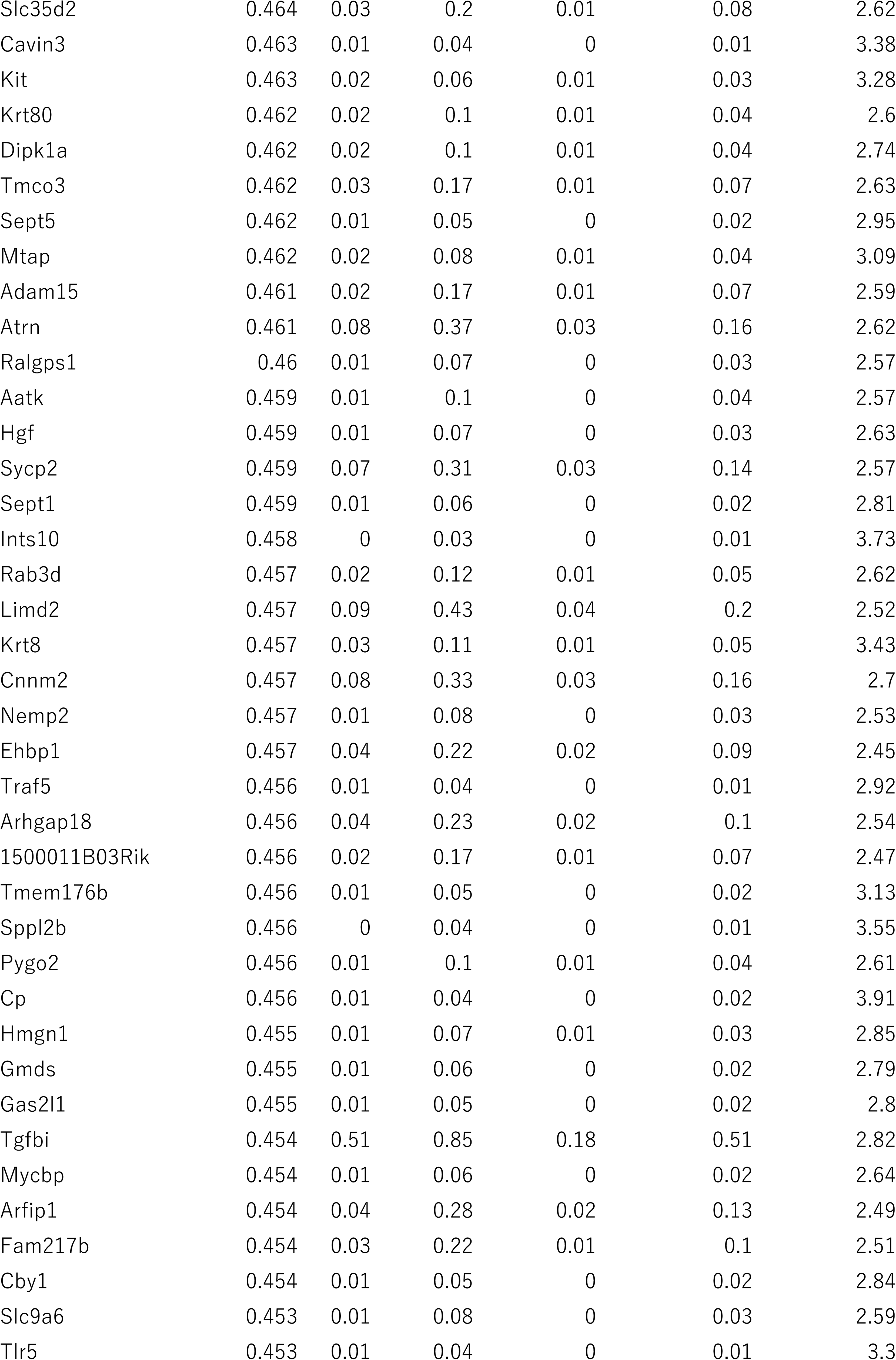
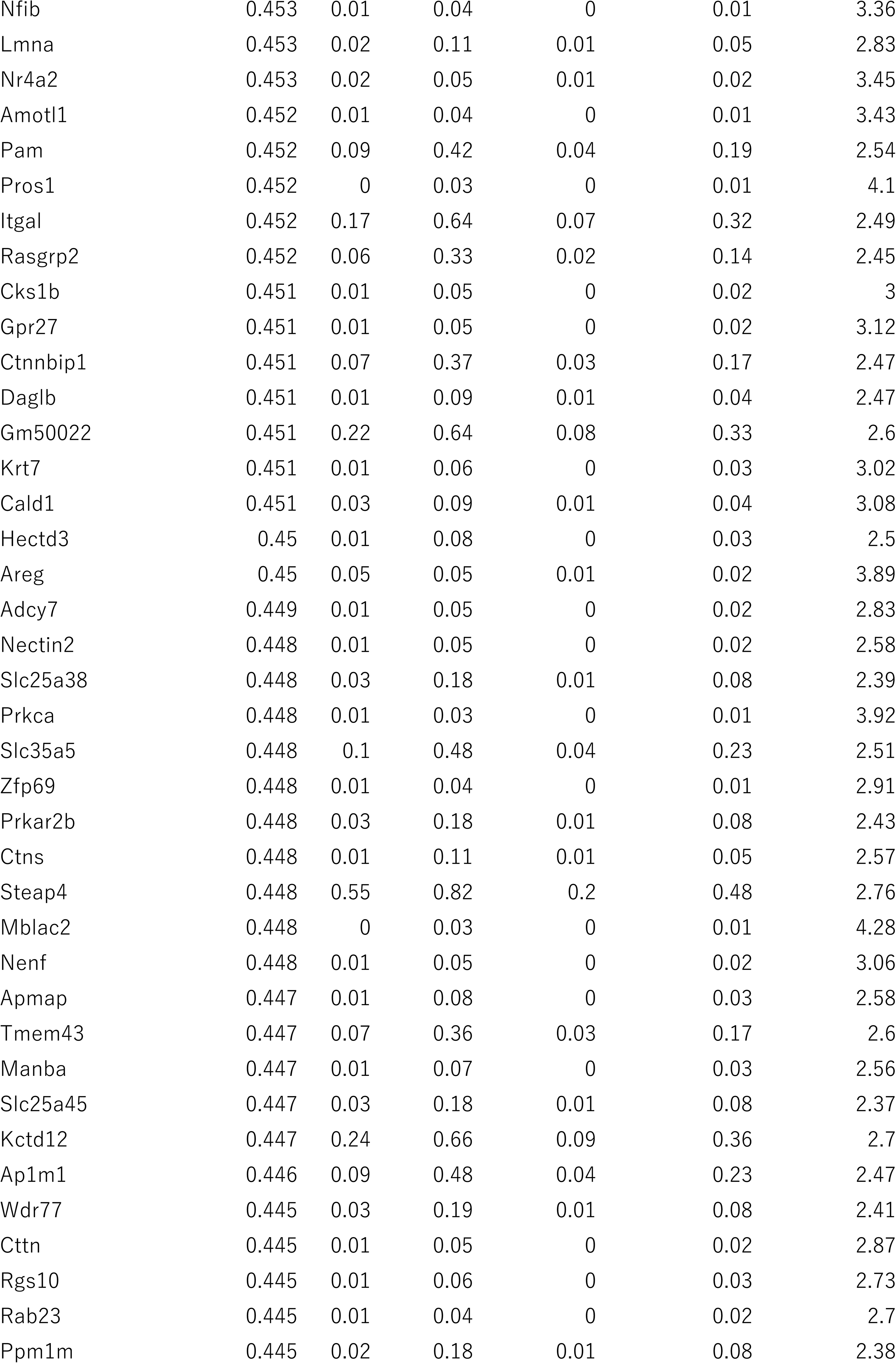
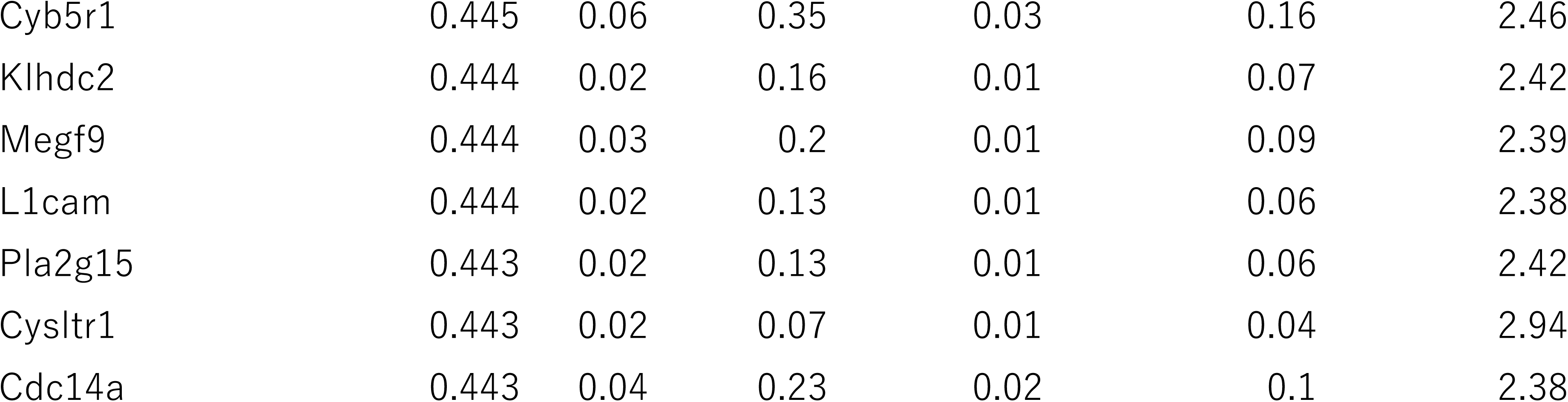
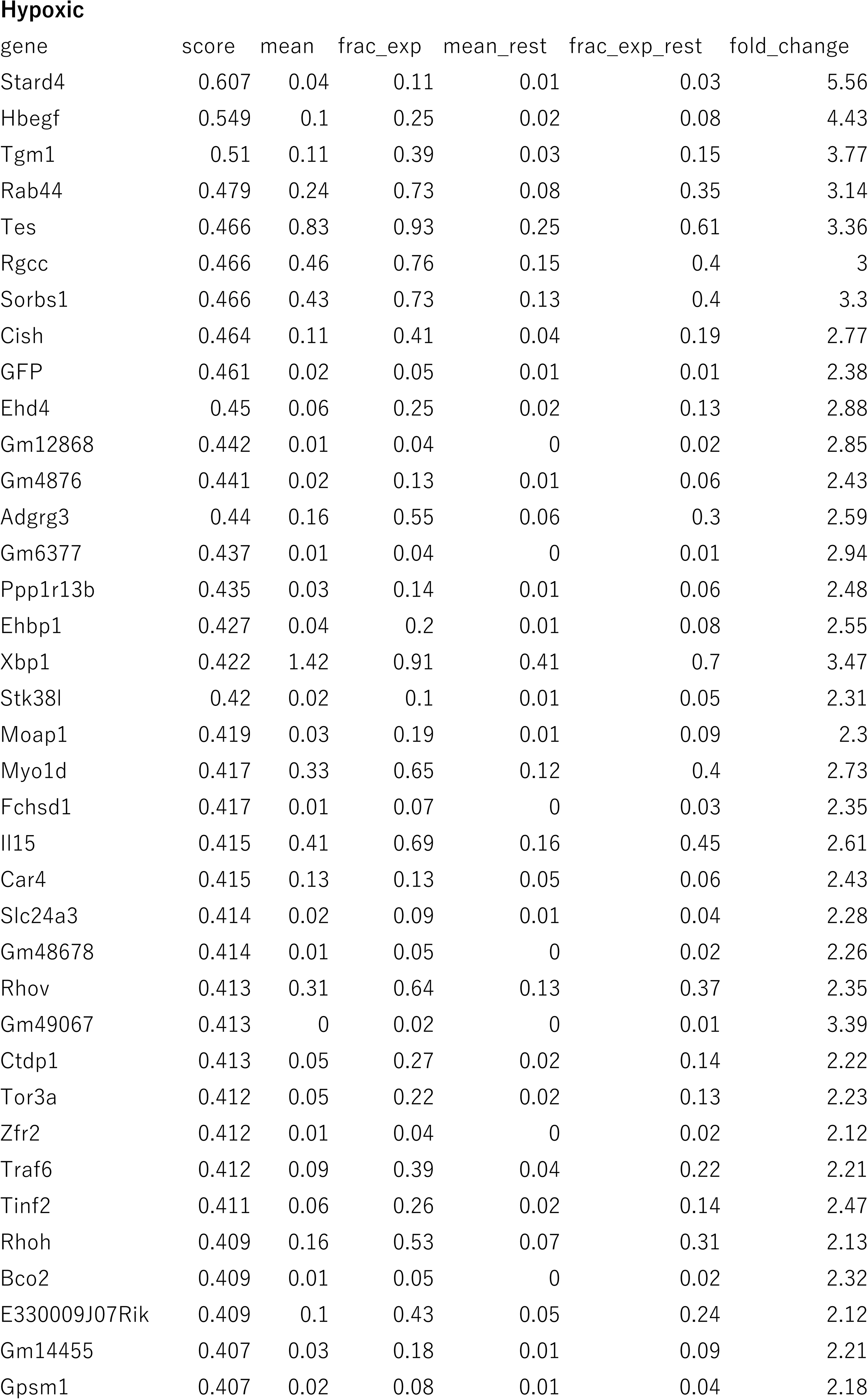
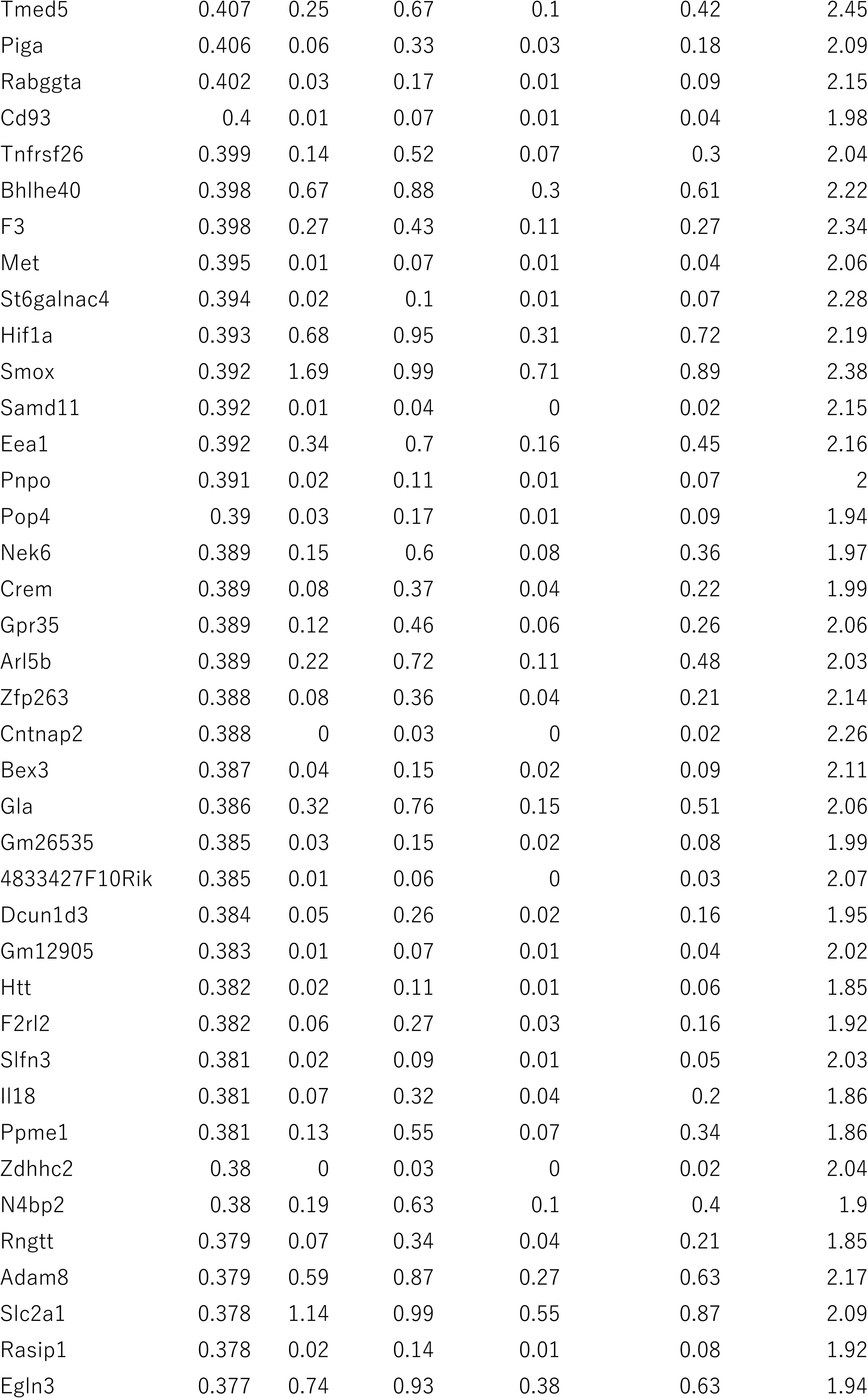
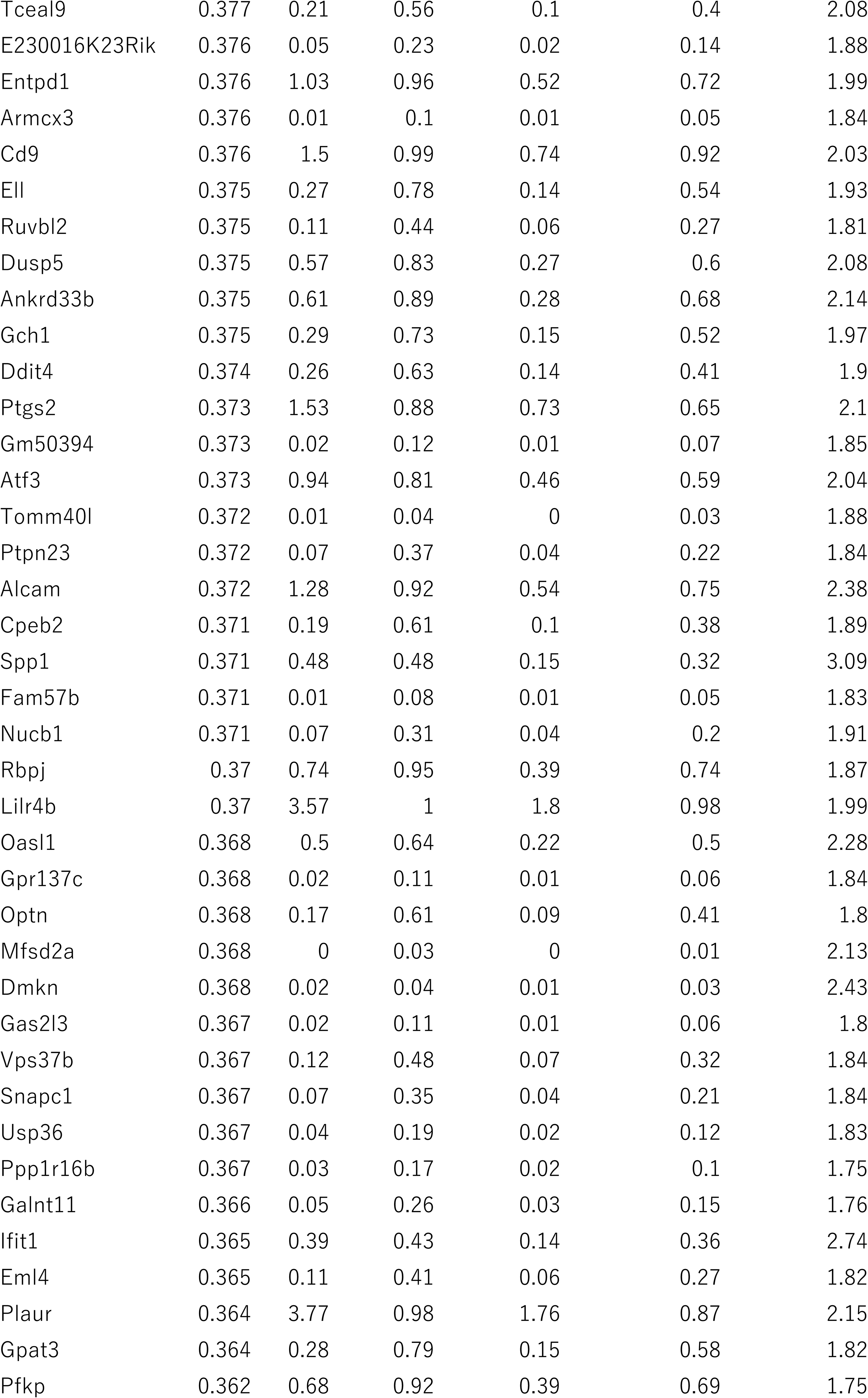
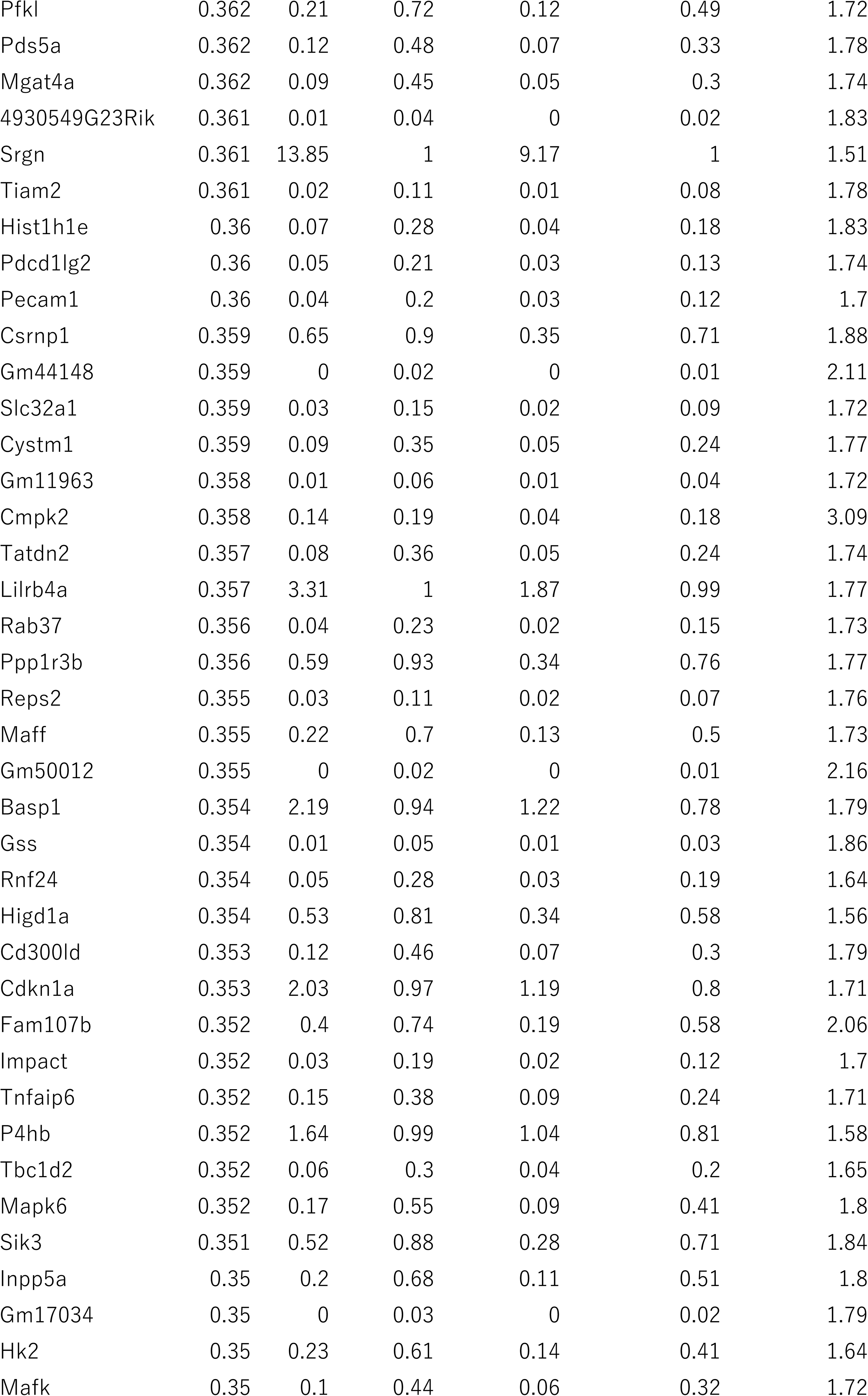
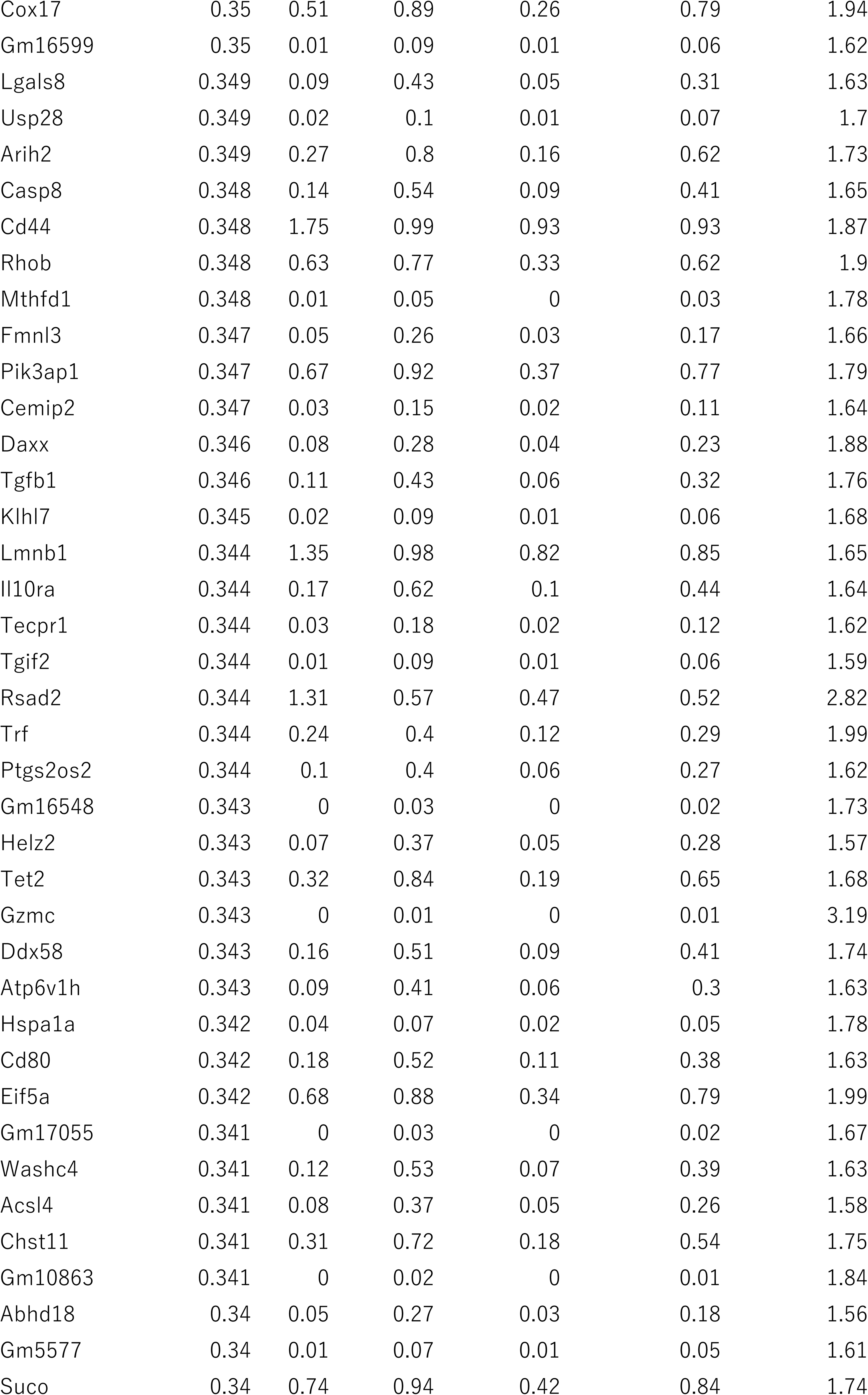
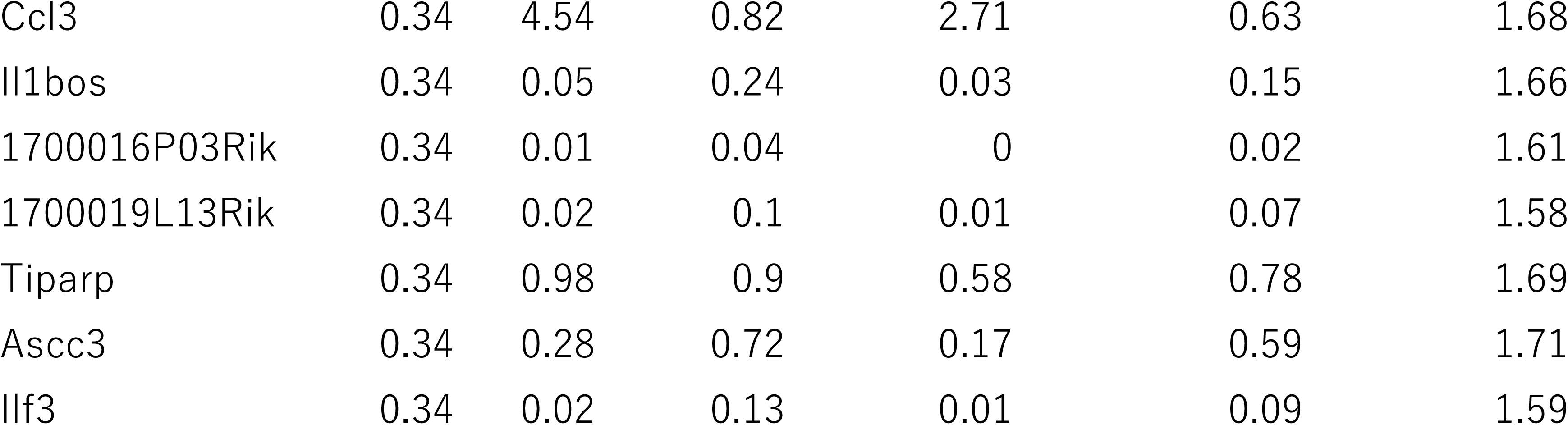
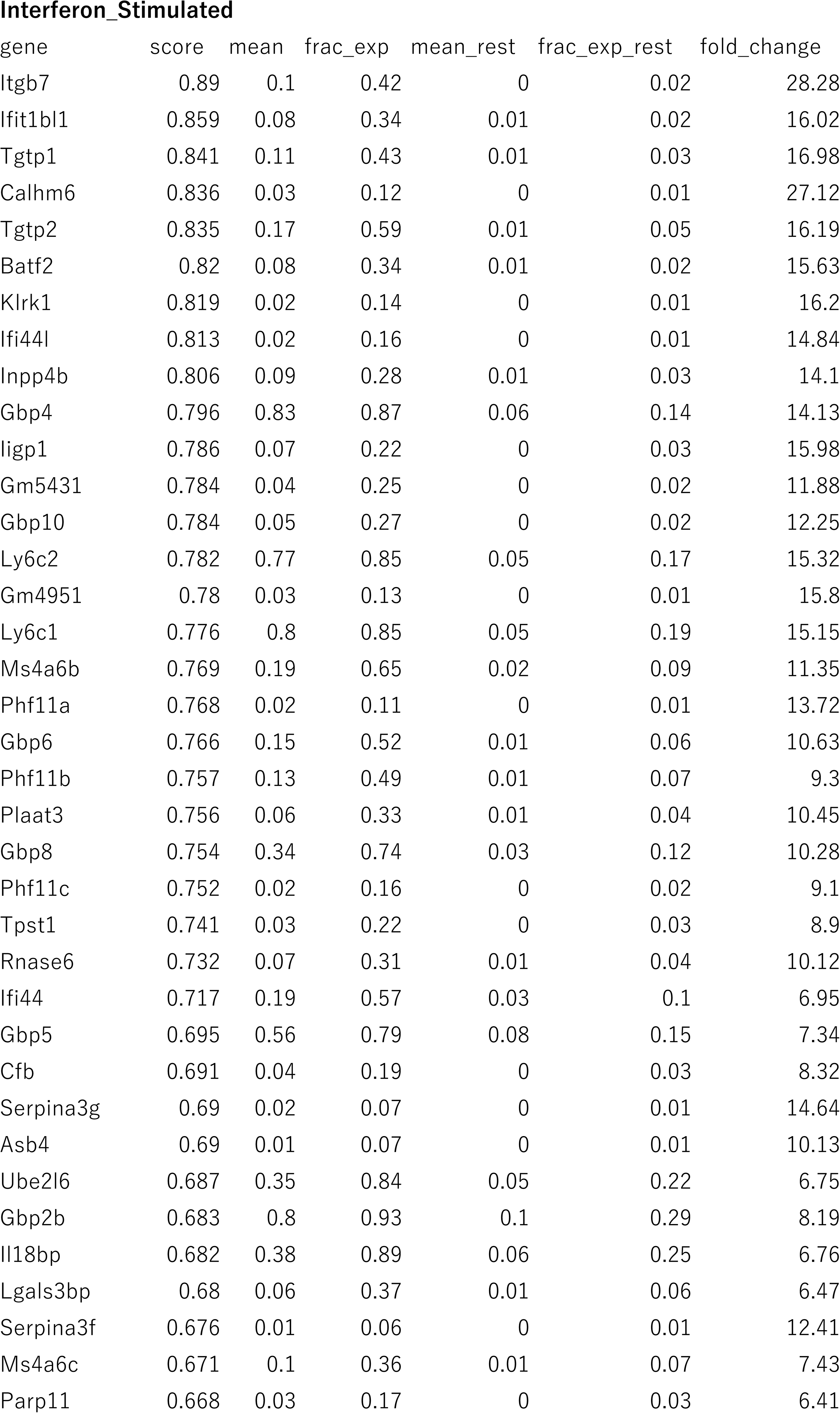
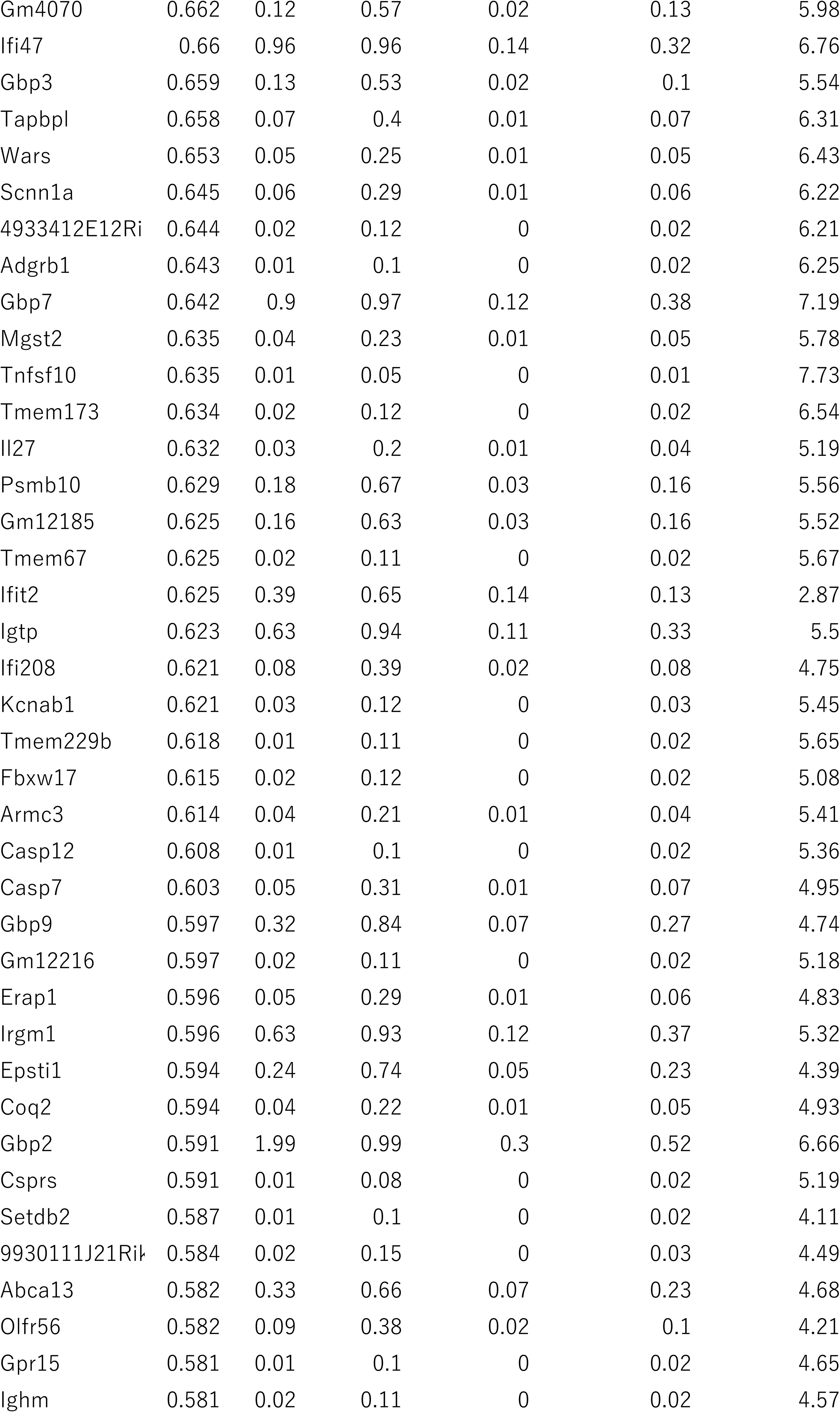
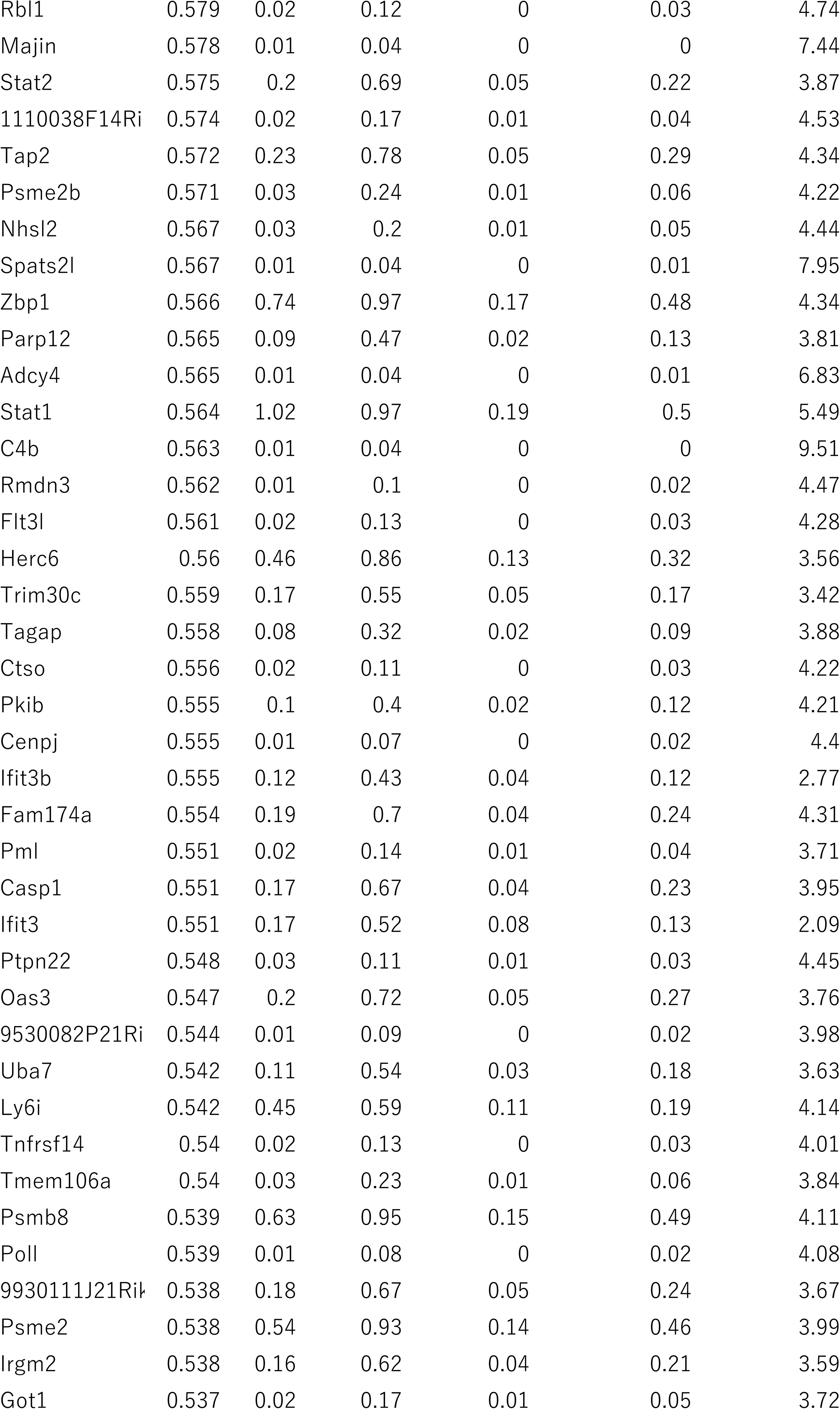
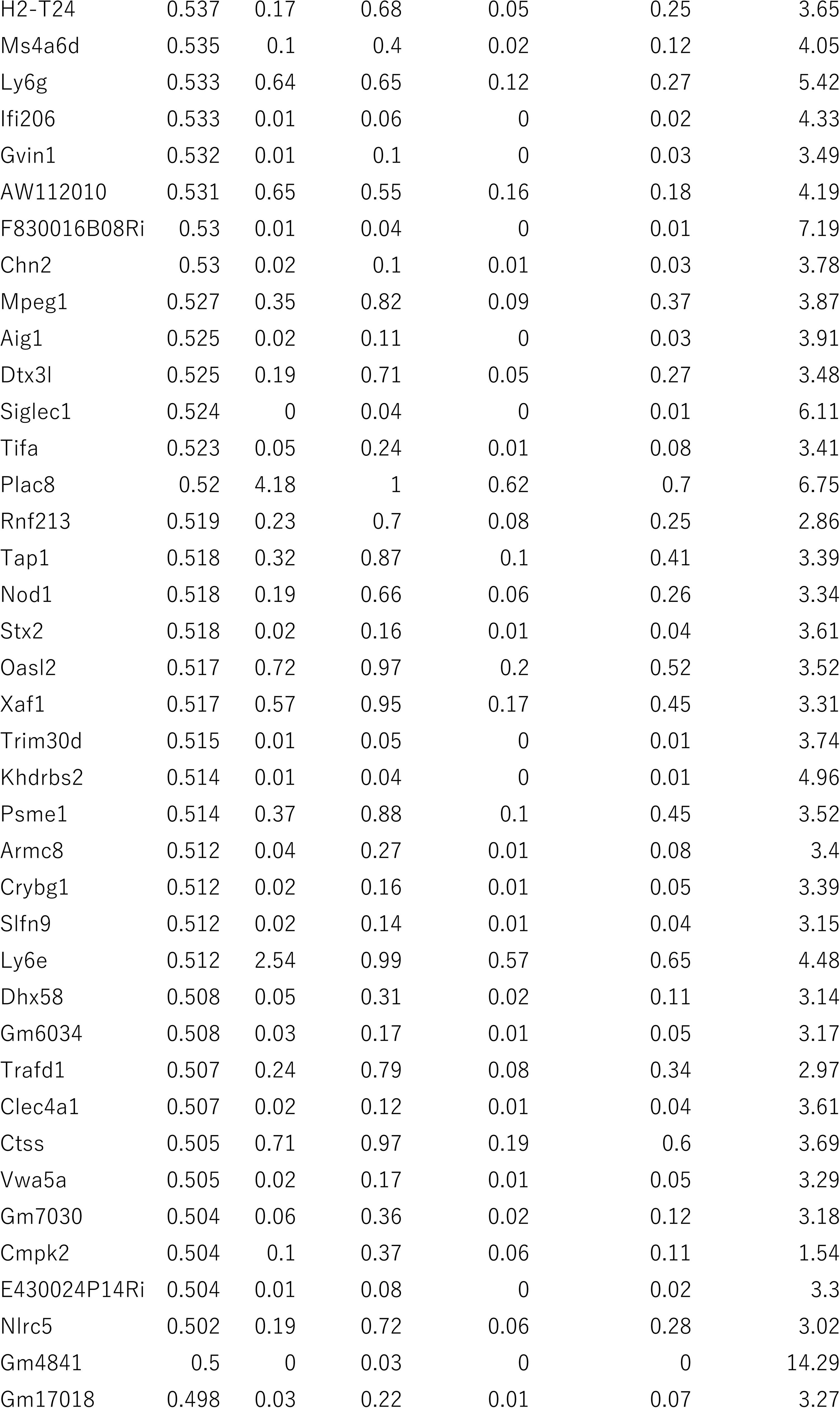
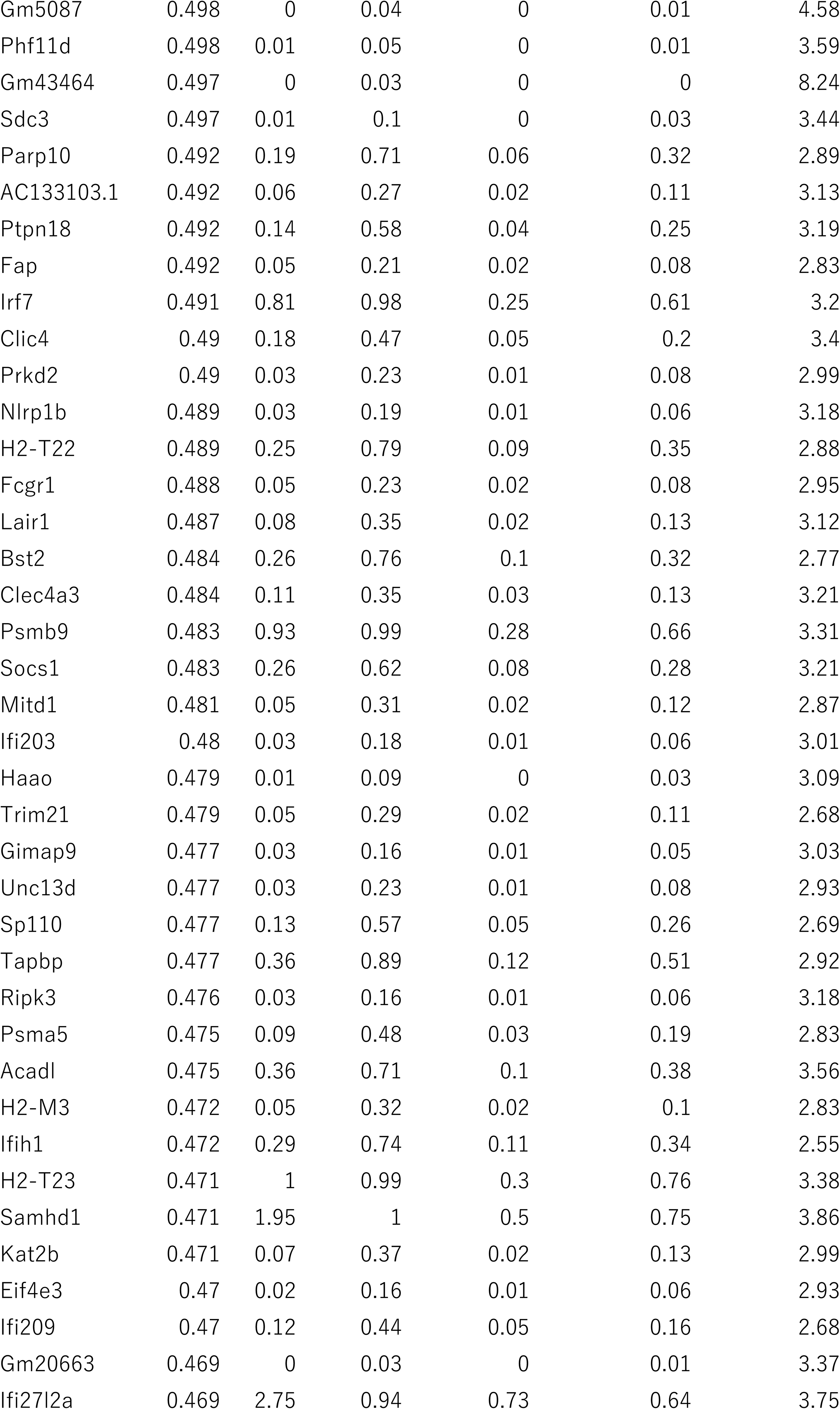
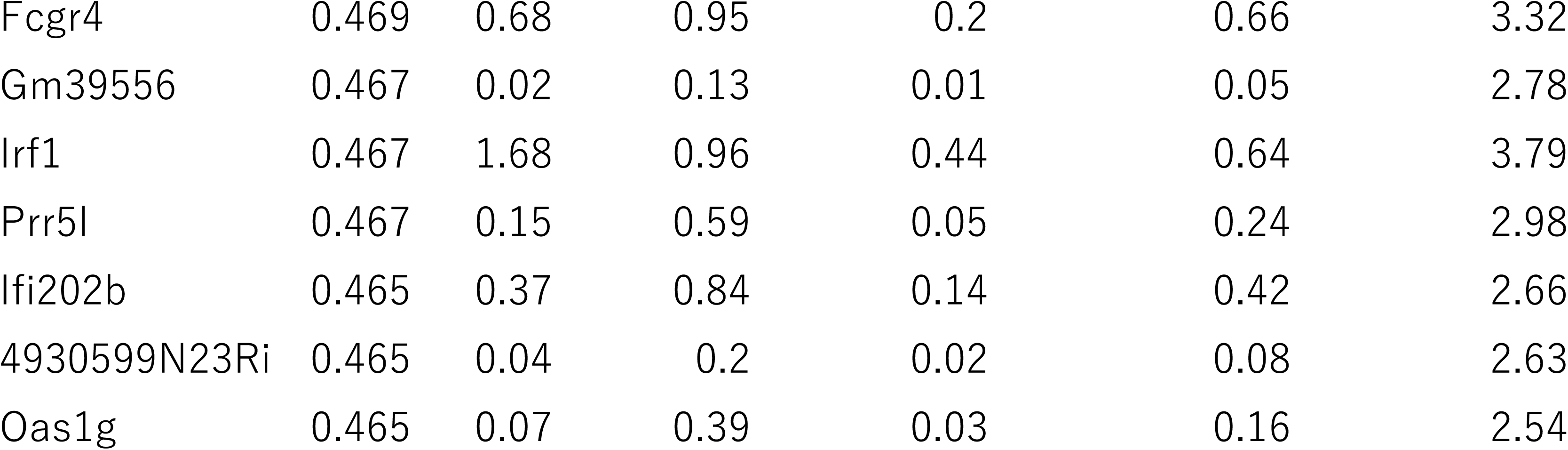
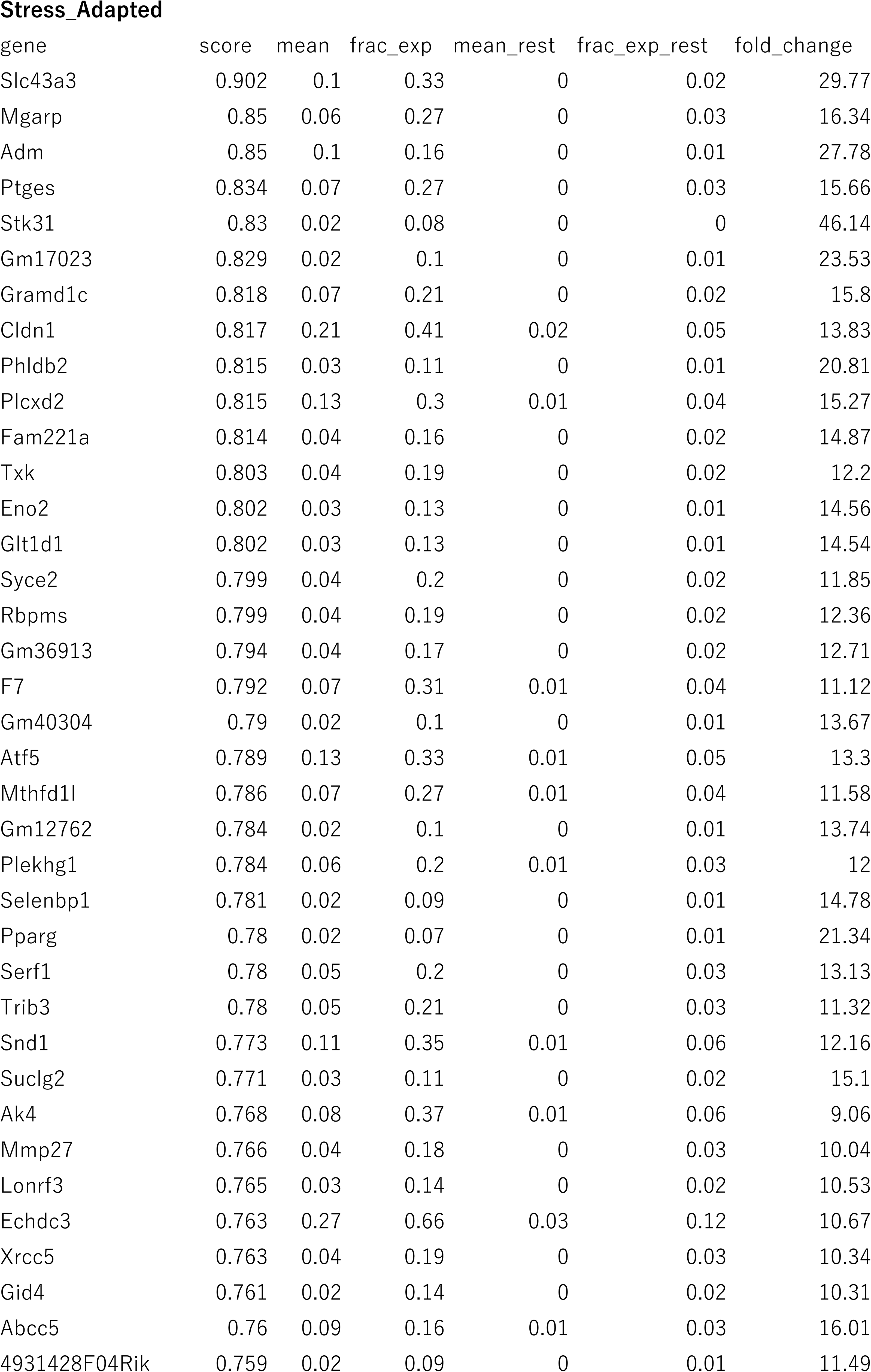
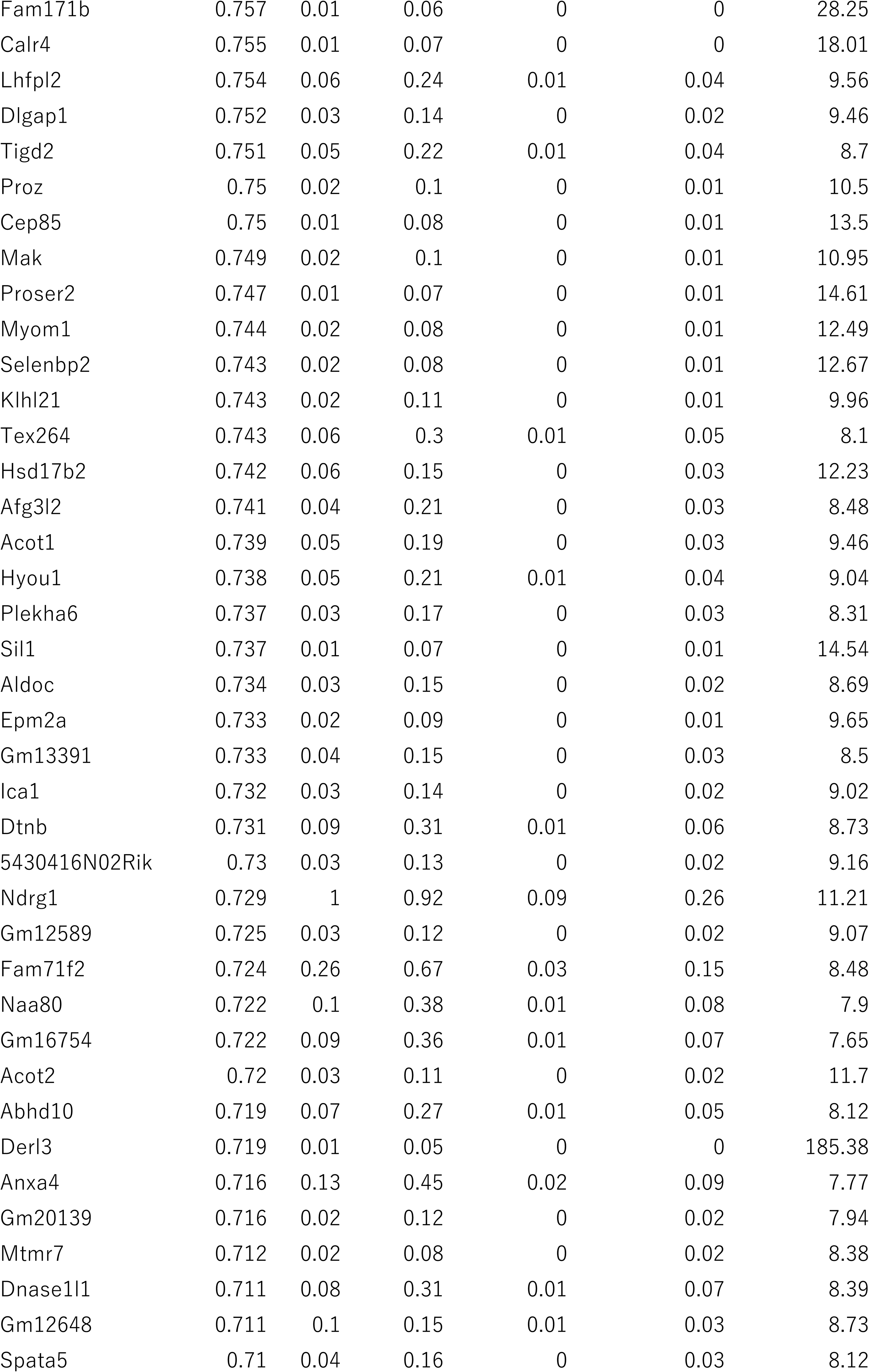
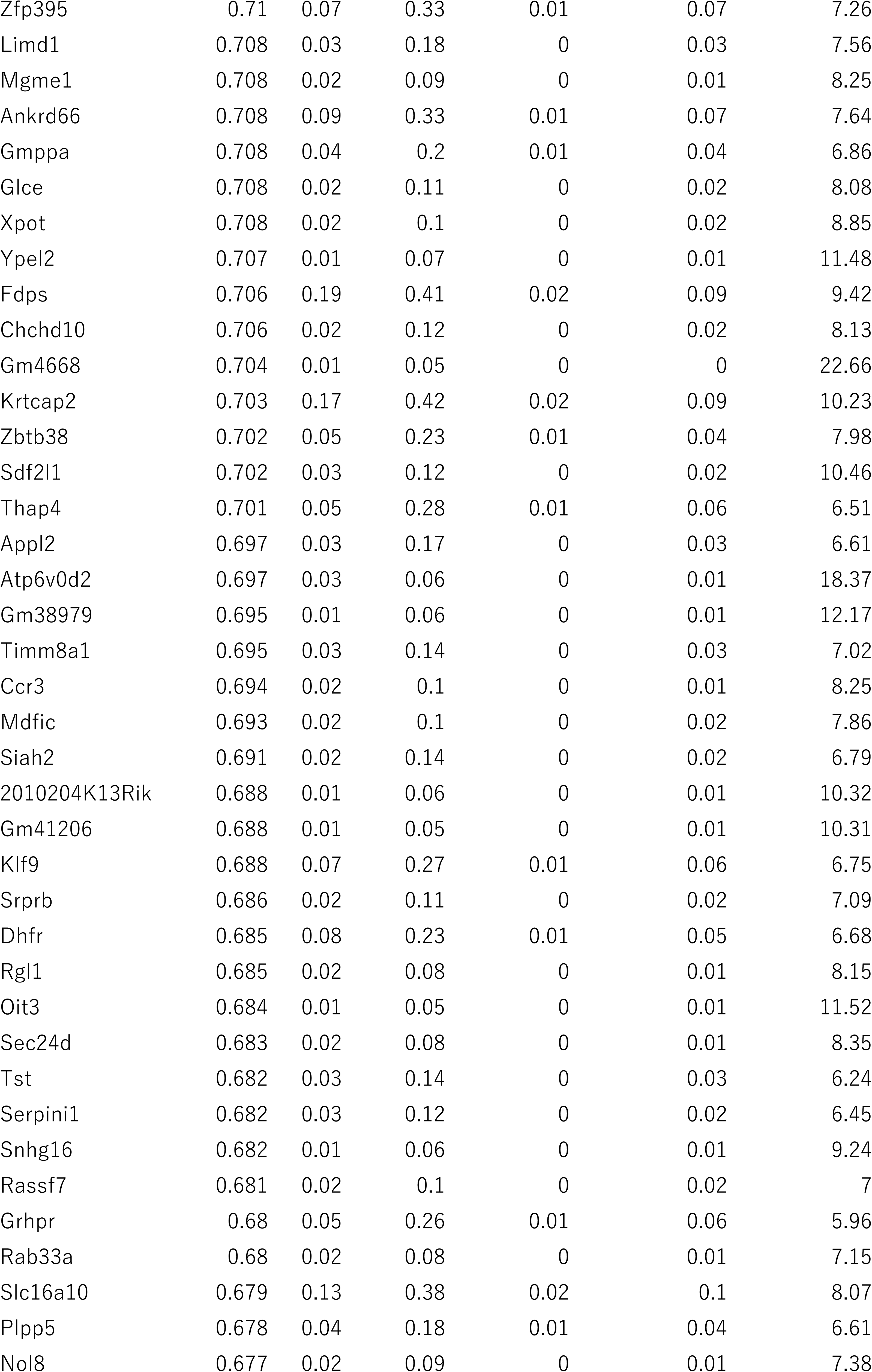
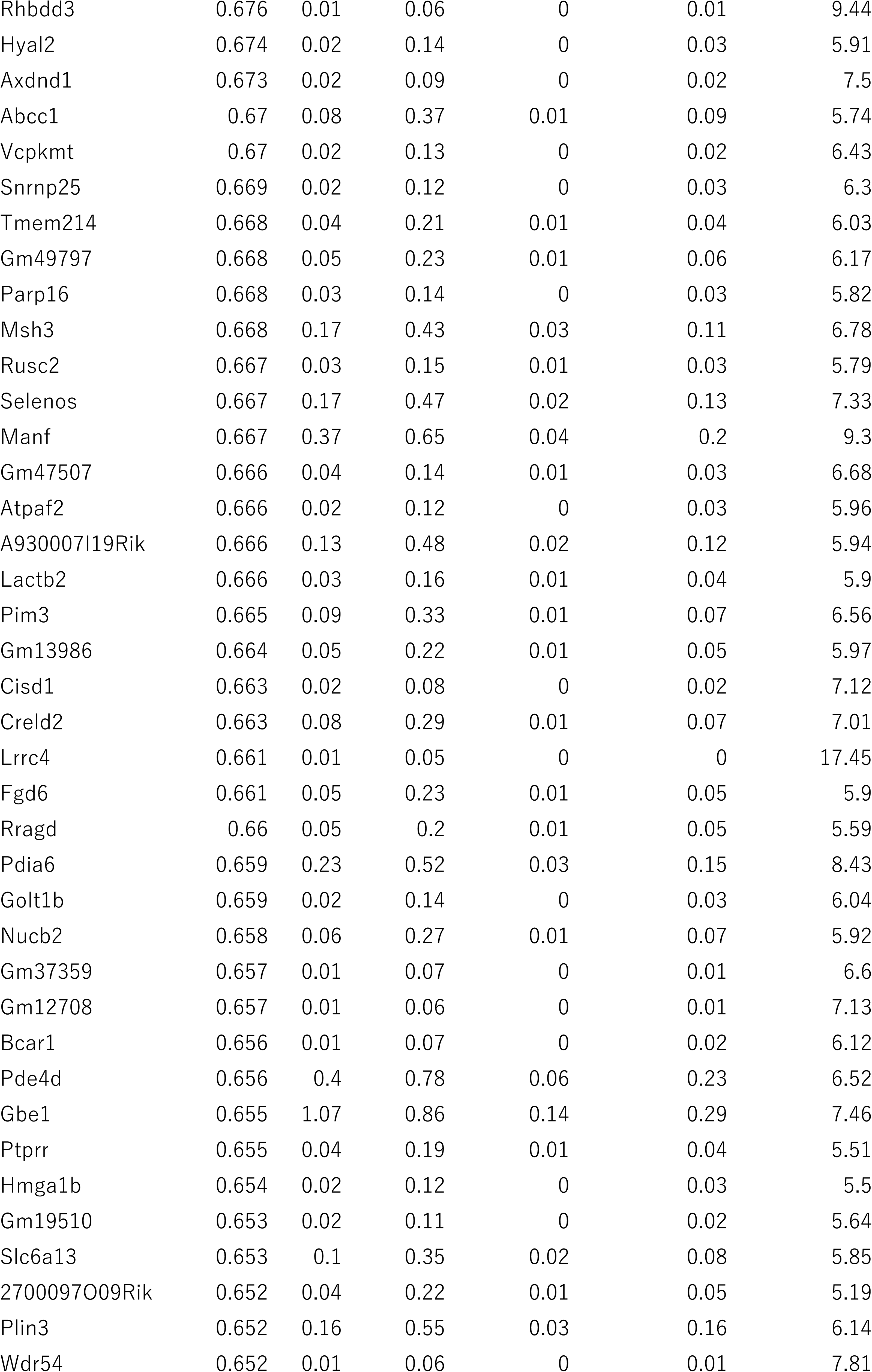
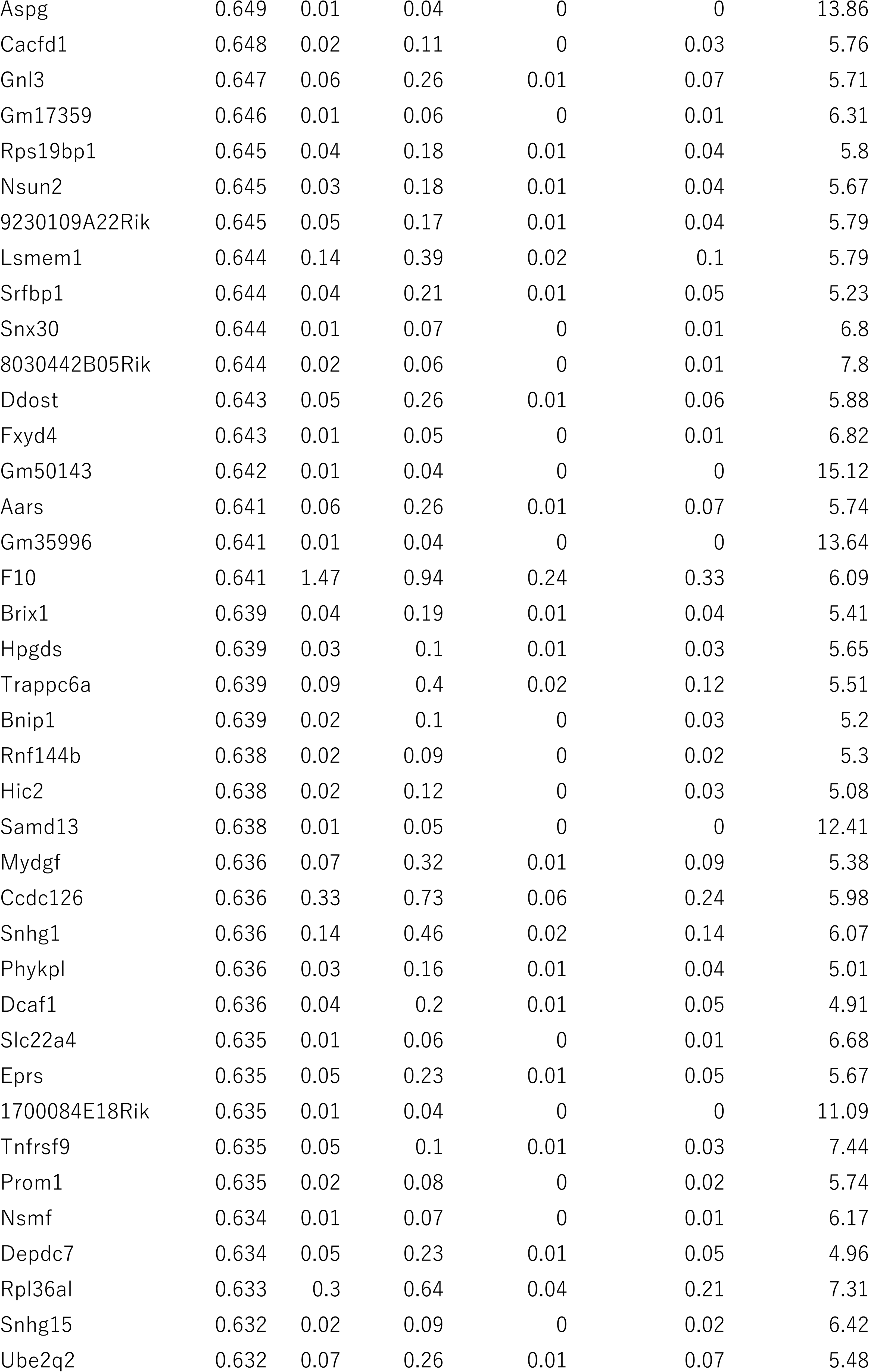
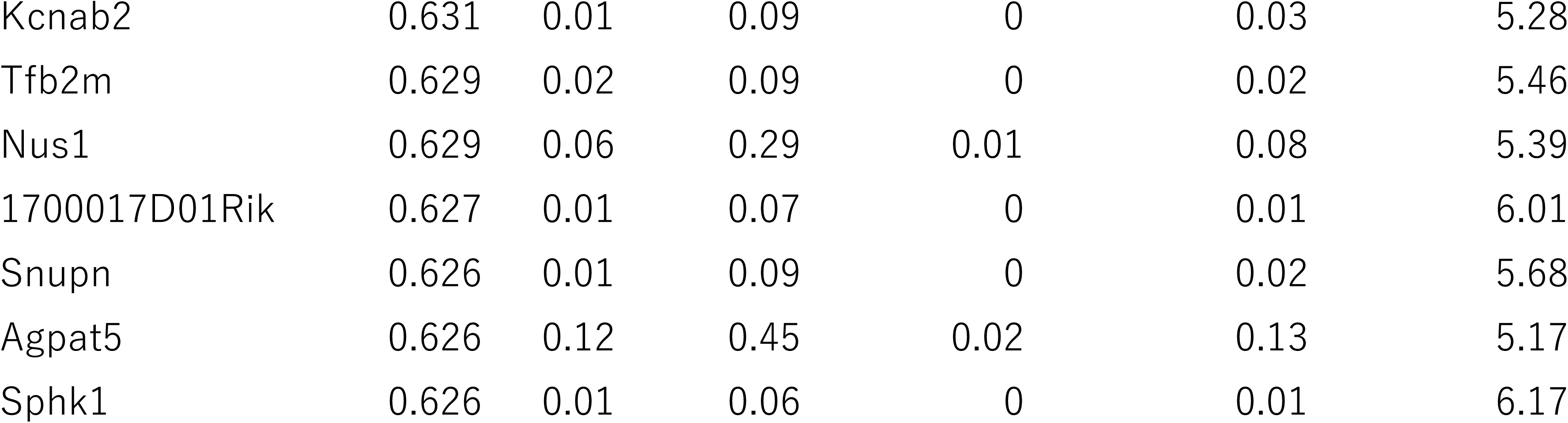
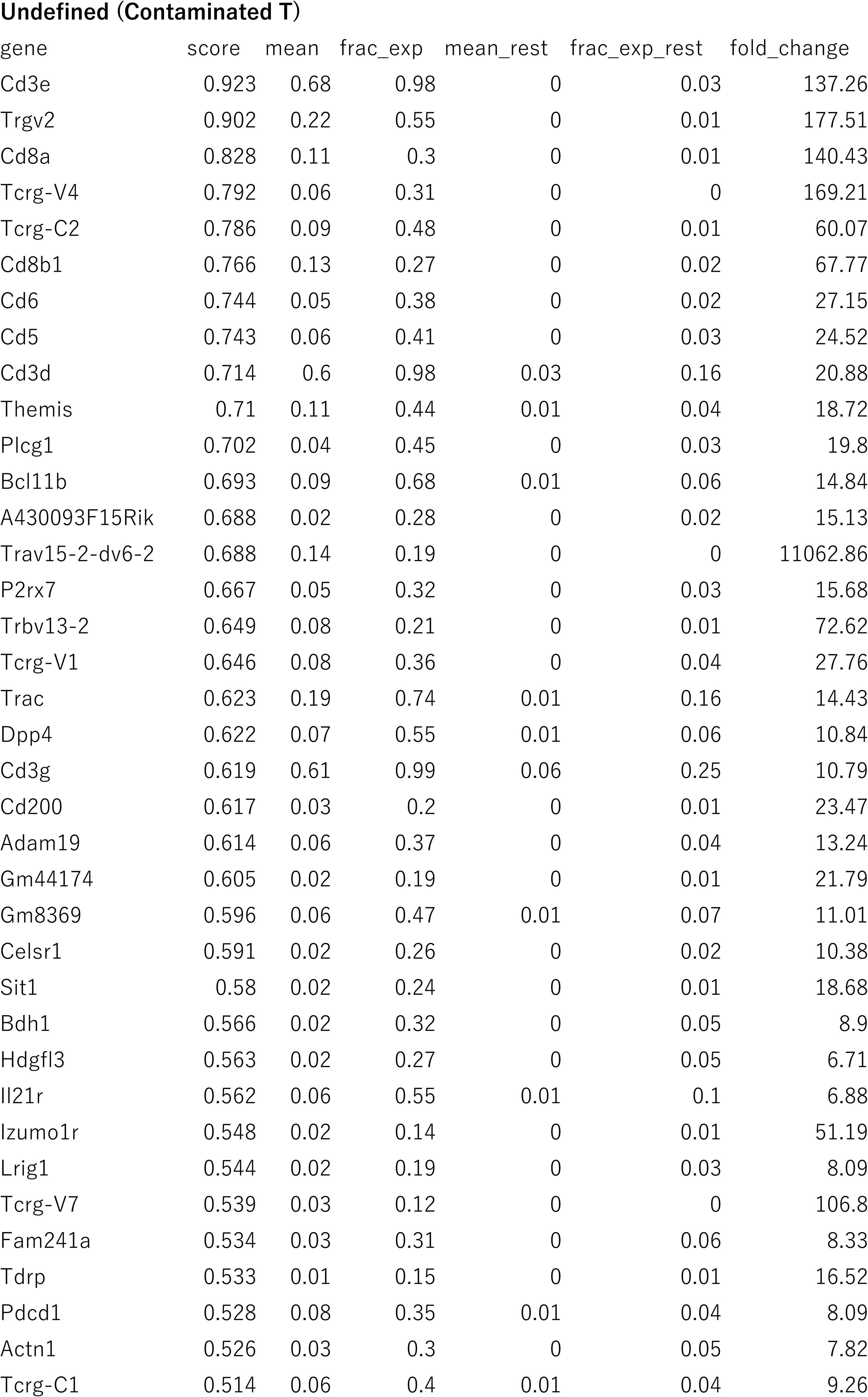
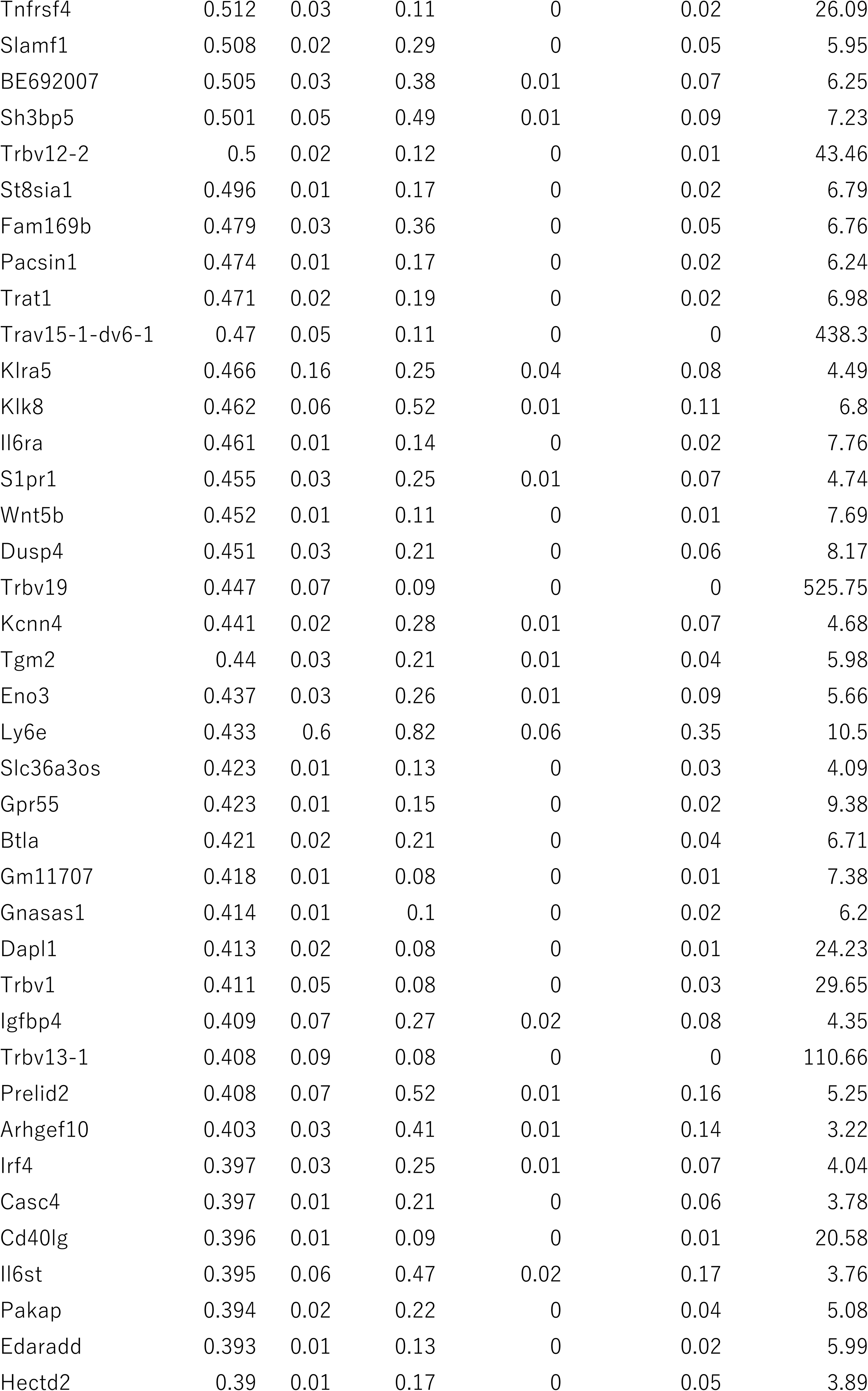
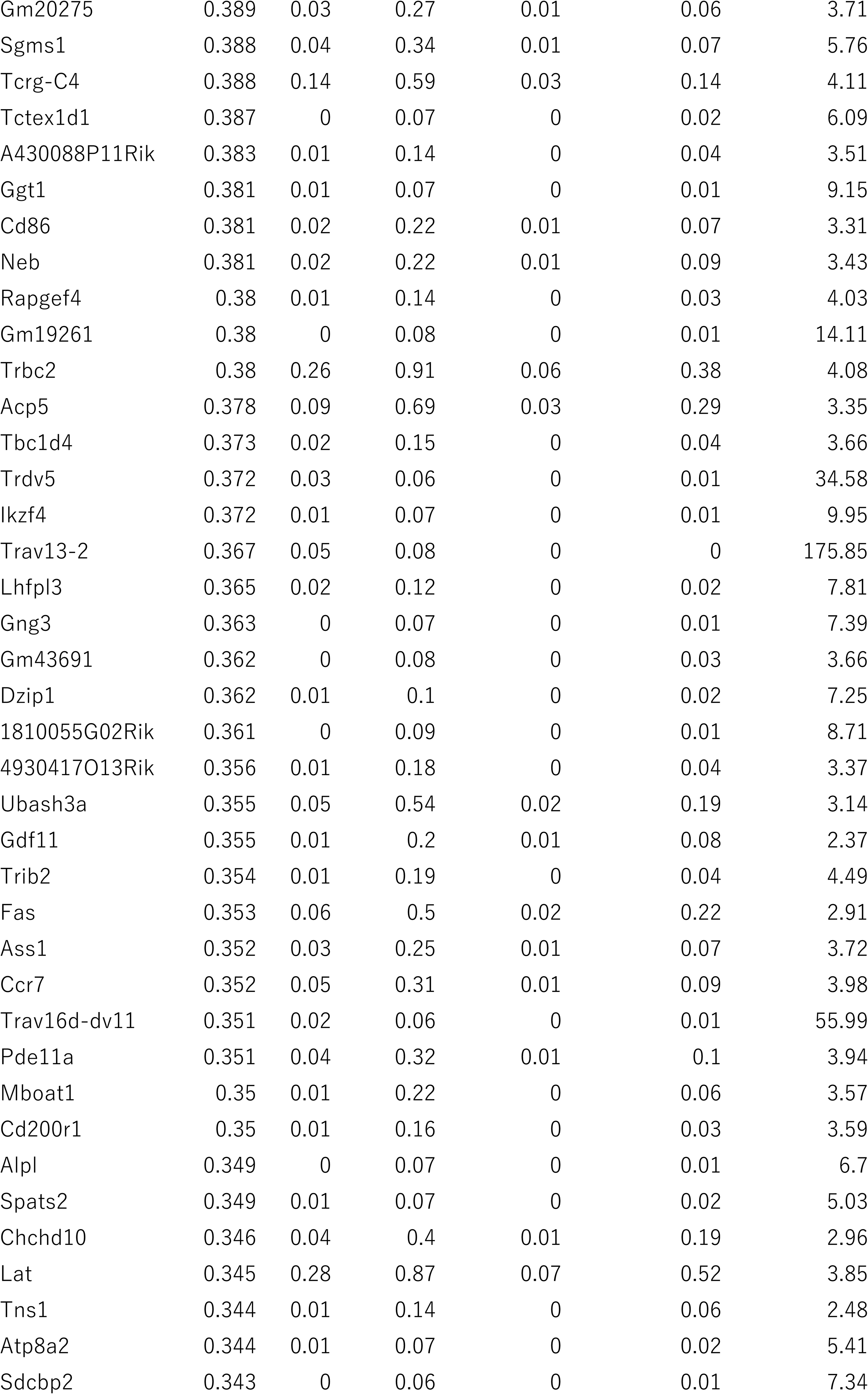
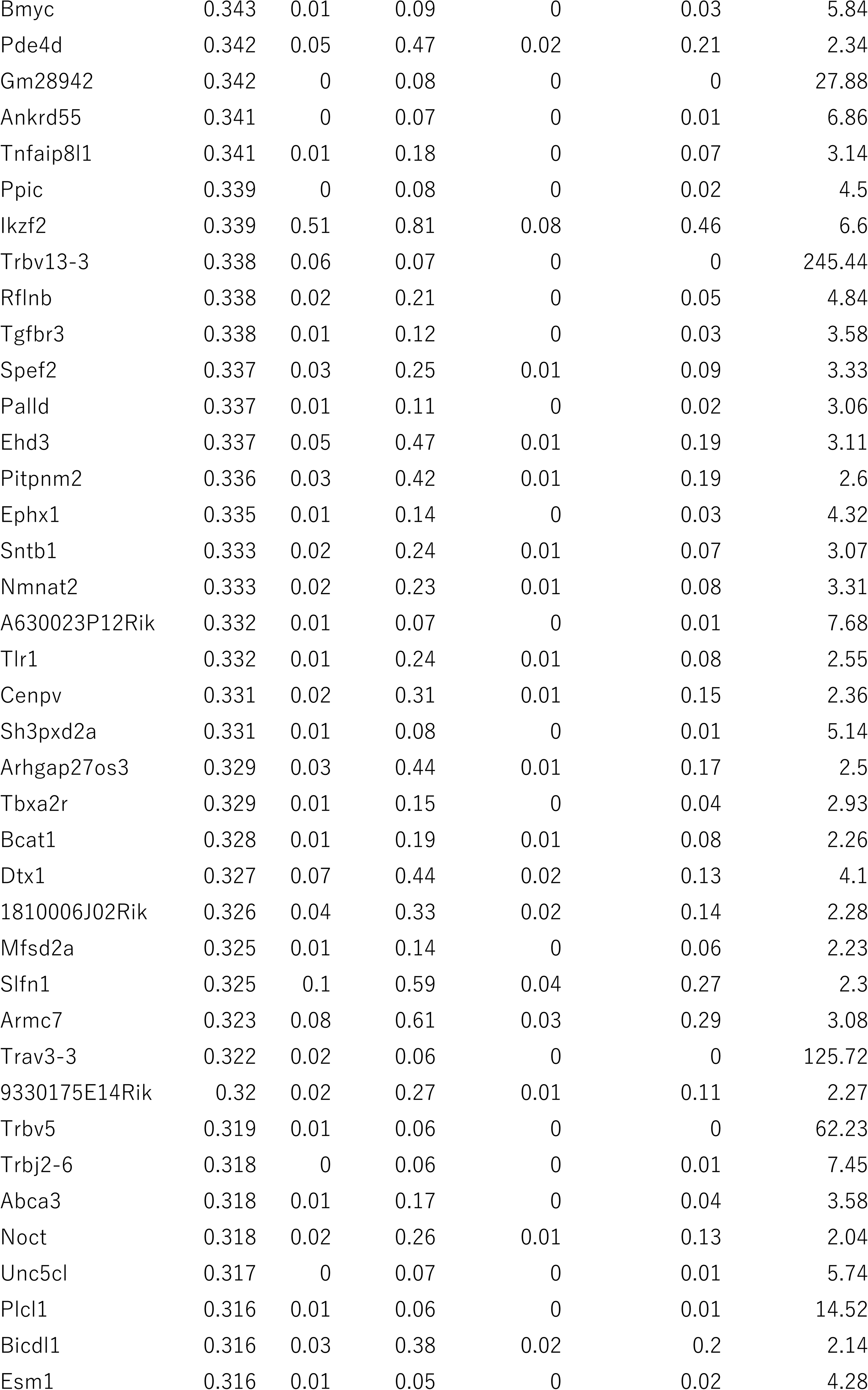
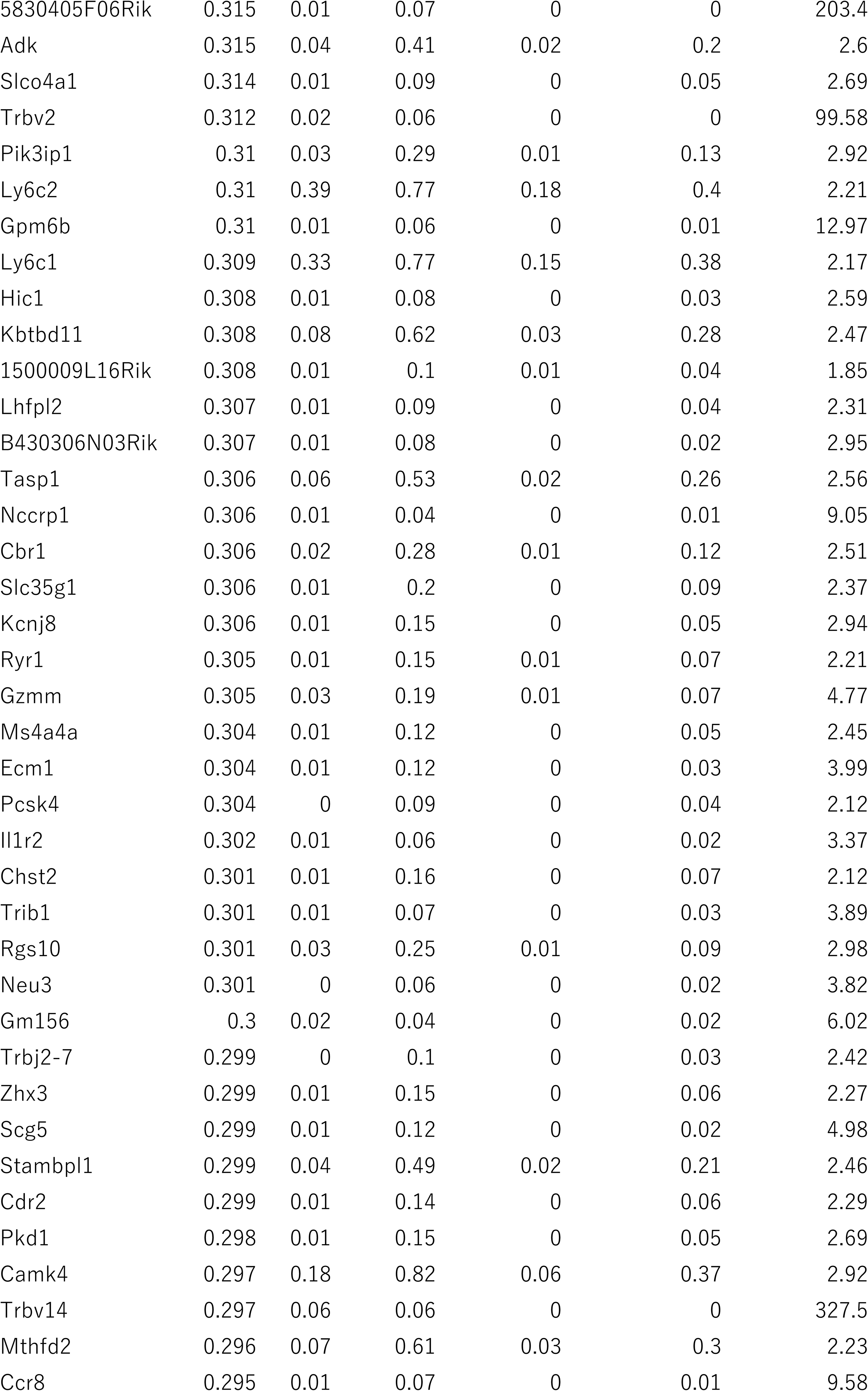
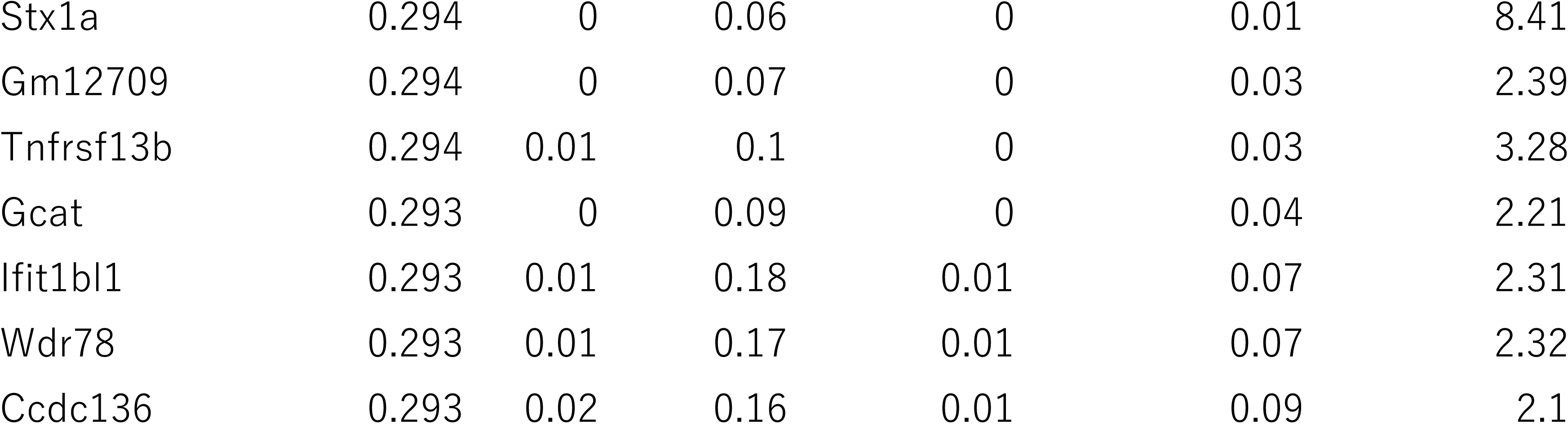
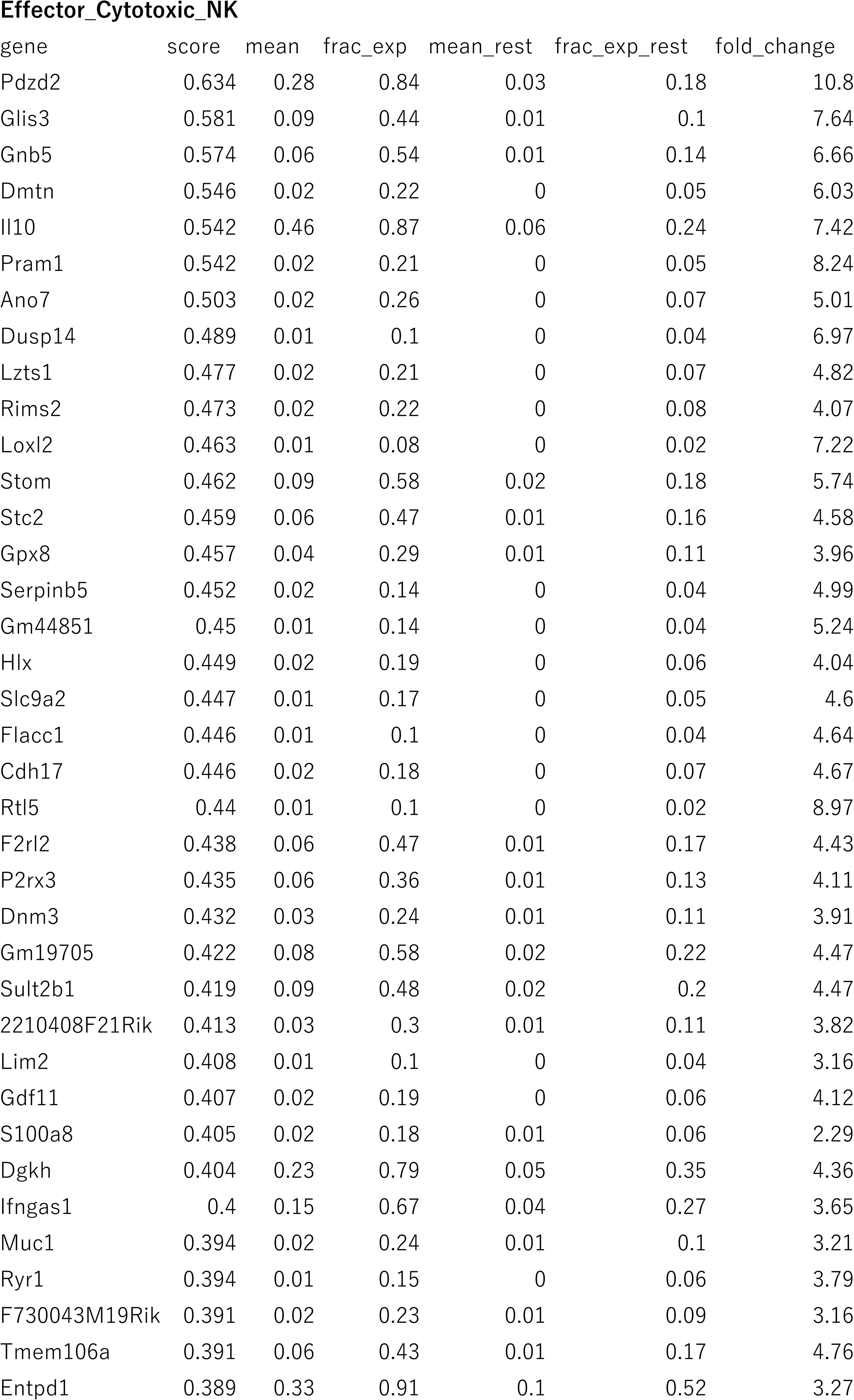
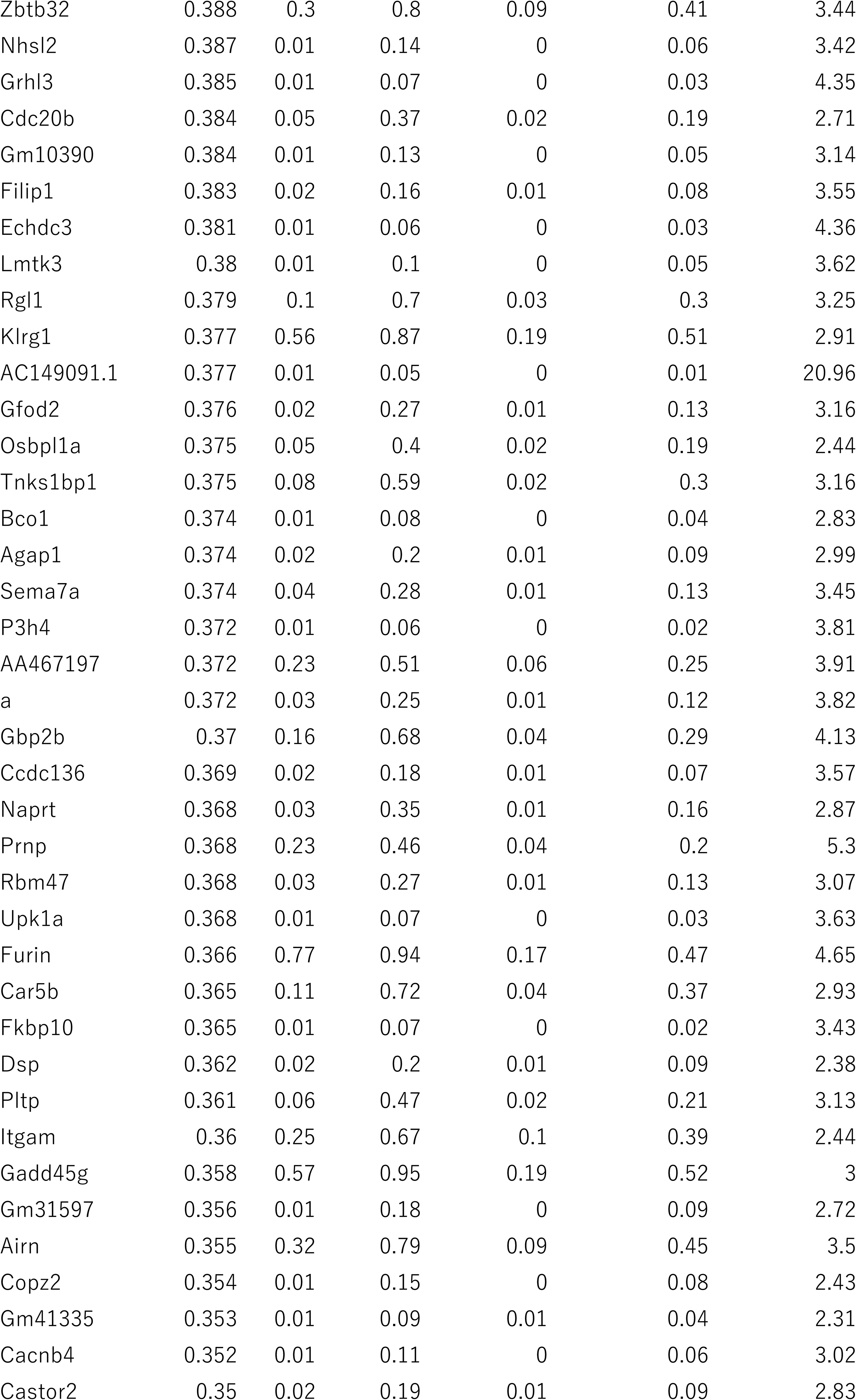
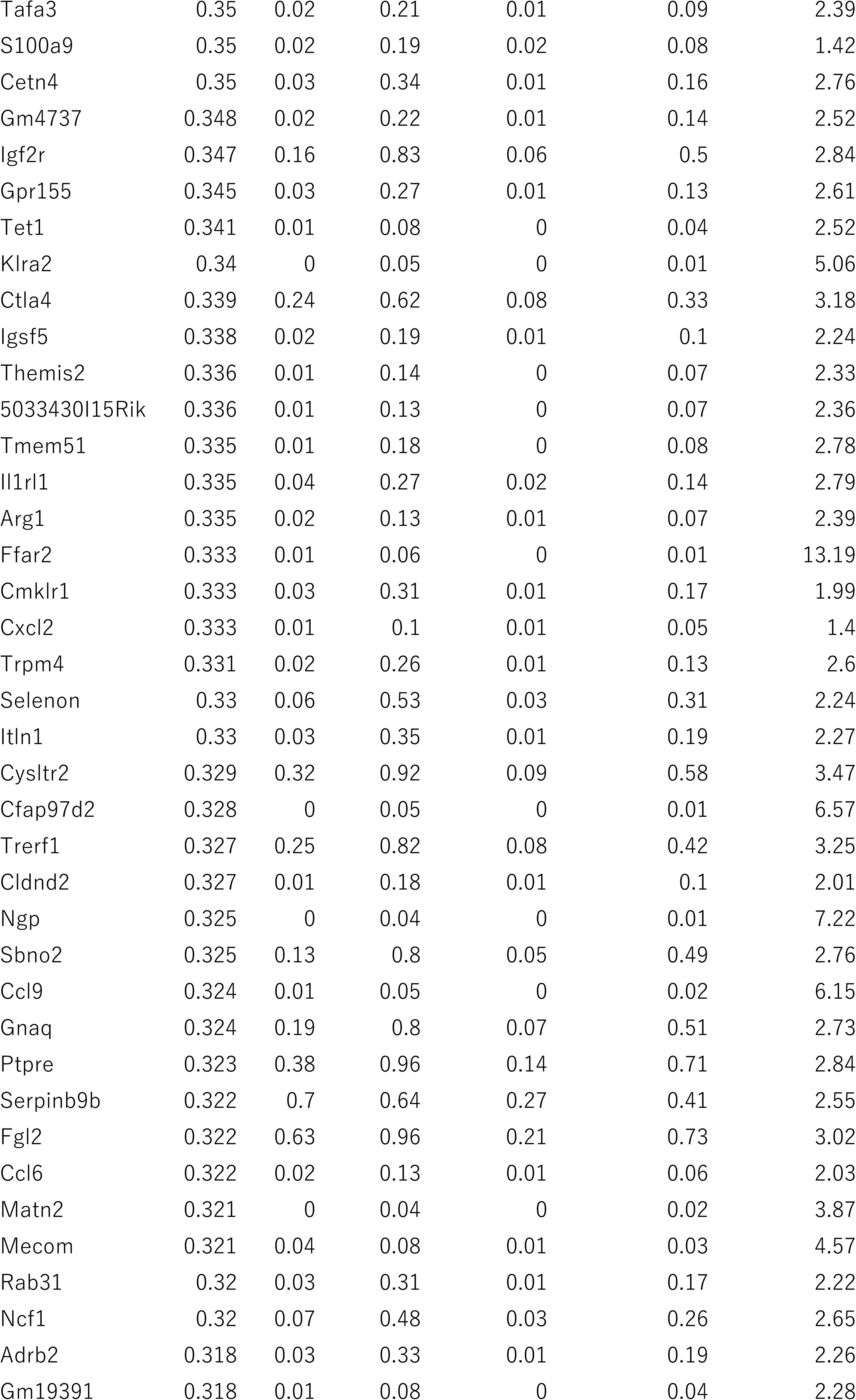
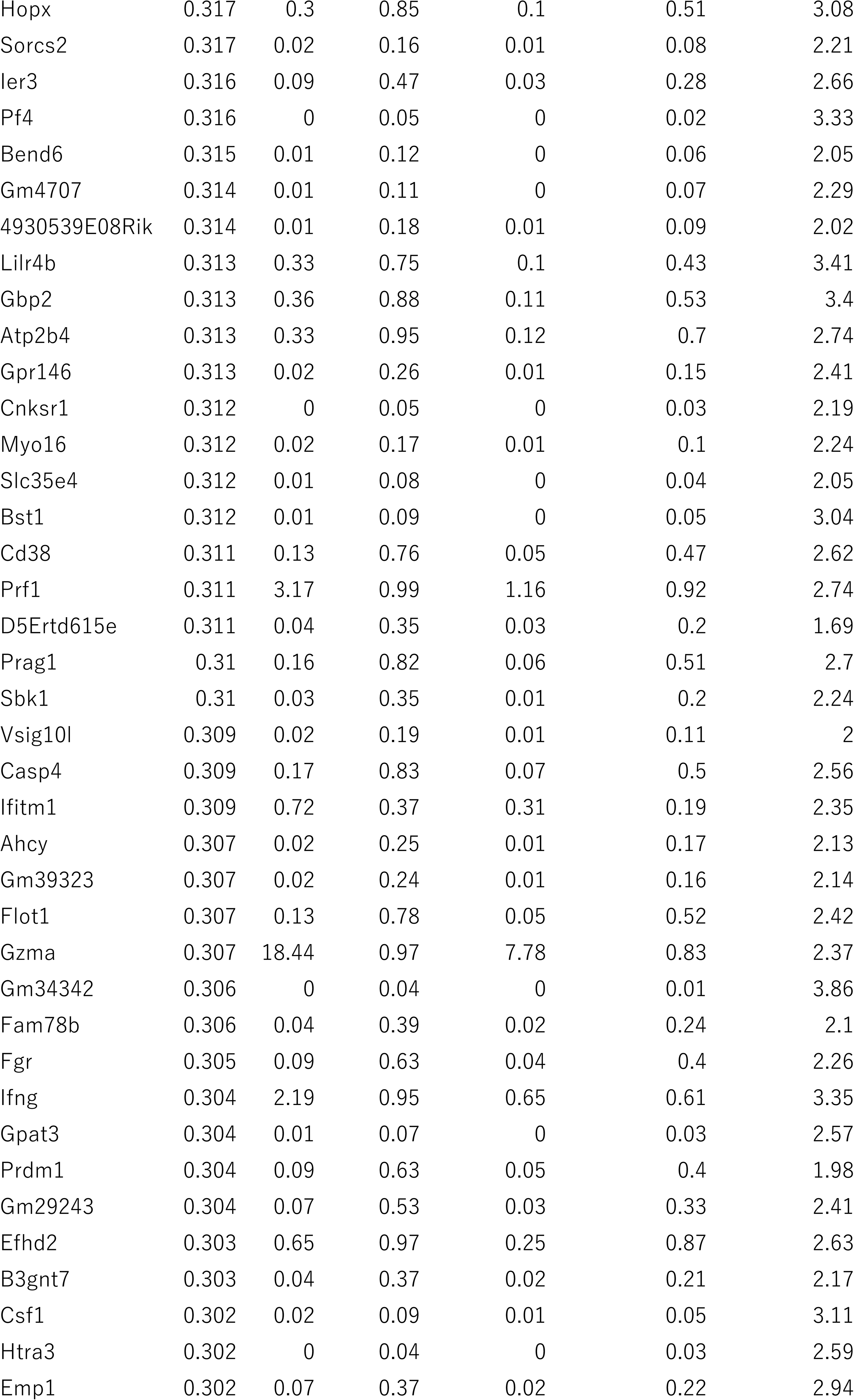
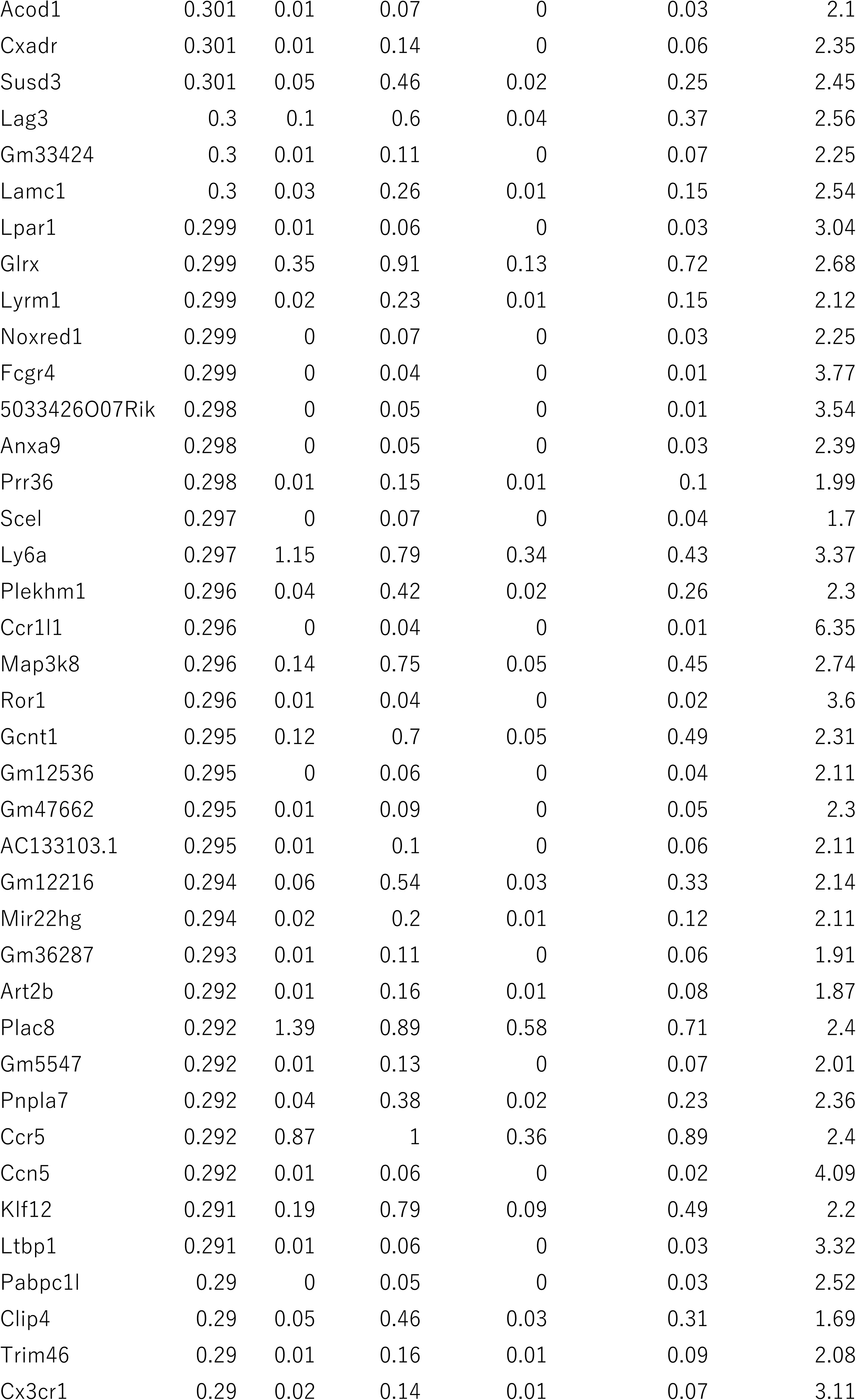
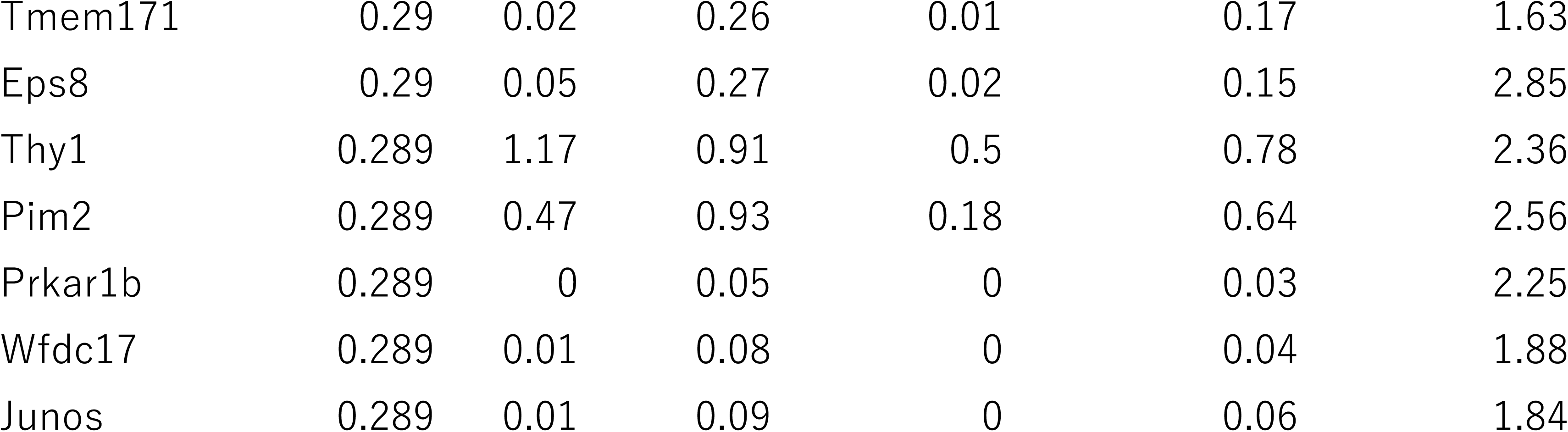
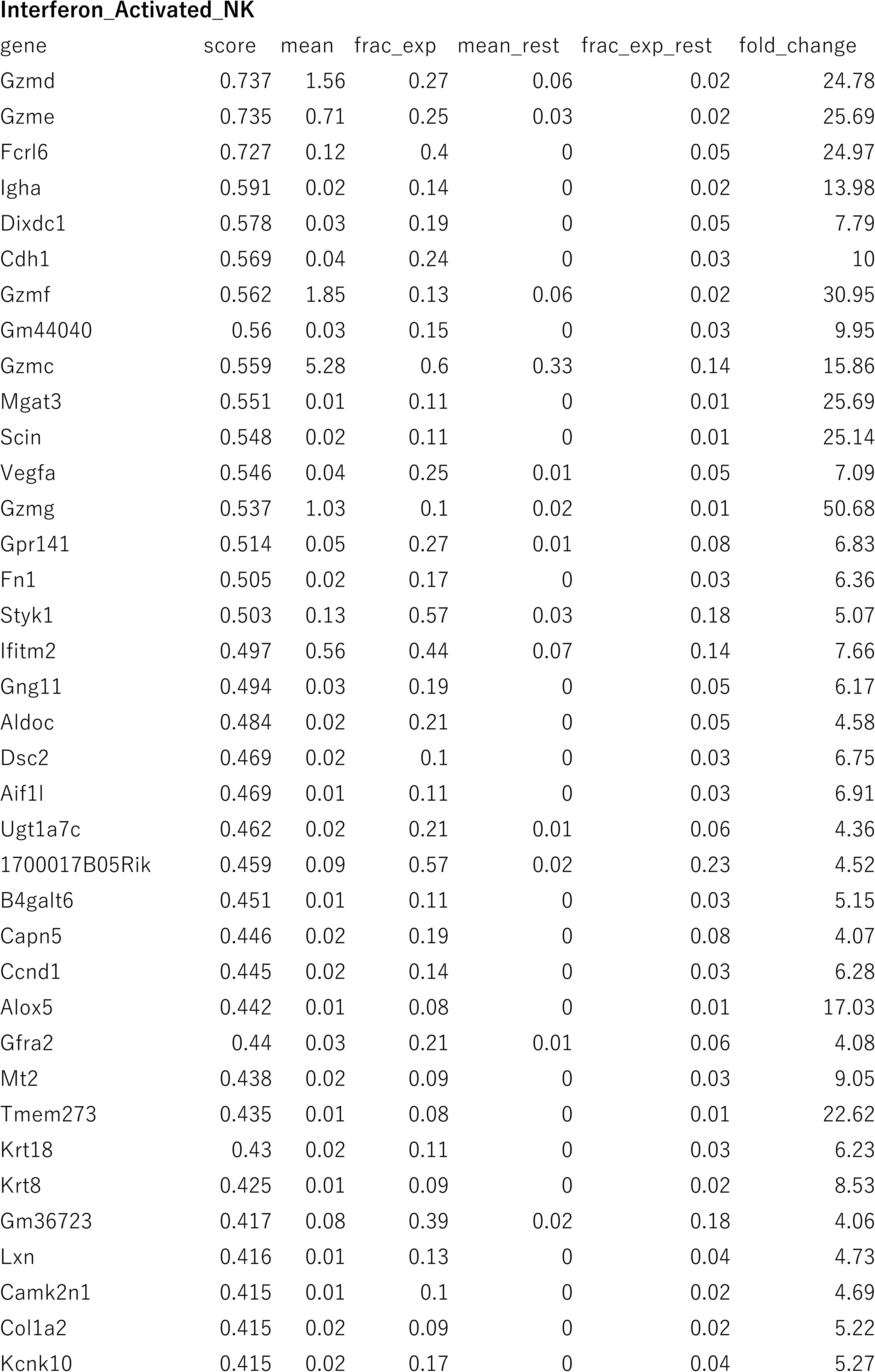
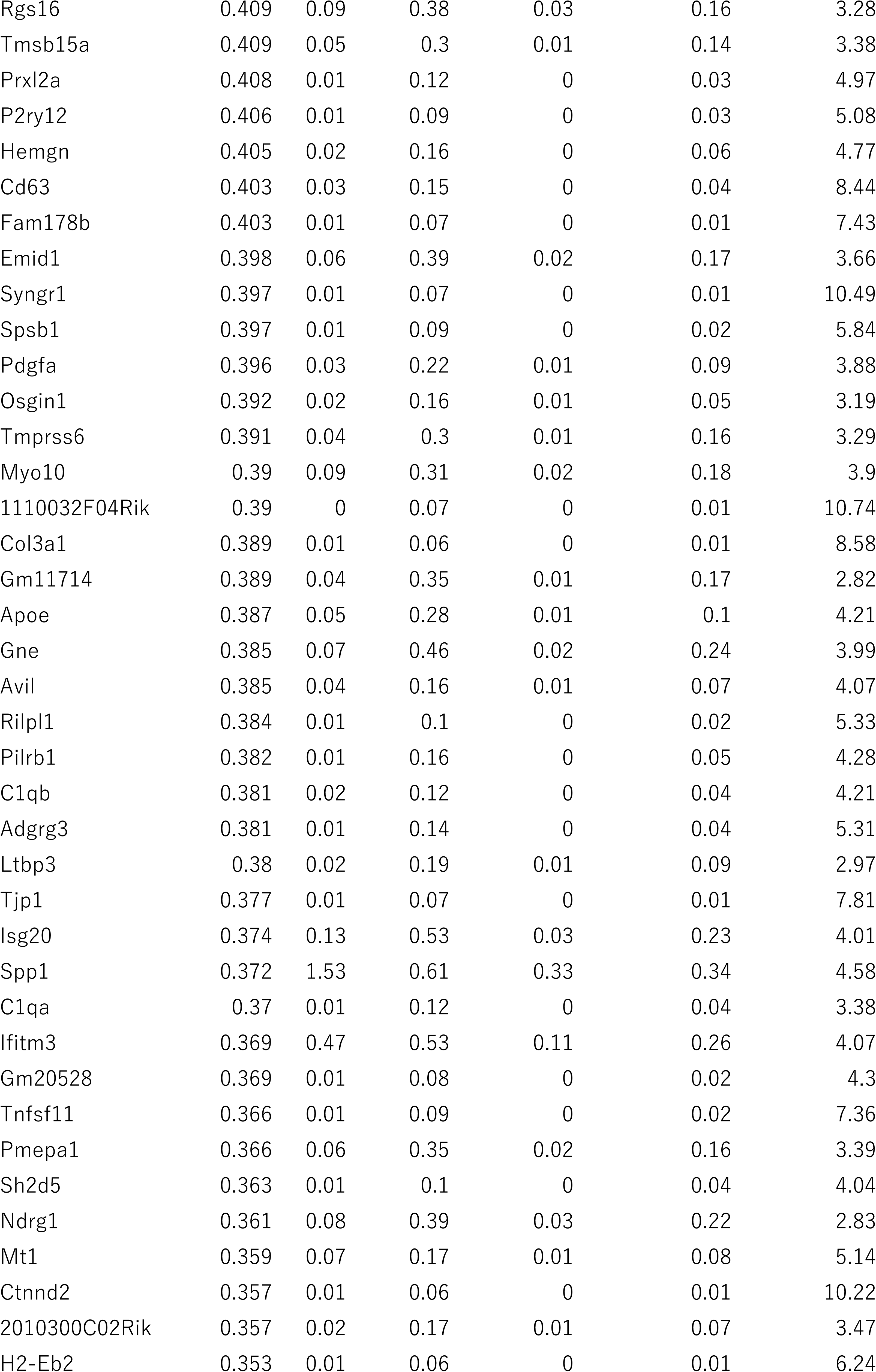
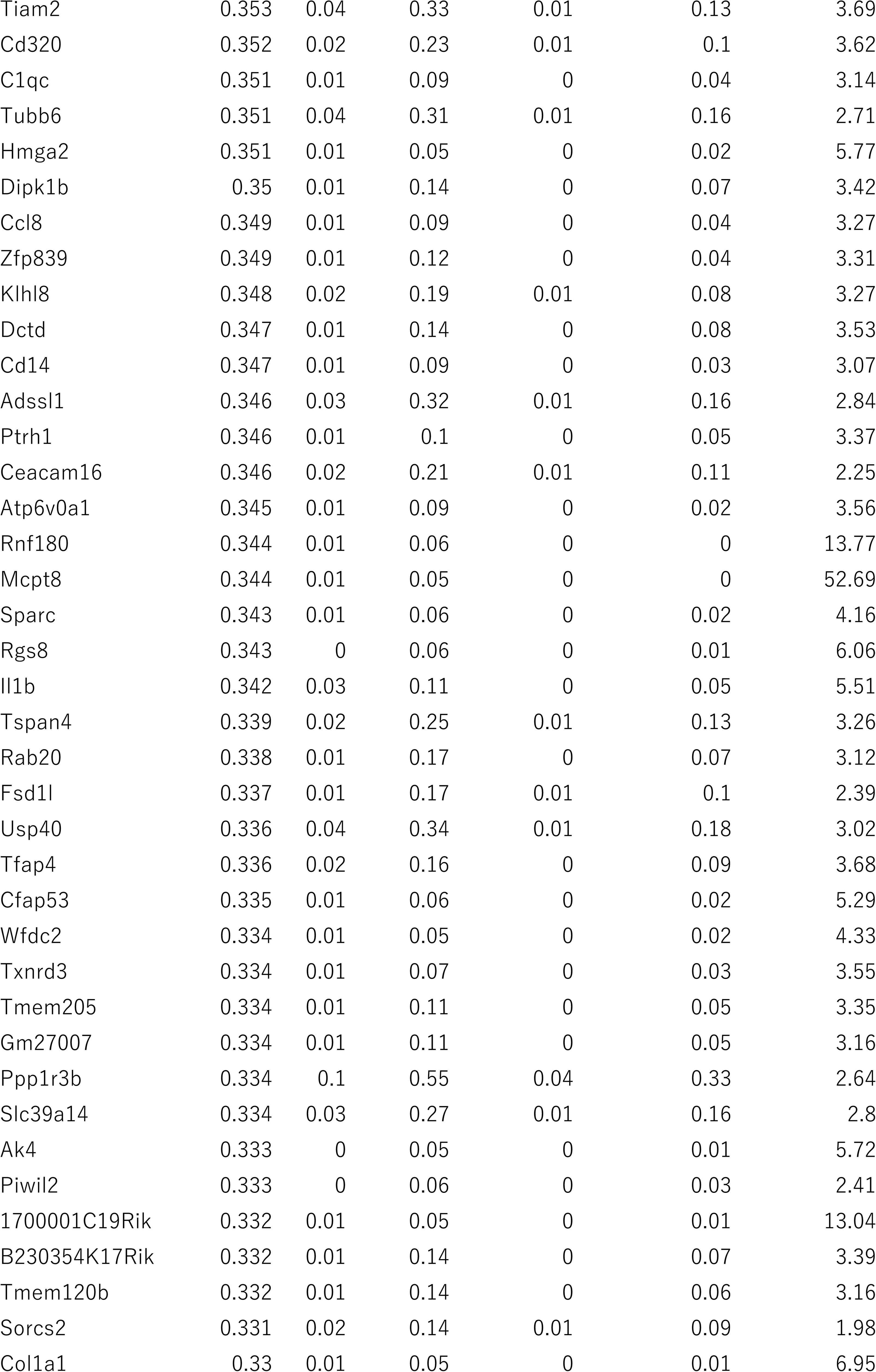
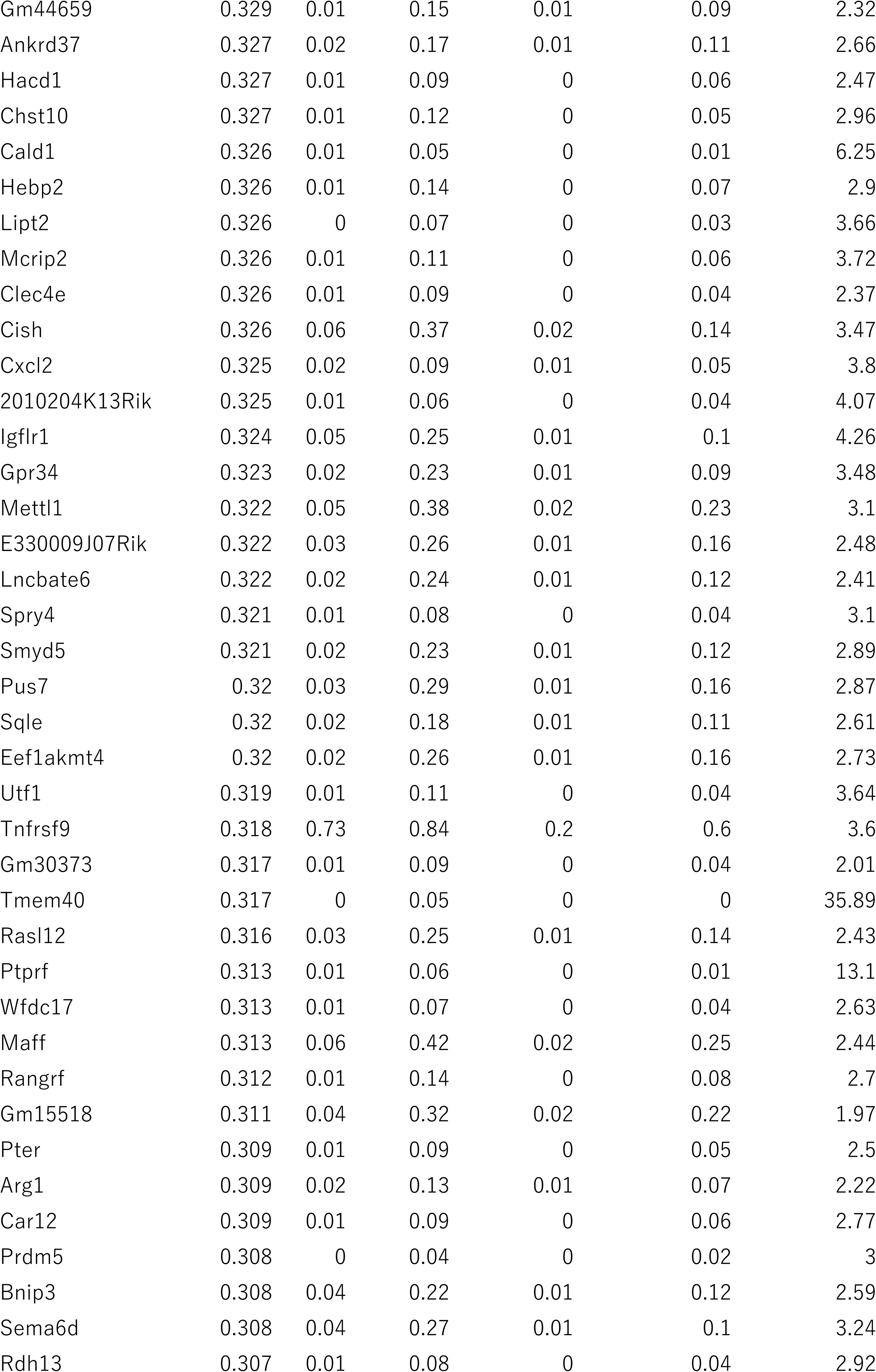
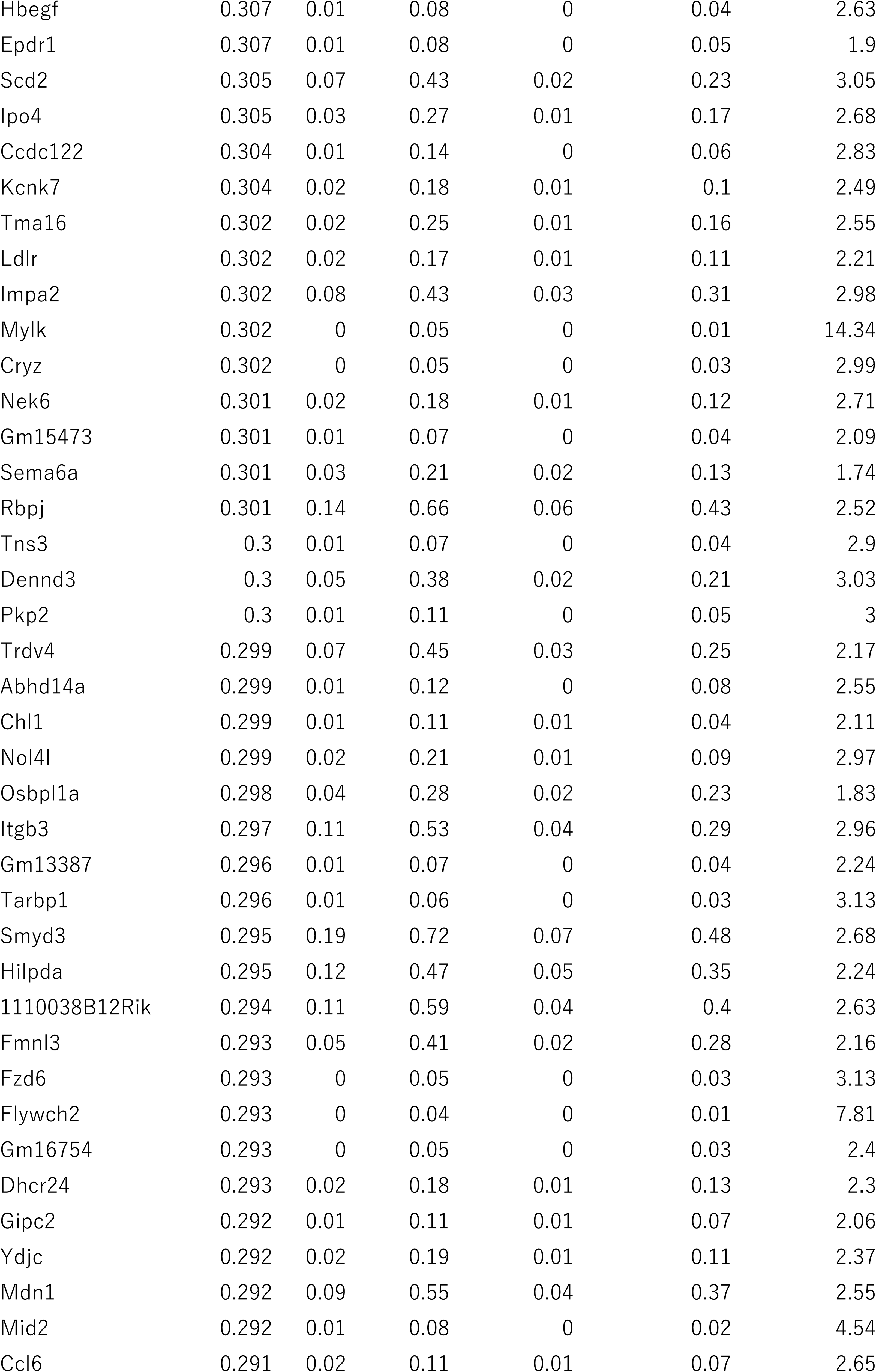
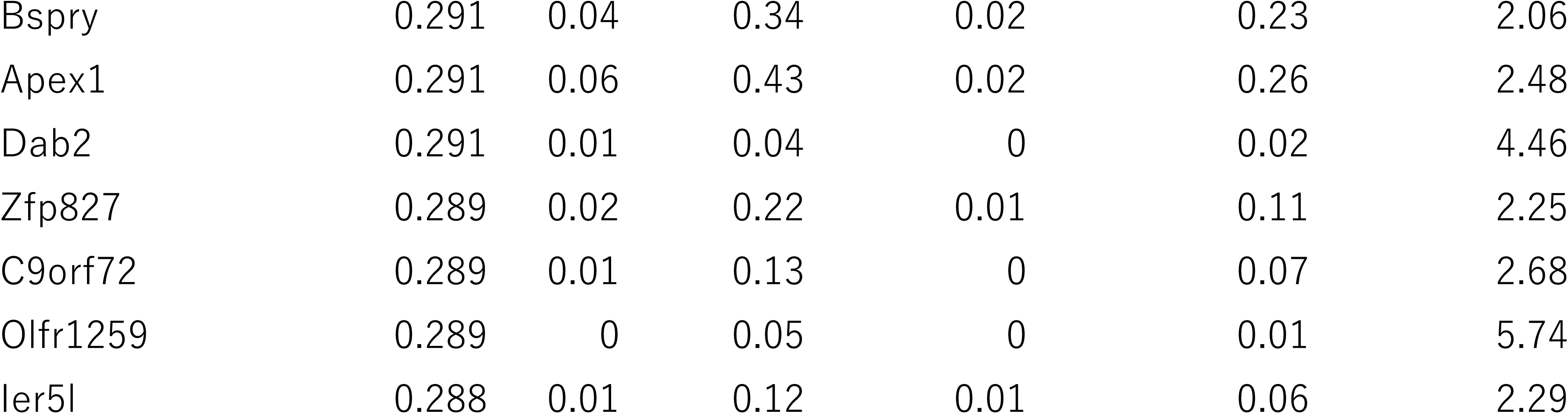
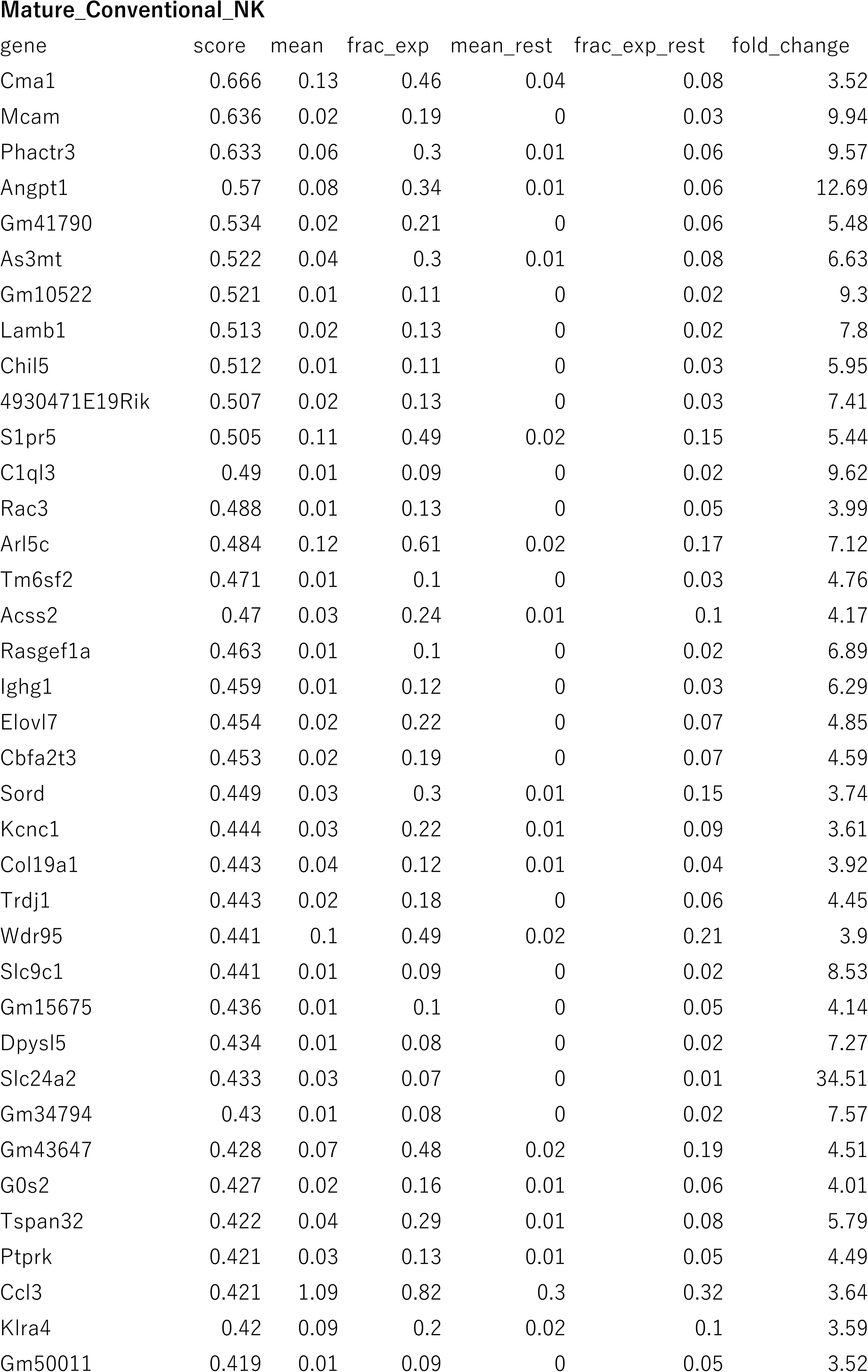
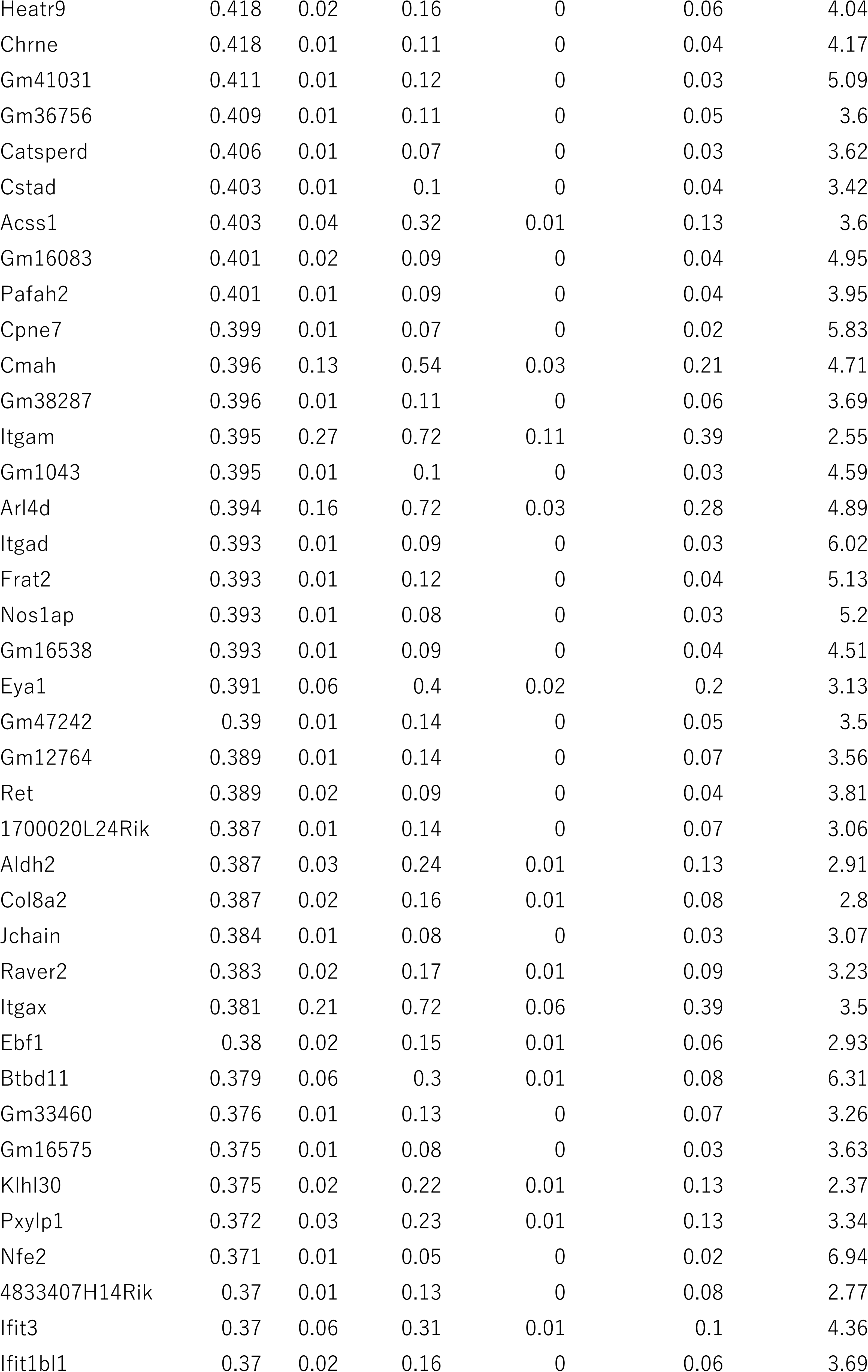
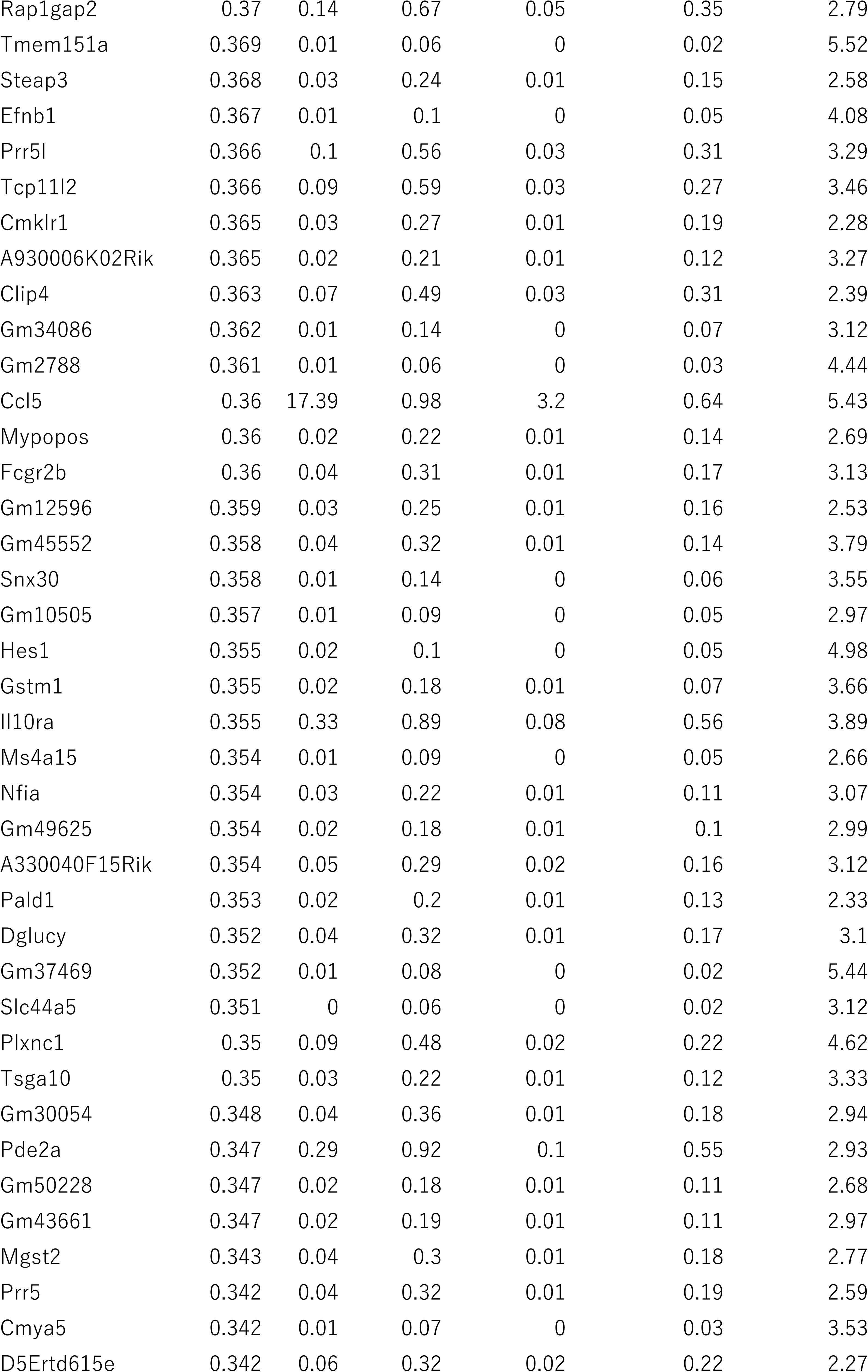
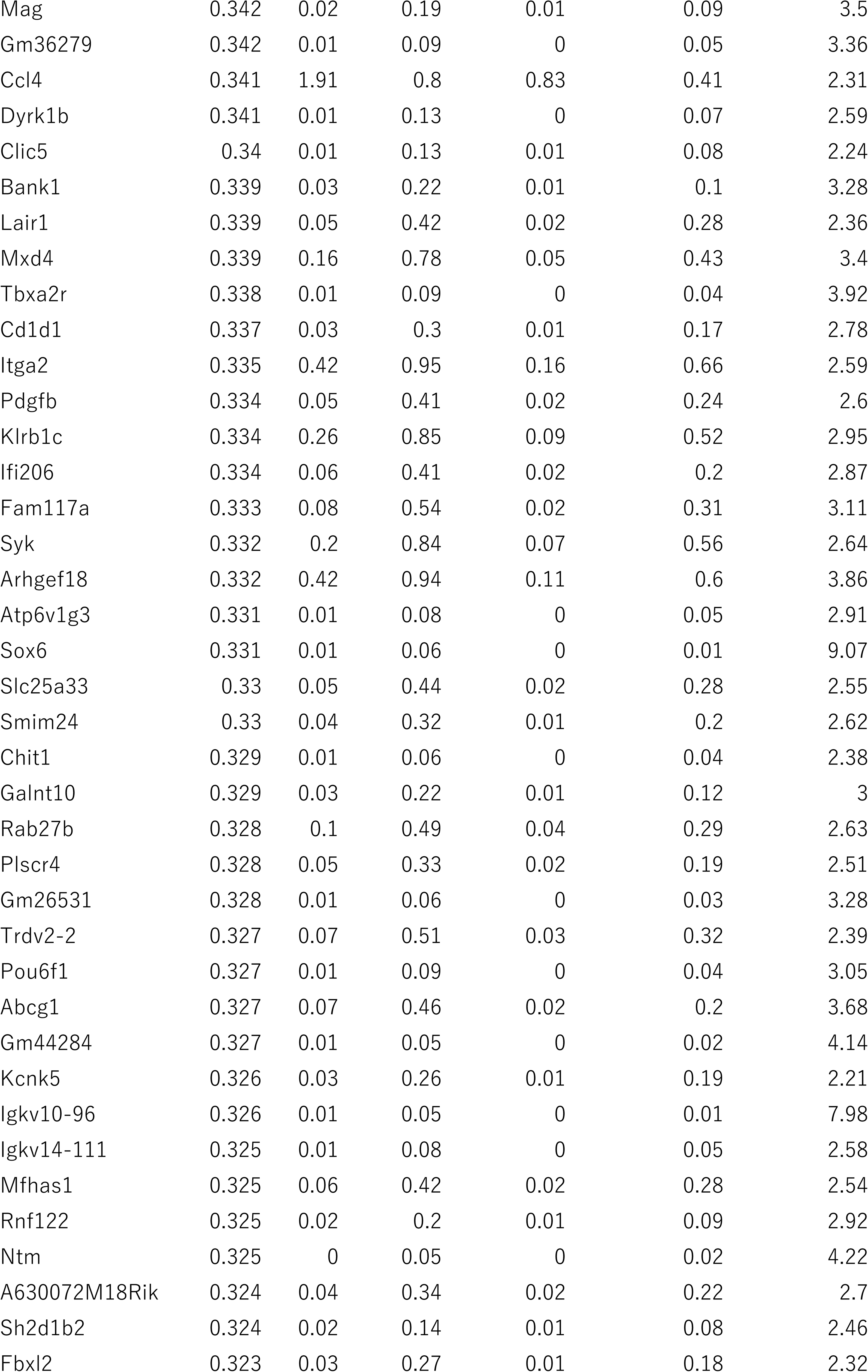
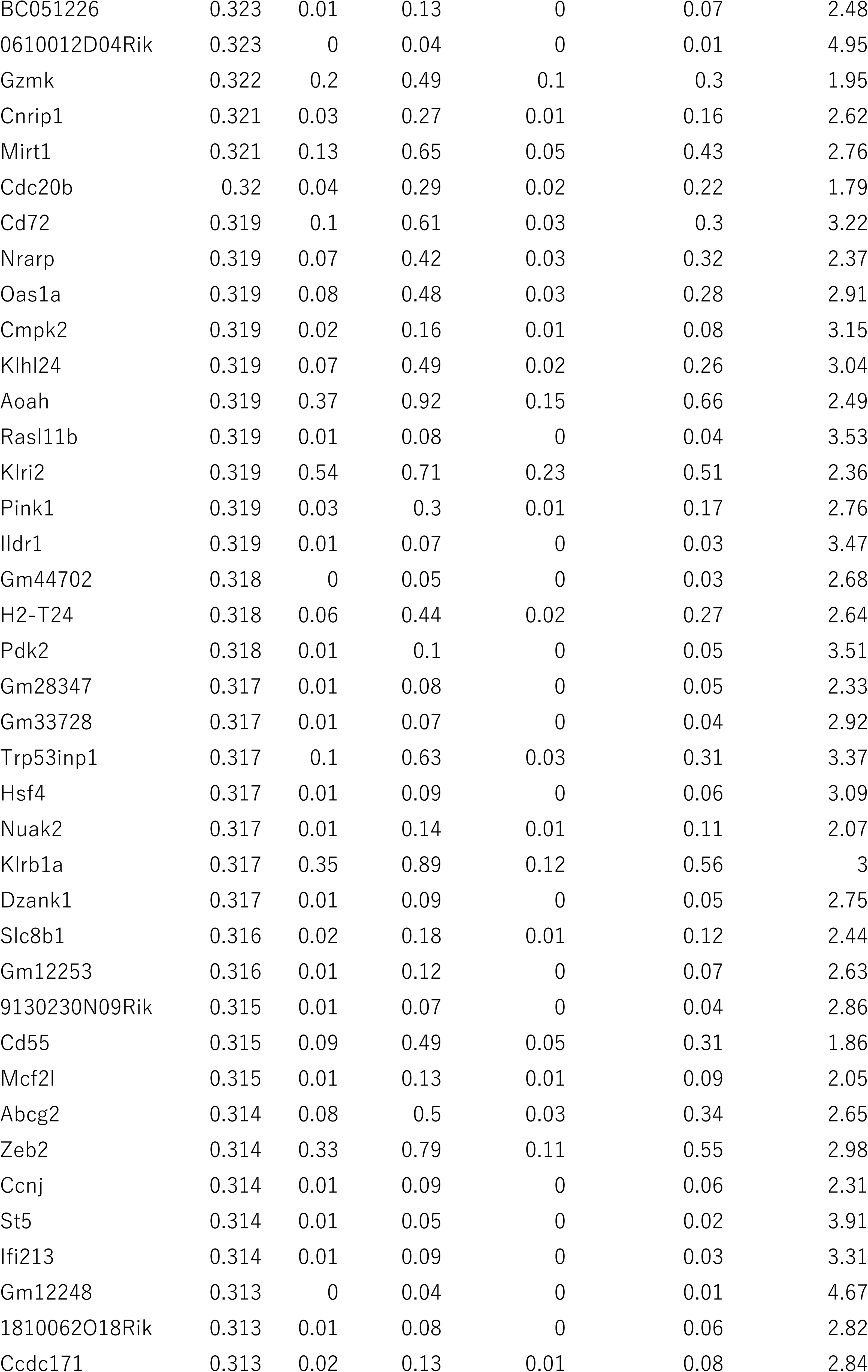
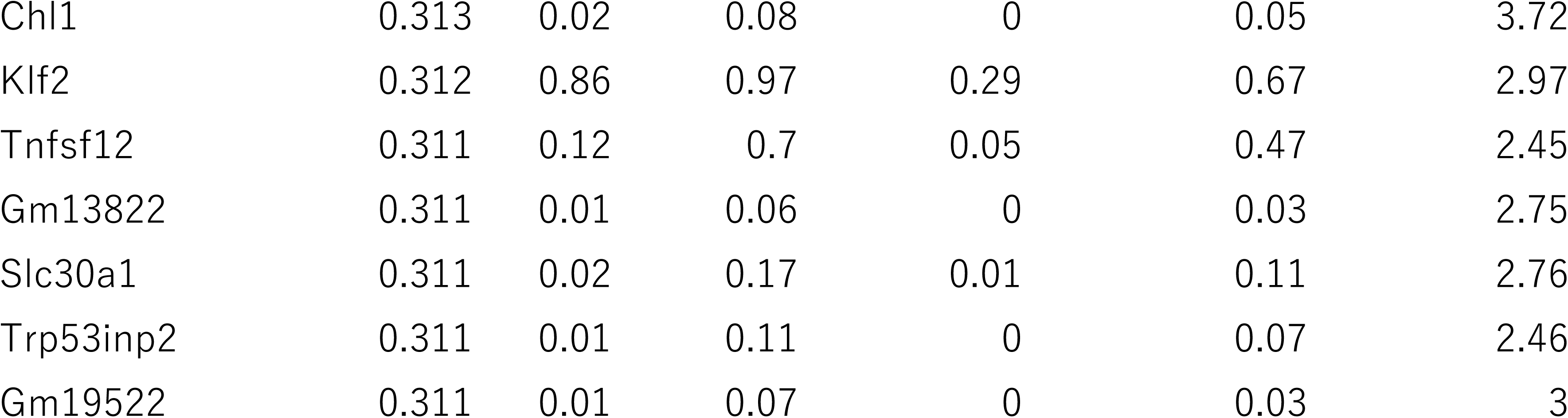
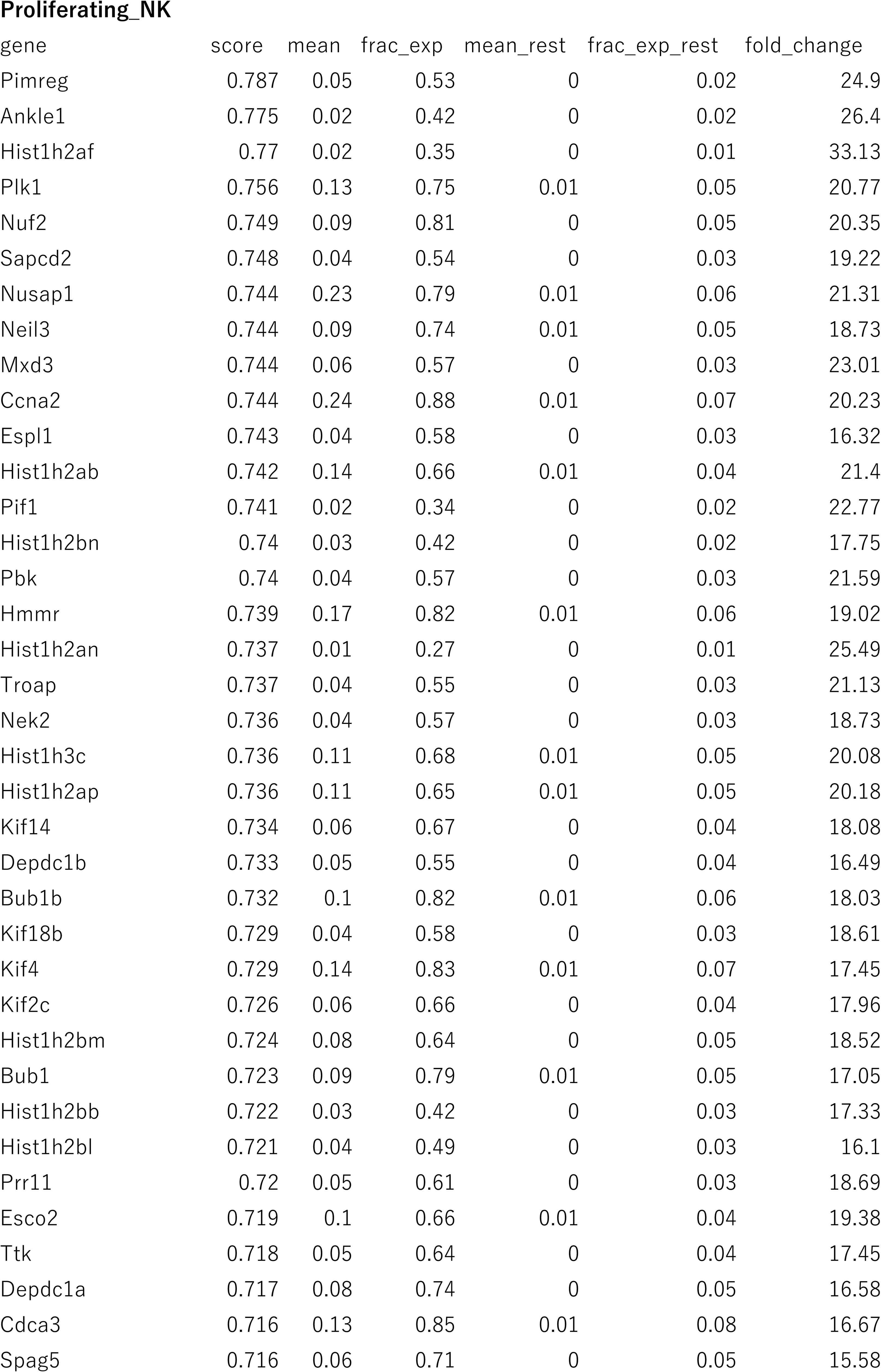
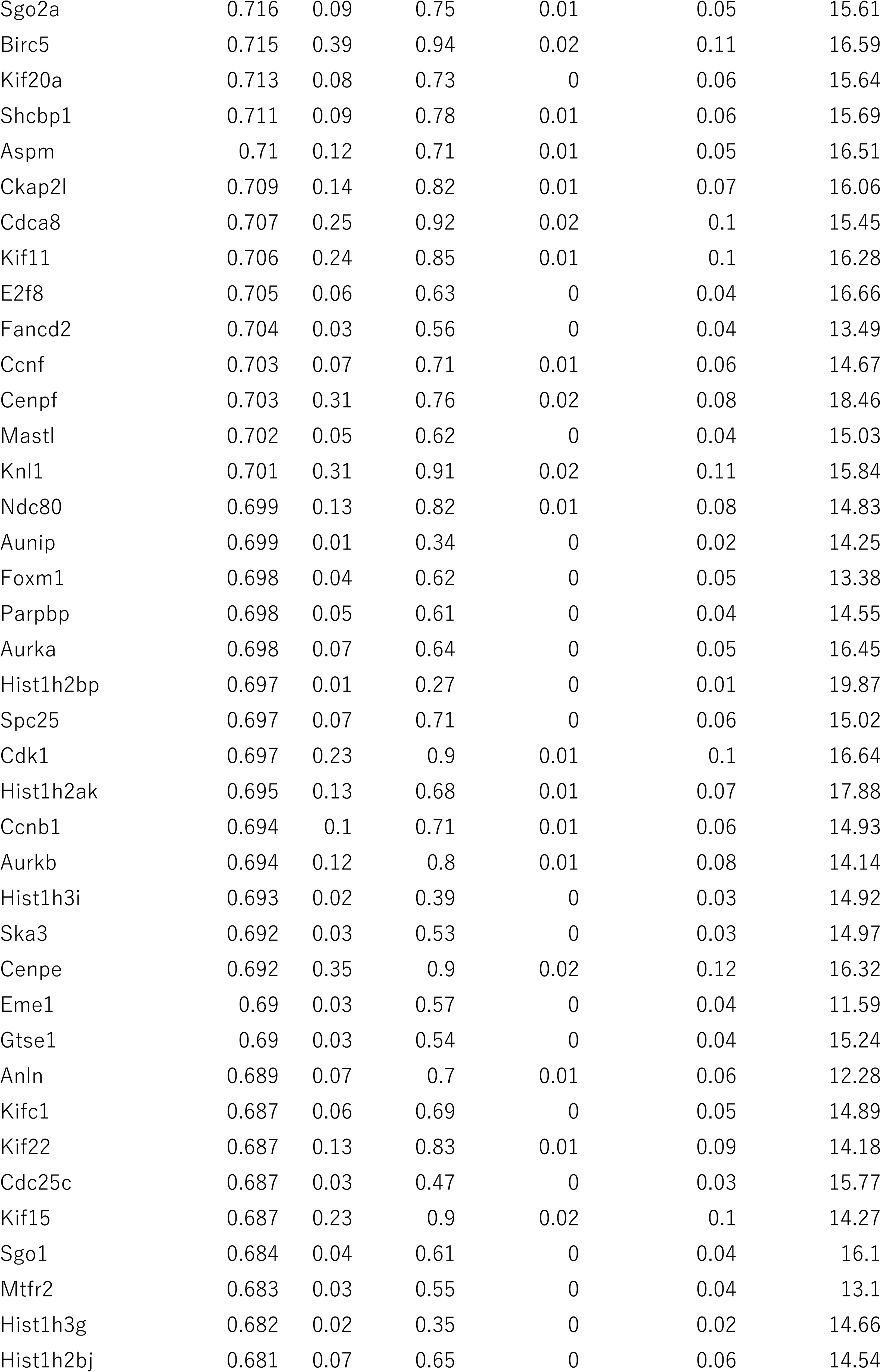
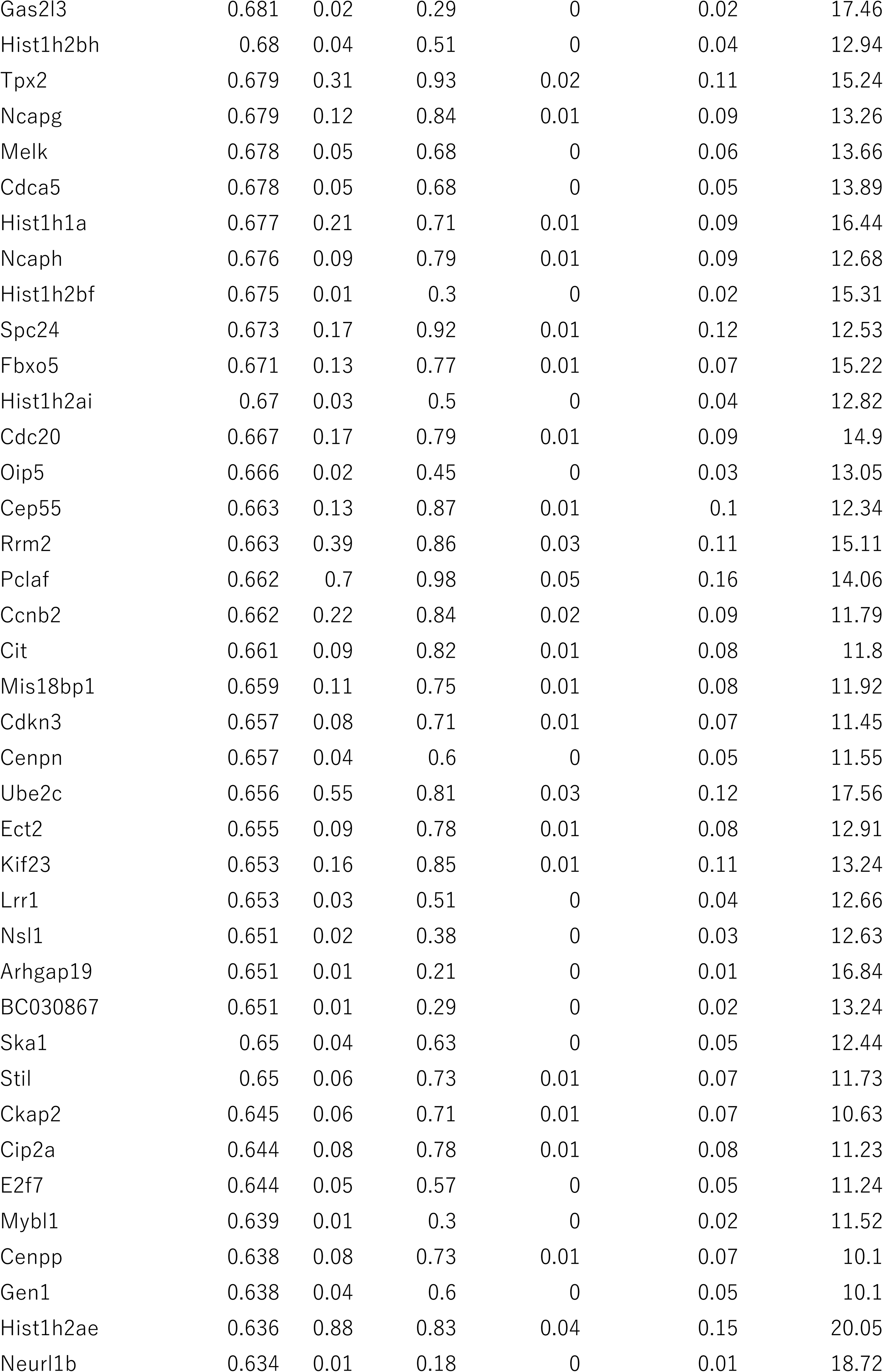
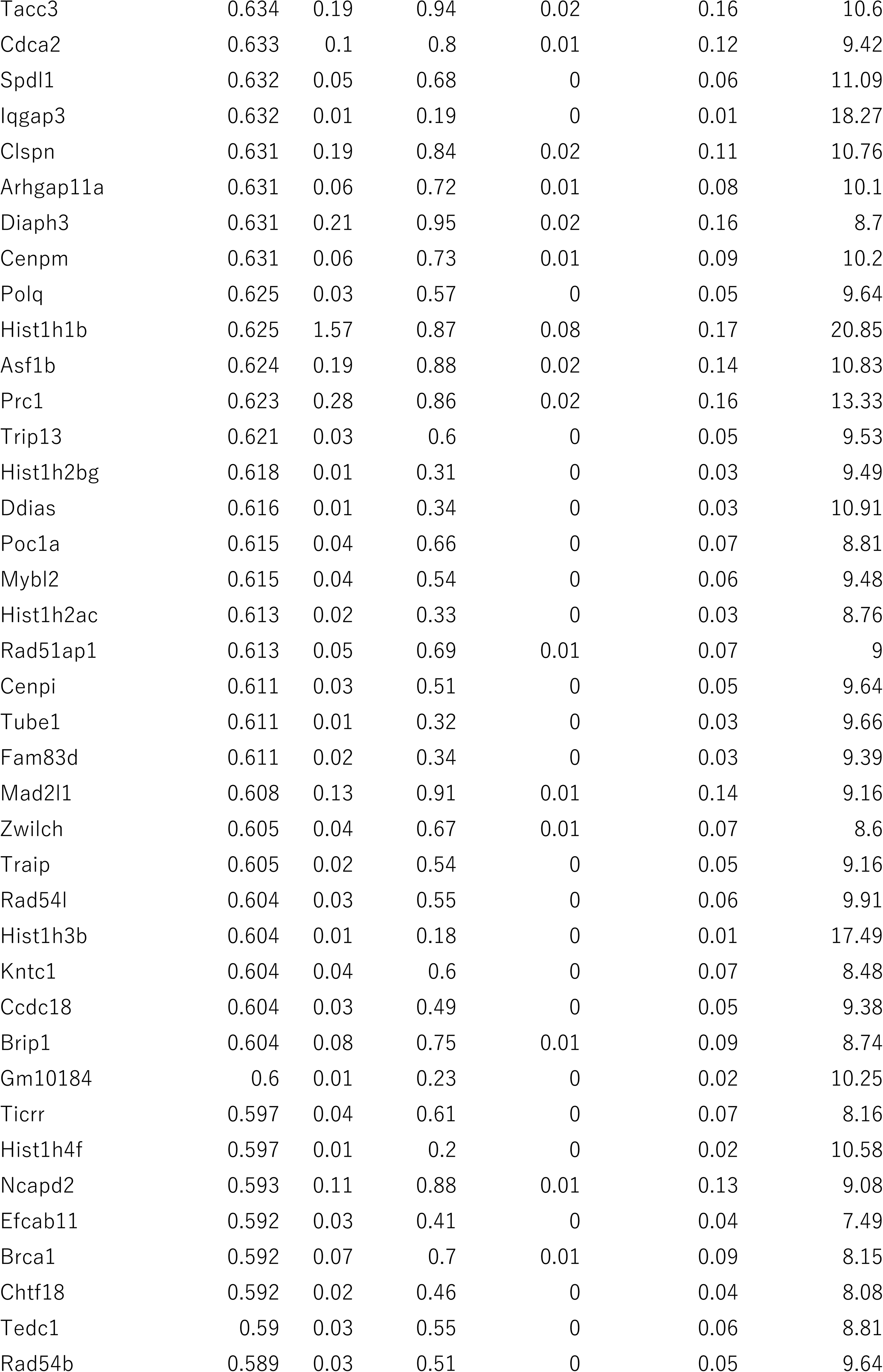
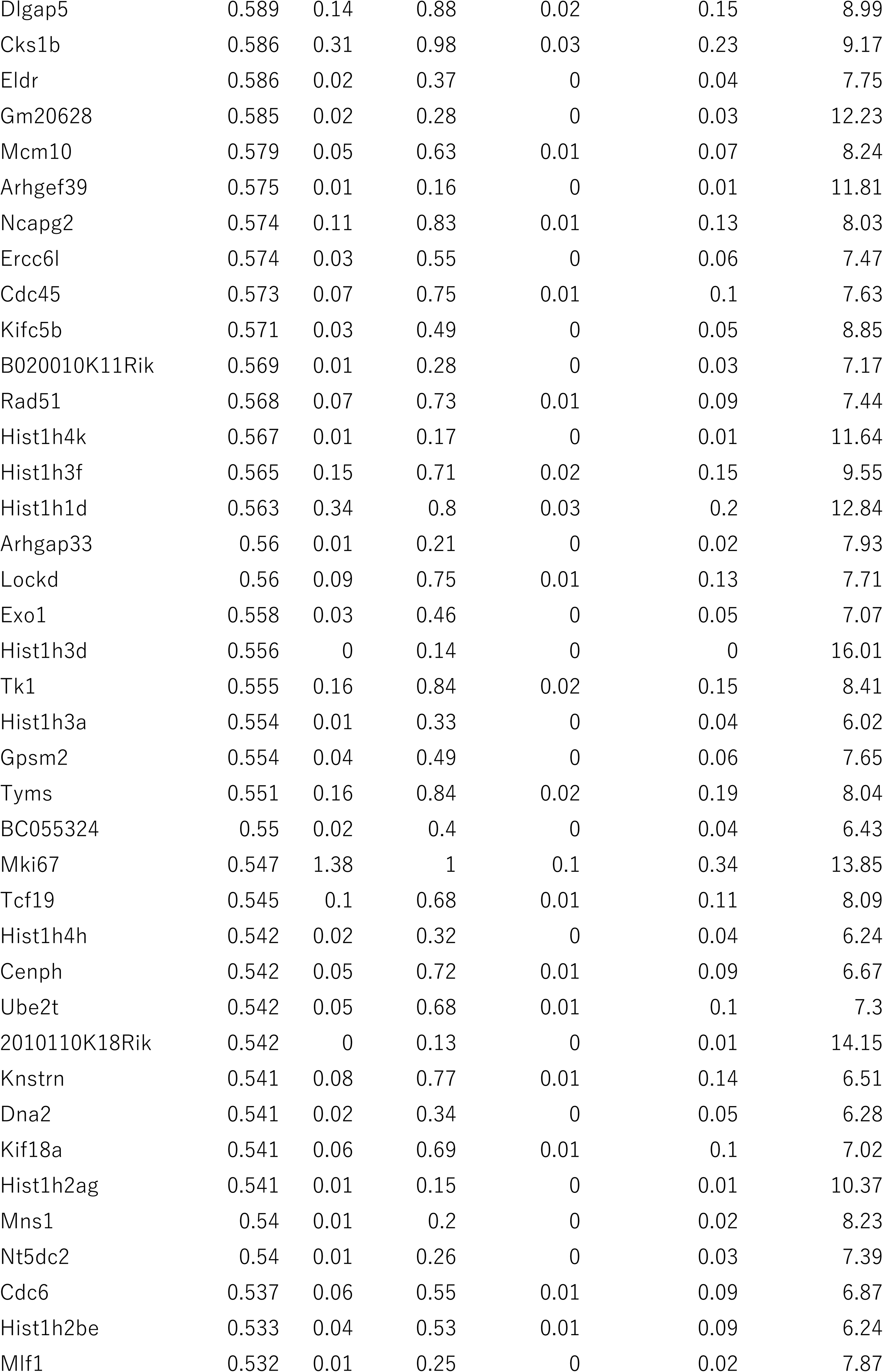
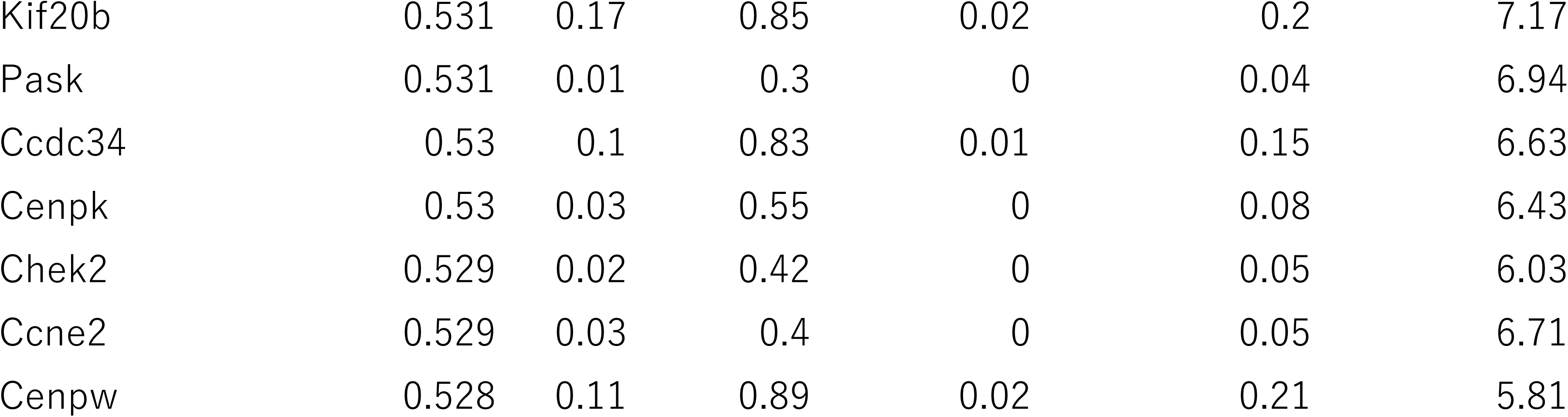
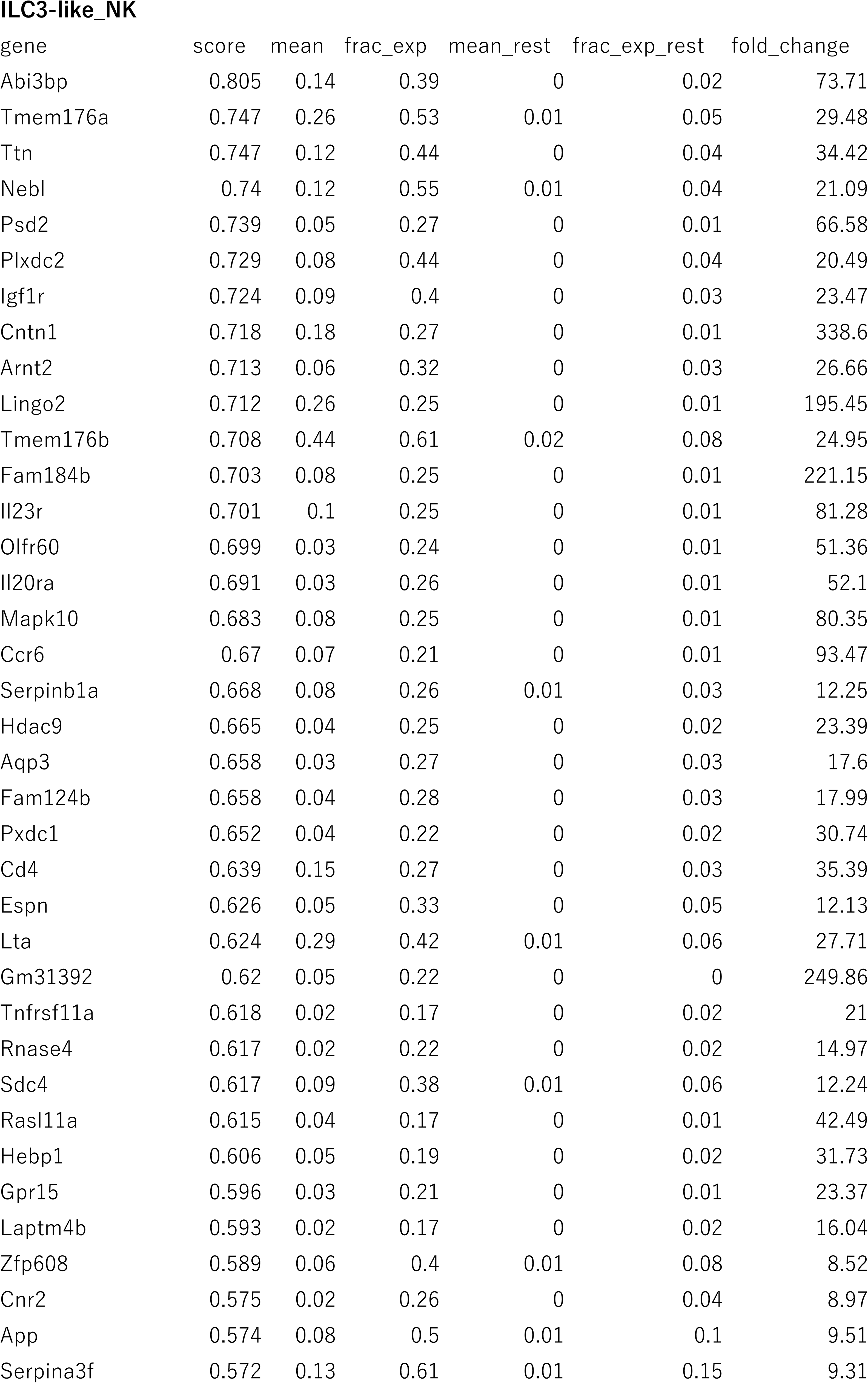
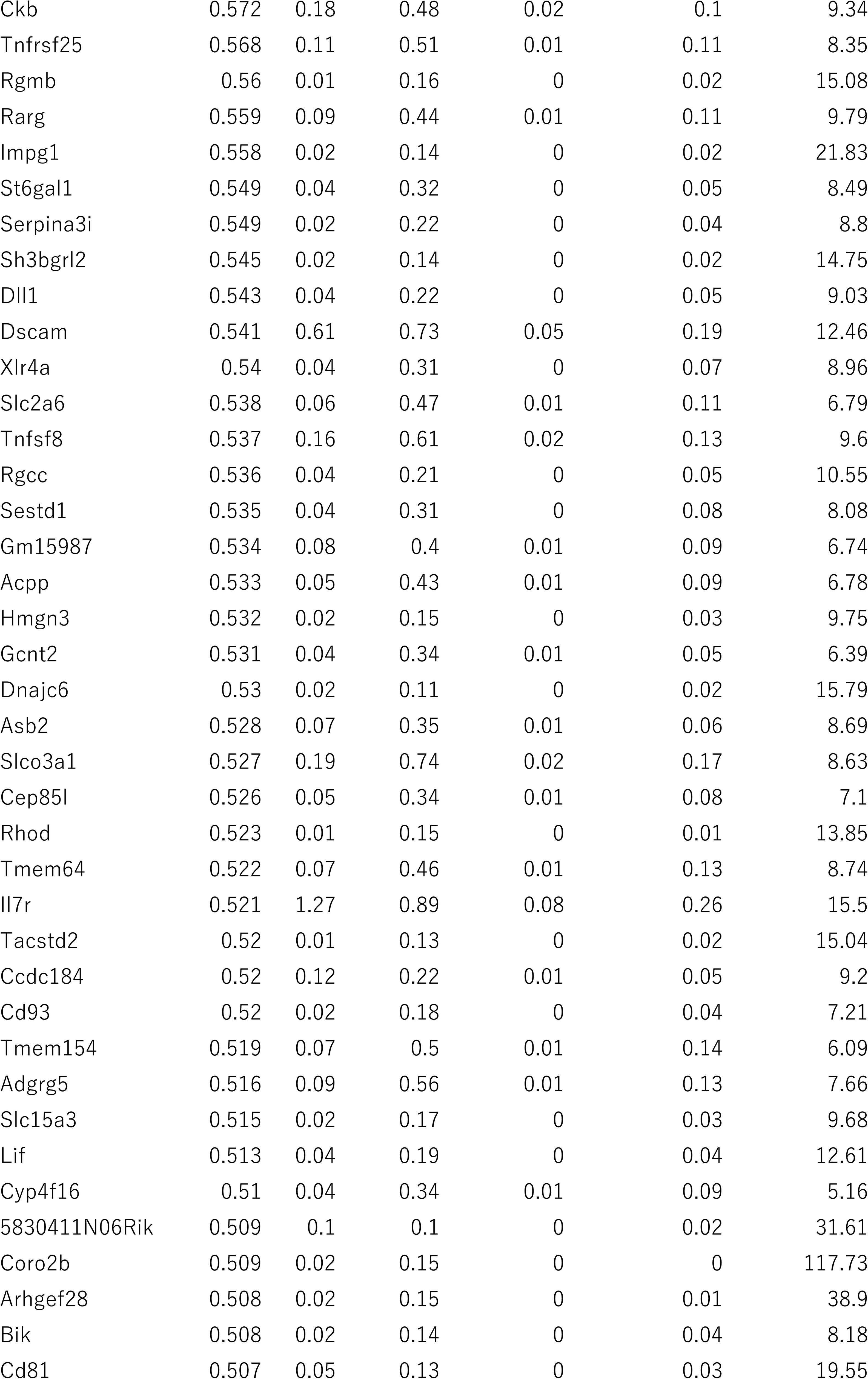
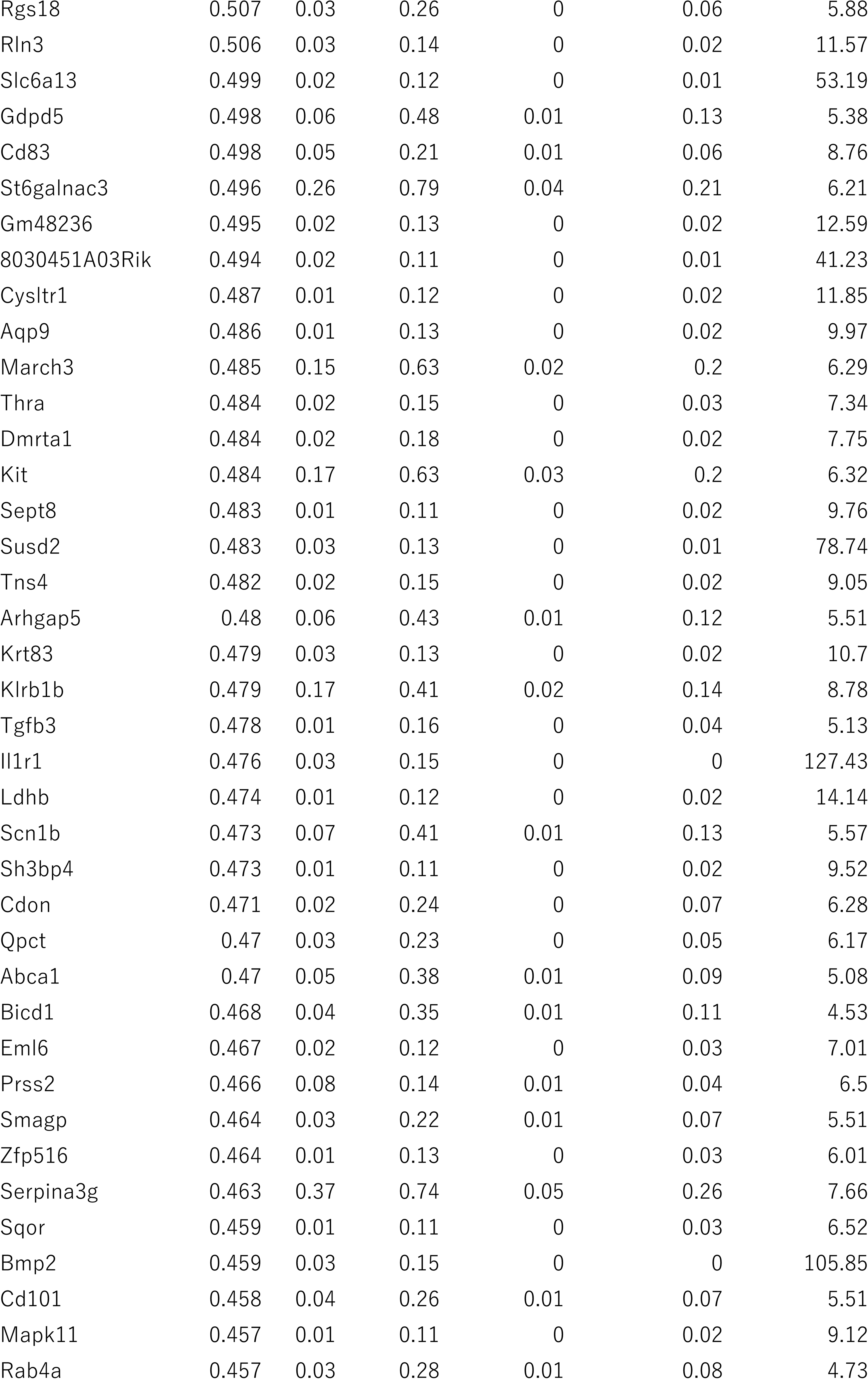
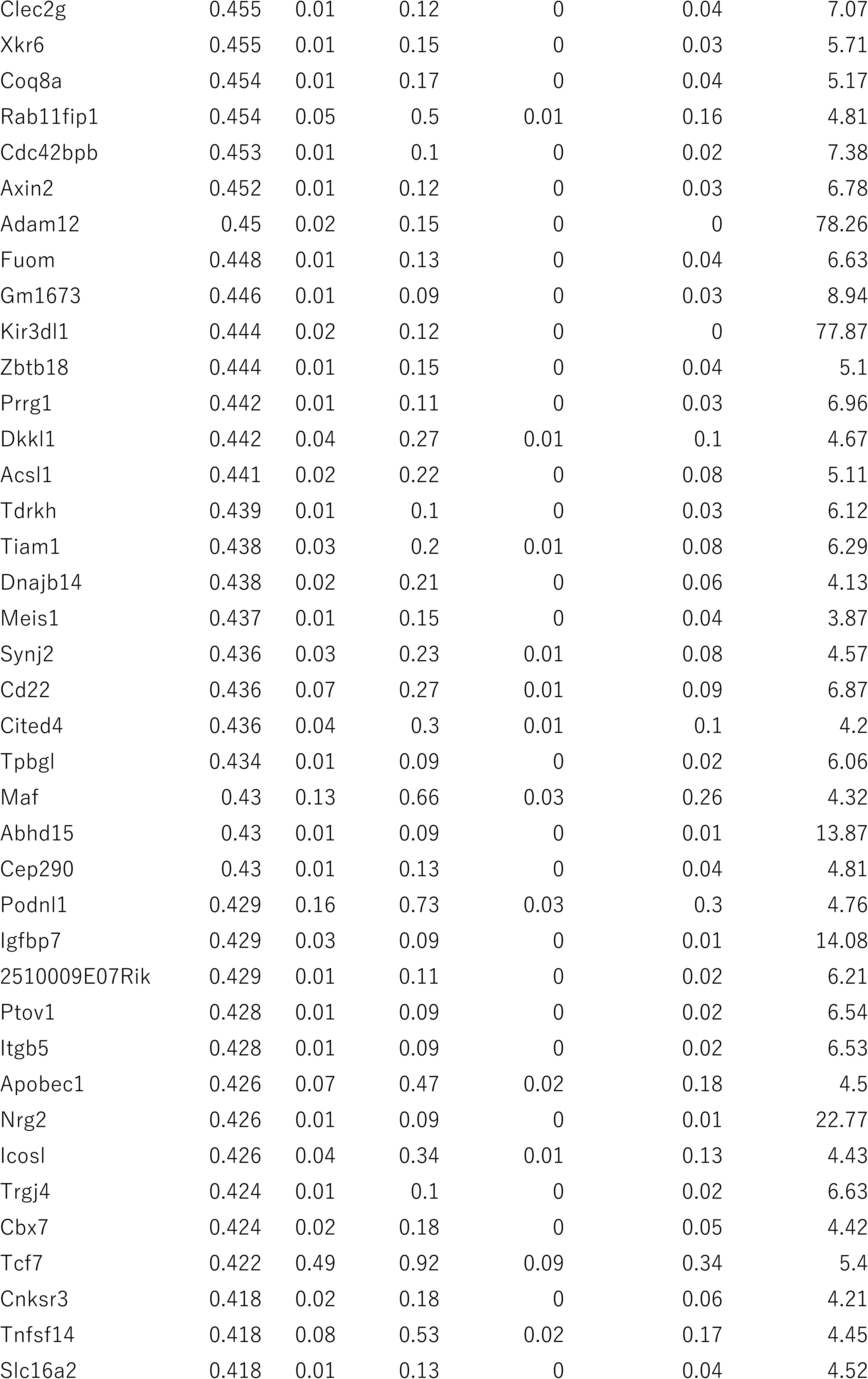
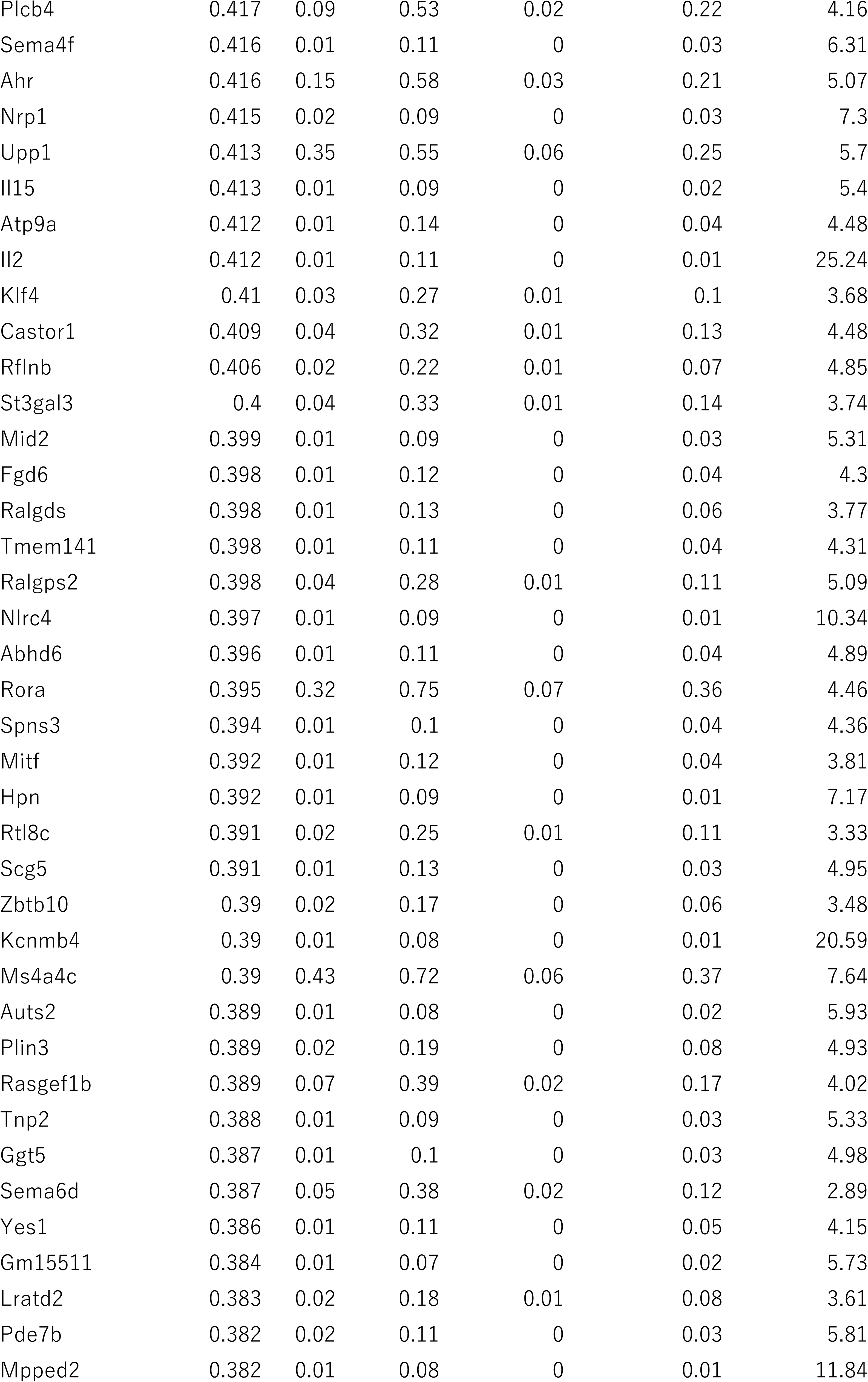
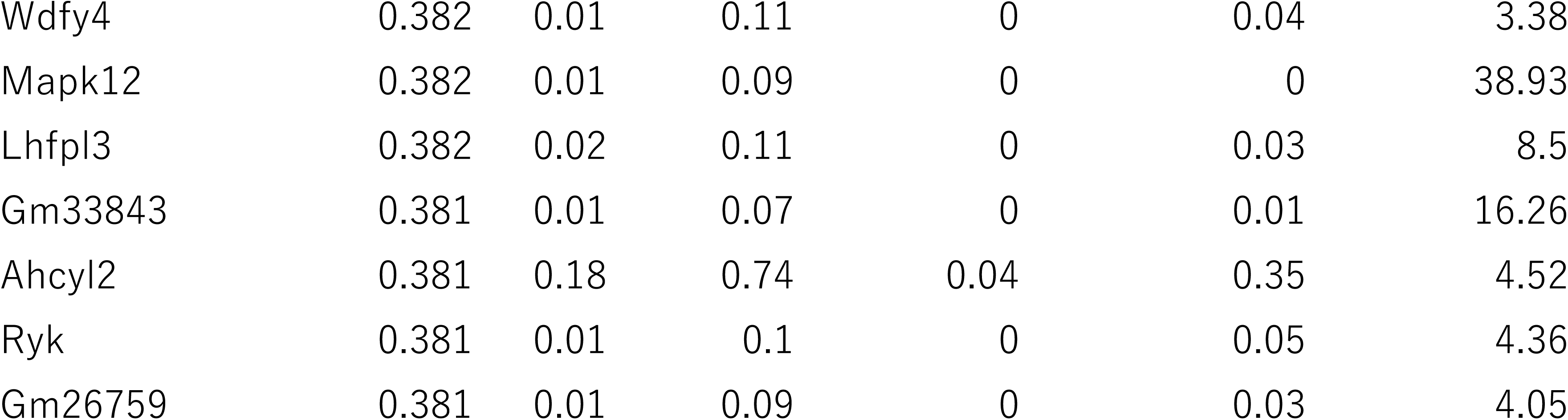
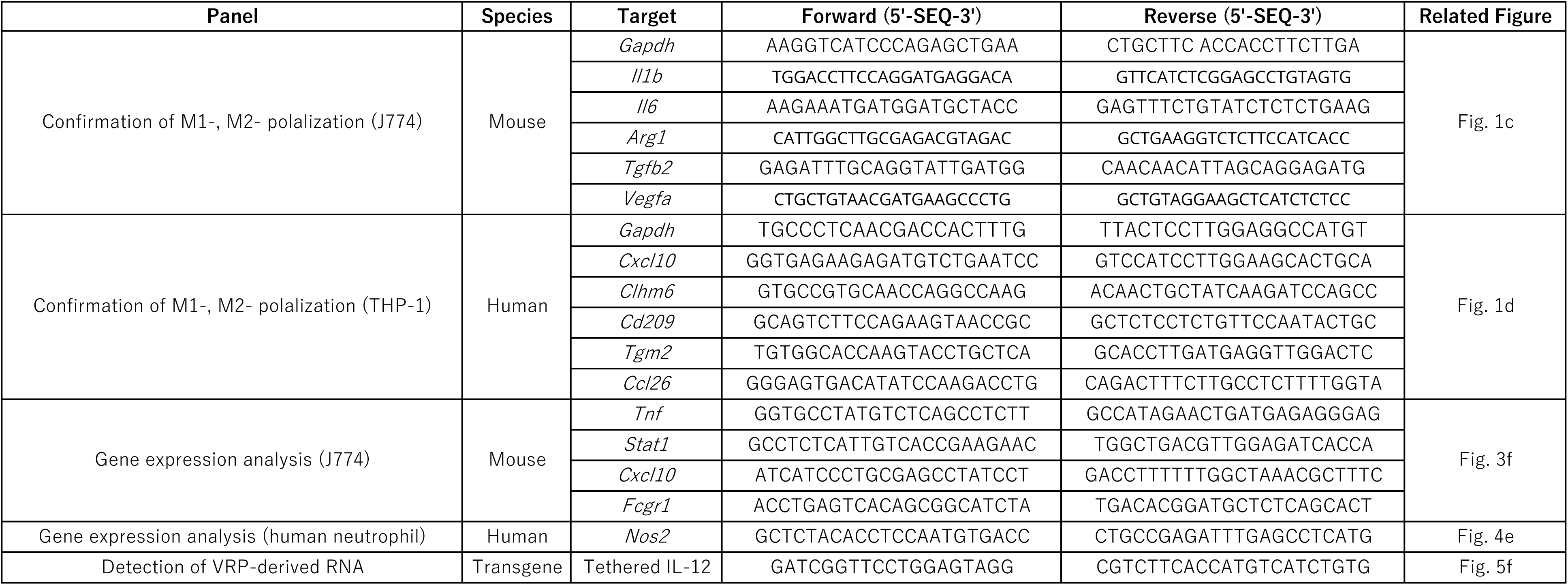

